# The *Draparnaldia* genome: alternative mechanisms for multicellularity and terrestrialization in green plants

**DOI:** 10.1101/2024.09.12.612648

**Authors:** Lenka Caisová, Ewout Crombez, Minerva Susana Trejo Arellano, Marta Gut, Tyler Scott Alioto, Jèssica Gómez-Garrido, Marc Dabad, Anna Esteve-Codina, Ivan Petřík, Aleš Pěnčík, Ondřej Novák, Yves Van de Peer, Beatriz Vicoso, Jiří Friml

**Affiliations:** Institute of Science and Technology Austria, 3400 Klosterneuburg, Austria; Biology Centre, Institute of Plant Molecular Biology, Czech Academy of Sciences, 37005 České Budějovice, Czech Republic; Department of Plant Biotechnology and Bioinformatics, Ghent University, 9052 Ghent, Belgium; Center for Plant Systems Biology, VIB, 9052 Ghent, Belgium; Centro Nacional de Análisis Genómico (CNAG), 08028 Barcelona, Spain; Universitat de Barcelona (UB), 08028 Barcelona, Spain; Laboratory of Growth Regulators, Institute of Experimental Botany, The Czech Academy of Sciences & Faculty of Science, Palacký University, 78371Olomouc, Czech Republic; Department of Biochemistry, Genetics and Microbiology, University of Pretoria, 0028 Pretoria, South-Africa; College of Horticulture, Academy for Advanced Interdisciplinary Studies, Nanjing Agricultural University, 210095 Nanjing, China

**Keywords:** *Draparnaldia*, green algae, terrestrialization, multicellularity, plant evolution, phytohormones, genome, model organism

## Abstract

Green plants contain two algal lineages: Streptophyte algae that diverged into land plants and Chlorophyte algae that are mostly aquatic. *Draparnaldia* is a Chlorophyte alga morphologically resembling mosses and living in both, the aquatic and terrestrial habitats. Because of its complex morphology and terrestrial adaptations, *Draparnaldia* can provide new insights into the evolution of multicellularity and terrestrialization in green plants. To develop *Draparnaldia* into a model, we *de novo* sequenced its genome and transcriptomes, and profiled its phytohormone repertoire. We found that 1) Expanded gene families in *Draparnaldia* with respect to unicellular *Chlamydomonas* are linked to multicellularity and abiotic stresses. 2) *Draparnaldia*’*s* terrestrial adaptations are reflected at both the morphological and molecular levels. 3) *Draparnaldia* synthesizes most of the phytohormones used by land plants to thrive in terrestrial habitats. All of this makes *Draparnaldia* a powerful model to uncover and study alternative evolutionary trajectories towards multicellularity and terrestrialization in plants.

## Introduction

Plant terrestrialization (the transition from water-to-land as the main plant habitat) is one of the major events in Earth history. It is generally accepted that land plants originated from the Streptophyte algae ^1^. Therefore, some Streptophyte algae (*Mesostigma*, *Klebsormidium*, *Chara*) and several early diverging land plants such as liverworts (*Marchantia*), mosses (*Physcomitrium*) as well as the major land plant model *Arabidopsis* are currently used to study plant terrestrialization (Figure 1A, B and Table S1 in the present study; ^2–7^). However, there is also a major sister lineage to Streptophyte algae and land plants, the Chlorophyte algae; some of which are developmentally complex and also possess adaptations to terrestrial habitats ^8–11^. Thus, a major mystery in plant evolution is why no land plant species evolved from Chlorophyte algae?

**Figure 1.**
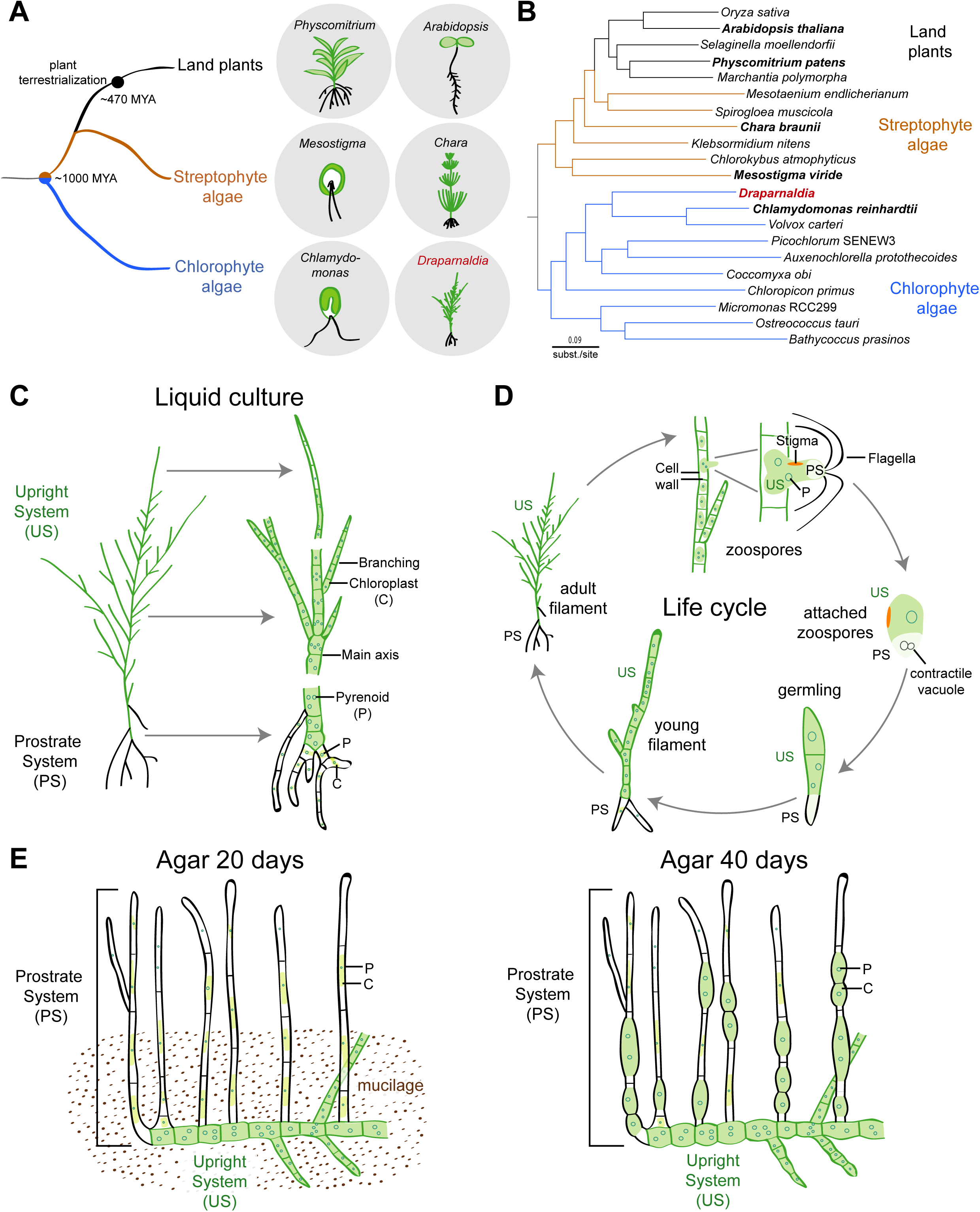
Phylogenetic position and morphology of *Draparnaldia*. **A** Schematic phylogeny of green plants consisting of three main lineages (left): Land plants (black), Streptophyte algae (brown) that diverged into land plants, and Chlorophyte algae (blue) a sister lineage of Streptophyte algae and land plants. On the right, cartoon representations of some of the species used in this study. *Draparnaldia*, a Chlorophyte alga, is labelled in red. MYA = millions of years ago. **B** Phylogenetic tree of 21 species of green plants using genome-wide concatenated orthologous genes (see also Table S1). Species drawn in Figure 1A are shown in bold in the tree. *Draparnaldia* is highlighted in red. Color code of the tree branches as in Figure 1A. **C** Morphology of *Draparnaldia* in liquid culture. The alga consists of the Upright System (US) and the Prostrate System (PS). In liquid culture the US grows upwards, whereas the Prostrate System (PS) grows downwards and anchors the alga to the substrate. The chloroplasts (C) in the PS are small in size and thus the PS is not green. Each chloroplast contains one or more pyrenoids (P). **D** Life cycle of *Draparnaldia* (in liquid culture). Only a development of zoospores, the main type of asexual reproduction, is shown. Zoospores are solely aquatic and have a clearly recognizable distinct structure of the US and the PS. **E** Morphology of *Draparnaldia* on agar. After transfer of *Draparnaldia* from liquid to solid medium, the alga loses its polarity: the US falls onto the substrate, among which the PS is formed and grows up. Agar 20 days (left): The US is surrounded by a mucilage produced by the alga. The PS is actively growing by apical growth and the chloroplast become enlarged in size. Agar 40 days (right): The mucilage disappears. The PS is no longer growing by apical growth. Instead, cells are becoming rounded. They are almost completely filled by chloroplasts and divide by intercalary growth.

To answer this question, we need to understand better the developmental complexity and terrestrialization in Chlorophyte algae as compared to its counterpart Streptophyte algae and earliest diverged land plants. The established models among Chlorophyte algae are limited to unicellular (e.g. *Chlamydomonas*, ^12^), colonial (*Volvox*, ^13^) and folious species (*Ulva*, ^14^). A multicellular filamentous Chlorophyte alga model that would morphologically resemble complex Streptophyte algae, is missing. To close this knowledge gap, we establish *Draparnaldia* as a new chlorophyte model. *Draparnaldia* has several unique features:

(*i*) It has a strategic phylogenetic position, being closely related to the unicellular *Chlamydomonas* (class Chlorophyceae), but is the most morphologically complex member of the order Chaetophorales, that contains species with filamentous body plans (see Figure 3 in Caisová 2020 ^15^). These make *Draparnaldia* an excellent model to study the evolution of complex bodies in chlorophytes.
(*ii*) It has remarkable adaptations to both, aquatic and terrestrial environment. Importantly, these adaptations can be induced under laboratory conditions. In Liquid culture (= mimicking the aquatic environment, Figure 1C, D), the Upright System (US) grows up and the Prostrate System (PS) grows down and attaches the alga to the substrate. In contrast, on Agar (= mimicking the terrestrial environment, Figure 1E), the polarity of the alga switches. The US falls on the substrate and the PS grows up. These unique adaptations of *Draparnaldia* to aquatic and terrestrial environments allow the study of terrestrialization in Chlorophyte algae.
(*iii*) It is easy to cultivate and it is fast growing. Its life cycle can be completed within 7-9 days (Figure 1D). Moreover, *Draparnaldia* reproduces via zoospores (unicellular flagellated stages), which can be induced simultaneously in large amounts allowing synchronous phenotypic screenings.
(*iv*) Importantly, there are protocols on highly efficient transient transformation, including the isolation and regeneration of the protoplasts and a list of antibiotics that can be used for selection of transformants ^16^.

Together, this makes *Draparnaldia* a uniquely suitable and easily accessible model to algal and plant biologists.

To further broaden this tool set and establish *Draparnaldia* as a genetic model, here we present a comprehensive assessment of *Draparnadia’s* genome, transcriptome and hormone repertoire. This represents a first genome reported for a morphologically complex Chlorophyte alga; additionally with adaptations to terrestrial-like environment. The *Draparnaldia* genome has the potential to shed light on the evolution of complex Chlorophyte algae and allow to better understand the traits that may have allowed for the rise of land plants.

## Results Taxonomic acts

Genus *Draparnaldia* currently contains 24 accepted species ^17,18^. The *Draparnaldia* CCAC 6921 strain from which the genome has been sequenced cannot be morphologically assigned to any of the 24 species and is therefore described here as the new species: *Draparnaldia erecta* Caisová & Friml sp. nov., Figure S1. Furthermore, we emended the description of the genus *Draparnaldia* (Roth) Bory 1808. Ann. Mus. Natl. Hist., 12: 399-409. emend. to include information on the morphology of the genus in culture and on the arrangement of the Prostrate System (= rhizoids) in water and dry environment. All taxonomic requirements necessary for valid species description and genus emendation are in Note S1.

## Genome assembly and functional annotations

The *Draparnaldia de novo* genome represents the first genome of a multicellular filamentous Chlorophyte alga. The genome assembly, sequenced using a combination of Nanopore and Illumina, has a scaffold N50 of 195 kb, N90 of 49 kb and a span of 238 Mb (Figure 2A, Figure S2, Table S2). A high completeness was confirmed with two independent methods: Merqury k-mer completeness is over 90%, and BUSCO gene completeness is 96.4% (Figure 2A, Figure S2B). The genomic GC content is 58% (Figure 2A).

**Figure 2.**
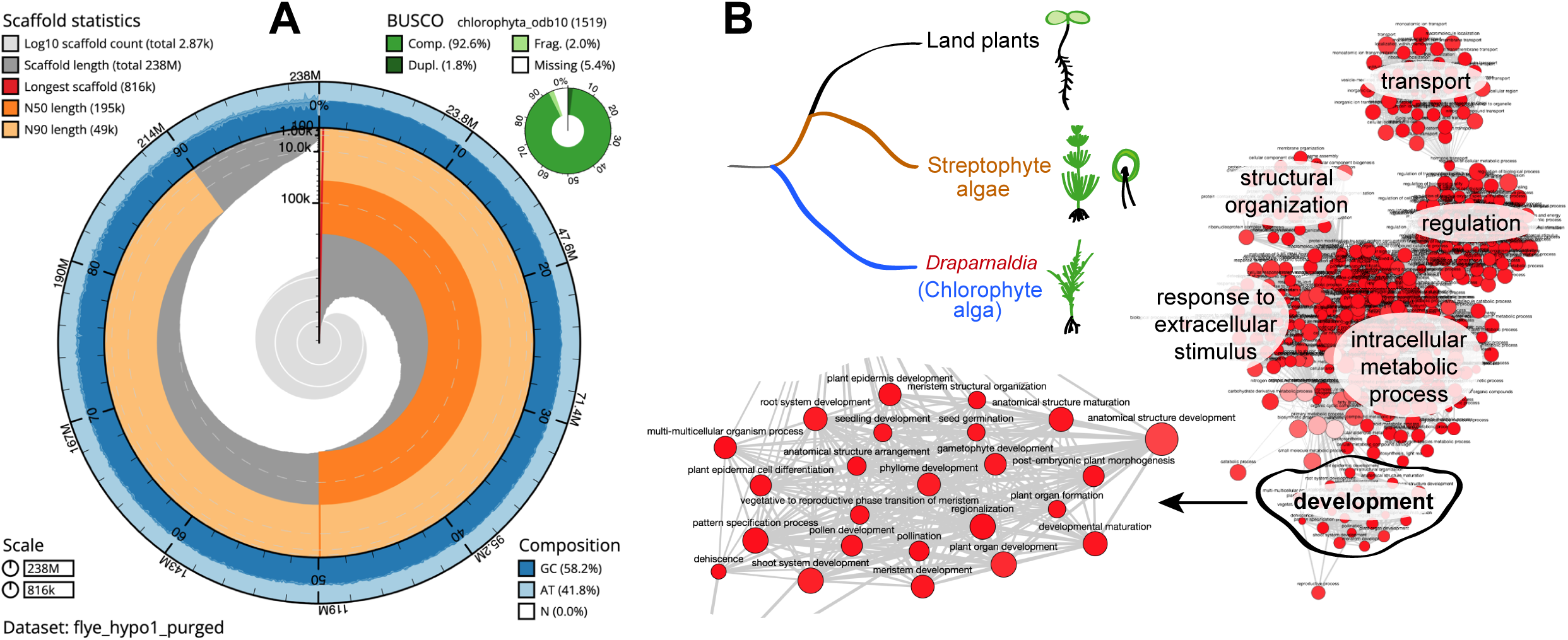
Genome assembly and functional annotations. **A** Snail plot and summary of assembly statistics for *Draparnadia*’s genome. The main plot is divided into 1,000 size-ordered bins around the circumference with each bin representing 0.1% of the 238 Mbp assembly. The distribution of scaffold lengths is shown in dark grey with the plot radius scaled to the longest scaffold present in the assembly (816 kb, shown in red). Orange and pale-orange arcs show the N50 and N90 scaffold lengths (195 kb and 49 kb, respectively). The pale grey spiral shows the cumulative scaffold count on a log scale with white scale lines showing successive orders of magnitude. The blue and pale-blue area around the outside of the plot shows the distribution of GC, AT and N percentages in the same bins as the inner plot. A summary of complete, fragmented, duplicated, and missing BUSCO genes in the chlorophyta_odb10 set is shown in the top right. **B** Conserved processes and pathways between the genomes of *Draparnaldia*, Streptophyte algae and land plants calculated with REVIGO. Top left: Tree schematic of the species used for functional clusters: *Draparnaldia*, Streptophyte algae (unicellular and multicellular species) and land plants. Right: Network representation of the common GO terms related to biological processes. Unconnected nodes were removed to provide clarity of the network. Six main functional clusters of GO terms have been identified: transport, structural organization, regulation, response to extracellular stimulus, intracellular metabolic process, and development. A zoom-in into the ‘development’ cluster is shown. The dataset used was imported from REVIGO (source data in Table S3).

Approximately 40% of the assembled genome consists of repeats, while approximately 20% of the genome are exons (for reference, ∼29% of the genome of the model plant *Arabidopsis thaliana* consists of protein-coding exons). Retrotransposons were identified 8x times more frequently than DNA transposons. LINEs are the most frequent transposons, while LTRs cover the longest region of the genome. The three most commonly occurring transposon families are RTE-X, Gypsy and I-Jockey, all retrotransposons.

RNA sequencing of three vegetative stages (Figure 1C-E) was used together with *ab initio* gene prediction and sequence conservation analyses to annotate the genome. In total, we annotated 19,081 protein-coding genes that produce 27,433 transcripts and encode for 26,190 unique protein products. Comparative genomics and sequence conservation analyses allowed us to assign functionality to 42% of the predicted proteins. Most (94%) of the annotated transcripts are multi-exonic, encoding 7.59 exons on average (Table S2). In addition, 15,487 non-coding transcripts were annotated, of which 10,037 and 5,450 are long and short non-coding RNA genes, respectively.

### Shared gene clusters between *Drapanaldia* and streptophytes

Within its life cycle, *Draparnaldia* morphologically resemble Streptophyte algae and mosses. Thus, to identify conserved processes and pathways related to the development between *Draparnaldia* and the streptophyte lineage, we searched for common orthogroups (= a group of genes across different species that descend from a common ancestral gene) among *Draparnaldia*, Streptophyte algae (*Chlorokybus*, *Mesostigma*, *Mesotaenium*, *Spirogloea*, *Chara*, *Klebsormidium*) and land plants (Figure 2B left panel, Table S3). Gene Ontology (GO) enrichment analyses of the conserved orthogroups among all compared species grouped into 6 significant GO clusters: ‘transport’, ‘structural organization’, ‘regulation’, ‘response to extracellular stimulus’, ‘intracellular metabolic process’ and ‘development’.

The ‘development’ cluster includes GO terms such as ‘shoot system development’ (GO:0048367), ‘phyllome development’ (GO:0048827), ‘root system development’ (GO:0022622) and ‘pollen development’ (GO:0009555). This is in agreement with the fact that *Draparnaldia* is developmentally complex and has two clearly distinguishable structures (the US (=Upright System) and PS (= Prostrate System)). From all *Draparnaldia* morphological structures, the PS (= Prostrate System) is the one that most resembles land plant organs morphologically and likely also functionally, specifically the tip-growing filamentous rooting structures (i.e. rhizoids and root hairs). Thus, we investigated the conserved orthogroups that were annotated with the GO term **‘**Root system development’ (GO:0022622) and also ‘Pollen development’ (GO:0009555), as it has been shown that the growth of filamentous rooting structures and pollen tubes, which also have tip growth, is controlled by overlapping gene regulatory networks ^19,20^. Associated with these GO terms, we identified orthogroups involved in processes that are fundamental for root-related tip growth such as MATE efflux family protein (OG0000040) regulating turgor pressure and RAS-related GTPases (OG0000067) controlling vesical secretion (=membrane trafficking) ^21^. Additionally, we identified Auxin-responsive GH3 family proteins (OG0000392), glutamate receptors (OG0000445) and ALBA (OG0002116) that are involved in the elongation of root-related structures ^22–24^. Interestingly, we also identified the PAS domain from the PAS/LOV protein (orthogroup OG0007343), which in *Arabidopsis* has been described as a blue light receptor and might play a role in the ‘root greening’, i. e. the development of chloroplasts in roots ^25^. This process resembles the ‘PS greening’ that occurs in *Draparnaldia* when it is transferred from the liquid culture to agar (compare Figure 1C and Figure 1E).

In summary, these analyses identified interesting genes and clusters in *Draparnaldia* providing potential molecular insights into an alternative evolution of morphological complexity and terrestrialization in chlorophytes versus streptophytes.

### Convergent and lineage-specific expansions associated with multicellularity in chlorophytes

Multicellularity evolved independently multiple times and likely required complex rearrangements of the genomes, such as expansions and contractions of functional networks and genes. Thus, we compared gene family evolution between the multicellular *Draparnaldia* and the unicellular *Chlamydomonas,* both belonging to the class Chlorophyceae. We were interested in orthogroups that were expanded or contracted in *Draparnaldia*, but not in *Chlamydomonas*. This direct comparison of unicellular and multicellular members within the same class provides a unique opportunity to pinpoint the pathways and gene families coopted for this alternative rise of multicellularity.

Our analyses revealed 24 expanded orthogroups in *Draparnaldia* with respect to *Chlamydomonas* (Figure S3), of which 22 were annotated with GO terms (Table S4). In addition, there was a single contracted orthogroup in *Draparnaldia* with respect to *Chlamydomonas* that was not significantly associated with any GO term enrichment. In the expanded orthogroups of *Draparnaldia*, we observed an overrepresentation of several GO terms associated with multicellularity and abiotic stress responses (Figure 3A). Some of those seem to mirror changes that occurred in land plants. For instance, among the expanded orthogroups are early light-inducible like proteins (ELIPs) (OG0000047). This core group of proteins expanded in *Draparnaldia* (*Draparnaldia* 12 copies, *Chlamydomonas* 7 copies) and in early divergent land plants (*Physcomitrium* and *Marchantia*). In *Physcomitrium*, ELIPs are hypothesized to be involved in the response to high-intensity light and drought stress ^26^. Thus, in *Draparnaldia*, ELIPs may reflect adaption to drought and to light stress when the alga is growing on agar, mimicking the terrestrial habitat. Another example of an expanded orthogroup in *Draparnaldia* is the ‘glycoside hydrolase’ (= cellulase) (OG0000039) with 11 copies in *Draparnaldia* and 2 copies in *Chlamydomonas*. In land plants, cellulase plays a potential role in elongation ^27^. Because *Draparnaldia* is multicellular and it has a Prostrate System with a tip growth, it might be that cellulose is involved in elongation of *Draparnaldia*’s cells, as is the case in land plants. Further work is required to experimentally identify the actual substrate for these putative enzymes.

To better characterize the convergent and divergent processes underlying the shift to multicellularity, we performed a similar analysis in streptophytes by inferring gene families’ expansions/contractions associated with the evolution of developmental complexity (Table S4). We identified 40/6 orthogroups that expanded/contracted in *Chara* with respect to *Mesostigma*, which are multi– and unicellular algae, respectively and live in comparable habitats as *Draparnaldia* and *Chlamydomonas* (Figure 3B). We did not, however, find any significantly overrepresented GO terms associated with these orthogroups. This could be explained either by:

(i) Multicellularity in Streptophyte algae evolved in different way than in Chlorophyte algae, or
(ii) The large evolutionary distance between *Chara* and *Mesostigma* reduced the power to detect gene expansion/contractions due to high sequence divergence.

**Figure 3.**
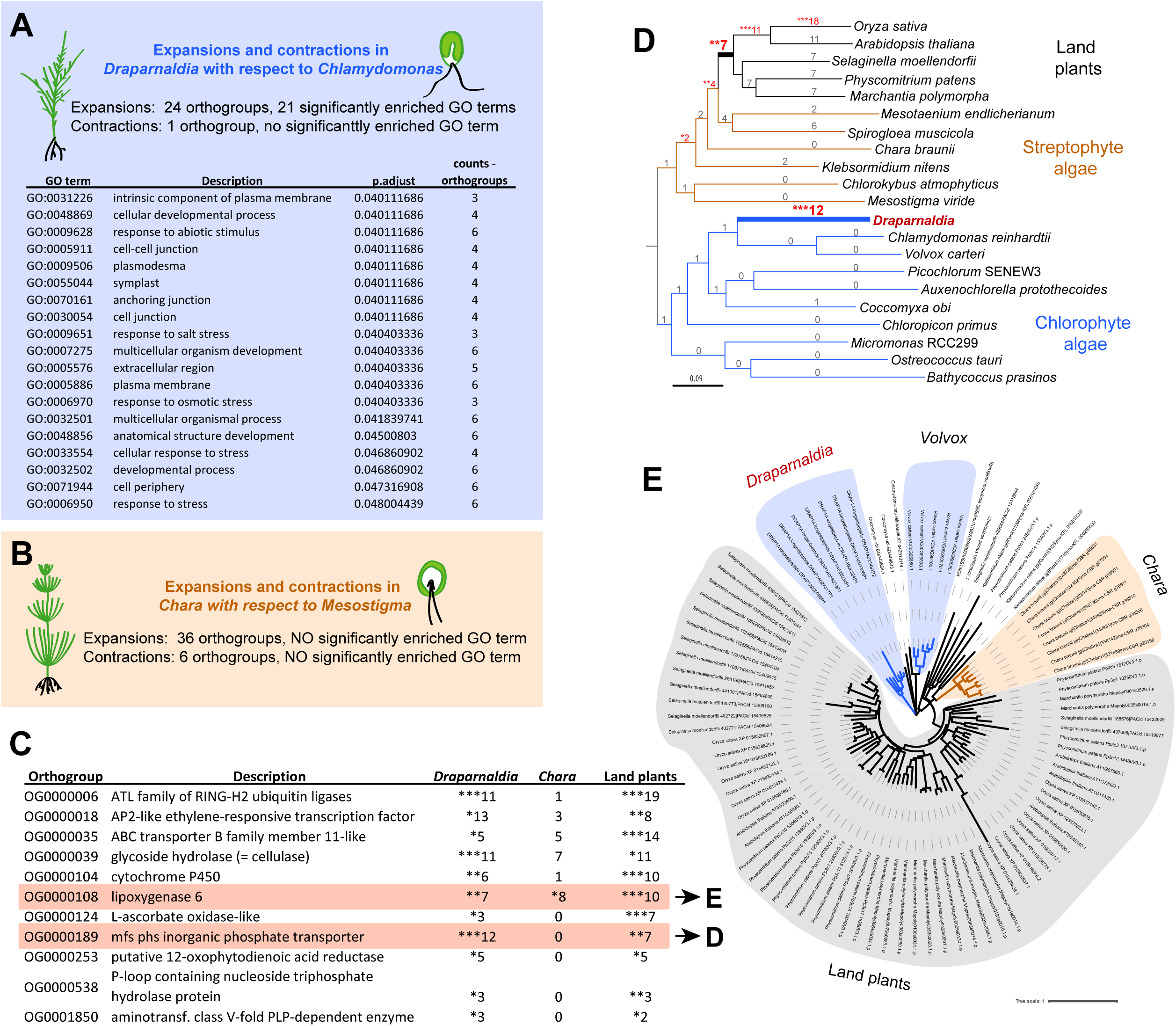
Convergent and lineage-specific expansions associated with multicellularity. **A** GO enrichment analysis of expanded and contracted orthogroups in *Draparnaldia* with respect to *Chlamydomonas*. Only the significant GO terms that have ≥ 3 orthogroups are shown. Significant GO terms that have ≤ 3 orthogroups can be found in Table S4. **B** No significant GO terms were identified among the expansions and contractions in *Chara* with respect to *Mesostigma*. **C** Shared orthogroup expansions identified between multicellular species (*Draparnaldia* – Chlorophyte alga, *Chara* – Streptophyte alga, and land plants). Source data for comparison in Figure S3. **D, E** Examples of shared expanded orthogroups: **D** Phylogenetic tree of the phosphate transporter with a high number of gene copies in *Draparnaldia* (in contrast to other chlorophytes), * p < 0.05, ** p < 0.01, p < 0.001. **E** Phylogenetic tree of Lipoxygenase, showing independent expansions in all four multicellular lineages, including *Volvox*. Only orthogroups that expanded to more than 3 copies were considered for the analyses.

Since the expanded/contracted orthogroups in *Chara* versus *Mesostigma* did not lead to any significant GO terms, we broadened our analysis and searched for orthogroup expansions and contractions that are shared by at least two of three independent multicellular ‘groups’: *Draparnaldia*, multicellular Streptophyte algae (*Klebsormidium* and *Chara*)) and land plants (Figure 3C, Table S4). The unicellular *Chlamydomonas* and *Mesostigma* were used as a control. We used the whole phylogeny to be able to pinpoint where the events in the ancestral branch leading to land plants happened. We identified one orthogroup that is contracted in both *Draparnaldia* and *Chara* (OG0000029, Table S4) and 11 orthogroups that are expanded in both *Draparnaldia* and land plants. Among the common expanded orthogroups are ‘mfs phs inorganic phosphate transporters’ (OG0000189), (Figure 3D). Inorganic phosphate (Pi) is an essential substrate for the healthy growth of land plants. The amount of phosphate (Pi) in soil is however limited ^28^. To optimize phosphate usage, land plants have evolved specialized transporters that are responsible for the Pi uptake by roots (reviewed by Młodzińska and Zboińska 2016 ^29^). In *Draparnaldia*, the high number of copies of phosphate transporters might serve as an adaptive strategy to cope with limited Pi availability when the alga is growing in the terrestrial habitat.

Moreover, we also identified one orthogroup lipoxygenase 6 (OG0000108) that is expanded in all three multicellular groups, *Draparnaldia*, *Chara* and land plants (Figure 3E). Plant lipoxygenases (LOXs) play an important role in response to biotic and abiotic stresses. Specifically, LOX6 contributes to the jasmonate synthesis, which makes plants less attractive to crustacean and more protective to drought ^30,31^. Both, *Draparnaldia* and *Chara* can live in habitats where they can be exposed to land plant pathogens.

In summary, *Draparnaldia* genome provides the first genomic insights in the evolution of multicellularity and body plan specification in chlorophytes and an independent reference to multicellularity evolution in the land plant lineage. Overall, this offers completely novel perspectives and opportunities for studies on the evolution of multicellularity in plants, in general.

### *Draparnaldia* transcriptome from aquatic and terrestrial habitats

*Draparnaldia* can be grown in liquid culture and on agar, which mimic its aquatic and terrestrial habitat, respectively. To investigate the gene expression changes involved in *Draparnaldia’s* transition between these two habitats, we compared the transcriptomes of the alga growing in Liquid culture versus Agar 20 days (= 20 days after being transferred to agar), and Agar 20 days versus Agar 40 days (= 40 days after transfer to agar), (Figure 4A). We identified a total of 2786 differentially expressed genes (DEG) between these three conditions (Table S5). There were 1414 up regulated genes and 1047 down regulated genes in Liquid culture compared to Agar 20 days, highlighting the extensive rewiring of the transcriptome needed for surviving in these different habitats. There were much fewer differences between the two agar cultures, as expected given that these corresponded to different time points but to the same environment: only 103 genes were upregulated and 222 down regulated when comparing Agar 20 days to Agar 40 days (Figure 4B).

**Figure 4.**
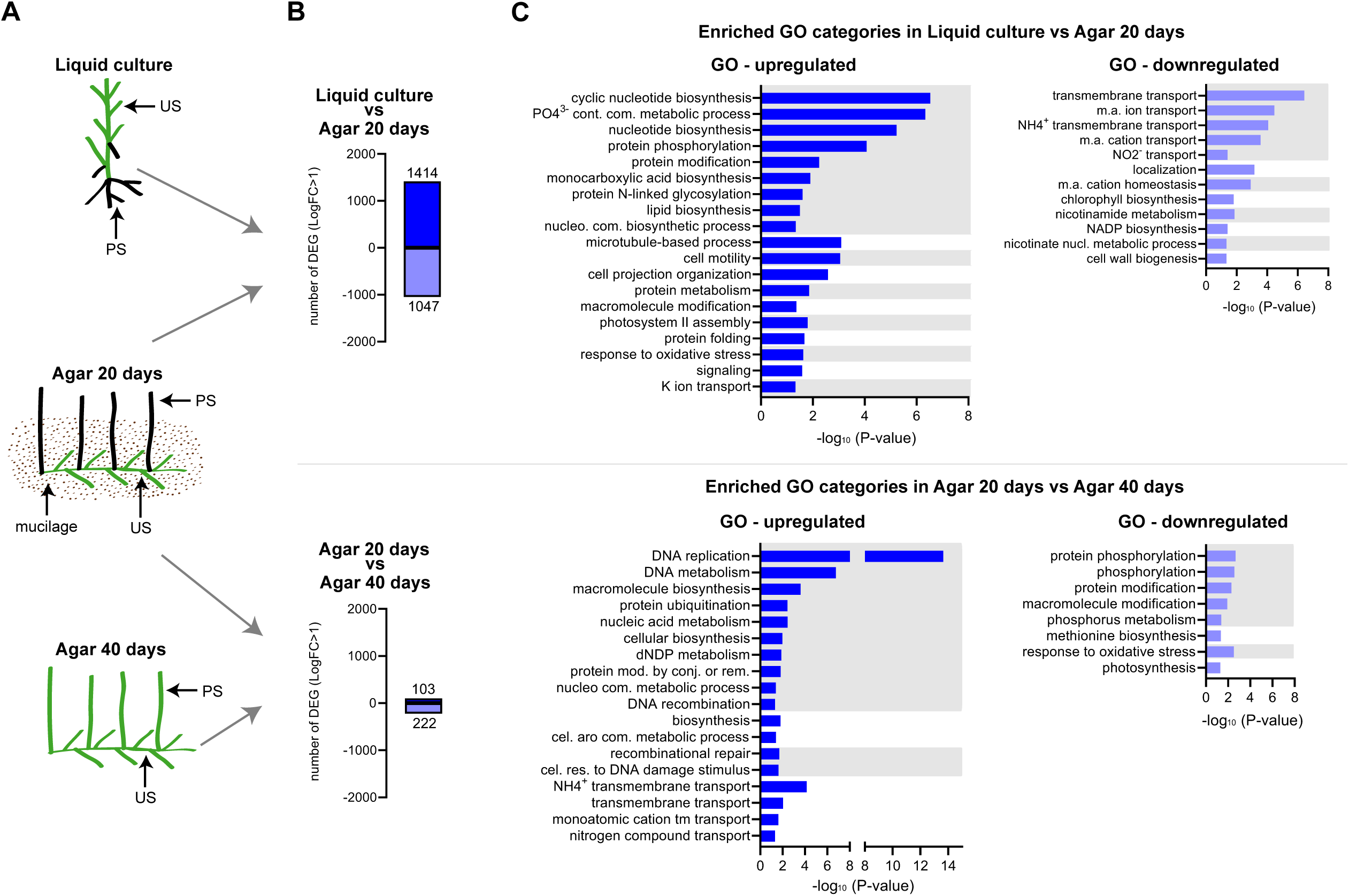
Draparnaldia transcriptome from aquatic and terrestrial habitats. **A** Three different morphologies (= growing conditions) of *Draparnaldia* used for gene expression analysis. **B** Absolute number of differentially expressed genes between different growing conditions. **C** Gene ontology (GO) enrichment of the differentially expressed genes in liquid culture versus agar. Only biological processes are shown. Up-regulated genes are depicted in dark blue, down-regulated genes in pale blue, and superclusters from the TreeMap view of REVIGO grouped by grey and white background. Abbreviations: aro. – aromatic, cel. – cellular, com. – compound, conj. – conjugation, cont. – containing, m.a. – monoatomicmod. – modification, nucl. – nucleotide, nucleo. – nucleobase-containing, rem. – removal, res. – response.

To evaluate the biological function of DEG between conditions, we conducted a Gene Ontology (GO) enrichment analysis (for more details see Material and Methods) (Figure 4C, Table S5). Among genes upregulated in Liquid culture (relative to Agar 20 days), where the Upright System grows up and Prostrate System grows down, GO terms, such as ‘cell motility’ (GO:0048870), ‘nucleotide biosynthesis’ (GO:0009165), lipid biosynthesis (GO:0008610) and ‘protein phosphorylation’ (GO:0006468) were enriched. This likely reflects the fact that in Liquid culture, the alga is reproducing and actively growing. In contrast, GO terms enriched among upregulated genes in Agar 20 days refer to ‘transport’ (GO:0055085, GO:0015707, GO:0072488) and ‘homeostasis’ (GO:0055080). This highlights *Draparnaldia*’s adaptations to the limited availability of nutrition and different osmotic pressure on land as compared to water. Among other GO terms enriched among genes upregulated in Agar 20 days are ‘cell wall biogenesis’ (GO:0042546) and ‘chlorophyll biosynthesis’ (GO:0015995). These terms are likely associated with the morphological change of the alga on agar, specifically with the upright growth of the PS and with the enlargement of the chloroplasts in size in the PS, as discussed previously. Biological processes such as DNA replication (GO:0006260) and metabolism (GO:0006259, GO:0090304), as well as biosynthesis processes (GO:0044249, GO:0009058), are enriched among genes upregulated at Agar 20 days (relative to Agar 40 days), suggesting that the alga is actively growing. In contrast, genes upregulated at Agar 40 days are enriched for protein phosphorylation (GO:0006468), response to oxidative stress (GO:0006979) and photosynthesis (GO:0015979), which are likely associated with the osmotic stress, higher temperature, drought and salinity.

Overall, transcriptome analyses of *Draparnaldia* growing in Liquid culture and Agar identified plausible processes and candidate genes with an important role in chlorophytes terrestrialization, which provides an essential base for comparative analyses with streptophytes.

### Protein-protein interaction network of differentially accumulated ‘Transmembrane transport’ transcripts

One of the prominent enriched GO terms among genes upregulated in Agar 20 days against Liquid culture was ‘Transmembrane transport’ (GO:0055085, Table S5). Transmembrane transport is considered to be one of the essential adaptations for the terrestrial life of plants, as it allows plants to deal with drought stress and limited availability of nutrition ^32,33^.We therefore searched for specific pathways and processes included in this GO enrichment. We analyzed potential protein-protein interactions of the orthogroups enriched for ‘Transmembrane transport’ in *Draparnaldia*, by looking at interactions of their *Arabidopsis* homologues in the STRING database (Methods, Table S6). This yielded a network of proteins consisting of three main clusters: nitrogen transport, ion homeostasis and hormone transport:

Figure 5A shows a cluster with ammonium transporters (AMT1s) and the urea-proton symporter DUR3, which in land plants are involved in nitrogen uptake under nitrogen-deficiency conditions^34,35^. We hypothesize that these components might have a similar function in *Draparnaldia* growing on agar, where the alga has only limited contact with the medium (only using the US, but not the PS) and hence has limited contact with nutrition (Figure 1E). Thus, the alga has to activate a signaling network dealing with limited nitrogen availability, compared to Liquid culture, where both the US and the PS are in contact with medium.

**Figure 5.**
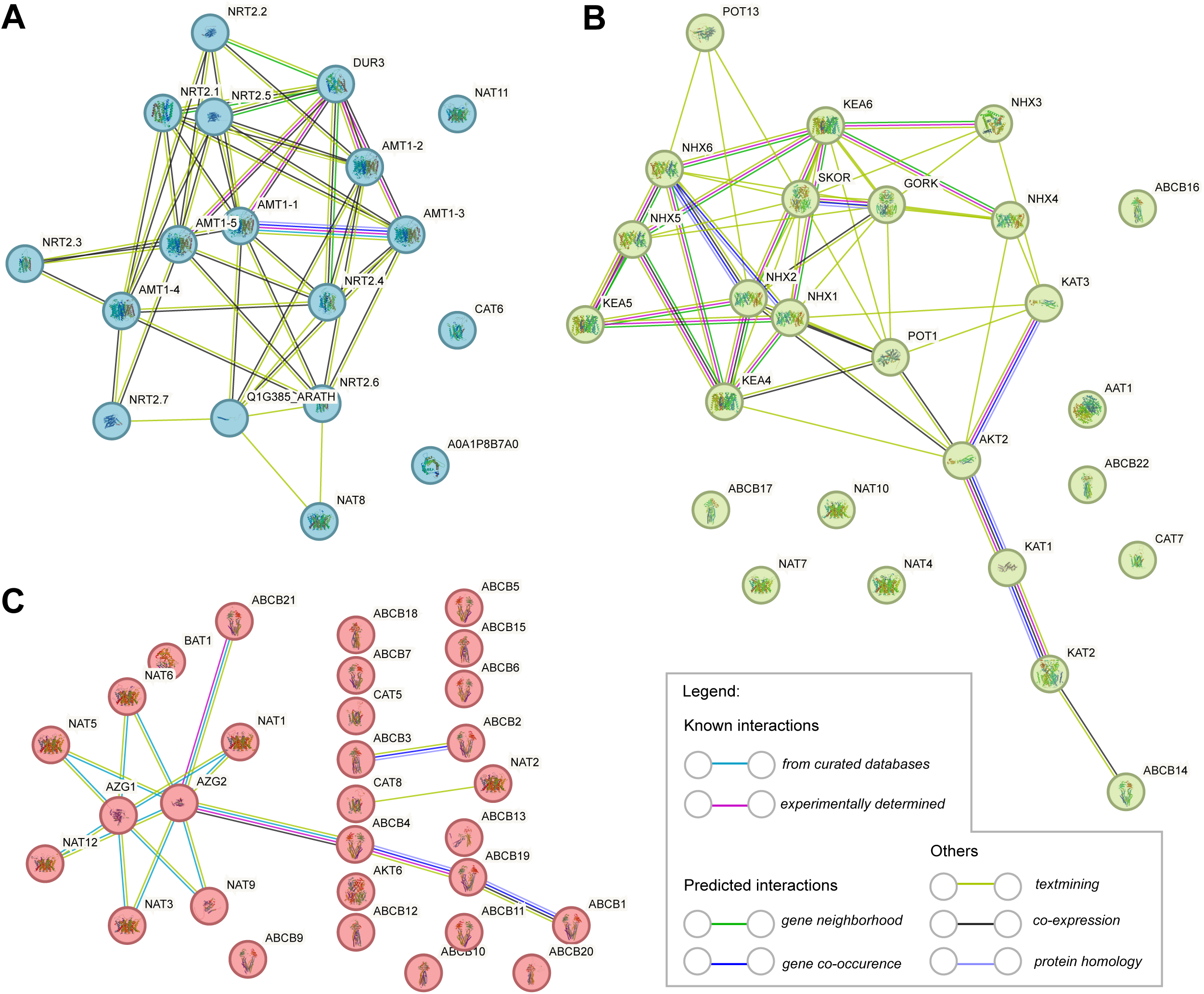
Protein-protein interaction network of differentially accumulated ‘Transmembrane transport’ transcripts. This GO term was upregulated in Agar 20 days but not in Liquid culture (Figure 4C). Three major clusters were detected by the STRING database based on known interaction from *Arabidopsis*. A Protein interactions involved in nitrogen transport. B Protein interactions of potassium channels and sodium/hydrogen exchanger. C. Protein interactions involved in cytokine and auxin transport.

Figure 5B shows cluster of proteins including potassium channels (KAT, KEA, SKOR, GORK) and sodium/hydrogen exchangers (NHX) that in land plants seem to be critical for maintaining proper cytosolic osmolarity in response to drought and salt stress ^36–38^. The function of these channels is probably conserved between *Draparnaldia* and land plants. As agar dries up, osmolarity increases. *Draparnaldia* maintains the osmotic homeostasis by up-regulating potassium channels and sodium/hydrogen exchangers that will pump Na^+^ into the vacuole.

Figure 5C shows interactions of proteins member of the nucleobase-ascorbate transporter (NAT) family ^39^ and proteins associated with hormone transport in land plants. For instance, the AZA-GUANINE RESISTANTs (AZGs) are cytokinin transporters ^40,41^ and ABCB family, for i which transporters for Brassinosteroid (ABCB19, ABCB1) and auxin (ABCB4) have been studied ^42,43^. In land plants, homologues of some of the transporters we found up-regulated in *Draparnaldia* are important for plant growth, development and resistance to stress. For example, the NAT12 plays an important role in regulating *Gossypium* (cotton) resistance to salt and drought stress ^44^. AZG proteins can mediate changes in the root architecture of *Arabidopsis*. Growth of *Arabidopsis* under drought or salt stress induces AZG expression that leads to either reduction or a lack of lateral roots ^40^. Similarly, *Draparnaldia* grown in agar has enhanced AZG expression in comparison to that grown in Liquid culture and it has mainly unbranched PS.

Taken together, in *Draparnaldia* these analyses identified groups of proteins related to nitrogen transport, ion homeostasis and hormone transport; all of which are important to sustain plant life on land. Thus, our analysis of the ‘membrane transporter’ GO term provided an example how our *Draparnaldia* genome and transcriptome data can be mined to obtain more specific insights into alternative adaptation to land environment.

### Differential expression of ethylene and abscisic acid signaling components in *Draparnaldia*

As differentially regulated transcripts in *Draparnaldia* code for proteins associated with the hormone transport, we investigated whether hormone signaling components are also differentially regulated between Liquid culture and Agar. We identified differentially regulated components of two hormonal pathways between these two environments: ethylene and abscisic acid.

Firstly, we found ethylene response transcription factor (ERF – DRAP1A013737) upregulated in Liquid culture compared to either Agar conditions (Figure 6 left panel). A phylogenetic tree of the ERFs in land plants showed that *Draparnaldia*’s ERF belongs to the same clade as *Arabidopsis* ERF71 and ERF73 (Figure S4A, Table S7). In *Arabidopsis* these two genes are downstream components of ethylene signaling and are involved in the hypoxic stress responses triggered during flooding ^45^. This is an interesting finding, because in the *Draparnaldia’s* transcriptome and genome we did not find homologues of the components of the ethylene signaling pathway upstream from EIN3/EIL, but the ethylene downstream components are present. Moreover, in Liquid culture *Draparnaldia* develops the more complex PS (longer, more branched and dense) than on Agar. This feature of development seems to resemble land plants, which under hypoxic stress produce adventitious roots ^46^ and longer root hairs ^47^ (Figure 6, schematics in grey background).

**Figure 6.**
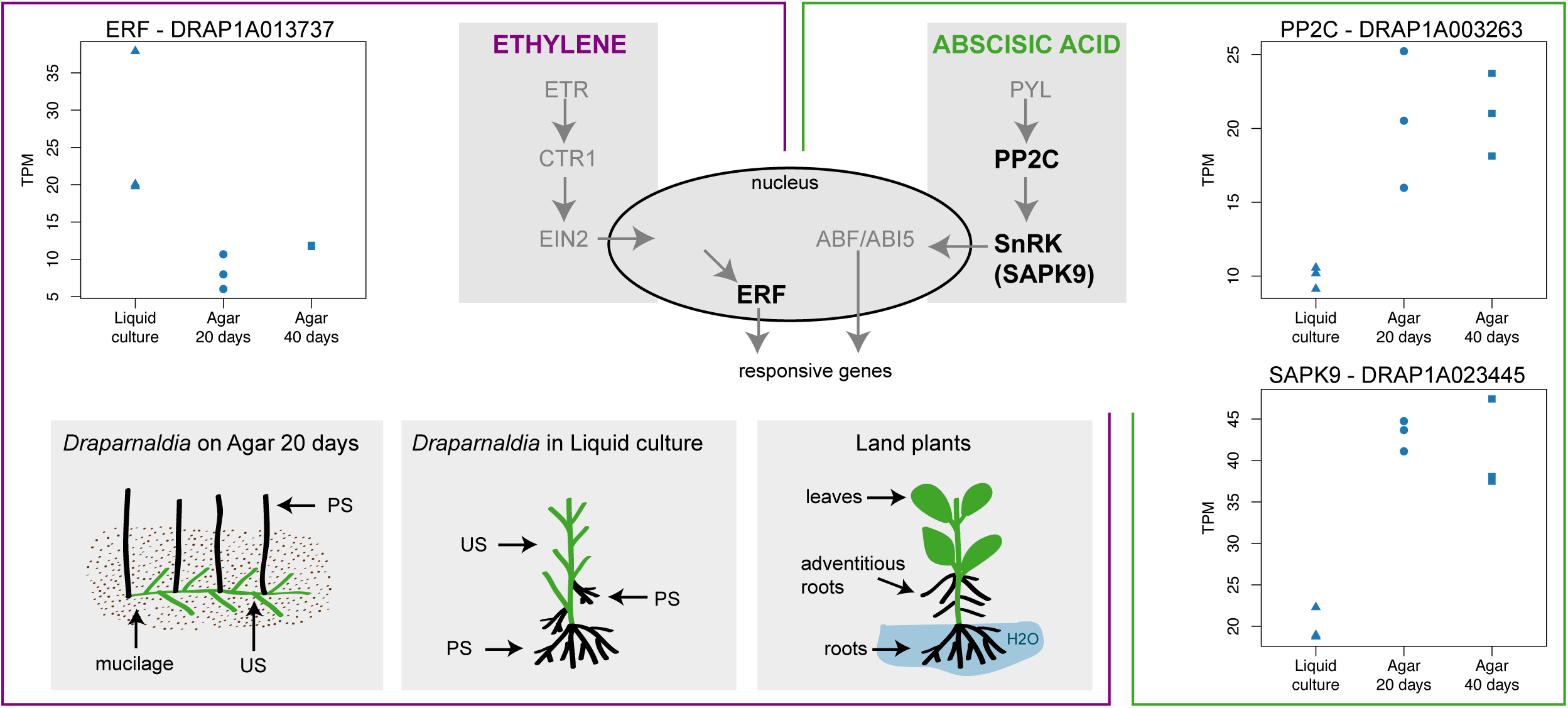
Differential expression of ethylene and abscisic acid signaling genes in *Draparnaldia*. Simplified schematics of the *Arabidopsis* abscisic acid (violet, left box) and ethylene (green, right box) pathways. Common players in both signaling pathways are shaded and encircled in the middle. In bold are the respective homologue enzymes from *Arabidopsis* that were differentially expressed in the *Draparnaldia* transcriptome. Genes in grey were not found in the *Draparnaldia* transcriptome, nor in the genome sequence. Plots of TPM gene counts of the relevant enzymes expressed in *Draparnaldia’s* Liquid culture, Agar 20 days, and Agar 40 days are shown. *Draparnaldia* homologues for PP2C and SAPK9 involved in ABA signaling were upregulated on Agar. Whereas *Draparnaldia* homologue for ethylene signaling (ERF) was upregulated in Liquid culture.

Abscisic acid (ABA) mediates drought stress response in land plants. The core ABA signaling cascade in land plants consists of the PYR/PYL ABA receptors, type 2C protein phosphatases (PP2Cs) clade A and Snf1-related protein kinases 2 (SnRK2s) ^48^. In *Draparnaldia*, we found DRAP1A003263 in the same orthogroup as PP2C genes involved in ABA signaling in *Arabidopsis* (Figure S4B, Table S7), ^49^. We also found a homologue of the SAPK9 (DRAP1A023445), which is a member of SnRK2s and is a positive regulator of ABA-mediated stress signaling in rice ^50^. Whereas the SAPK9 was upregulated on Agar compared to Liquid culture as we might expect, the PP2Cs was accumulated at higher levels when growing on Agar compared to Liquid culture. This is in contrast to how the PP2Cs function in *Arabidopsis* after ABA treatment. A role of ABA in algae remains enigmatic and crucial parts of a pathway (the receptors) are missing, except the PYL homologous proteins found in Zygnematophyceae algae ^51^. Exogenous ABA was shown to increase stress resistance in *Chlamydomonas*, but an exact mechanism is unclear ^52,53^. *Draparnaldia* as a model organism might help to clarify mechanism of ABA signaling. For example, type 2C protein phosphatases (PP2Cs) are evolutionary conserved. PP2Cs have much less copies in algae and lower plants then those in higher plants (Figure S4B, Table S7). It has been hypothesized that this expansion of PP2C family may correlate with an exposure to abiotic stresses on land ^48^. Interestingly, *Draparnaldia* has the highest number of gene copies from clade A of PP2Cs when compared to the gene copies of other chlorophyte algae analyzed (Figure S4B). As the clade A is in *Arabidopsis* involved in ABA signaling, it would be interesting to see if the higher number of PP2C genes in *Draparnaldia* is related to its ability to cope with draught stress as compared to other Chlorophyte algae.

The differential regulation of ethylene and ABA downstream signaling components between aquatic and terrestrial environment provide initial insights into common strategies how both Streptophyte and Chlorophyte algae adapt to terrestrial existence.

### Presence of phytohormones in *Draparnaldia* versus *Klebsormidium*

To complement our genome and transcriptome insights and get a more complete picture of hormones in *Draparnaldia*, we (1) experimentally surveyed the repertoire of plant hormones in *Draparnaldia*’s biomass and medium using tandem mass spectrometry (Figure 7A, blue background; and Table S8) and (2) searched for homologues of known genes involved in hormones biosynthesis, transport and signaling in *Draparnaldia*’s genome (Figure 7B, blue background; and Table S8). To have a comparison between multicellular Chlorophyte and Streptophyte alga, we also performed the comparative analyses with *Klebsormidium* (Figure 7, orange background). We have detected the presence of following hormones in *Draparnaldia* and *Klebsormidium*:

**Figure 7.**
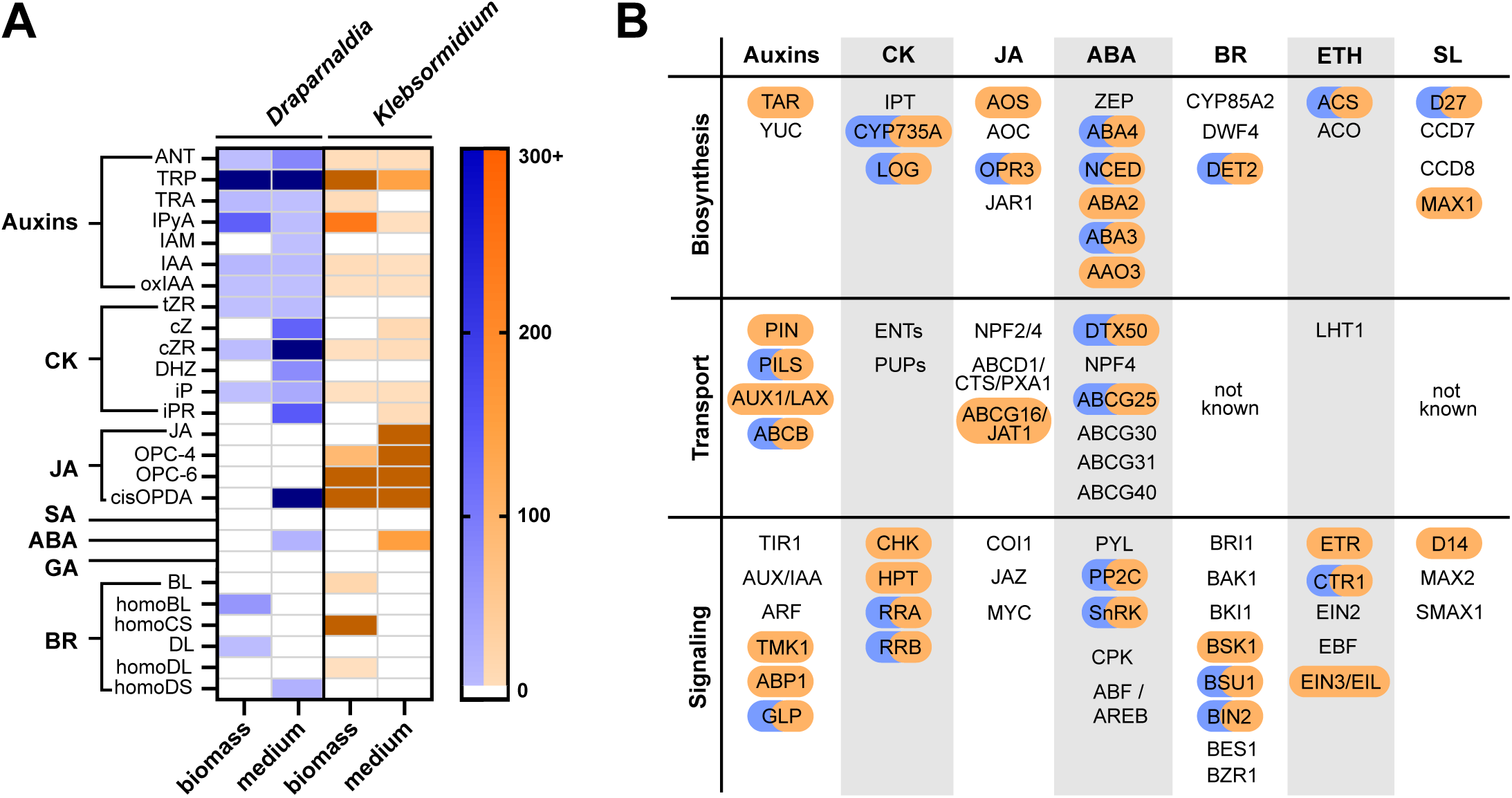
Presence of phytohormones in *Draparnaldia* versus *Klebsormidium*. **A** Heat map of abundance of the endogenous phytohormone compounds detected in the biomass and in the growth media of *Draparnaldia* (blue)and *Klebsormidium* (brown). Darker shades represent higher abundance of the compound, whereas lighter shades represent lower abundance. No color indicates that the compound was not detected. Concentration of phytohormones in biomass has been quantified in pmol*g^-1^ FW and in the growth media in pmol*ml^-1^. CK: cytokinins, JA: jasmonic acid, SA: salicylic acid, ABA: abscisic acid, GA: gibberellins, BR: brassinosteroids. The raw data for the heat map can be found in Table S8. **B** Reconstruction of the main plant phytohormone biosynthesis, transport and signaling pathways with the enzymes found in the genomes of *Draparnaldia* (blue) and *Klebsormidium* (brown). Colored boxes indicate the presence of genes in the pathways, no color indicates the absence of genes. All searches were done using HMMER (E-value < 1 x 10^-3^). Raw data in Table S9.

**Auxin** – There are four auxin biosynthetic pathways characterized in *Arabidopsis* based on their main intermediate products (IPyA, IAN, IAM, TRA). All metabolites from these pathways, except IAM, were detected in both *Draparnaldia* and *Klebsormidium*. However, the homologues of genes (TAR, YUC) encoding the enzymes of the main *Arabidopsis* IPyA pathway, were absent in *Draparnaldia*. In *Klebsormidium*, only TAR genes were present. Notably, all auxin’s conjugates were missing in both algae and for degradation pathways, only the first oxidative product (oxIAA) was detected. From four plant families involved in auxin transport, two homologues (PILS, ABCB) were present in *Draparnaldia*, whereas homologues of all transporters (PIN, PILS, AUX1/LAX, ABCB) were found in *Klebsormidium*. Regarding the signaling, two mechanisms were described in *Arabidopsis*. The homologs of proteins from the canonical TIR1/AFBs-Aux/IAA-ARF nuclear signaling pathway ^54^ were completely absent in both algal groups. From the cell-surface ABP1/ABL-TMK pathway ^55^, we recovered several GERMIN-LIKE PROTEINS, which might be potential candidates for ABP1-LIKE (ABL) auxin receptors in *Draparnaldia*. In *Klebsormidium*, all three gene families were present. These suggests that *Draparnaldia* has alternative mechanisms auxin biosynthesis and large part of transport and signaling as compared to land plant lineage.

**Cytokinins** – In *Draparnaldia*, we detected the DHZ-, tZR-, iP-, iPR-, cZ-, cZR– and MeS-type CKs. In *Klebsormidium*, all but tZR, DHZ were also found. The fact that we were unable to quantify tZR and DHZ in *Klebsormidium* could be explained by the growth conditions or the age of the culture, as previous reports have successfully detected them in this alga ^56^. Moreover, two CK-plant related homologous genes involved in biosynthesis (CYP735A, LOG) were identified in both algae. CKs conjugates were absent in both algae. No homologues of plant genes involved in CK transport were found in *Draparnaldia* or *Klebsormidium* genomes. From the CKs signaling pathways, only potential response regulators (RRA, RRB) were found in *Draparnaldia*. In contrast, genes for the complete multistep phosphorylation pathway (CHK, HPT, RRA, RRB) were present in *Klebsormidium*. Thus, despite the fact that *Draparnaldia* can produce all major types of cytokinins, it evolved an alternative signaling mechanisms.

**Jasmonate** – In *Draparnaldia* only cis-OPDA, a biosynthetic precursor of JA was detected. It is unclear however, whether it is involved in the jasmonate pathway in *Draparnaldia*, as OPDA is also a signaling molecule with functions independent of jasmonates ^57^. Furthermore, OPR3 was the sole JA biosynthetic gene present in *Draparnaldia*. Interestingly, *Draparnaldia* had the highest number of OPR3 gene copies from all of the assessed green algae, comparable in number to *Arabidopsis*. It is not clear if OPR3 is involved in JA synthesis, as JA biosynthesis can also occur via an OPR3-independent route ^58^. In contrast, JA and most of its precursors (cisOPDA, OPC-6, OPC-4) and two biosynthetic genes (AOS, OPR3) were detected in *Klebsormidium*. Both algae lacked the genes encoding for the transport (except ABCG16/JAT1 in *Klebsormidium*), signaling components, and their respective hormone receptors. Thus, neither metabolomics nor the genome supports a role of JA as signaling molecule in *Draparnaldia*.

**ABA** – only ABA was detected in both algae, the ABA biosynthetic precursors and conjugates were undetectable in our analyses. From the biosynthetic pathway, three genes (ABA4, NCED, ABA3) out of the six known in *Arabidopsis* were present in *Draparnaldia*. The complete pathway, except for ZEP, is encoded in the *Klebsormidium* genome. The potential homologues of the ABA transport mechanisms, DTX50 and ABCG25 were present in both algae. As for signaling, only the signal mediators PP2C and SnRK, out of the expected PYR1/PYL-PP2C-SnRK2-CPK-AREB/ABF, were present both in *Draparnaldia* (as previously discuss in the transcriptome changes and in *Klebsormidium*). Thus, despite ABA is present both in Streptophyte algae and Chlorophyte algae, its role and signaling mechanism remain obscure in both algal lineages.

**Brassinosteroids** – We detected three types of BRs in both algae: homo BL, DL, homo DS (*Draparnaldia*) and BL, homo CS, homo DL (*Klebsormidium*). They encode for a gene homologue of DET2 that acts in the BRs biosynthetic pathway. The homologues of the BRs transport pathway from *Arabidopsis* (ABCB19, ABCB1) were found also in *Draparnaldia*. Regarding BR signaling, (BRI1-BAK1_BKI1-BSK1-BSU1-BIN2-BES1-BZR1), the nucleocytoplasmic regulators BSU1 and BIN2 (downstream BRs signaling) were present in both algae. In addition, the receptor-like cytoplasmic kinase BSK1 (upstream BRs signaling) was present only in *Klebsormidium.* This suggests that Brassinosteroids are present and use a similar signaling mechanism in both algal lineages.

**Ethylene** and **Strigolactones** – for these hormones, measurements in algae were not performed in this study. **Ethylene**: From the two-step biosynthetic pathway, only ACS homologue is present in both *Draparnaldia* and *Klebsormidium*. From the *Arabidopsis* canonical signaling pathway (ETR-CTR1-EIN2-EBF-EIN3/EIL-ERFs), only the potential homologue of the CTR1 protein kinase was present in *Draparnaldia,* whereas *Klebsormidium* contains also the ETR receptor and the transcription factors EIN3/EIL. **Strigolactones**: from four genes of the *Arabidopsis* biosynthetic pathway, D27 was present in both algae and MAX1 only in *Klebsormidium*. A complete canonical signaling pathway (D14-MAX2-SMAX1) was lacking in both algae, only the receptor D14 was present in *Klebsormidium*. The identity of the strigolactone transporters remains obscure in the green lineage.

## Discussion

### Independent evolution of multicellularity in two green algae lineages

Both the Chlorophyte and the Streptophyte green algal lineages contain unicellular– and multicellular species. Among them, the unicellular-multicellular transition in volvocine algae (Chlorophyte algae) has been the focus of most of the research on the evolution of multicellularity. Surprisingly, a comparison of the unicellular *Chlamydomonas* and the colonial *Volvox* revealed that their genomes are remarkable similar. It seems that the colonial morphology of *Volvox* could be explained by only very few changes such as the cell-cell adhesion of ECM proteins that are enriched in *Volvox* ^59^. Here, we provide additional knowledge on the origin/evolution of multicellularity in chlorophyte green algae by comparing the genome of the multicellular filamentous *Draparnaldia* with the genome of unicellular *Chlamydomonas*. (1) We found a number of expanded orthogroups that could potentially explain the multicellularity and developmental complexity in *Draparnaldia* with respect to *Chlamydomonas.* (2) We found that many developmental processes enriched in *Draparnaldia* were conserved in Streptophyte algae and land plants. A similar trend was also observed by Nelson et al. 2024 ^60^ between marine Chlorophyte algae and land plants.

Altogether, our insights suggest that the major rewiring of functional networks from the last common ancestor of green plants was required to allow multicellularity in Chlorophyte algae. In fact, there are only few Chlorophyte lineages containing solely multicellular species, including the Chaetophorales with *Draparnaldia* ^15,61^. The majority of Chlorophyte lineages contain algae with different degree of morphological complexity, ranging from unicellular algae to multicellular filaments. And these lineages often include algae that within one species can switch between a unicellular stage and simple clusters of cells (= simple multicellularity) ^62–64^. This raises a question: What drives the multicellularity?

### Drivers of multicellularity in algae: Insights from *Draparnaldia*

It has been repeatedly shown that ‘simple’ multicellularity (i.e. simple clusters of cells) can be driven by environmental stresses ^65^. However, this multicellularity is often reversible as it might be beneficial for the organism to become unicellular again. To become permanently multicellular, additional genetic changes that promote division of labor between cells might be needed as suggested ^66^. Intriguingly, this concept of origin of multicellularity is remarkably similar to our findings. Here, we provide an independent example from the chlorophyte lineage. We show that multicellularity is characterized by the expansion of gene families with GO terms related to both abiotic stresses and multicellularity/development when we compare *Draparnaldia* to the unicellular *Chlamydomonas*. The fact that *Draparnaldia* shows functional expansions that could be related to multicellularity and to abiotic stresses, and at the same time it has the potential to change its morphology depending on the aquatic versus terrestrial environment (Figure 1), makes this alga a unique and powerful system to study the environmental, genetic, and epigenetic drivers of multicellular evolution in Chlorophyte algae.

### The plant terrestrialization: Insights from *Draparnaldia*

Although multicellularity evolved multiple times, the transition from water to land occurred successfully only once, specifically from the Streptophyte algae ^51,67^. What traits allowed Streptophyte algae to give rise to land plants is an open question. Nevertheless, traits such as filamentous growth, branching, rooting structures, apical growth and plasmodesmata, are being considered as key morphological features necessary for plant terrestrialization ^7^. We show that these features are also present in *Draparnaldia* (Caisová 2020 ^15^ and Figure 1). Moreover, *Draparnaldia* lives in the same habitats as many Streptophyte algae (e.g. *Klebsormidium*, *Chara*, and some Zygnematophyceae). It also possesses similar adaptations to changing environment as the Streptophyte alga *Chara* (i.e. polarity switch of rhizoids on aquatic versus dry environment ^68^) or land plants (‘transmembrane transport’ related genes upregulated in *Draparnaldia* transcriptome Agar 20 days with respect to the Liquid culture). In addition, also *Draparnaldia* phytohormones-related components are upregulated in terrestrial environment. This is significant since in land plants, phytohormones gained much more importance; especially in development and coping with biotic and abiotic stresses and therefore, it has been suggested that they may play an important role in plant terrestrialization ^69^. Despite all these shared features, *Draparnaldia* and their relatives did not manage the complete transition to land habitat and the reasons for this can be now further explored.

In summary, *Draparnaldia* represents a powerful model system that will allow us to study development of multicellularity and terrestrialization in Chlorophyte algae and compare it with the development of multicellularity and terrestrialization in streptophytes. This will help us to understand how the water-to-land plant transition happened and what are the traits unique for streptophytes that allowed them to give a rise to land plants.

## Acknowledgements

We thank Fyodor Kondrashov for his critical advice on sequencing strategies, Rodrigo Redondo for his help with the HMW DNA extraction, Michaela Mrvková for technical assistance with the phytohormone analyses, Karolina Fučíková and Fabio Rindi for their advice curating the taxonomic classifications, Josef Mravec for his comments on cellulase, Elizabeth Hollwey and Cláudia Ribeiro for helpful discussions. Financial support was provided by the Institute of Science and Technology Austria (IC1023IPC03 to LC, JF and IC1031IPC01 to BV); the Vetenskapsrådet (grant number 2020-06424 to MSTA), Internal Grant Agency of Palacky University (IGA_PrF_2024_013 to IP). Institutional support to CNAG (to MG, TSA, JGG, MD, AEC) was provided by the Spanish Ministry of Science and Innovation through the Instituto de Salud Carlos III, and by the Generalitat de Catalunya through the Departament de Salut and the Departament de Recerca i Universitats.

## Author contributions

B.V., J.F., L.C. conceptualized the study. A.E.C., A.P., E.C., L.C., I.P., J.G.G., M.G., T.S.A. provided resources and materials. M.G. prepared libraries and sequenced genome and transcriptomes. J.G.G., T.S.A. generated the draft genome. A.E.C., A.P., B.V., E.C., I.P., J.F., J.G.G., L.C., M.G., M.D., M.S.T.A., O.N., T.S.A., Y.V.de P. analyzed data. I.P., A.P., O.N. performed and interpreted measurements of phytohormones. B.V., J.F., L.C., M.S.T.A. wrote the manuscript with input from all of the authors.

## Supplemental figures

**Figure S1. Related to Figure 1. Morphology of Draparnaldia growing in Liquid culture and on Agar. A-G** *Draparnaldia* growing in Liquid culture. **A-C** 1, 2 and 4 days old germlings, respectively. **D-E** 5 and 8 days old young filaments. **F-G** 11 days old adult plant. H-J. 20 days old *Draparnaldia* growing on agar. **K-N** 40 days old *Draparnaldia* growing on agar. **K-L** Macroscoscopic view of the Prostrate system (PS). **M-N** Detail view of the Prostrate System at 50 μm magnification where chloroplast can be seen enlarged in size (**M**) and cells become rounded (**N**). Abbreviations. US: Upright System, PS: Prostrate System, M: mucilage.

**Figure S2. Related to Figure 2A. Workflow and statistics of the genome assembly pipeline. A** CLAWS workflow rule graph. **B** K-mer copy number spectrum produced by Merqury is a stacked k-mer distribution broken down by the number of copies of distinct k-mers from the read set present in the assembled genome. The number of k-mers present in the assembly but not in the reads is indicated by the bar graph at the origin of the graph. **C** Cumulative length of scaffolds categorized by BLAST hits to major t. **D** Workflow of the genome annotation process.

**Figure S3. Related to Figure 3. Twenty four orthogroups expanded in *Draparnaldia* and not in *Chlamydomonas***. Phylogenetic trees of significantly expanded orthogroups in the multicellular chlorophyte *Draparnaldia* with respect to the unicellular chlorophyte *Chlamydomonas*.

**Figure S4. Related to Figure 6. Phylogenetic gene tree of the enzymes involved in the ethylene and abscisic acid signaling pathways in the green lineage**. **A** Phylogenetic placement of *Draparnaldia* ethylene response transcritpion factors (ERFs) with respect to the *Arabidopsis* ERFs. **B** Gene tree of ABA phosphatases in the land plants *Arabidopsis thaliana*, *Oryza sativa*, *Marchantia polymorpha*, *Physcomitrium patens*, and in the green algae *Draparnaldia*, *Auxenochlorella protothecoides*, *Picochlorum SENEW*, *Coccomyxa obi*, *Chlamydomonas reinhardtii*, *Volvox carteri*, *Selaginella moellendorffii*, *Klebsormidium nitens*, *Spirogloea muscicola*, and *Mesotaenium endlicherianum*.

## Supplemental Tables

**Table S1. Related to Figure 1. Summary of genome statistics of the species used for this study.**

**Table S2a. Related to Figure 2A. Statistics for all assemblies.**

**Table S2b. Related to Figure 2A. Assembly pipeline configuration.**

**Table S2c. Related to Figure 2A. DRAP1A annotation statistics.**

**Table S3. Related to Figure 2B. Overlap of orthogroups between *Draparnaldia*, streptophyte algae, and land plants**.

**Table S4a. Related to Figure 3. Enriched GO terms in orthogroups expanded in *Draparnaldia* with respect to *Chlamydomonas*.**

**Table S4b. Related to Figure 3. Enriched Pfam domains in orthogroups expanded in *Draparnaldia* with respect to *Chlamydomonas***.

**Table S4c. Related to Figure 3. Enriched GO terms in orthogroups contracted in *Draparnaldia* with respect to *Chlamydomonas*. GO terms were not significant**.

**Table S4d. Related to Figure 3. Enriched GO terms in orthogroups expanded in *Chara* with respect to *Mesostigma***.

**Table S4e. Related to Figure 3. Enriched Pfam domains in orthogroups expanded in *Chara* with respect to *Mesostigma***.

**Table S4f. Related to Figure 3. Enriched GO terms in orthogroups contracted in *Chara* with respect to *Mesostigma***.

**Table S4g. Related to Figure 3. Enriched Pfam domains in orthogroups contracted in *Chara* with respect to *Mesostigma***.

**Table S5a. Related to Figure 4. DEG agar_culture_20_days_vs_liquid_culture.**

**Table S5b. Related to Figure 4. DEG agar_culture_40_days_vs_agar_culture_20_days.**

**Table S5c. Related to Figure 4. DEG agar_culture_40_days_vs_liquid_culture.**

**Table S5d. Related to Figure 4. GO enrichment of downregulated genes in agar_culture_20_days_vs_liquid_culture.**

**Table S5e. Related to Figure 4. GO enrichment of upregulated genes in agar_culture_20_days_vs_liquid_culture.**

**Table S5f. Related to Figure 4. GO enrichment of downregulated genes in agar_culture_40_days_vs_agar_culture_20_days.**

**Table S5g. Related to Figure 4. GO enrichment of upregulated genes in agar_culture_40_days_vs_agar_culture_20_days.**

**Table S6. Related to Figure 5. Correspondence between the gene name used of the protein-protein interaction network and the respective Gene IDs in *Draparnaldia* and *Arabidopsis***.

**Table S7. Related to Figure 6 and Figure 7. Absolute count numbers of conserved transcription factors among land plants and algae, including *Draparnaldia***.

**Table S8a. Related to Figure 7. Detection statistics of the cytokinins both in the alga and in the media from *Draparnaldia* and *Klebsormidium nitens***.

**Table S8b. Related to Figure 7. Detection statistics of the 2MeS cytokinins both in the alga and in the media from *Draparnaldia* and *Klebsormidium nitens***.

**Table S8c. Related to Figure 7. Detection statistics of the jasmonates both in the alga and in the media from *Draparnaldia* and *Klebsormidium nitens***.

**Table S8d. Related to Figure 7. Detection statistics of the abscisates both in the alga and in the media from *Draparnaldia* and *Klebsormidium nitens***.

**Table S8e. Related to Figure 7. Detection statistics of the gibberellins both in the alga and in the media from *Draparnaldia* and *Klebsormidium nitens***.

**Table S8f. Related to Figure 7. Detection statistics of the brassinosteroids both in the alga and in the media from *Draparnaldia* and *Klebsormidium nitens***.

**Table S8g. Related to Figure 7. Four biological replicates for the detection of auxin in two independent samples of *Klebsormidium nitens* in medium with or without the algae**.

**Table S8h. Related to Figure 7. Auxin metabolites detected in *Klebsormidium nitens* samples in medium with or without the algae**.

**Table S8i. Related to Figure 7. Four biological replicates for the detection of auxin in two independent samples of *Draparnaldia* in medium with or without the algae**.

**Table S8j. Related to Figure 7. Four biological replicates for the detection of auxin in two independent samples of *Draparnaldia* in medium with or without the algae**.

**Table S8k. Related to Figure 7. List of compounds for the hormonomics detection.**

**Table S8l. Related to Figure 7. Gene copies encoding members of the synthesis, signaling and transport networks of the phytohormones in the different green species used in this study.**

## Supplemental Notes

**Note S1. Related to Figure 1. Taxonomic acts for *Draparnaldia***

## STAR Methods

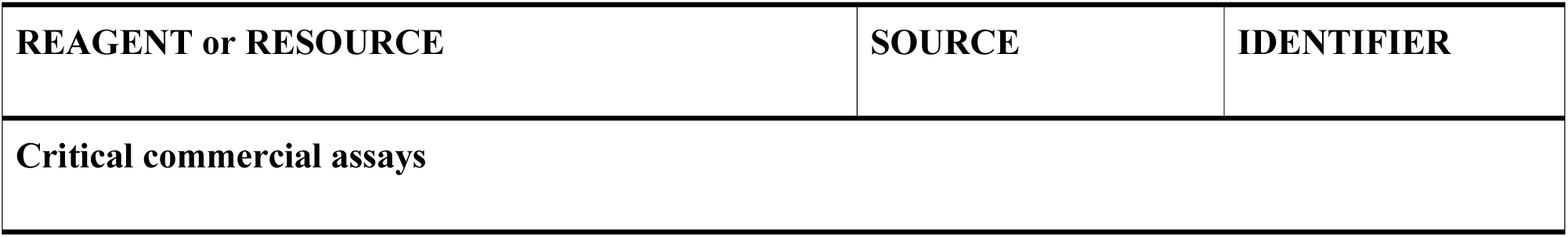
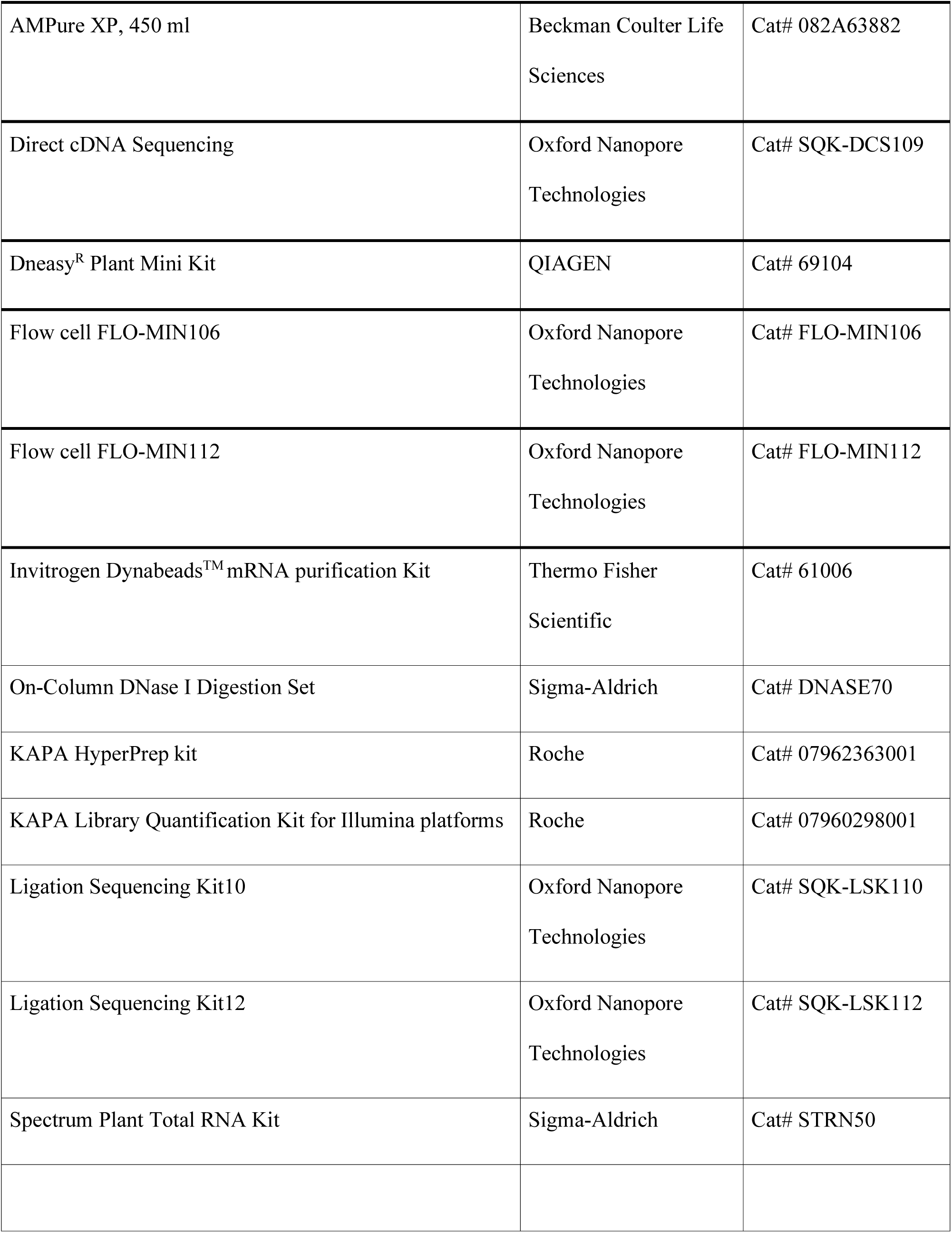
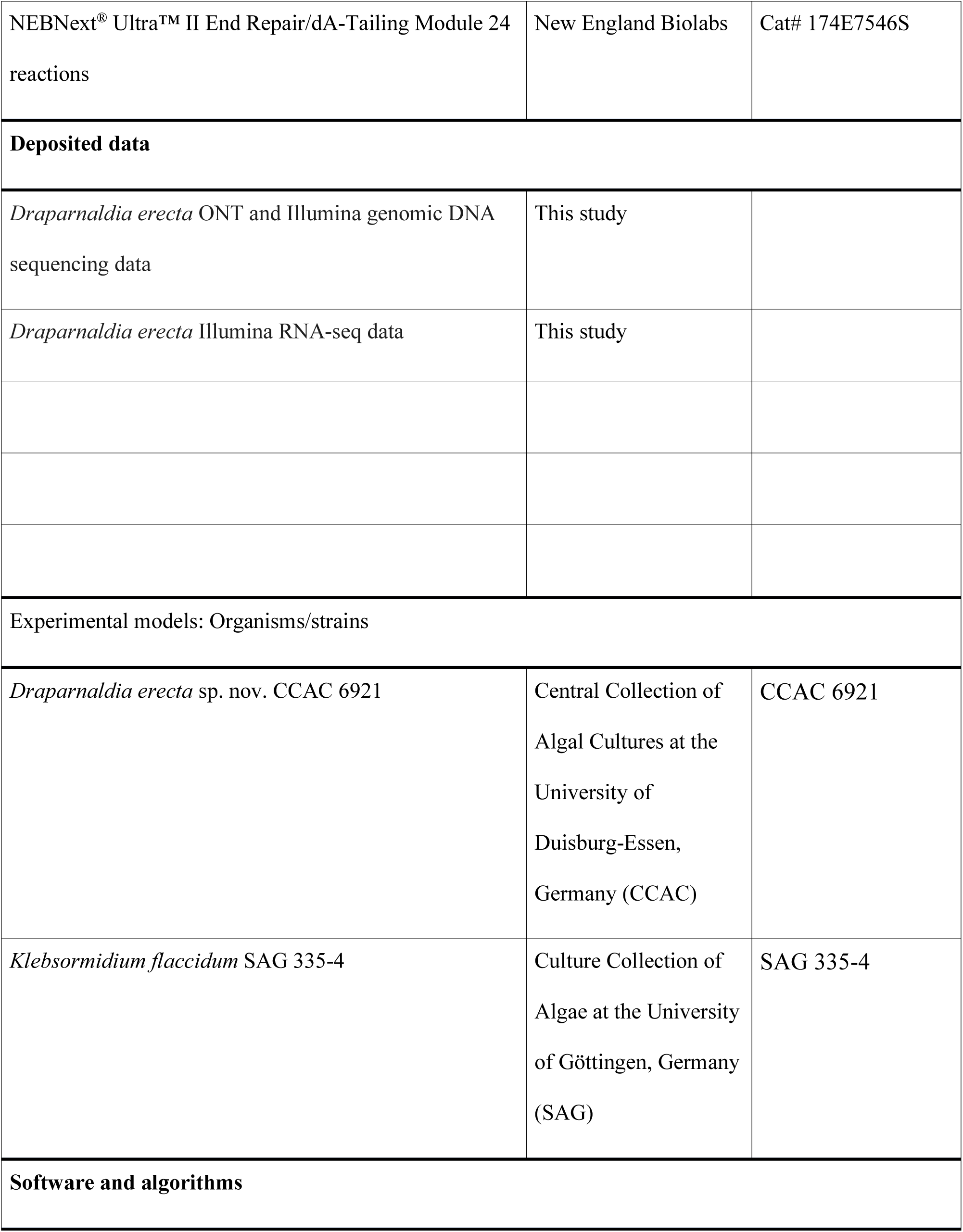
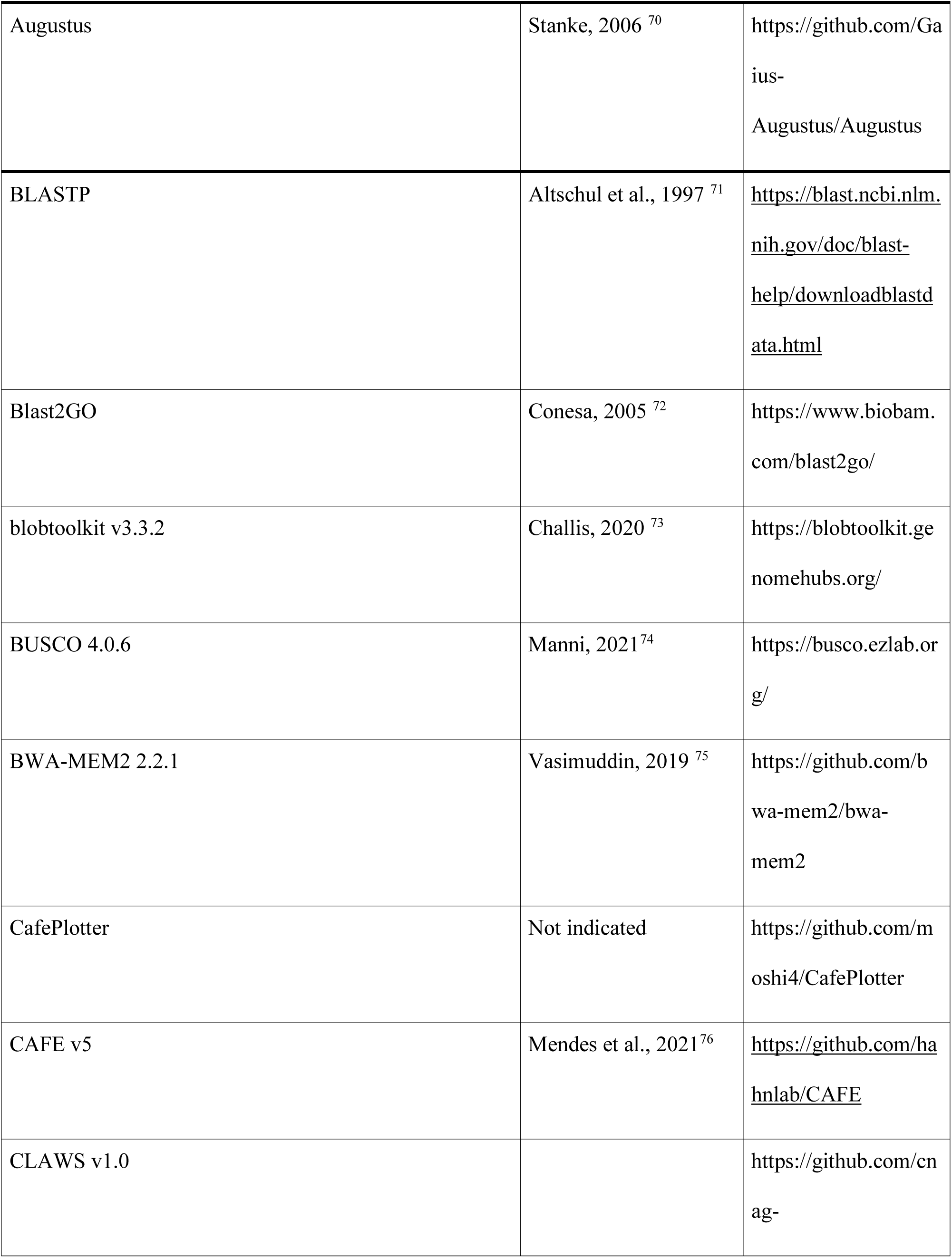
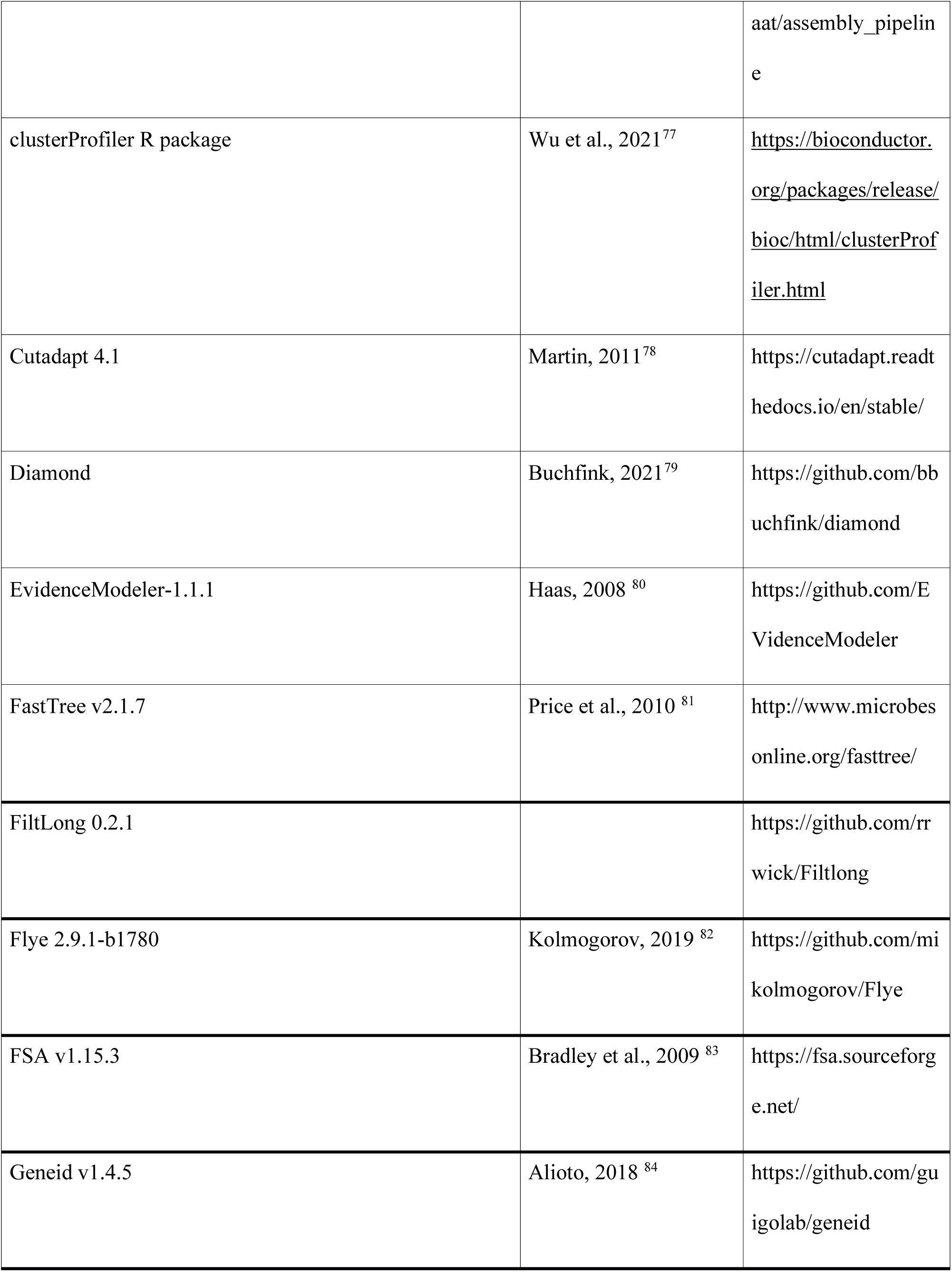
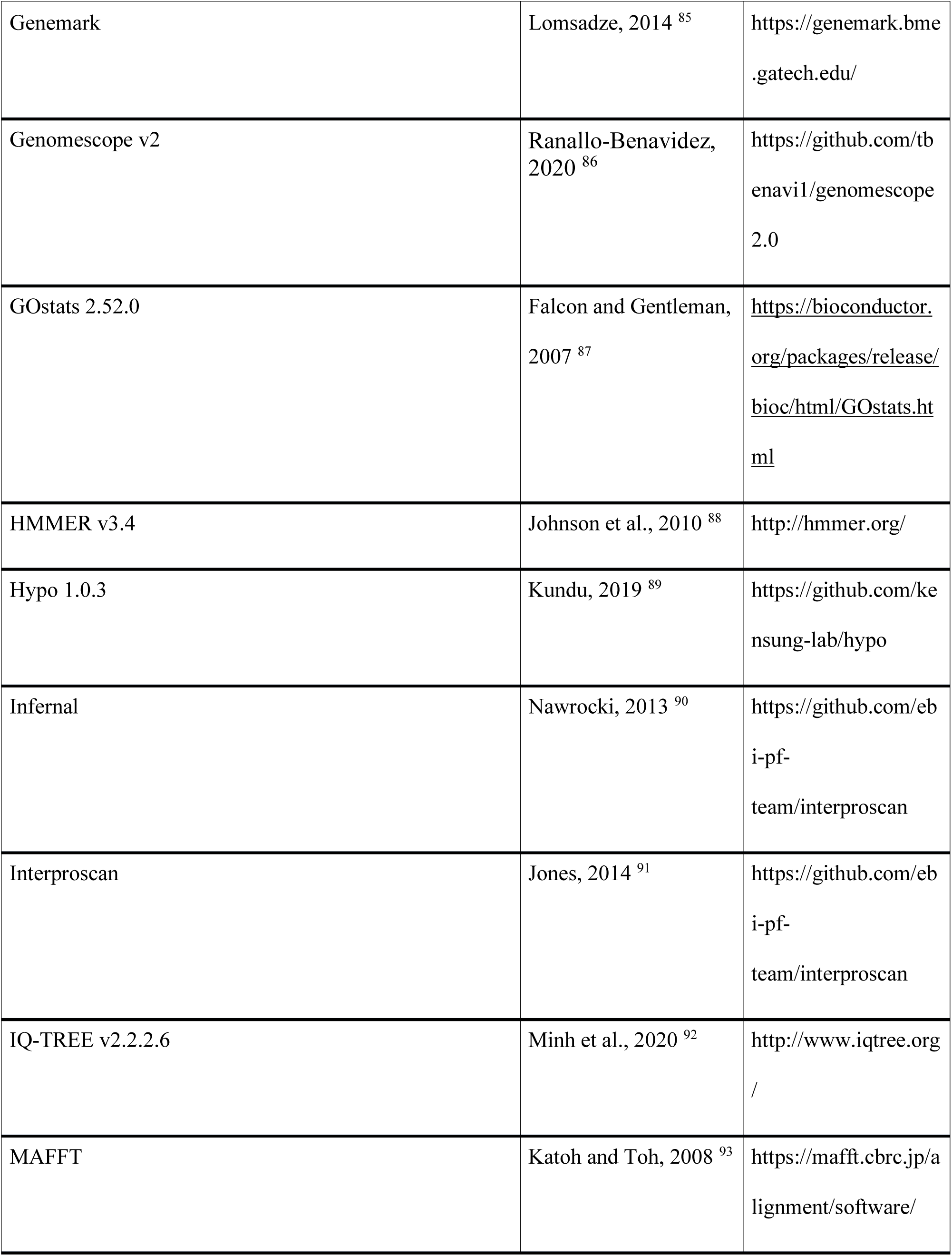
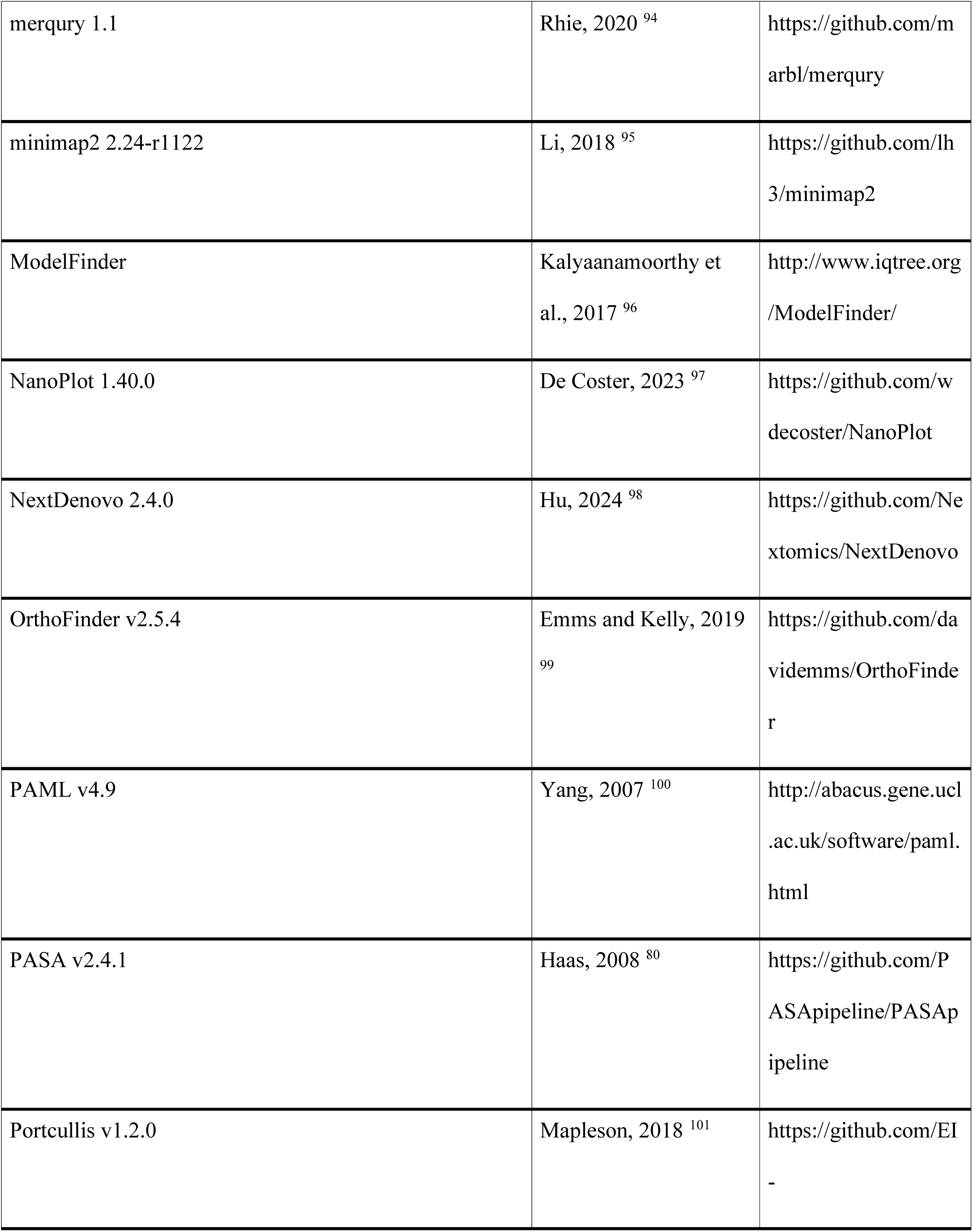
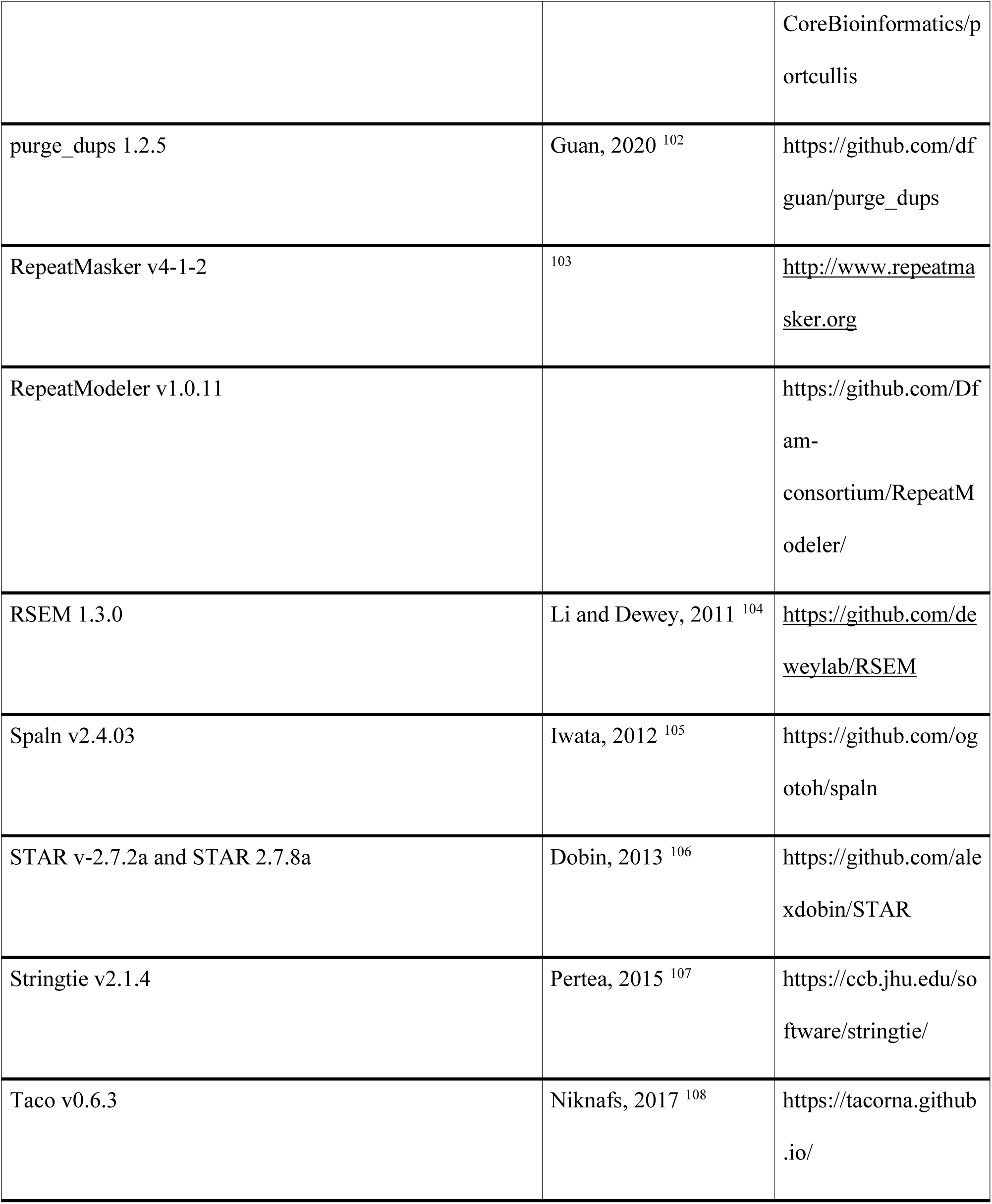
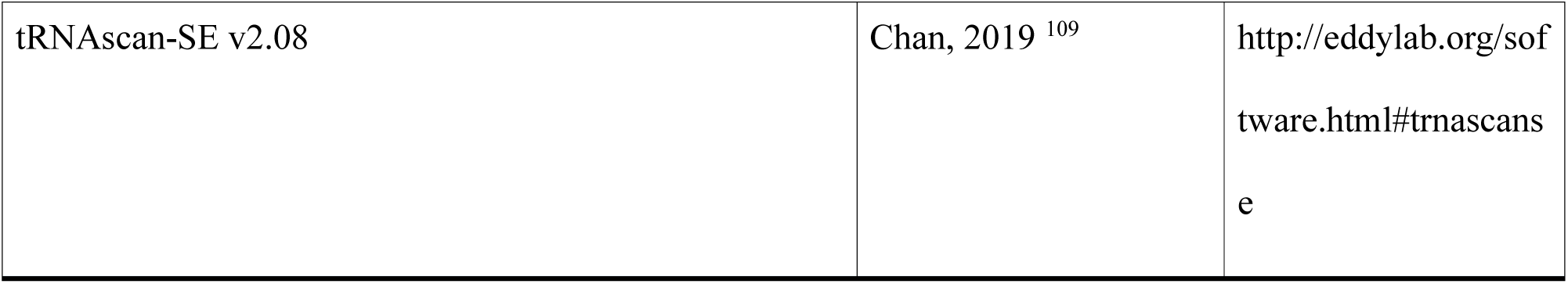
Key resources table.

### Contact for Reagent and Resource Sharing

Further information and requests for resources and reagents should be directed to and will be fulfilled by the lead contact, Jiří Friml.

## Method Details

### Biomass growth and harvest for DNA and RNA extraction

The standart maintanace, growth conditions and propagation of *Draparnaldia* was described in Caisová and Jobe 2019 ^16^.

For this study, biomass grown in three different conditions was used: Liquid culture, Agar 20 days and Agar 40 days. Liquid culture (Figure 1C) was inoculated by 50 mL of a stock culture and grown for seven days in the aerated flask, for details see Caisová and Jobe 2019. After seven days the culture contains all stages of *Draparnaldia’s* life cycle shown in Figure 1D.

Agar 20 days (Figure 1E left): harvested biomass from Liquid culture was inoculated on 1% agar plates and grown for 20 days. Agar 40 days (Figure 1E right): Agar 20 days was grown additional 20 days.

Biomass from Liquid culture was harvested using a cells trainer (40 μm). Biomass growing on Agar 20 days and Agar 40 days was harvested using spatula. To remove rest of media, biomass harvested from Liquid culture was washed with sterile deionized water. The harvested biomass was squeeze-dry using the filter paper, weighted and 50-60 mg of biomass was transferred into 1.5 mL (for gDNA and RNA extractions) and 2 mL (for HMW gDNA extraction) Eppendorf tubes. Tubes with biomass were freezed in liquid nitrogen and stored at –70 degree freezer. Before extraction, biomass was grained in tubes by using glass stick with a rough surface. All extractions were performed directly in 1.5 and 2 mL tubes.

### HMW gDNA extraction

Biomass from Liquid culture was used for HMW gDNA extraction using a modified CTAB protocol. The following steps were applied:

1. add a mixture of 180 µL TE buffer, 140 µL 5% SDS and 5 µL proteinase K solution to the grained frozen biomass in 2 mL tube, invert the tube (slowly, several times)
2. incubation @ 65°C, 20‘, during this time invert the tube 3-4 times
3. add 100 µL 5 M NaCl and 1 mL CTAB buffer (pre-warmed @65°C)
4. incubation @ 60°C, 25‘, invert the tube 3-4 times
5. centrifuge (5,000 g, 10‘) & transfer supernatant into new 2 ml Eppendorf tubes (max. 800 µL per tube)
6. add equal volume of 24:1 chloroform: isoamyl alcohol (800uL per epi)
7. incubation for 10 min at platform rocker.
8. separate phases by centrifugation (7,000 g, 5’)
9. transfer the upper, aqueous phase carefully into a new 2mL Eppendorf tube
10. add 10 µL RNAse A (10 mg/mL) and incubate at 37 °C for 25‘
11. repeat steps 6 – 8.
12. transfer the upper, aqueous phase carefully into a new 2mL Eppendorf tube
13. add 10% of 3M sodium acetate (pH 5.2, filtered 0.22), 2/3 (0.6-0.7) volume(s) of RT isopropanol. This means for 600 µL of the upper phase add 60 µL sodium acetate (no mixing) and 1000 µL of isopropanol.
14. carefully invert by hand, several times –> white filaments will be produced and they will clock together
15. centrifuge (3,000 g for 5’) and pour the supernatant into a new 2 mL Eppendorf tube
16. wash pellet with 1 mL 70% RT ethanol by carefully inverting few times the Eppendorf tube.
17. centrifuge (3,000 g for 5’) & discard the supernatant carefully
18. repeat the washing step
19. air-dry the pellet at the bench (for 10-15 min – until there are no drops on the walls of Eppendorf tube),
20. dissolve the pellet in 50 – 200 µl Buffer EB (from Qiagen). This is 10mM Tris-Cl, pH 8.5. Keep the dissolved HMW gDNA for 1-2 hour at the room temperature and then at 4 degrees overnight. After that measure the concentration and OD.
21. Store it at 4 degrees

Useful notes: 1) Room Temperature Isopropanol has to be used. Cold isopropanol will break long fragments of DNA and all salts (and other contaminates) will be accumulated with DNA. 2) The same holds true for 70% Ethanol (washing step). 3) To collect DNA, a low speed (3,000 g) centrifugation needs to be applied. Low speed will allow to collect mostly long and clean DNA fragments. 4) 70% Ethanol for washing step has to be prepared freshly. 5) Perform all step with centrifugation and washing at the room temperature. 6) To avoid breaking DNA filaments, use tips with a broad opening and do not vortex your samples.

gDNA extraction for Illumina sequencing was done from Liquid culture. DNA was extracted using Dneasy^R^ Plant Mini Kit (QIAGEN, Catalog No 69104) by standard protocol. Only additional two washes with AW2 were performed between steps 9 and 10 to remove chlorophyll.

## RNA extraction

Total RNA was extracted from three different conditions (Liquid culture, Agar 20 days and Agar 40 days) in three biological replicates. Extraction was performed using the Spectrum Plant Total RNA Kit (Sigma-Aldrich, Catalog No STRN50). Official protocol was followed with a ‘protocol A’ in Step 4 Bind RNA to Column, including the optional on-column Dnase Digestion step. In On column Dnase Digestion, firstly, washing with Wash Solution I was done (step 1 in the protocol). Secondly, additional two washes were performed with Wash Solution 2 to remove chlorophyll of the alga. After that, a digestion mixture was applied (step 3 in the protocol). Other steps remain the same.

Poly A RNA was extracted from tree conditions (see above), in one biological replicate. This extraction was done in two steps: Firstly, total RNA was isolated as described above. Secondly, poly A RNA was pull down from total RNA using the Invitrogen Dynabeads^TM^ mRNA purification Kit (Catalog No. 61006). All steps in the protocol were followed, no modifications were done.

## Long-Read Whole Genome Library Preparation and Sequencing

HMW gDNA extracted from *Draparnaldia* was quality-controlled for long-read sequencing, considering purity, quantity, and integrity. The sequencing libraries were prepared using the ligation 1D sequencing kit 10 or upgraded kit 12 chemistry (SQK-LSK110 kit SQK-LSK112, respectively) from Oxford Nanopore Technologies (ONT). In brief, 3.0 μg of the gDNA underwent end-repair and adenylation using the NEBNext UltraII End Repair/dA-Tailing Module (NEB), followed by ligation of sequencing adaptors. The ligation product was purified using 0.4X AMPure XP Beads and eluted in Elution Buffer (ONT).

The WGS long-read sequencing run was performed on a GridIon Mk1 (ONT) using a flow cell R9.4.1 (FLO-MIN106) compatible with sequencing ligation kit 10 or flow cell R10.4 (FLO-MIN112) compatible with sequencing ligation kit 12. The sequencing data was collected for 72 hours. The quality parameters of the sequencing runs were monitored in real time using the MinKNOW platform version 21.10.8, resp. 21.11.7, and the basecalling was performed using Guppy version 5.1.17, resp. 5.1.13.

## Long-Read Direct cDNA Library Preparation and Sequencing

Enriched polyA-RNA fraction from different conditions (Liquid culture, Agar 20 days and Agar 40 days) was used for the direct cDNA library preparation protocol using the Direct cDNA Sequencing Kit (SQK-DCS109) from ONT, following the manufacturer’s instructions.

The sequencing runs were performed on GridION Mk1 (ONT) using a flow cell R9.4.1 FLO-MIN106D (ONT), and the sequencing data were collected for 72 hours. The quality parameters of the sequencing runs were monitored in real time by the MinKNOW platform version 21.11.7, and the basecalling was performed with Guppy version 5.1.13.

## Short-Read: Whole Genome Sequencing (WGS) Library Preparation

The short-insert paired-end library for whole genome sequencing was prepared using the PCR-free protocol and the KAPA HyperPrep kit (Roche). After end-repair and adenylation, Illumina platform-compatible adaptors with unique dual indexes and unique molecular identifiers (Integrated DNA Technologies) were ligated. The sequencing libraries were quality controlled on an Agilent 2100 Bioanalyzer using the DNA 7500 assay (Agilent) to assess size and quantified using the Kapa Library Quantification Kit for Illumina platforms (Roche).

## Short-Read: RNASeq Library Preparation

Total RNA from different conditions (Liquid culture, Agar 20 days and Agar 40 days) was used to prepare the RNASeq libraries. The KAPA Stranded mRNA-Seq Illumina Platforms Kit (Roche) was used, following the manufacturer’s recommendations, with 500 ng of total RNA as the input material. Library quality was assessed on an Agilent 2100 Bioanalyzer using the DNA 7500 assay.

## Short-Read Sequencing

The RNASeq and WGS libraries were sequenced on Illumina NovaSeq 6000 with a read length of 2×101bp or 2×151bp respectively, following the manufacturer’s protocol for dual indexing. Image analysis, base calling, and quality scoring of the run were processed using the manufacturer’s software, Real Time Analysis (RTA 3.4.4).

## Nuclear Genome Assembly

The DNA sequencing produced 14.6 Gb of ONT reads (read N50=10.4 kb) base-called with the high-accuracy model of Guppy and 24.4 Gb of Illumina PE 2×151bp reads. These data were used with the CNAG Snakemake assembly pipeline CLAWS v1.0 (https://github.com/cnag-aat/assembly_pipeline) (flowchart with complete assembly workflow shown in Figure S2A). In brief, the Illumina reads were adapter-trimmed with Cutadapt v4.1 ^78^ and the ONT reads were filtered with FiltLong v0.2.1 (https://github.com/rrwick/Filtlong), removing reads shorter than one kilobase as well as low quality reads (<80% accurate). Then the filtered ONT reads are assembled with both Flye v2.9. ^82^ and NextDenovo v2.4.0 ^98^. The assemblies were polished with both ONT and Illumina PE reads using Hypo ^89^ and haplotypic and other artificially duplicated sequence were removed using purge_dups v1.2.5 ^102^. Evaluations were performed at every step, using BUSCO v4.0.6 ^74^ with chlorophyta_odb10, Merqury v1.1 ^94^ to estimate the consensus accuracy (QV) and k-mer statistics, and fasta-stats.py to generate contiguity statistics. Our best assembly, was obtained with Flye (see assembly metrics in Table S2). First, the filtered ONT reads (FiltLong) longer than 1 kb were assembled with Flye. BlobToolKit v3.3.2 ^73^ was used to detect possible contamination and produce figures. The specific parameters and versions used to assemble this sample are also listed in Table S2. Possible contamination in the assembly was assessed by BLAST using BlobTools. Most sequence is classified as Chlorophyta (Figure S2C). Potential contamination indicated by hits to other taxa were manually confirmed as algal sequences after protein-coding gene annotation and remain in the assembly.

## Genome annotation

Repeats present in the *Draparnaldia* genome assembly were annotated with RepeatMasker v4-1-2 (http://www.repeatmasker.org) using the custom repeat library available for “algae”. Moreover, a new repeat library specific for our assembly was made with RepeatModeler v1.0.11. After excluding those repeats that were part of repetitive protein families (performing a BLAST ^110^ search against Uniprot) from the resulting library, RepeatMasker was run again with this new library in order to annotate the specific repeats.

The gene annotation of the *Draparnaldia* genome assembly was obtained by combining transcript alignments, protein alignments and *ab initio* gene predictions. A flowchart of the annotation process is shown in Figure S2D.

Firstly, RNA from three different conditions was obtained and sequenced with both Illumina RNAseq and ONT direct cDNAseq. After sequencing, the short and long reads were aligned to the genome using, respectively, STAR ^106^ v-2.7.2a and MINIMAP2 ^95^ v2.14 with the splice option. Transcript models were subsequently generated using Stringtie ^107^ v2.1.4 on each BAM file and then all the models produced were combined using TACO ^108^ v0.6.3. High-quality junctions to be used during the annotation process were obtained by running Portcullis ^101^ v1.2.0 after mapping with STAR and MINIMAP2. Finally, PASA assemblies were produced with PASA ^80^ v2.4.1. The *TransDecoder* program, which is part of the PASA package, was run on the PASA assemblies to detect coding regions in the transcripts. Secondly, the complete proteomes of *Chlamydomonas reinhardtii, Ostreococus tauri* and *Micromonas commoda* were downloaded from Uniprot in July 2022 and aligned to the genome using Spaln ^105^ v2.4.03. *Ab initio* gene predictions were performed on the repeat-masked genome assembly with three different programs: GeneID ^84^ v1.4, Augustus ^70^ v3.3.4 and Genemark-ES ^85^ v2.3e with and without incorporating evidence from the RNAseq data. The protein models previously produced by Transdecoder on the PASA assemblies were filtered by length and match to the protein and long read evidence. The resulting models were used to train Geneid and Augustus before gene prediction. Genemark, which runs in a self-trained mode, was not previously trained. Finally, all the data were combined into consensus CDS models using EvidenceModeler-1.1.1 (EVM) ^80^. Additionally, UTRs and alternative splicing forms were annotated via two rounds of PASA annotation updates. Functional annotation was performed on the annotated proteins with Blast2go ^72^. First, a Diamond Blastp ^79^ search was made against the nr database (last accessed August 2022). Furthermore, Interproscan ^91^ was run to detect protein domains on the annotated proteins. All these data were combined by Blast2go, which produced the final functional annotation results.

The annotation of ncRNAs was obtained by running the following steps. First, the program cmsearch ^111^ v1.1 that is part of the Infernal ^90^ package was run against the RFAM database of RNA families ^112^ v12.0. Additionally, tRNAscan-SE ^109^ v2.08 was run in order to detect the transfer RNA genes present in the genome assembly. Identification of lncRNAs was done by first filtering the set of PASA-assemblies that had not been included in the annotation of protein-coding genes to retain those longer than 200bp and not covered more than 80% by a small ncRNA. The resulting transcripts were clustered into genes using shared splice sites or significant sequence overlap as criteria for designation as the same gene.

## Orthogroup and phylogenetic tree inference

To conduct phylogenetic and comparative genomic analyses for the *Draparnaldia* genome, a set of twenty genomes of plant and algal species was selected (Table S1). This selection included 9 chlorophyte algae (*Chlamydomonas reinhardtii*, *Volvox carteri*, *Picochlorum* SENEW3, *Auxenochlorella protothecoides*, *Coccomyxa obi*, *Chloropicon primus*, *Micromonas* RCC299, *Ostreococcus tauri* and *Bathycoccus prasinos*), 6 streptophyte algae (*Mesotaenium endlicherianum*, *Spirogloea muscicola*, *Chara braunii*, *Klebsormidium nitens*, *Chlorokybus atmophyticus* and *Mesostigma viride*), and 5 land plants (*Oryza sativa*, *Arabidopsis thaliana*, *Selaginella moellendorfii*, *Physcomitrium patens* and *Marchantia polymorpha*). Orthogroups across these species were inferred using OrthoFinder v2.5.4 ^99^ employing a Markov Cluster Inflation (MCI) parameter of 3 and using multiple sequence alignments inferred with MAFFT ^93^ to generate gene trees (option –M msa). Subsequently, the species tree alignment from the OrthoFinder output was used to infer a phylogenetic tree with IQ-TREE v2.2.2.6 ^92^. Hereby, 1000 ultrafast bootstrap replicates were obtained and ModelFinder ^96^ was utilized to select the best fitting substitution model. A fossil-calibrated ultrametric tree was obtained using MCMCTREE in PAML v4.9 ^100^ from concatenated multiple sequence alignments of ten low-copy orthogroups for which only one sequence was retained per species (OG0004968, 5010, 5040, 5144, 5161, 5169, 5208, 5217, 5269, 5306). Fossil data was obtained from Morris et al. (2018)^113^. MCMCTREE was run using the approximate likelihood calculation for protein data whereby 20 MCMC chains were executed, each for 40,000 generations with a burn-in of 4,000 generations. Priors were set as in Dos Reis et al. (2017) ^114^. This was repeated twice to evaluate convergence by plotting the posterior means of the different runs, plotting the infinite-site plot, and calculating Geweke’s convergence diagnostic and PSRF diagnostic. All metrics indicated convergence.

## Analysis of orthogroup expansions and contractions

Expanded and contracted orthogroups were inferred using CAFE v5 ^76^. Hereby, the orthogroups obtained with OrthoFinder and the fossil-calibrated ultrametric tree obtained with MCMCTREE were used as input (see above). First, orthogroups were filtered out if there was one or more species that had ≥ 100 gene copies, as this may lead to parameter estimates that are non-informative ICAFE was run twice with base settings (i.e. the stochastic birth-death model) to estimate a single birth-death (λ) parameter for the whole tree. Both runs converged to the same likelihood and λ parameter. Next, the estimated λ was utilized to also analyze the orthogroups that had ≥ 100 gene copies. CafePlotter (https://github.com/moshi4/CafePlotter) was utilized to visualize expansions and contractions of the orthogroups on the phylogenetic tree. Orthogroups where the probability under the stochastic birth-death model (null model) of the branch leading to *Draparnaldia* was less than 0.05, were regarded as significantly expanded/contracted. Furthermore, expanded/contracted orthogroups that were also expanded/contracted in the closely related unicellular *Chlamydomonas reinhardtii* were filtered out, as these orthogroups do not represent expansions/contractions related to the morphological complexity and multicellularity of *Draparnaldia*. In addition, significant expanded/contracted orthogroups were inferred in the multicellular streptophyte *Chara braunii* that were not expanded/contracted in the closely related unicellular *Mesostigma viride*.

## Gene Ontology/Pfam annotation and enrichment analyses for orthogroups

Pfam domains were searched in the proteomes of all selected species using the HMMSEARCH program from HMMER v3.4 ^88^. A Pfam domain was associated with a protein if the E-value was lower than 1 x 10^-3^. If a minimum of 50% of the proteins within an orthogroup possessed the Pfam domain, the orthogroup was annotated with the respective Pfam domain. GO terms were associated to Pfam domains by using the Pfam2GO mappings from the GO database. As such, an orthogroup annotated with a Pfam domain was also annotated with the GO terms of the respective Pfam domain. In addition, GO annotation data of *Draparnaldia*, *Arabidopsis thaliana*, *Micromonas* RCC299 and *Physcomitrium patens* was used to further annotate the orthogroups. GO annotation data for *A. thaliana* and *Micromonas* RCC299 were acquired from PLAZA 5.0 ^115^, while data for *P. patens* was acquired from Phytozome 13 ^116^. GO and Pfam enrichment analyses were performed in R using the clusterProfiler package ^77^. Enrichment analyses were performed for orthogroups that were expanded/contracted in *Draparnaldia* (and not in *C. reinhardtii*) and in *C. braunii* (and not in *M. viride*), and orthogroups that were only present in specific species, e.g. multicellular species. Adjusted p-values were computed utilizing the Benjamini-Hochberg (BH) method. Significant enrichment was defined as having an adjusted p-value below 0.05.

## RNA-seq quantification and differential expression

RNA-seq short reads were mapped against the in-house assembled *Draparnaldia* genome using STAR software version 2.7.8a ^106^ with ENCODE parameters. Annotated genes were quantified with RSEM v1.3.0 ^104^ using the the in-house *Draparnaldia* annotation. Differential expression analysis was performed with limma R package ^117^, using TMM normalization. The voom function ^118^ was used to transform the count data into log2-counts per million (logCPM), estimate mean-variance relationship and to compute observation-level weights. These voom-transformed counts were used to fit the linear models. Genes were considered differentially expressed (DEG) with an adjusted p-value < 0.05 and absolute fold change |FC| > 1.5. A Gene Ontology enrichment analysis was performed with the DEG using GOstats ^87^. With the aim of assessing sample similarities, a multidimensional scaling (MDS) plot was generated using the top 500 most variable genes. Heatmaps with the DEG were plotted using the voom-transformed counts and using pheatmap R package scaling by row. Volcano plots were performed with the Enhancedvolcano R Package.

To be able to visualize the expression patterns of individual genes, we estimated expression values in TPM (Transcript Per Million) for each gene from their total read count, the length of their longest transcript, and the sum of mapped read counts per sample. TPM values were quantile-normalized across samples with NormalyzerDE ^119^ before plotting.

## Genetic and protein-protein interactions

For each GO term enriched in the differential expression analysis in *Drapanaldia*, we first extracted all the genes annotated for the GO term of interest. We then obtained the set of their *A. thaliana* orthologs from the Orthofinder results (see section “Orthogroup and phylogenetic tree inference”). The final set of *A. thaliana* genes corresponding to each GO-enriched term was then directly used as input for StringDB to extract the genetic and protein-protein interactions between them.

## Quantification of plant hormones

Two algae *Draparnaldia erecta* CCAC 6921 (chlorophyte) and *Klebsormidium flaccidum* SAG 335-4 (streptophyte) were used for quantification of plant hormones. From streptophyte algae we selected *K. flaccidum* SAG 335-4 because it was axenic, contrary to *K. nitens* NIES-2285 (used in the comparative analyses of this study). The presence of unidentified bacteria in a non-axenic culture could potentially cofound the hormone detection, specifically auxin. We also did not select *Chara* (multicellular streptophyte used for identification of expansions and contractions of orthogroups), because of its slow growth made it impossible to obtain enough biomass for hormone quantification. Seven days old liquid culture of both, *D. erecta* and *K. flaccidum* was used for analyses (see above for details). The biomass was separated from media with a strainer and both biomass and media were freezed in liquid nitrogen and stored at –70 degree freezer.

The concentrations of the plant hormones in the algae were determined by the target metabolomic approach. The biomass of *Draparnaldia* and *K. flaccium* was weighed into 2 mL microtubes in three independent replicates, each per 15 mg FW. Moreover, three replicates of 0.2 mL liquid media were prepared. The samples were extracted in 60% acetonitrile (v/v) containing the stable isotopically labelled internal standard mixture. The crude extract was purified by Oasis® HLB extraction cartridges 30 mg/1 mL (Waters, Milford, CT USA). The flowthrough fraction was collected into the glass tube and evaporated to dryness in the SpeedVac concentrator (RC1010 Centrivap Jouan, ThermoFisher, USA). The samples were dissolved in 40 µl of 25% acetonitrile (v/v) and transferred into the glass insert enclosed in the liquid chromatography vial. Then, 5 µl of the sample were injected into the Acquity UPLC® CSH^TM^ C18 1.7 µm 2.1×150 mm chromatographic column (Waters, Milford, CT USA). The gradient elution and tandem mass spectrometry detection in the multiple reaction monitoring mode were performed according to the previously published methodology ^120^. The final concentrations were calculated by the isotope dilution method.

### Identification of proteins involved in phytohormone biosynthesis, transport and signaling

First, BLASTP searches were conducted for a selection of phytohormone proteins in *Draparnaldia* and the other studied species ^71^. Hits with (1) an E-value lower than 1 x 10^-10^, (2) a protein similarity higher or equal to 50% and (3) an alignment coverage higher or equal to 60% were considered candidate phytohormone proteins. Relevant protein domains were verified with HMMSEARCH from the HMMER v3.4 package ^88^. (E-value < 1 x 10^-3^) and in accordance to the specific phytohormone Pfam domains from PLAZA v5 ^115^, TAIR ^121^, InterPro ^122,123^. To resolve ambiguity of proteins with high homology but belonging to distinct phytohormone classes, multiple sequence alignments were constructed using FSA v1.15.3 ^83^, and gene trees were constructed using IQ-TREE v2.2.2.6 ^92^ (with 1000 ultra-fast bootstrap replicates) or using FastTree v2.1.7 ^81^.

### Transcription factor identification

The proteomes of *Draparnaldia* and the other studied species were scanned for HMM profiles, extracted from Wang et al. (2019) ^35^ with HMMSEARCH from the HMMER v3.4 package ^88^. Hits with an E-value lower than 1 x 10^-10^ were considered candidate transcription factors. Those were validated by identifying the required Pfam domains (E-value < 1 x 10^-5^), following the domain rules according to the TAPSCAN database ^124^.

