## Supplemental Figure S1 for "The *Draparnaldia* genome: alternative mechanisms for multicellularity and terrestrialization in green plants"

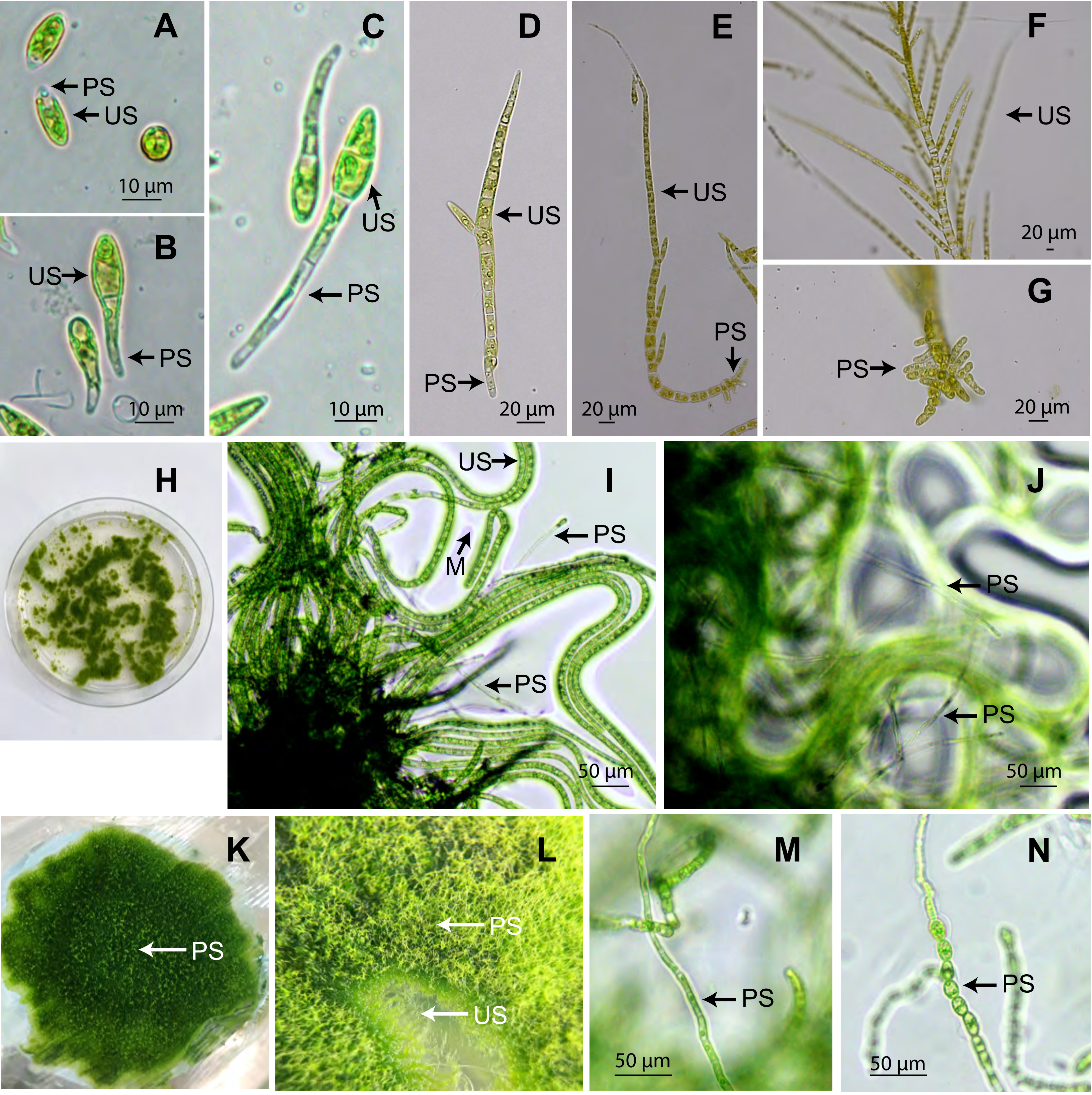


**Figure S1. Related to Figure 1. Morphology of *Draparnaldia* growing in liquid culture and on agar. A-G.** *Draparnaldia* growing in liquid culture. **A-C.** 1, 2 and 4 days old germlings, respectively. **D-E.** 5 and 8 days old young filaments. **F-G.** 11 days old adult plant. **H-J.** 20 days old *Draparnaldia* growing on agar. **K-N.** 40 days old *Draparnaldia* growing on agar. **K-L.** Macroscoscopic view of the Prostrate system (PS). **M-N.** Detail view of the Prostrate System at 50 µm magnification where chloroplast can be seen enlagerged in size (**M**) and cells become rounded (**N**). Abbreviations. US: Upright System, PS: Prostrate System, M: mucilage.
