## Supplemental Figure S2 for "The *Draparnaldia* genome: alternative mechanisms for multicellularity and terrestrialization in green plants"

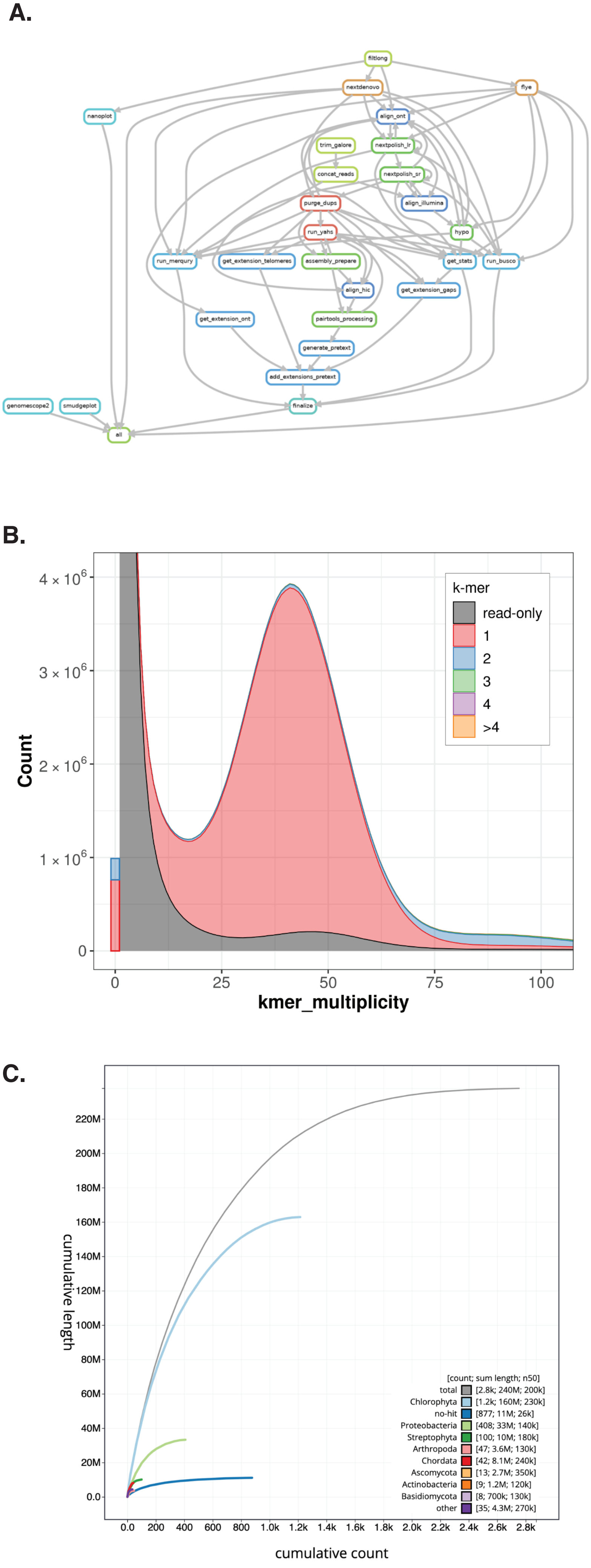


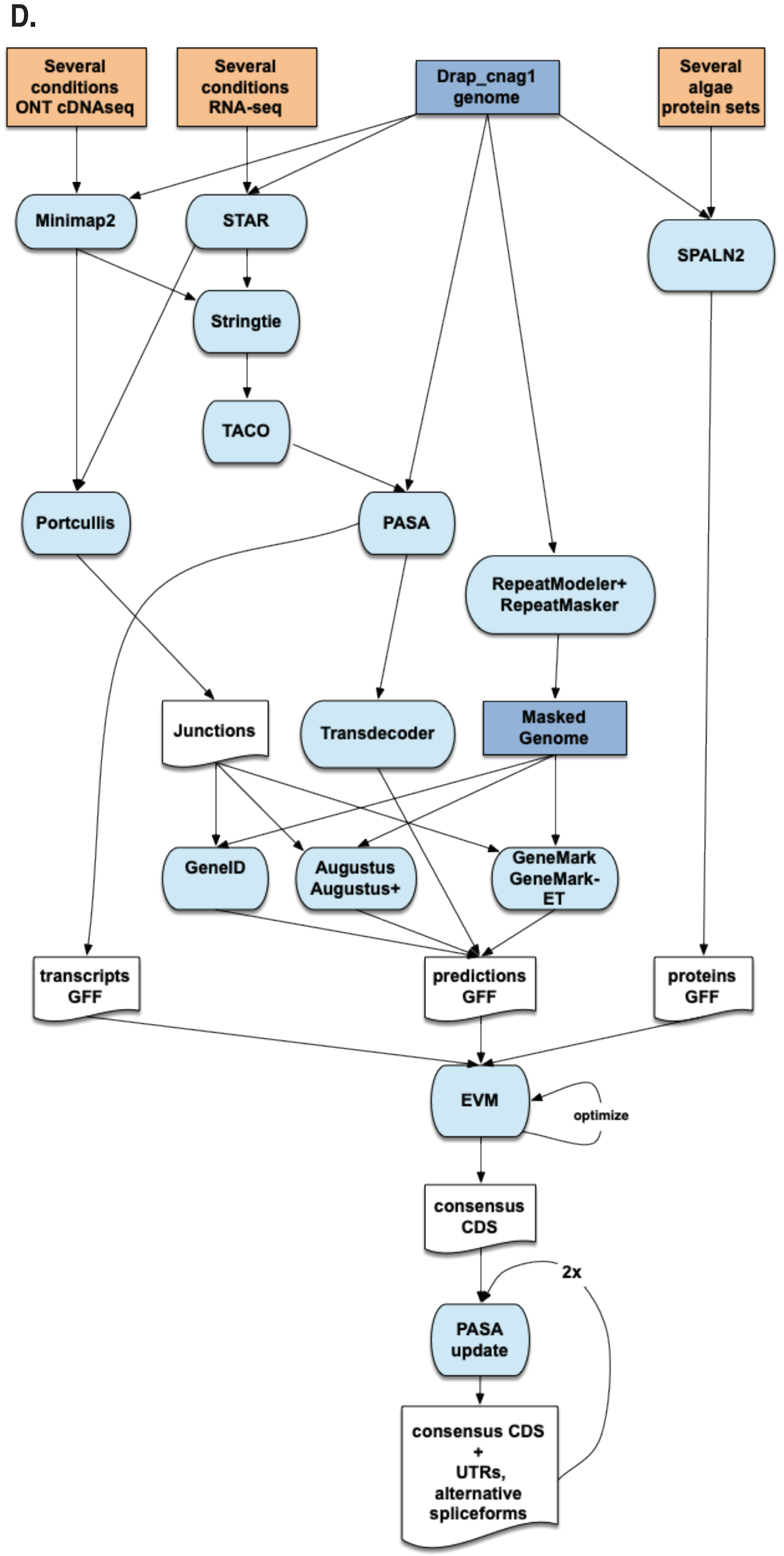


**Figure S2. Related to Figure 2A. Workflow and statistics of the genome assembly pipeline. A.** CLAWS workflow rule graph. **B.** K-mer copy number spectrum produced by Merqury is a stacked k-mer distribution broken down by the number of copies of distinct k-mers from the read set present in the assembled genome. The number of k-mers present in the assembly but not in the reads is indicated by the bar graph at the origin of the graph. **C.** Cumulative length of scaffolds categorized by BLAST hits to major t. **D.** Workflow of the genome annotation process.
