## Supplemental Figure S3 for "The *Draparnaldia* genome: alternative mechanisms for multicellularity and terrestrialization in green plants"

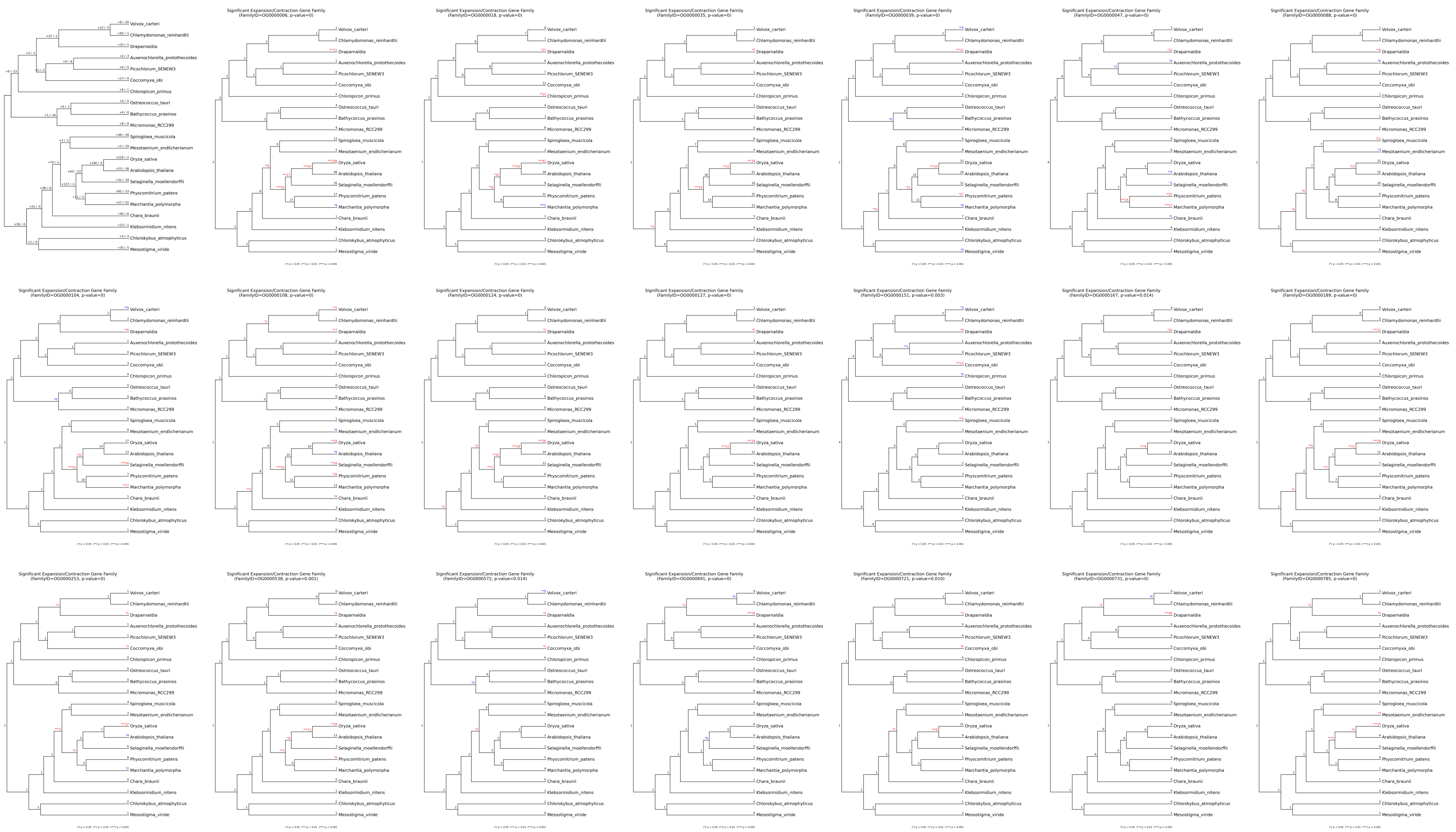

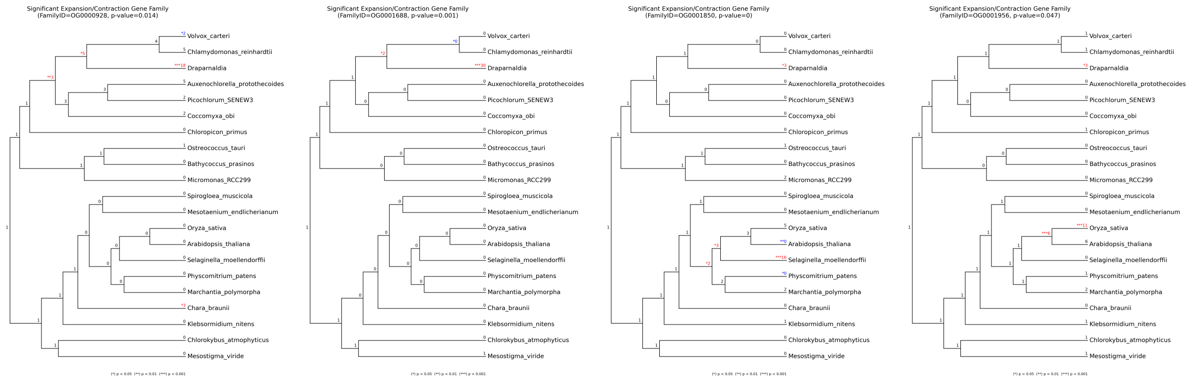

**Figure S3. Related to Figure 3. Twenty four orthogroups expanded in *Draparnaldia* and not in *Chlamydomonas*.** Phylogenetic trees of significantly expanded orthogroups in the multicellular chlorophyte *Draparnaldia* with respect to the unicellular chlorophyte *Chlamydomonas*.
