## Supplemental Figure S4 for "The *Draparnaldia* genome: alternative mechanisms for multicellularity and terrestrialization in green plants"

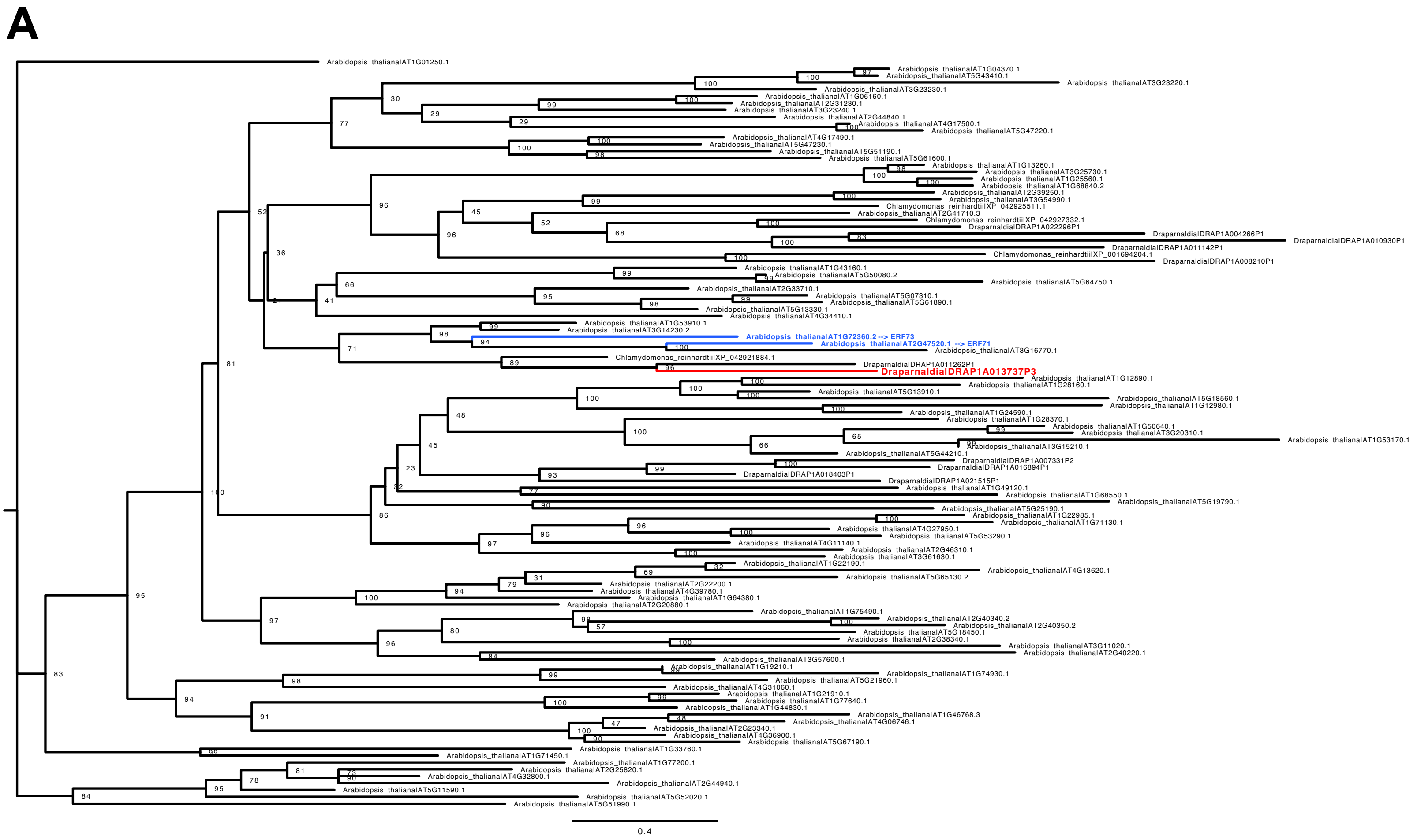


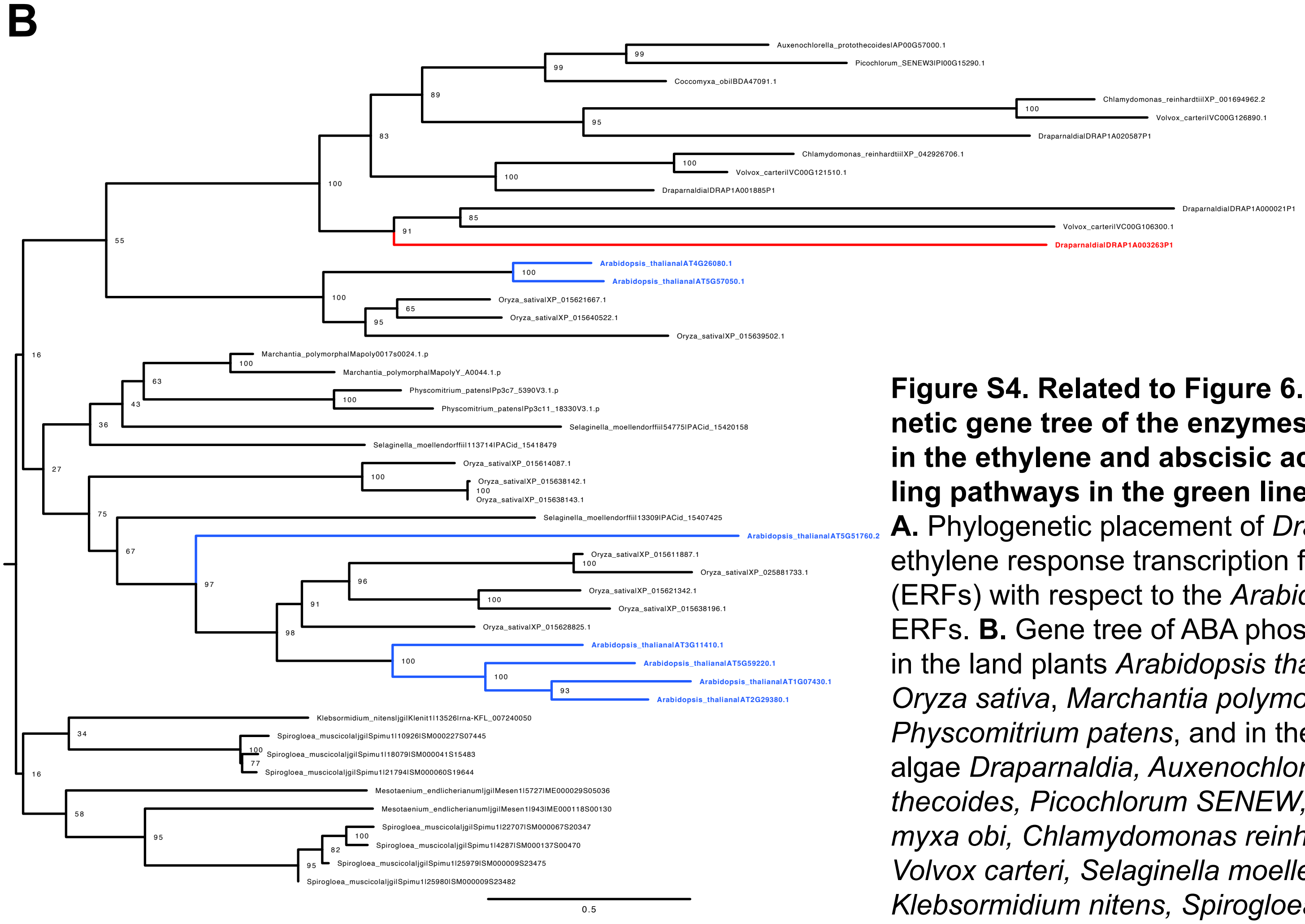


**Figure S4. Related to Figure 6. Phylogenetic gene tree of the enzymes involved in the ethylene and abscisic acid signalling pathways in the green lineage.**
**A.** Phylogenetic placement of *Draparnaldia* ethylene response transcription factors (ERFs) with respect to the *Arabidopsis* ERFs. **B.** Gene tree of ABA phosphatases in the land plants *Arabidopsis thaliana*, *Oryza sativa*, *Marchantia polymorpha*, *Physcomitrium patens*, and in the green algae *Draparnaldia, Auxenochlorella protothecoides, Picochlorum SENEW, Coccomyxa obi, Chlamydomonas reinhardtii, Volvox carteri, Selaginella moellendorffii, Klebsormidium nitens, Spirogloea muscicola, and Mesotaenium endlicherianum.*
