## Supplemental Note S1 for "The *Draparnaldia* genome: alternative mechanisms for multicellularity and terrestrialization in green plants"

**Note S1. Related to Figure 1. Taxonomic acts for *Draparnaldia***

Genus *Draparnaldia* contains 24 accepted species (Algae Base 2024, Index Nominum Algarum 2024). The *Draparnaldia* CCAC 6921 strain from which the genome has been sequenced cannot be morphologically assigned to any of the 24 species and is therefore described here as the new species: *Draparnaldia erecta* Caisová & Friml sp. nov. We also emended the description of the genus *Draparnaldia* (Roth) Bory 1808. Ann. Mus. Natl. Hist., 12: 399-409. emend. to include information on the morphology of the genus in culture and on the arrangement of the Prostrate System (= rhizoids) in water and dry environment.

***Draparnaldia* (Roth) Bory 1808. Ann. Mus. Natl. Hist., 12: 399-409. emend.**

EMENDED DIAGNOSIS: Plants consist of the Upright System (UP) and the Prostrate System (PS), both uniseriate and branched. Spatial orientation of the PS can differ in liquid and solid media in *Draparnaldia erecta*. The US consists of a main axis and branches with intercalary growth. The PS consists of rhizoids with apical growth. Branching arises directly below a cross wall (US, PS) or in the middle of a cell (PS). Branching is alternate, opposite, spiral or in fascicles, mostly primary, secondary or tertiary. US terminal cells usually bear long multicellular hairs. Cells are uninucleate, large barrel–shaped, cylindrical, rectangular (main axis in nature and occasionally in culture) or rectangular (rhizoids, branching, main axes in culture). Chloroplast is single, parietal, band-shaped with one or several pyrenoids. It is either large (US, and PS on agar) or reduced in size, making cells appear colorless (PS in liquid culture). Asexual reproduction via zoospores or aplanospores (one zoospore and aplanospore per filament cell), or via fragmentation of branches. Sexual reproduction isogamous (not observed in culture). Habitat: shallow freshwater, at least partially running waters with fluctuating water levels. Plants can be embedded in a soft mucilage in both nature and culture (agar).

LECTOTYPE SPECIES: *Draparnaldia mutabilis* (Roth) Bory 1808. Ann. Mus. Natl. Hist., 12: 399-409.
Type designated in: Hazen 1902. Mem. Torrey Bot. Club., 11: 135-250.

***Draparnaldia erecta* Caisová & Friml sp. nov.**
DIAGNOSIS: With the characters of the genus. The Prostrate System (= rhizoids) grows up when the alga is not in contact with liquid medium.

HOLOTYPE: Material of the authentic strain CCAC 6921 deposited in the xxx herbarium under xxx number.

TYPE LOCALITY: Italy, Sardinia, a bank of the river 'Rio Picocca', isolated from a leaf surface of *Salix* species.

ETYMOLOGY: The species is named after the erected Prostrate System that is typical for *Draparnaldia* when growing on agar.

AUTHENTIC STRAIN: CCAC 6921.
