## Supplemental Table S1 for "The *Draparnaldia* genome: alternative mechanisms for multicellularity and terrestrialization in green plants"

**Table S1. Related to Figure 1. Summary of genome statistics of the species used for this study**

| name | species | taxon | num_coding_genes | genome_size(Mb) | average_gene_size | median_gene_size | average_num_exons_per_gene | median_num_exons_per_gene | downloaded_from | assembly_level | publication_year | description |
|---|---|---|---|---|---|---|---|---|---|---|---|---|
| **Draparnaldia** | DRAP1A.longestpeptide | chlorophyte | 19081 | 237.9 | 5584 | 4240 | 7.59 | NA | species_of_interest | scaffold | 2022 | species of interest |
| **Galdieria sulphuraria** | Galdieria_sulphuraria | rhodophyte | 7156 | 13.7 | NA | NA | NA | NA | Pico-PLAZA3.0 | scaffold | 2013 | extremophilic red algae, HGT |
| **Cyanidioschyzon merolae 10D** | Cyanidioschyzon_merolae | rhodophyte | 4803 | 16.55 | NA | NA | NA | NA | NCBI | complete | 2007 | primitive unicellular red algae |
| **Chondrus crispus** | Chondrus_crispus | rhodophyte | 9808 | 105 | 1238 | 984 | 1.7 | 1 | JGI PhycoCosm | scaffold | 2013 | marine red algae, gene-dense regions surrounded by repeat-rich regions dominated by transposable elements. Despite its fairly large size, this genome shows features typical of compact gen |
| **Ostreococcus tauri** | Ostreococcus_tauri | chlorophyte | 7668 | 13 | NA | NA | NA | NA | Pico-PLAZA3.0 | chromosome | 2014 | smallest free-living photosynthetic eukaryote |
| **Chlamydomonas reinhardtii** | Chlamydomonas_reinhardti | chlorophyte | 19527 | 111.1 | NA | NA | NA | NA | NCBI | chromosome | 2018 | unicellular, cilia |
| **Volvox carteri** | Volvox_carteri | chlorophyte | 14971 | 138 | NA | NA | NA | NA | Pico-PLAZA3.0 | scaffold | 2010 | spherical colonies (“multicellular”) |
| **Bathycoccus prasinos** | Bathycoccus_prasinos | chlorophyte | 7826 | 15 | NA | NA | NA | NA | Pico-PLAZA3.0 | chromosome | 2012 | unicellular, marine |
| **Micromonas commoda RCC299** | Micromonas_RCC299 | chlorophyte | 10164 | 20.9 | NA | NA | NA | NA | Pico-PLAZA3.0 | complete | 2009 | motile, unicellular, global |
| **Picochlorum SENEW3** | Picochlorum_SENEW3 | chlorophyte | 6953 | 13.5 | NA | NA | NA | NA | Pico-PLAZA3.0 | scaffold | 2015 | haloterant, one of smallest and gene dense eukaryotic genomes |
| **Auxenochlorella protothecoides** | Auxenochlorella_protothecc | chlorophyte | 6810 | 22.9 | NA | NA | NA | NA | Pico-PLAZA3.0 | scaffold | 2014 | unicellular, oil bodies |
| **Coccomyxa sp. Obi** | Coccomyxa_obi | chlorophyte | 11588 | 50.41 | NA | NA | NA | NA | NCBI | chromosome | 2021 | unicellular, cold climate |
| **Chloropicon primus** | Chloropicon_primus | chlorophyte | 8627 | 17.58 | NA | NA | NA | NA | NCBI | complete | 2022 | unicellular |
| **Chara braunii S276** | Chara_braunii | streptophyte | 35672 | 1700 | 16622 | 4283 | 3.61 | 2 | JGI PhycoCosm | scaffold | 2018 | one of the most morphologically complex extant Charophyta |
| **Mesostigma viride NIES-296** | Mesostigma_viride | streptophyte | 29795 | 442 | 8254 | 4231 | 5.91 | 3 | JGI PhycoCosm | scaffold | 2020 | unicellular biflagellate freshwater charophyte algae covered by an outer layer of basket-like scales |
| **Mesotaenium endlicherianum SAG 12.9** | Mesotaenium_endlicherianu | streptophyte | 11080 | 174 | 5852 | 4456 | 6.75 | 5 | JGI PhycoCosm | scaffold | 2019 | green alga found in subaerial/terrestrial environments associated with Bryophytes |
| **Klebsormidium nitens NIES-2285** | Klebsormidium_nitens | streptophyte | 16132 | 104 | 4965 | 4157 | 6.88 | 6 | JGI PhycoCosm | scaffold | 2014 | widely spread terrestial and aquatic. early diverging lineage of charophytes, multicellular non-branching filaments without differentiated or specialized cells. |
| **Chlorokybus atmophyticus CCAC 0220** | Chlorokybus_atmophyticus | streptophyte | 9300 | 74.34 | 2555 | 2201 | 8.03 | 7 | JGI PhycoCosm | scaffold | 2020 | subaerial/terrestrial environments, sarcinoid cell packets where each cell has its own cell wall, occasionally producing biflagellate, scaly zoospores |
| **Spirogloea muscicola CCAC 0214** | Spirogloea_muscicola | streptophyte | 27137 | 171 | 3351 | 2667 | 7.81 | 6 | JGI PhycoCosm | scaffold | 2019 | green alga found in subaerial/terrestrial environments associated with Bryophytes |
| **Physcomitrium patens v3.3** | Physcomitrium_patens | bryophyte | 32926 | 473 | NA | NA | NA | NA | Phytozome13 | chromosome | 2018 | species of moss, which is a basal lineage of land plants, having diverged before the acquisition of well-developed vasculature. |
| **Marchantia polymorpha v3.1** | Marchantia_polymorpha | bryophyte | 19287 | 225 | NA | NA | NA | NA | Phytozome13 | scaffold | 2017 | species of liverwort, M. polymorpha occupies a critical node in the evolution of land plants from their algal ancestors, with extant liverworts possibly retaining features of ancestral land plants. |
| **Selaginella moellendorffii v1.0** | Selaginella_moellendorffii | lycophyte | 22273 | 210 | NA | NA | NA | NA | Phytozome13 | scaffold | 2011 | the lycophytes sit between the bryophytes and the euphyllophytes |
| **Arabidopsis thaliana** | Arabidopsis_thaliana | dicot | 27655 | 135 | NA | NA | NA | NA | Dicots PLAZA 5.0 | chromosome | 2017 | dicot model organism |
| **Oryza sativa ssp. japonica** | Oryza_sativa | monocot | 42580 | 373.8 | NA | NA | NA | NA | NCBI | chromosome | 2015 | monocot model organism |
| **Cyanophora paradoxa CCMP329** | Cyanophora_paradoxa | glaucophyte | 25518 | 98 | 2332 | 1730 | 10.22 | 8 | JGI PhycoCosm | contig | 2019 | glaucophyte |
