## Supplemental Table S2 for "The *Draparnaldia* genome: alternative mechanisms for multicellularity and terrestrialization in green plants"

**Table S2a. Related to Figure 2A.** Statistics for all assemblies.

| assembly | N50 | L50 | span | num seqs | BUSCO | QV | k-mer comp. |
|---|---|---|---|---|---|---|---|
| flye | 185,940 | 403 | 247,448,887 | 4258 | C:95.9% [S:93.5%,D:2.4%],F:0.8%,M:3.3%,n:1519 | 33.2 | 92.7 |
| nextdenovo | 408,205 | 156 | 273,107,403 | 906 | C:88.0% [S:86.3%,D:1.7%],F:0.4%,M:11.6%,n:1519 | 31.9 | 84.5 |
| flye hypo | 185,959 | 403 | 247,504,853 | 4258 | C:96.5% [S:94.2%,D:2.3%],F:0.7%,M:2.8%,n:1519 | 35.9 | 93.3 |
| **flye hypo purged** | **195,149** | **378** | **237,927,493** | **2874** | **C:96.4% [S:94.4%,D:2.0%],F:0.7%,M:2.9%,n:1519** | **36.4** | **92.4** |
| nextdenovo hypo | 408,909 | 156 | 273,263,006 | 906 | C:88.3% [S:86.5%,D:1.8%],F:0.5%,M:11.2%,n:1519 | 35.8 | 85.8 |
| nextdenovo hypo purged | 432,865 | 144 | 263,054,452 | 829 | C:87.8% [S:86.2%,D:1.6%],F:0.5%,M:11.7%,n:1519 | 36.0 | 85.1 |

**Table S2b. Related to Figure 2A.** Assembly pipeline configuration

| Rule | Programs and Versions | Parameters/Notes |
|---|---|---|
| genomescope2 | Genomescope v2 | k=18 p=2 |
| filtLong | FiltLong 0.2.1 | --min length 1000 --min mean q 80 |
| trim galore | Cutadapt 4.1 | --gzip -q 20 --paired --retain unpaired |
| nanoplot | NanoPlot 1.40.0 | --plots dot |
| concat reads | gzip 1.5 | zcat command and redirection |
| align illumina | BWA-MEM2 2.2.1 | bwa-mem2 mem -Y (use soft clipping for supplementary alignments) |
| align_ont | minimap2 2.24-r1122 | -ax map-ont |
| purge dups | purge dups 1.2.5 |  |
| run busco | BUSCO 4.0.6 | genome mode using the chlorophyta odb10 database |
| run merqury | merqury 1.1 | k=18 |
| blobtools | blobtoolkit v3.3.2 | |
| hypo | Hypo 1.0.3 | |
| Flye | Flye 2.9.1-b1780 | --nano-raw -i 2 --scaffold -g 466m |
| Nextdenovo | NextDenovo 2.4.0 | read_cutoff = 1k genome_size = 466m seed_depth = 45 seed_cutoff = |

**Table S2c. Related to Figure 2A.** DRAP1A annotation statistics

| | |
|---|---|
| Number of protein-coding genes | 27,433 |
| Median gene length (bp) | 4,240 |
| Number of transcripts | 27,433 |
| Number of exons | 154,004 |
| Number of coding exons | 146,360 |
| Median UTR length (bp) | 1,546 |
| Median intron length (bp) | 376 |
| Exons/transcript | 7.59 |
| Transcripts/gene | 1.44 |
| Multi-exonic transcripts | 94% |
| Gene density (gene/Mb) | 80.2 |
