## Supplemental Table S4 for "The *Draparnaldia* genome: alternative mechanisms for multicellularity and terrestrialization in green plants"

**Table S4a. Related to Figure 3. Enriched GO terms in orthogroups expanded in Draparnaldia with respect to Chlamydomonas.**

| ID | Description | GeneRatio | BgRatio | pvalue | p.adjust | qvalue | geneID | Count |
|---|---|---|---|---|---|---|---|---|
| GO:0031226 | intrinsic component of plasma membrane | 3/22 | 56/13359 | 0.000101549321747445 | 0.0401116859452821 | 0.0256212822071869 | OG0000035/OG0000124/OG0000127 | 3 |
| GO:0010218 | response to far red light | 2/22 | 16/13359 | 0.000306364313681248 | 0.0401116859452821 | 0.0256212822071869 | OG0000035/OG0000047 | 2 |
| GO:0005887 | integral component of plasma membrane | 2/22 | 23/13359 | 0.000641423450863759 | 0.0401116859452821 | 0.0256212822071869 | OG0000035/OG0000127 | 2 |
| GO:0051607 | defense response to virus | 2/22 | 23/13359 | 0.000641423450863759 | 0.0401116859452821 | 0.0256212822071869 | OG0000124/OG0000167 | 2 |
| GO:0140546 | defense response to symbiont | 2/22 | 23/13359 | 0.000641423450863759 | 0.0401116859452821 | 0.0256212822071869 | OG0000124/OG0000167 | 2 |
| GO:0048869 | cellular developmental process | 4/22 | 266/13359 | 0.000847210374204637 | 0.0401116859452821 | 0.0256212822071869 | OG0000035/OG0000167/OG0000721/OG0001956 | 4 |
| GO:0009628 | response to abiotic stimulus | 6/22 | 726/13359 | 0.000889138224304288 | 0.0401116859452821 | 0.0256212822071869 | OG0000035/OG0000047/OG0000124/OG0000167/OG0000721/OG0001956 | 6 |
| GO:0009699 | phenylpropanoid biosynthetic process | 2/22 | 29/13359 | 0.00102318023375165 | 0.0401116859452821 | 0.0256212822071869 | OG0000124/OG0000721 | 2 |
| GO:0009615 | response to virus | 2/22 | 34/13359 | 0.00140677497935591 | 0.0401116859452821 | 0.0256212822071869 | OG0000124/OG0000167 | 2 |
| GO:0010029 | regulation of seed germination | 2/22 | 35/13359 | 0.00149054799493739 | 0.0401116859452821 | 0.0256212822071869 | OG0000047/OG0000167 | 2 |
| GO:0009637 | response to blue light | 2/22 | 36/13359 | 0.00157665547673467 | 0.0401116859452821 | 0.0256212822071869 | OG0000035/OG0000047 | 2 |
| GO:0009698 | phenylpropanoid metabolic process | 2/22 | 36/13359 | 0.00157665547673467 | 0.0401116859452821 | 0.0256212822071869 | OG0000124/OG0000721 | 2 |
| GO:0008271 | secondary active sulfate transmembrane transporter activity | 1/22 | 1/13359 | 0.00164682985253406 | 0.0401116859452821 | 0.0256212822071869 | OG0000127 | 1 |
| GO:0008447 | L-ascorbate oxidase activity | 1/22 | 1/13359 | 0.00164682985253406 | 0.0401116859452821 | 0.0256212822071869 | OG0000124 | 1 |
| GO:0010329 | auxin efflux transmembrane transporter activity | 1/22 | 1/13359 | 0.00164682985253406 | 0.0401116859452821 | 0.0256212822071869 | OG0000035 | 1 |
| GO:0019854 | L-ascorbic acid catabolic process | 1/22 | 1/13359 | 0.00164682985253406 | 0.0401116859452821 | 0.0256212822071869 | OG0000124 | 1 |
| GO:0035336 | long-chain fatty-acyl-CoA metabolic process | 1/22 | 1/13359 | 0.00164682985253406 | 0.0401116859452821 | 0.0256212822071869 | OG0000721 | 1 |
| GO:0042084 | 5-methyltetrahydrofolate-dependent methyltransferase activity | 1/22 | 1/13359 | 0.00164682985253406 | 0.0401116859452821 | 0.0256212822071869 | OG0000721 | 1 |
| GO:0050062 | long-chain-fatty-acyl-CoA reductase activity | 1/22 | 1/13359 | 0.00164682985253406 | 0.0401116859452821 | 0.0256212822071869 | OG0000721 | 1 |
| GO:0052716 | hydroquinone:oxygen oxidoreductase activity | 1/22 | 1/13359 | 0.00164682985253406 | 0.0401116859452821 | 0.0256212822071869 | OG0000124 | 1 |
| GO:0080019 | fatty-acyl-CoA reductase (alcohol-forming) activity | 1/22 | 1/13359 | 0.00164682985253406 | 0.0401116859452821 | 0.0256212822071869 | OG0000721 | 1 |
| GO:0080160 | selenate transport | 1/22 | 1/13359 | 0.00164682985253406 | 0.0401116859452821 | 0.0256212822071869 | OG0000127 | 1 |
| GO:0102965 | alcohol-forming fatty acyl-CoA reductase activity | 1/22 | 1/13359 | 0.00164682985253406 | 0.0401116859452821 | 0.0256212822071869 | OG0000721 | 1 |
| GO:0005911 | cell-cell junction | 4/22 | 322/13359 | 0.00171891852284634 | 0.0401116859452821 | 0.0256212822071869 | OG0000035/OG0000124/OG0000127/OG0000721 | 4 |
| GO:0009506 | plasmodesma | 4/22 | 322/13359 | 0.00171891852284634 | 0.0401116859452821 | 0.0256212822071869 | OG0000035/OG0000124/OG0000127/OG0000721 | 4 |
| GO:0055044 | symplast | 4/22 | 322/13359 | 0.00171891852284634 | 0.0401116859452821 | 0.0256212822071869 | OG0000035/OG0000124/OG0000127/OG0000721 | 4 |
| GO:0070161 | anchoring junction | 4/22 | 322/13359 | 0.00171891852284634 | 0.0401116859452821 | 0.0256212822071869 | OG0000035/OG0000124/OG0000127/OG0000721 | 4 |
| GO:0030054 | cell junction | 4/22 | 323/13359 | 0.00173858700691625 | 0.0401116859452821 | 0.0256212822071869 | OG0000035/OG0000124/OG0000127/OG0000721 | 4 |
| GO:1900140 | regulation of seedling development | 2/22 | 39/13359 | 0.00184891272684474 | 0.0404033364384395 | 0.0258075735438623 | OG0000047/OG0000167 | 2 |
| GO:0009651 | response to salt stress | 3/22 | 160/13359 | 0.00219663065678811 | 0.0404033364384395 | 0.0258075735438623 | OG0000124/OG0000721/OG0001956 | 3 |
| GO:0007275 | multicellular organism development | 6/22 | 876/13359 | 0.00234810087442321 | 0.0404033364384395 | 0.0258075735438623 | OG0000035/OG0000047/OG0000124/OG0000167/OG0000721/OG0001956 | 6 |
| GO:0005576 | extracellular region | 5/22 | 617/13359 | 0.00282797756943533 | 0.0404033364384395 | 0.0258075735438623 | OG0000035/OG0000124/OG0000572/OG0000721/OG0000785 | 5 |
| GO:0044550 | secondary metabolite biosynthetic process | 2/22 | 50/13359 | 0.00302327143007577 | 0.0404033364384395 | 0.0258075735438623 | OG0000124/OG0000721 | 2 |
| GO:0005886 | plasma membrane | 6/22 | 926/13359 | 0.00310888695890322 | 0.0404033364384395 | 0.0258075735438623 | OG0000035/OG0000124/OG0000127/OG0000167/OG0000721/OG0000785 | 6 |
| GO:0008172 | S-methyltransferase activity | 1/22 | 2/13359 | 0.00329107073763935 | 0.0404033364384395 | 0.0258075735438623 | OG0000721 | 1 |
| GO:0008559 | ABC-type xenobiotic transporter activity | 1/22 | 2/13359 | 0.00329107073763935 | 0.0404033364384395 | 0.0258075735438623 | OG0000035 | 1 |
| GO:0008705 | methionine synthase activity | 1/22 | 2/13359 | 0.00329107073763935 | 0.0404033364384395 | 0.0258075735438623 | OG0000721 | 1 |
| GO:0009111 | vitamin catabolic process | 1/22 | 2/13359 | 0.00329107073763935 | 0.0404033364384395 | 0.0258075735438623 | OG0000124 | 1 |
| GO:0010315 | auxin export across the plasma membrane | 1/22 | 2/13359 | 0.00329107073763935 | 0.0404033364384395 | 0.0258075735438623 | OG0000035 | 1 |
| GO:0010328 | auxin influx transmembrane transporter activity | 1/22 | 2/13359 | 0.00329107073763935 | 0.0404033364384395 | 0.0258075735438623 | OG0000035 | 1 |
| GO:0010541 | acropetal auxin transport | 1/22 | 2/13359 | 0.00329107073763935 | 0.0404033364384395 | 0.0258075735438623 | OG0000035 | 1 |
| GO:0035337 | fatty-acyl-CoA metabolic process | 1/22 | 2/13359 | 0.00329107073763935 | 0.0404033364384395 | 0.0258075735438623 | OG0000721 | 1 |
| GO:0042365 | water-soluble vitamin catabolic process | 1/22 | 2/13359 | 0.00329107073763935 | 0.0404033364384395 | 0.0258075735438623 | OG0000124 | 1 |
| GO:0046271 | phenylpropanoid catabolic process | 1/22 | 2/13359 | 0.00329107073763935 | 0.0404033364384395 | 0.0258075735438623 | OG0000124 | 1 |
| GO:0046274 | lignin catabolic process | 1/22 | 2/13359 | 0.00329107073763935 | 0.0404033364384395 | 0.0258075735438623 | OG0000124 | 1 |
| GO:0050687 | negative regulation of defense response to virus | 1/22 | 2/13359 | 0.00329107073763935 | 0.0404033364384395 | 0.0258075735438623 | OG0000124 | 1 |
| GO:0060919 | auxin import into cell | 1/22 | 2/13359 | 0.00329107073763935 | 0.0404033364384395 | 0.0258075735438623 | OG0000035 | 1 |
| GO:0071490 | cellular response to far red light | 1/22 | 2/13359 | 0.00329107073763935 | 0.0404033364384395 | 0.0258075735438623 | OG0000047 | 1 |
| GO:0080161 | auxin transmembrane transporter activity | 1/22 | 2/13359 | 0.00329107073763935 | 0.0404033364384395 | 0.0258075735438623 | OG0000035 | 1 |
| GO:1902358 | sulfate transmembrane transport | 1/22 | 2/13359 | 0.00329107073763935 | 0.0404033364384395 | 0.0258075735438623 | OG0000127 | 1 |
| GO:1903173 | fatty alcohol metabolic process | 1/22 | 2/13359 | 0.00329107073763935 | 0.0404033364384395 | 0.0258075735438623 | OG0000721 | 1 |
| GO:1903175 | fatty alcohol biosynthetic process | 1/22 | 2/13359 | 0.00329107073763935 | 0.0404033364384395 | 0.0258075735438623 | OG0000721 | 1 |
| GO:0006970 | response to osmotic stress | 3/22 | 185/13359 | 0.0033148248161568 | 0.0404033364384395 | 0.0258075735438623 | OG0000124/OG0000721/OG0001956 | 3 |
| GO:0032501 | multicellular organismal process | 6/22 | 948/13359 | 0.00349743963596233 | 0.041839740830216 | 0.0267250748011156 | OG0000035/OG0000047/OG0000124/OG0000167/OG0000721/OG0001956 | 6 |
| GO:0009411 | response to UV | 2/22 | 56/13359 | 0.00377805726189257 | 0.0443749998396837 | 0.0283444678786964 | OG0000035/OG0000047 | 2 |
| GO:0048856 | anatomical structure development | 6/22 | 969/13359 | 0.0039016248904214 | 0.0450080299859326 | 0.0287488149820524 | OG0000035/OG0000047/OG0000124/OG0000167/OG0000721/OG0001956 | 6 |
| GO:0033554 | cellular response to stress | 4/22 | 427/13359 | 0.00476534877910173 | 0.0468609020529363 | 0.0299323343730667 | OG0000047/OG0000127/OG0000167/OG0001956 | 4 |
| GO:0032502 | developmental process | 6/22 | 1009/13359 | 0.00476793922103548 | 0.0468609020529363 | 0.0299323343730667 | OG0000035/OG0000047/OG0000124/OG0000167/OG0000721/OG0001956 | 6 |
| GO:0009970 | cellular response to sulfate starvation | 1/22 | 3/13359 | 0.00493272653188803 | 0.0468609020529363 | 0.0299323343730667 | OG0000127 | 1 |
| GO:0043473 | pigmentation | 1/22 | 3/13359 | 0.00493272653188803 | 0.0468609020529363 | 0.0299323343730667 | OG0000035 | 1 |
| GO:0043476 | pigment accumulation | 1/22 | 3/13359 | 0.00493272653188803 | 0.0468609020529363 | 0.0299323343730667 | OG0000035 | 1 |
| GO:0043478 | pigment accumulation in response to UV light | 1/22 | 3/13359 | 0.00493272653188803 | 0.0468609020529363 | 0.0299323343730667 | OG0000035 | 1 |
| GO:0043479 | pigment accumulation in tissues in response to UV light | 1/22 | 3/13359 | 0.00493272653188803 | 0.0468609020529363 | 0.0299323343730667 | OG0000035 | 1 |
| GO:0043480 | pigment accumulation in tissues | 1/22 | 3/13359 | 0.00493272653188803 | 0.0468609020529363 | 0.0299323343730667 | OG0000035 | 1 |
| GO:0043481 | anthocyanin accumulation in tissues in response to UV light | 1/22 | 3/13359 | 0.00493272653188803 | 0.0468609020529363 | 0.0299323343730667 | OG0000035 | 1 |
| GO:0070141 | response to UV-A | 1/22 | 3/13359 | 0.00493272653188803 | 0.0468609020529363 | 0.0299323343730667 | OG0000047 | 1 |
| GO:0071491 | cellular response to red light | 1/22 | 3/13359 | 0.00493272653188803 | 0.0468609020529363 | 0.0299323343730667 | OG0000047 | 1 |
| GO:0071492 | cellular response to UV-A | 1/22 | 3/13359 | 0.00493272653188803 | 0.0468609020529363 | 0.0299323343730667 | OG0000047 | 1 |
| GO:0071944 | cell periphery | 6/22 | 1021/13359 | 0.00505397316391651 | 0.0473169081723198 | 0.0302236076316594 | OG0000035/OG0000124/OG0000127/OG0000167/OG0000721/OG0000785 | 6 |
| GO:0006950 | response to stress | 6/22 | 1027/13359 | 0.00520171944213102 | 0.0480044394230949 | 0.030662767237825 | OG0000047/OG0000124/OG0000127/OG0000167/OG0000721/OG0001956 | 6 |
| GO:1901617 | organic hydroxy compound biosynthetic process | 2/22 | 68/13359 | 0.00552227996274134 | 0.0502449698018438 | 0.0320939028227518 | OG0000124/OG0000721 | 2 |
| GO:0030154 | cell differentiation | 3/22 | 232/13359 | 0.00623911658576919 | 0.0511491989721894 | 0.0326714779160799 | OG0000035/OG0000167/OG0001956 | 3 |
| GO:0009845 | seed germination | 2/22 | 74/13359 | 0.00650877735628915 | 0.0511491989721894 | 0.0326714779160799 | OG0000047/OG0000167 | 2 |
| GO:0008272 | sulfate transport | 1/22 | 4/13359 | 0.00657180110633393 | 0.0511491989721894 | 0.0326714779160799 | OG0000127 | 1 |
| GO:0009806 | lignan metabolic process | 1/22 | 4/13359 | 0.00657180110633393 | 0.0511491989721894 | 0.0326714779160799 | OG0000124 | 1 |
| GO:0009807 | lignan biosynthetic process | 1/22 | 4/13359 | 0.00657180110633393 | 0.0511491989721894 | 0.0326714779160799 | OG0000124 | 1 |
| GO:0015116 | sulfate transmembrane transporter activity | 1/22 | 4/13359 | 0.00657180110633393 | 0.0511491989721894 | 0.0326714779160799 | OG0000127 | 1 |
| GO:0015301 | anion:anion antiporter activity | 1/22 | 4/13359 | 0.00657180110633393 | 0.0511491989721894 | 0.0326714779160799 | OG0000127 | 1 |
| GO:0016682 | oxidoreductase activity, acting on diphenols and related substances as donors, oxygen as acceptor | 1/22 | 4/13359 | 0.00657180110633393 | 0.0511491989721894 | 0.0326714779160799 | OG0000124 | 1 |
| GO:0046365 | monosaccharide catabolic process | 1/22 | 4/13359 | 0.00657180110633393 | 0.0511491989721894 | 0.0326714779160799 | OG0000124 | 1 |
| GO:0046688 | response to copper ion | 1/22 | 4/13359 | 0.00657180110633393 | 0.0511491989721894 | 0.0326714779160799 | OG0000124 | 1 |
| GO:0071486 | cellular response to high light intensity | 1/22 | 4/13359 | 0.00657180110633393 | 0.0511491989721894 | 0.0326714779160799 | OG0000047 | 1 |
| GO:0140323 | solute:anion antiporter activity | 1/22 | 4/13359 | 0.00657180110633393 | 0.0511491989721894 | 0.0326714779160799 | OG0000127 | 1 |
| GO:0019748 | secondary metabolic process | 2/22 | 76/13359 | 0.00685419812216125 | 0.0527120474632877 | 0.0336697451614938 | OG0000124/OG0000721 | 2 |
| GO:0048646 | anatomical structure formation involved in morphogenesis | 2/22 | 77/13359 | 0.00702998896942936 | 0.0534279161676632 | 0.0341270052757438 | OG0000035/OG0000721 | 2 |
| GO:0009605 | response to external stimulus | 4/22 | 484/13359 | 0.00740663752115437 | 0.0546655744220136 | 0.0349175577210841 | OG0000035/OG0000124/OG0000127/OG0000167 | 4 |
| GO:0019439 | aromatic compound catabolic process | 2/22 | 80/13359 | 0.00756960431122284 | 0.0546655744220136 | 0.0349175577210841 | OG0000124/OG0000167 | 2 |
| GO:0009639 | response to red or far red light | 2/22 | 81/13359 | 0.00775353514892631 | 0.0546655744220136 | 0.0349175577210841 | OG0000035/OG0000047 | 2 |
| GO:0022804 | active transmembrane transporter activity | 2/22 | 82/13359 | 0.00793948461302868 | 0.0546655744220136 | 0.0349175577210841 | OG0000035/OG0000127 | 2 |
| GO:1901361 | organic cyclic compound catabolic process | 2/22 | 82/13359 | 0.00793948461302868 | 0.0546655744220136 | 0.0349175577210841 | OG0000124/OG0000167 | 2 |
| GO:0009536 | plastid | 8/22 | 1899/13359 | 0.00820459899070351 | 0.0546655744220136 | 0.0349175577210841 | OG0000035/OG0000047/OG0000124/OG0000127/OG0000167/OG0000572/OG0000721/OG0001956 | 8 |
| GO:0010023 | proanthocyanidin biosynthetic process | 1/22 | 5/13359 | 0.00820829832652525 | 0.0546655744220136 | 0.0349175577210841 | OG0000124 | 1 |
| GO:0010540 | basipetal auxin transport | 1/22 | 5/13359 | 0.00820829832652525 | 0.0546655744220136 | 0.0349175577210841 | OG0000035 | 1 |
| GO:0016679 | oxidoreductase activity, acting on diphenols and related substances as donors | 1/22 | 5/13359 | 0.00820829832652525 | 0.0546655744220136 | 0.0349175577210841 | OG0000124 | 1 |
| GO:0042908 | xenobiotic transport | 1/22 | 5/13359 | 0.00820829832652525 | 0.0546655744220136 | 0.0349175577210841 | OG0000035 | 1 |
| GO:0071484 | cellular response to light intensity | 1/22 | 5/13359 | 0.00820829832652525 | 0.0546655744220136 | 0.0349175577210841 | OG0000047 | 1 |
| GO:1900367 | positive regulation of defense response to insect | 1/22 | 5/13359 | 0.00820829832652525 | 0.0546655744220136 | 0.0349175577210841 | OG0000124 | 1 |
| GO:0090351 | seedling development | 2/22 | 84/13359 | 0.00831741335564908 | 0.0548270308954011 | 0.0350206878132593 | OG0000047/OG0000167 | 2 |
| GO:0051716 | cellular response to stimulus | 5/22 | 810/13359 | 0.00898018450344098 | 0.0585979716083118 | 0.0374293708171065 | OG0000035/OG0000047/OG0000127/OG0000167/OG0001956 | 5 |
| GO:0050896 | response to stimulus | 7/22 | 1545/13359 | 0.00968624946754059 | 0.0623340729992402 | 0.0398158002537106 | OG0000035/OG0000047/OG0000124/OG0000127/OG0000167/OG0000721/OG0001956 | 7 |
| GO:0010277 | chlorophyllide a oxygenase [overall] activity | 1/22 | 6/13359 | 0.00984222205251162 | 0.0623340729992402 | 0.0398158002537106 | OG0000151 | 1 |
| GO:0010345 | suberin biosynthetic process | 1/22 | 6/13359 | 0.00984222205251162 | 0.0623340729992402 | 0.0398158002537106 | OG0000721 | 1 |
| GO:0071456 | cellular response to hypoxia | 2/22 | 94/13359 | 0.0103259721159515 | 0.0647628930767446 | 0.0413672056152581 | OG0000167/OG0001956 | 2 |
| GO:0036294 | cellular response to decreased oxygen levels | 2/22 | 95/13359 | 0.0105375756065827 | 0.0648311794462137 | 0.0414108234363953 | OG0000167/OG0001956 | 2 |
| GO:0071453 | cellular response to oxygen levels | 2/22 | 95/13359 | 0.0105375756065827 | 0.0648311794462137 | 0.0414108234363953 | OG0000167/OG0001956 | 2 |
| GO:0042221 | response to chemical | 5/22 | 855/13359 | 0.0112130279225601 | 0.0667741458170757 | 0.0426518904355445 | OG0000035/OG0000124/OG0000167/OG0000721/OG0001956 | 5 |
| GO:0010117 | photoprotection | 1/22 | 7/13359 | 0.0114735761388474 | 0.0667741458170757 | 0.0426518904355445 | OG0000047 | 1 |
| GO:0010380 | regulation of chlorophyll biosynthetic process | 1/22 | 7/13359 | 0.0114735761388474 | 0.0667741458170757 | 0.0426518904355445 | OG0000047 | 1 |
| GO:0034644 | cellular response to UV | 1/22 | 7/13359 | 0.0114735761388474 | 0.0667741458170757 | 0.0426518904355445 | OG0000047 | 1 |
| GO:0048226 | Casparian strip | 1/22 | 7/13359 | 0.0114735761388474 | 0.0667741458170757 | 0.0426518904355445 | OG0000124 | 1 |
| GO:2000068 | regulation of defense response to insect | 1/22 | 7/13359 | 0.0114735761388474 | 0.0667741458170757 | 0.0426518904355445 | OG0000124 | 1 |
| GO:0001666 | response to hypoxia | 2/22 | 102/13359 | 0.0120724489792139 | 0.0696321610765374 | 0.0444774436076302 | OG0000167/OG0001956 | 2 |
| GO:1901615 | organic hydroxy compound metabolic process | 2/22 | 103/13359 | 0.012299305377025 | 0.0699514663202831 | 0.0446813993768144 | OG0000124/OG0000721 | 2 |
| GO:0048046 | apoplast | 2/22 | 105/13359 | 0.0127586473326649 | 0.0699514663202831 | 0.0446813993768144 | OG0000124/OG0000721 | 2 |
| GO:0006555 | methionine metabolic process | 1/22 | 8/13359 | 0.0131023644346041 | 0.0699514663202831 | 0.0446813993768144 | OG0000721 | 1 |
| GO:0009086 | methionine biosynthetic process | 1/22 | 8/13359 | 0.0131023644346041 | 0.0699514663202831 | 0.0446813993768144 | OG0000721 | 1 |
| GO:0009531 | secondary cell wall | 1/22 | 8/13359 | 0.0131023644346041 | 0.0699514663202831 | 0.0446813993768144 | OG0000124 | 1 |
| GO:0010494 | cytoplasmic stress granule | 1/22 | 8/13359 | 0.0131023644346041 | 0.0699514663202831 | 0.0446813993768144 | OG0000167 | 1 |
| GO:0050688 | regulation of defense response to virus | 1/22 | 8/13359 | 0.0131023644346041 | 0.0699514663202831 | 0.0446813993768144 | OG0000124 | 1 |
| GO:1901001 | negative regulation of response to salt stress | 1/22 | 8/13359 | 0.0131023644346041 | 0.0699514663202831 | 0.0446813993768144 | OG0000124 | 1 |
| GO:1901463 | regulation of tetrapyrrole biosynthetic process | 1/22 | 8/13359 | 0.0131023644346041 | 0.0699514663202831 | 0.0446813993768144 | OG0000047 | 1 |
| GO:0036293 | response to decreased oxygen levels | 2/22 | 107/13359 | 0.0132254534974561 | 0.0700298603225954 | 0.0447314734340189 | OG0000167/OG0001956 | 2 |
| GO:0070482 | response to oxygen levels | 2/22 | 108/13359 | 0.0134616402081517 | 0.0707009721501302 | 0.0451601451570002 | OG0000167/OG0001956 | 2 |
| GO:0010187 | negative regulation of seed germination | 1/22 | 9/13359 | 0.0147285907833753 | 0.0743333566098472 | 0.0474803255516704 | OG0000167 | 1 |
| GO:0016620 | oxidoreductase activity, acting on the aldehyde or oxo group of donors, NAD or NADP as acceptor | 1/22 | 9/13359 | 0.0147285907833753 | 0.0743333566098472 | 0.0474803255516704 | OG0000721 | 1 |
| GO:0048859 | formation of anatomical boundary | 1/22 | 9/13359 | 0.0147285907833753 | 0.0743333566098472 | 0.0474803255516704 | OG0000035 | 1 |
| GO:0090056 | regulation of chlorophyll metabolic process | 1/22 | 9/13359 | 0.0147285907833753 | 0.0743333566098472 | 0.0474803255516704 | OG0000047 | 1 |
| GO:0090691 | formation of plant organ boundary | 1/22 | 9/13359 | 0.0147285907833753 | 0.0743333566098472 | 0.0474803255516704 | OG0000035 | 1 |
| GO:0016491 | oxidoreductase activity | 4/22 | 601/13359 | 0.0155456114861736 | 0.0776732303605909 | 0.0496136651480392 | OG0000124/OG0000151/OG0000253/OG0000721 | 4 |
| GO:0022857 | transmembrane transporter activity | 3/22 | 326/13359 | 0.0157235675776992 | 0.0776732303605909 | 0.0496136651480392 | OG0000035/OG0000127/OG0000189 | 3 |
| GO:0019438 | aromatic compound biosynthetic process | 4/22 | 605/13359 | 0.0158954528239091 | 0.0776732303605909 | 0.0496136651480392 | OG0000047/OG0000124/OG0000167/OG0000721 | 4 |
| GO:0006817 | phosphate ion transport | 1/22 | 10/13359 | 0.0163522590232823 | 0.0776732303605909 | 0.0496136651480392 | OG0000127 | 1 |
| GO:0008361 | regulation of cell size | 1/22 | 10/13359 | 0.0163522590232823 | 0.0776732303605909 | 0.0496136651480392 | OG0000035 | 1 |
| GO:0019852 | L-ascorbic acid metabolic process | 1/22 | 10/13359 | 0.0163522590232823 | 0.0776732303605909 | 0.0496136651480392 | OG0000124 | 1 |
| GO:0098661 | inorganic anion transmembrane transport | 1/22 | 10/13359 | 0.0163522590232823 | 0.0776732303605909 | 0.0496136651480392 | OG0000127 | 1 |
| GO:1901401 | regulation of tetrapyrrole metabolic process | 1/22 | 10/13359 | 0.0163522590232823 | 0.0776732303605909 | 0.0496136651480392 | OG0000047 | 1 |
| GO:0048580 | regulation of post-embryonic development | 2/22 | 125/13359 | 0.0177545170378635 | 0.0811943982489062 | 0.0518628061162966 | OG0000047/OG0000167 | 2 |
| GO:0009067 | aspartate family amino acid biosynthetic process | 1/22 | 11/13359 | 0.017973372986987 | 0.0811943982489062 | 0.0518628061162966 | OG0000721 | 1 |
| GO:0010584 | pollen exine formation | 1/22 | 11/13359 | 0.017973372986987 | 0.0811943982489062 | 0.0518628061162966 | OG0000721 | 1 |
| GO:0016903 | oxidoreductase activity, acting on the aldehyde or oxo group of donors | 1/22 | 11/13359 | 0.017973372986987 | 0.0811943982489062 | 0.0518628061162966 | OG0000721 | 1 |
| GO:0050779 | RNA destabilization | 1/22 | 11/13359 | 0.017973372986987 | 0.0811943982489062 | 0.0518628061162966 | OG0000167 | 1 |
| GO:0061157 | mRNA destabilization | 1/22 | 11/13359 | 0.017973372986987 | 0.0811943982489062 | 0.0518628061162966 | OG0000167 | 1 |
| GO:0080022 | primary root development | 1/22 | 11/13359 | 0.017973372986987 | 0.0811943982489062 | 0.0518628061162966 | OG0000124 | 1 |
| GO:2000026 | regulation of multicellular organismal development | 2/22 | 127/13359 | 0.0182932833969885 | 0.0820657019059343 | 0.0524193500849377 | OG0000047/OG0000167 | 2 |
| GO:0009507 | chloroplast | 7/22 | 1761/13359 | 0.0192679341016069 | 0.083265730132191 | 0.0531858664034852 | OG0000035/OG0000047/OG0000127/OG0000167/OG0000572/OG0000721/OG0001956 | 7 |
| GO:0099402 | plant organ development | 3/22 | 353/13359 | 0.0194125459235258 | 0.083265730132191 | 0.0531858664034852 | OG0000035/OG0000124/OG0000167 | 3 |
| GO:0000097 | sulfur amino acid biosynthetic process | 1/22 | 12/13359 | 0.019591936501692 | 0.083265730132191 | 0.0531858664034852 | OG0000721 | 1 |
| GO:0003727 | single-stranded RNA binding | 1/22 | 12/13359 | 0.019591936501692 | 0.083265730132191 | 0.0531858664034852 | OG0000167 | 1 |
| GO:0009635 | response to herbicide | 1/22 | 12/13359 | 0.019591936501692 | 0.083265730132191 | 0.0531858664034852 | OG0000721 | 1 |
| GO:0010025 | wax biosynthetic process | 1/22 | 12/13359 | 0.019591936501692 | 0.083265730132191 | 0.0531858664034852 | OG0000721 | 1 |
| GO:0010030 | positive regulation of seed germination | 1/22 | 12/13359 | 0.019591936501692 | 0.083265730132191 | 0.0531858664034852 | OG0000047 | 1 |
| GO:0010166 | wax metabolic process | 1/22 | 12/13359 | 0.019591936501692 | 0.083265730132191 | 0.0531858664034852 | OG0000721 | 1 |
| GO:1901362 | organic cyclic compound biosynthetic process | 4/22 | 648/13359 | 0.0199777936987217 | 0.0843506845057138 | 0.0538788794626688 | OG0000047/OG0000124/OG0000167/OG0000721 | 4 |
| GO:0043207 | response to external biotic stimulus | 3/22 | 360/13359 | 0.0204414739779735 | 0.0850952663937507 | 0.0543544800820438 | OG0000035/OG0000124/OG0000167 | 3 |
| GO:0051707 | response to other organism | 3/22 | 360/13359 | 0.0204414739779735 | 0.0850952663937507 | 0.0543544800820438 | OG0000035/OG0000124/OG0000167 | 3 |
| GO:0009607 | response to biotic stimulus | 3/22 | 362/13359 | 0.0207409646311452 | 0.0850952663937507 | 0.0543544800820438 | OG0000035/OG0000124/OG0000167 | 3 |
| GO:0044419 | biological process involved in interspecies interaction between organisms | 3/22 | 365/13359 | 0.0211947988671434 | 0.0850952663937507 | 0.0543544800820438 | OG0000035/OG0000124/OG0000167 | 3 |
| GO:0046189 | phenol-containing compound biosynthetic process | 1/22 | 13/13359 | 0.0212079533891546 | 0.0850952663937507 | 0.0543544800820438 | OG0000124 | 1 |
| GO:0071483 | cellular response to blue light | 1/22 | 13/13359 | 0.0212079533891546 | 0.0850952663937507 | 0.0543544800820438 | OG0000047 | 1 |
| GO:1901568 | fatty acid derivative metabolic process | 1/22 | 13/13359 | 0.0212079533891546 | 0.0850952663937507 | 0.0543544800820438 | OG0000721 | 1 |
| GO:1901570 | fatty acid derivative biosynthetic process | 1/22 | 13/13359 | 0.0212079533891546 | 0.0850952663937507 | 0.0543544800820438 | OG0000721 | 1 |
| GO:0009066 | aspartate family amino acid metabolic process | 1/22 | 14/13359 | 0.0228214274656896 | 0.0888110972459969 | 0.0567279617409659 | OG0000721 | 1 |
| GO:0010928 | regulation of auxin mediated signaling pathway | 1/22 | 14/13359 | 0.0228214274656896 | 0.0888110972459969 | 0.0567279617409659 | OG0000035 | 1 |
| GO:0015293 | symporter activity | 1/22 | 14/13359 | 0.0228214274656896 | 0.0888110972459969 | 0.0567279617409659 | OG0000127 | 1 |
| GO:0061014 | positive regulation of mRNA catabolic process | 1/22 | 14/13359 | 0.0228214274656896 | 0.0888110972459969 | 0.0567279617409659 | OG0000167 | 1 |
| GO:1903313 | positive regulation of mRNA metabolic process | 1/22 | 14/13359 | 0.0228214274656896 | 0.0888110972459969 | 0.0567279617409659 | OG0000167 | 1 |
| GO:0009409 | response to cold | 2/22 | 144/13359 | 0.0231479847240037 | 0.0895425037826731 | 0.0571951466234445 | OG0000047/OG0000167 | 2 |
| GO:0009526 | plastid envelope | 3/22 | 385/13359 | 0.0243616474483487 | 0.0923000362704563 | 0.0589565165683866 | OG0000047/OG0000127/OG0000721 | 3 |
| GO:0009958 | positive gravitropism | 1/22 | 15/13359 | 0.0244323625421796 | 0.0923000362704563 | 0.0589565165683866 | OG0000035 | 1 |
| GO:0010208 | pollen wall assembly | 1/22 | 15/13359 | 0.0244323625421796 | 0.0923000362704563 | 0.0589565165683866 | OG0000721 | 1 |
| GO:1901682 | sulfur compound transmembrane transporter activity | 1/22 | 15/13359 | 0.0244323625421796 | 0.0923000362704563 | 0.0589565165683866 | OG0000127 | 1 |
| GO:0048518 | positive regulation of biological process | 3/22 | 393/13359 | 0.0256973027589974 | 0.0939795113181867 | 0.0600292788605657 | OG0000047/OG0000124/OG0000167 | 3 |
| GO:0051239 | regulation of multicellular organismal process | 2/22 | 153/13359 | 0.025911191263413 | 0.0939795113181867 | 0.0600292788605657 | OG0000047/OG0000167 | 2 |
| GO:0000741 | karyogamy | 1/22 | 16/13359 | 0.0260407624240796 | 0.0939795113181867 | 0.0600292788605657 | OG0001956 | 1 |
| GO:0002213 | defense response to insect | 1/22 | 16/13359 | 0.0260407624240796 | 0.0939795113181867 | 0.0600292788605657 | OG0000124 | 1 |
| GO:0010197 | polar nucleus fusion | 1/22 | 16/13359 | 0.0260407624240796 | 0.0939795113181867 | 0.0600292788605657 | OG0001956 | 1 |
| GO:0043488 | regulation of mRNA stability | 1/22 | 16/13359 | 0.0260407624240796 | 0.0939795113181867 | 0.0600292788605657 | OG0000167 | 1 |
| GO:0072348 | sulfur compound transport | 1/22 | 16/13359 | 0.0260407624240796 | 0.0939795113181867 | 0.0600292788605657 | OG0000127 | 1 |
| GO:1902074 | response to salt | 1/22 | 16/13359 | 0.0260407624240796 | 0.0939795113181867 | 0.0600292788605657 | OG0000167 | 1 |
| GO:0009559 | embryo sac central cell differentiation | 1/22 | 17/13359 | 0.0276466309114263 | 0.0960200191869966 | 0.0613326503524567 | OG0001956 | 1 |
| GO:0009809 | lignin biosynthetic process | 1/22 | 17/13359 | 0.0276466309114263 | 0.0960200191869966 | 0.0613326503524567 | OG0000124 | 1 |
| GO:0016628 | oxidoreductase activity, acting on the CH-CH group of donors, NAD or NADP as acceptor | 1/22 | 17/13359 | 0.0276466309114263 | 0.0960200191869966 | 0.0613326503524567 | OG0000721 | 1 |
| GO:0016702 | oxidoreductase activity, acting on single donors with incorporation of molecular oxygen, incorporation of two atoms of oxygen | 1/22 | 17/13359 | 0.0276466309114263 | 0.0960200191869966 | 0.0613326503524567 | OG0000108 | 1 |
| GO:0042542 | response to hydrogen peroxide | 1/22 | 17/13359 | 0.0276466309114263 | 0.0960200191869966 | 0.0613326503524567 | OG0000124 | 1 |
| GO:0043487 | regulation of RNA stability | 1/22 | 17/13359 | 0.0276466309114263 | 0.0960200191869966 | 0.0613326503524567 | OG0000167 | 1 |
| GO:1901000 | regulation of response to salt stress | 1/22 | 17/13359 | 0.0276466309114263 | 0.0960200191869966 | 0.0613326503524567 | OG0000124 | 1 |
| GO:0006637 | acyl-CoA metabolic process | 1/22 | 18/13359 | 0.0292499717988456 | 0.0999760940849431 | 0.0638595875530352 | OG0000721 | 1 |
| GO:0010927 | cellular component assembly involved in morphogenesis | 1/22 | 18/13359 | 0.0292499717988456 | 0.0999760940849431 | 0.0638595875530352 | OG0000721 | 1 |
| GO:0035383 | thioester metabolic process | 1/22 | 18/13359 | 0.0292499717988456 | 0.0999760940849431 | 0.0638595875530352 | OG0000721 | 1 |
| GO:0009834 | plant-type secondary cell wall biogenesis | 1/22 | 19/13359 | 0.0308507888755587 | 0.103262225977259 | 0.065958599613957 | OG0000124 | 1 |
| GO:0022891 | | 1/22 | 19/13359 | 0.0308507888755587 | 0.103262225977259 | 0.065958599613957 | OG0000127 | 1 |
| GO:0034605 | cellular response to heat | 1/22 | 19/13359 | 0.0308507888755587 | 0.103262225977259 | 0.065958599613957 | OG0000047 | 1 |
| GO:0061013 | regulation of mRNA catabolic process | 1/22 | 19/13359 | 0.0308507888755587 | 0.103262225977259 | 0.065958599613957 | OG0000167 | 1 |
| GO:0006979 | response to oxidative stress | 2/22 | 170/13359 | 0.0314774760287583 | 0.104816750075144 | 0.0669515496650747 | OG0000124/OG0000167 | 2 |
| GO:0002832 | negative regulation of response to biotic stimulus | 1/22 | 20/13359 | 0.0324490859253892 | 0.105869239938391 | 0.0676238260646069 | OG0000124 | 1 |
| GO:0015103 | inorganic anion transmembrane transporter activity | 1/22 | 20/13359 | 0.0324490859253892 | 0.105869239938391 | 0.0676238260646069 | OG0000127 | 1 |
| GO:0032102 | negative regulation of response to external stimulus | 1/22 | 20/13359 | 0.0324490859253892 | 0.105869239938391 | 0.0676238260646069 | OG0000124 | 1 |
| GO:0047484 | regulation of response to osmotic stress | 1/22 | 20/13359 | 0.0324490859253892 | 0.105869239938391 | 0.0676238260646069 | OG0000124 | 1 |
| GO:0070887 | cellular response to chemical stimulus | 3/22 | 433/13359 | 0.0329665331214176 | 0.107016986916763 | 0.0683569478106094 | OG0000035/OG0000167/OG0001956 | 3 |
| GO:1901566 | organonitrogen compound biosynthetic process | 3/22 | 437/13359 | 0.0337475151354384 | 0.107282848319484 | 0.0685267664025382 | OG0000047/OG0000167/OG0000721 | 3 |
| GO:0048229 | gametophyte development | 2/22 | 177/13359 | 0.0338962811696773 | 0.107282848319484 | 0.0685267664025382 | OG0000721/OG0001956 | 2 |
| GO:0000096 | sulfur amino acid metabolic process | 1/22 | 21/13359 | 0.0340448667267712 | 0.107282848319484 | 0.0685267664025382 | OG0000721 | 1 |
| GO:0002683 | negative regulation of immune system process | 1/22 | 21/13359 | 0.0340448667267712 | 0.107282848319484 | 0.0685267664025382 | OG0000124 | 1 |
| GO:0032989 | cellular component morphogenesis | 1/22 | 21/13359 | 0.0340448667267712 | 0.107282848319484 | 0.0685267664025382 | OG0000721 | 1 |
| GO:0050777 | negative regulation of immune response | 1/22 | 21/13359 | 0.0340448667267712 | 0.107282848319484 | 0.0685267664025382 | OG0000124 | 1 |
| GO:0009808 | lignin metabolic process | 1/22 | 22/13359 | 0.0356381350527581 | 0.110154235617616 | 0.0703608609452591 | OG0000124 | 1 |
| GO:0010200 | response to chitin | 1/22 | 22/13359 | 0.0356381350527581 | 0.110154235617616 | 0.0703608609452591 | OG0000167 | 1 |
| GO:0015995 | chlorophyll biosynthetic process | 1/22 | 22/13359 | 0.0356381350527581 | 0.110154235617616 | 0.0703608609452591 | OG0000047 | 1 |
| GO:0048767 | root hair elongation | 1/22 | 22/13359 | 0.0356381350527581 | 0.110154235617616 | 0.0703608609452591 | OG0000035 | 1 |
| GO:0009624 | response to nematode | 1/22 | 23/13359 | 0.0372288946710259 | 0.110320486043499 | 0.0704670531677554 | OG0000035 | 1 |
| GO:0010114 | response to red light | 1/22 | 23/13359 | 0.0372288946710259 | 0.110320486043499 | 0.0704670531677554 | OG0000047 | 1 |
| GO:0010181 | FMN binding | 1/22 | 23/13359 | 0.0372288946710259 | 0.110320486043499 | 0.0704670531677554 | OG0000253 | 1 |
| GO:0017148 | negative regulation of translation | 1/22 | 23/13359 | 0.0372288946710259 | 0.110320486043499 | 0.0704670531677554 | OG0000167 | 1 |
| GO:0033865 | nucleoside bisphosphate metabolic process | 1/22 | 23/13359 | 0.0372288946710259 | 0.110320486043499 | 0.0704670531677554 | OG0000721 | 1 |
| GO:0033875 | ribonucleoside bisphosphate metabolic process | 1/22 | 23/13359 | 0.0372288946710259 | 0.110320486043499 | 0.0704670531677554 | OG0000721 | 1 |
| GO:0034032 | purine nucleoside bisphosphate metabolic process | 1/22 | 23/13359 | 0.0372288946710259 | 0.110320486043499 | 0.0704670531677554 | OG0000721 | 1 |
| GO:0034249 | negative regulation of cellular amide metabolic process | 1/22 | 23/13359 | 0.0372288946710259 | 0.110320486043499 | 0.0704670531677554 | OG0000167 | 1 |
| GO:0042910 | xenobiotic transmembrane transporter activity | 1/22 | 23/13359 | 0.0372288946710259 | 0.110320486043499 | 0.0704670531677554 | OG0000035 | 1 |
| GO:0007165 | signal transduction | 3/22 | 459/13359 | 0.0382176377873821 | 0.110466425005062 | 0.0705602714713774 | OG0000035/OG0000167/OG0000691 | 3 |
| GO:0009719 | response to endogenous stimulus | 3/22 | 459/13359 | 0.0382176377873821 | 0.110466425005062 | 0.0705602714713774 | OG0000035/OG0000167/OG0001956 | 3 |
| GO:0009725 | response to hormone | 3/22 | 461/13359 | 0.0386386243062403 | 0.110466425005062 | 0.0705602714713774 | OG0000035/OG0000167/OG0001956 | 3 |
| GO:0000932 | P-body | 1/22 | 24/13359 | 0.038817149343884 | 0.110466425005062 | 0.0705602714713774 | OG0000167 | 1 |
| GO:0009522 | photosystem I | 1/22 | 24/13359 | 0.038817149343884 | 0.110466425005062 | 0.0705602714713774 | OG0000047 | 1 |
| GO:0009813 | flavonoid biosynthetic process | 1/22 | 24/13359 | 0.038817149343884 | 0.110466425005062 | 0.0705602714713774 | OG0000124 | 1 |
| GO:0018958 | phenol-containing compound metabolic process | 1/22 | 24/13359 | 0.038817149343884 | 0.110466425005062 | 0.0705602714713774 | OG0000124 | 1 |
| GO:0022892 | | 1/22 | 24/13359 | 0.038817149343884 | 0.110466425005062 | 0.0705602714713774 | OG0000127 | 1 |
| GO:1903311 | regulation of mRNA metabolic process | 1/22 | 24/13359 | 0.038817149343884 | 0.110466425005062 | 0.0705602714713774 | OG0000167 | 1 |
| GO:0048827 | phyllome development | 2/22 | 192/13359 | 0.039317118557115 | 0.111398502578493 | 0.0711556346924704 | OG0000035/OG0000167 | 2 |
| GO:0022622 | root system development | 2/22 | 195/13359 | 0.0404390123667152 | 0.114076864580341 | 0.0728664346024011 | OG0000035/OG0000124 | 2 |
| GO:0048364 | root development | 2/22 | 196/13359 | 0.0408157067684074 | 0.114638898140831 | 0.0732254327378294 | OG0000035/OG0000124 | 2 |
| GO:0006779 | porphyrin-containing compound biosynthetic process | 1/22 | 26/13359 | 0.0419861588758128 | 0.11690973549041 | 0.0746759268571625 | OG0000047 | 1 |
| GO:0042626 | ATPase-coupled transmembrane transporter activity | 1/22 | 26/13359 | 0.0419861588758128 | 0.11690973549041 | 0.0746759268571625 | OG0000035 | 1 |
| GO:0015698 | inorganic anion transport | 1/22 | 27/13359 | 0.0435669212327259 | 0.119255216594665 | 0.07617409956837 | OG0000127 | 1 |
| GO:0016052 | carbohydrate catabolic process | 1/22 | 27/13359 | 0.0435669212327259 | 0.119255216594665 | 0.07617409956837 | OG0000124 | 1 |
| GO:0036464 | cytoplasmic ribonucleoprotein granule | 1/22 | 27/13359 | 0.0435669212327259 | 0.119255216594665 | 0.07617409956837 | OG0000167 | 1 |
| GO:0043531 | ADP binding | 1/22 | 27/13359 | 0.0435669212327259 | 0.119255216594665 | 0.07617409956837 | OG0000691 | 1 |
| GO:0009812 | flavonoid metabolic process | 1/22 | 28/13359 | 0.0451451936399329 | 0.120015617660068 | 0.0766598046647333 | OG0000124 | 1 |
| GO:0022853 | active ion transmembrane transporter activity | 1/22 | 28/13359 | 0.0451451936399329 | 0.120015617660068 | 0.0766598046647333 | OG0000127 | 1 |
| GO:0031348 | negative regulation of defense response | 1/22 | 28/13359 | 0.0451451936399329 | 0.120015617660068 | 0.0766598046647333 | OG0000124 | 1 |
| GO:0032535 | regulation of cellular component size | 1/22 | 28/13359 | 0.0451451936399329 | 0.120015617660068 | 0.0766598046647333 | OG0000035 | 1 |
| GO:0033014 | tetrapyrrole biosynthetic process | 1/22 | 28/13359 | 0.0451451936399329 | 0.120015617660068 | 0.0766598046647333 | OG0000047 | 1 |
| GO:0035770 | ribonucleoprotein granule | 1/22 | 28/13359 | 0.0451451936399329 | 0.120015617660068 | 0.0766598046647333 | OG0000167 | 1 |
| GO:0090066 | regulation of anatomical structure size | 1/22 | 28/13359 | 0.0451451936399329 | 0.120015617660068 | 0.0766598046647333 | OG0000035 | 1 |
| GO:0007154 | cell communication | 3/22 | 493/13359 | 0.045702752793241 | 0.120999911083745 | 0.0772885206857224 | OG0000035/OG0000127/OG0000167 | 3 |
| GO:0009926 | auxin polar transport | 1/22 | 29/13359 | 0.0467209798330106 | 0.122690052732215 | 0.0783680962539159 | OG0000035 | 1 |
| GO:0031331 | positive regulation of cellular catabolic process | 1/22 | 29/13359 | 0.0467209798330106 | 0.122690052732215 | 0.0783680962539159 | OG0000167 | 1 |
| GO:0005215 | transporter activity | 2/22 | 212/13359 | 0.0470239013309599 | 0.122985588096357 | 0.0785568690463937 | OG0000035/OG0000127 | 2 |
| GO:0006767 | water-soluble vitamin metabolic process | 1/22 | 30/13359 | 0.0482942835422134 | 0.12479242867308 | 0.0797109858886218 | OG0000124 | 1 |
| GO:0006997 | nucleus organization | 1/22 | 30/13359 | 0.0482942835422134 | 0.12479242867308 | 0.0797109858886218 | OG0001956 | 1 |
| GO:0010119 | regulation of stomatal movement | 1/22 | 30/13359 | 0.0482942835422134 | 0.12479242867308 | 0.0797109858886218 | OG0001956 | 1 |
| GO:0055085 | transmembrane transport | 3/22 | 508/13359 | 0.0492248392293559 | 0.12669022367396 | 0.0809231997395995 | OG0000035/OG0000127/OG0000189 | 3 |
| GO:0009630 | gravitropism | 1/22 | 31/13359 | 0.0498651084924778 | 0.126822283803703 | 0.0810075529591849 | OG0000035 | 1 |
| GO:0009896 | positive regulation of catabolic process | 1/22 | 31/13359 | 0.0498651084924778 | 0.126822283803703 | 0.0810075529591849 | OG0000167 | 1 |
| GO:0015994 | chlorophyll metabolic process | 1/22 | 31/13359 | 0.0498651084924778 | 0.126822283803703 | 0.0810075529591849 | OG0000047 | 1 |
| GO:0009266 | response to temperature stimulus | 2/22 | 221/13359 | 0.0506610009373764 | 0.126938857435577 | 0.0810820142003361 | OG0000047/OG0000167 | 2 |
| GO:1901700 | response to oxygen-containing compound | 3/22 | 514/13359 | 0.0506709098297823 | 0.126938857435577 | 0.0810820142003361 | OG0000124/OG0000167/OG0001956 | 3 |
| GO:0009561 | megagametogenesis | 1/22 | 32/13359 | 0.0514334584034297 | 0.126938857435577 | 0.0810820142003361 | OG0001956 | 1 |
| GO:0046165 | alcohol biosynthetic process | 1/22 | 32/13359 | 0.0514334584034297 | 0.126938857435577 | 0.0810820142003361 | OG0000721 | 1 |
| GO:0071489 | cellular response to red or far red light | 1/22 | 32/13359 | 0.0514334584034297 | 0.126938857435577 | 0.0810820142003361 | OG0000047 | 1 |
| GO:0080147 | root hair cell development | 1/22 | 32/13359 | 0.0514334584034297 | 0.126938857435577 | 0.0810820142003361 | OG0000035 | 1 |
| GO:0098656 | anion transmembrane transport | 1/22 | 32/13359 | 0.0514334584034297 | 0.126938857435577 | 0.0810820142003361 | OG0000127 | 1 |
| GO:0016887 | ATP hydrolysis activity | 2/22 | 223/13359 | 0.0514829421797542 | 0.126938857435577 | 0.0810820142003361 | OG0000035/OG0000538 | 2 |
| GO:0006778 | porphyrin-containing compound metabolic process | 1/22 | 33/13359 | 0.0529993369893921 | 0.128230605599803 | 0.0819071164984888 | OG0000047 | 1 |
| GO:0044528 | regulation of mitochondrial mRNA stability | 1/22 | 33/13359 | 0.0529993369893921 | 0.128230605599803 | 0.0819071164984888 | OG0000731 | 1 |
| GO:0046658 | anchored component of plasma membrane | 1/22 | 33/13359 | 0.0529993369893921 | 0.128230605599803 | 0.0819071164984888 | OG0000124 | 1 |
| GO:0048284 | organelle fusion | 1/22 | 33/13359 | 0.0529993369893921 | 0.128230605599803 | 0.0819071164984888 | OG0001956 | 1 |
| GO:0060918 | auxin transport | 1/22 | 33/13359 | 0.0529993369893921 | 0.128230605599803 | 0.0819071164984888 | OG0000035 | 1 |
| GO:0009521 | photosystem | 1/22 | 34/13359 | 0.0545627479593936 | 0.129111850482668 | 0.0824700104109593 | OG0000047 | 1 |
| GO:0009606 | tropism | 1/22 | 34/13359 | 0.0545627479593936 | 0.129111850482668 | 0.0824700104109593 | OG0000035 | 1 |
| GO:0009629 | response to gravity | 1/22 | 34/13359 | 0.0545627479593936 | 0.129111850482668 | 0.0824700104109593 | OG0000035 | 1 |
| GO:0009644 | response to high light intensity | 1/22 | 34/13359 | 0.0545627479593936 | 0.129111850482668 | 0.0824700104109593 | OG0000047 | 1 |
| GO:0009914 | hormone transport | 1/22 | 34/13359 | 0.0545627479593936 | 0.129111850482668 | 0.0824700104109593 | OG0000035 | 1 |
| GO:0033013 | tetrapyrrole metabolic process | 1/22 | 34/13359 | 0.0545627479593936 | 0.129111850482668 | 0.0824700104109593 | OG0000047 | 1 |
| GO:0009755 | hormone-mediated signaling pathway | 2/22 | 231/13359 | 0.0548193106053435 | 0.129245527923547 | 0.0825553966857267 | OG0000035/OG0000167 | 2 |
| GO:0002684 | positive regulation of immune system process | 1/22 | 35/13359 | 0.0561236950171715 | 0.130416931586665 | 0.0833036291053814 | OG0000124 | 1 |
| GO:0002833 | positive regulation of response to biotic stimulus | 1/22 | 35/13359 | 0.0561236950171715 | 0.130416931586665 | 0.0833036291053814 | OG0000124 | 1 |
| GO:0010224 | response to UV-B | 1/22 | 35/13359 | 0.0561236950171715 | 0.130416931586665 | 0.0833036291053814 | OG0000047 | 1 |
| GO:0050778 | positive regulation of immune response | 1/22 | 35/13359 | 0.0561236950171715 | 0.130416931586665 | 0.0833036291053814 | OG0000124 | 1 |
| GO:0006766 | vitamin metabolic process | 1/22 | 36/13359 | 0.0576821818611839 | 0.132137196745833 | 0.0844024460230836 | OG0000124 | 1 |
| GO:0048443 | stamen development | 1/22 | 36/13359 | 0.0576821818611839 | 0.132137196745833 | 0.0844024460230836 | OG0000035 | 1 |
| GO:0048466 | androecium development | 1/22 | 36/13359 | 0.0576821818611839 | 0.132137196745833 | 0.0844024460230836 | OG0000035 | 1 |
| GO:0048582 | positive regulation of post-embryonic development | 1/22 | 36/13359 | 0.0576821818611839 | 0.132137196745833 | 0.0844024460230836 | OG0000047 | 1 |
| GO:0003700 | DNA-binding transcription factor activity | 2/22 | 239/13359 | 0.0582316488829913 | 0.132575730024017 | 0.0846825585292725 | OG0000018/OG0000167 | 2 |
| GO:0050789 | regulation of biological process | 5/22 | 1313/13359 | 0.058284067069382 | 0.132575730024017 | 0.0846825585292725 | OG0000035/OG0000047/OG0000124/OG0000167/OG0001956 | 5 |
| GO:0032103 | positive regulation of response to external stimulus | 1/22 | 38/13359 | 0.0607917896753761 | 0.136941505396541 | 0.0874711913238456 | OG0000124 | 1 |
| GO:0032870 | cellular response to hormone stimulus | 2/22 | 245/13359 | 0.0608393375368534 | 0.136941505396541 | 0.0874711913238456 | OG0000035/OG0000167 | 2 |
| GO:0071495 | cellular response to endogenous stimulus | 2/22 | 245/13359 | 0.0608393375368534 | 0.136941505396541 | 0.0874711913238456 | OG0000035/OG0000167 | 2 |
| GO:0043603 | cellular amide metabolic process | 2/22 | 248/13359 | 0.062158402602626 | 0.137923030953487 | 0.0880981393739073 | OG0000167/OG0000721 | 2 |
| GO:0005516 | calmodulin binding | 1/22 | 39/13359 | 0.0623429180161272 | 0.137923030953487 | 0.0880981393739073 | OG0001956 | 1 |
| GO:0009640 | photomorphogenesis | 1/22 | 39/13359 | 0.0623429180161272 | 0.137923030953487 | 0.0880981393739073 | OG0000035 | 1 |
| GO:0031349 | positive regulation of defense response | 1/22 | 39/13359 | 0.0623429180161272 | 0.137923030953487 | 0.0880981393739073 | OG0000124 | 1 |
| GO:0051537 | 2 iron, 2 sulfur cluster binding | 1/22 | 39/13359 | 0.0623429180161272 | 0.137923030953487 | 0.0880981393739073 | OG0000151 | 1 |
| GO:0009867 | jasmonic acid mediated signaling pathway | 1/22 | 40/13359 | 0.0638916008842685 | 0.139497993147508 | 0.0891041442297914 | OG0000167 | 1 |
| GO:0048764 | trichoblast maturation | 1/22 | 40/13359 | 0.0638916008842685 | 0.139497993147508 | 0.0891041442297914 | OG0000035 | 1 |
| GO:0048765 | root hair cell differentiation | 1/22 | 40/13359 | 0.0638916008842685 | 0.139497993147508 | 0.0891041442297914 | OG0000035 | 1 |
| GO:0010033 | response to organic substance | 3/22 | 567/13359 | 0.0643504674271784 | 0.139497993147508 | 0.0891041442297914 | OG0000035/OG0000167/OG0001956 | 3 |
| GO:0031967 | organelle envelope | 3/22 | 567/13359 | 0.0643504674271784 | 0.139497993147508 | 0.0891041442297914 | OG0000047/OG0000127/OG0000721 | 3 |
| GO:0031975 | envelope | 3/22 | 567/13359 | 0.0643504674271784 | 0.139497993147508 | 0.0891041442297914 | OG0000047/OG0000127/OG0000721 | 3 |
| GO:0048469 | cell maturation | 1/22 | 41/13359 | 0.0654378419519572 | 0.140487475962427 | 0.0897361749670382 | OG0000035 | 1 |
| GO:0009791 | post-embryonic development | 3/22 | 573/13359 | 0.0659996377067537 | 0.140487475962427 | 0.0897361749670382 | OG0000035/OG0000047/OG0000167 | 3 |
| GO:0048583 | regulation of response to stimulus | 2/22 | 257/13359 | 0.0661750306982961 | 0.140487475962427 | 0.0897361749670382 | OG0000035/OG0000124 | 2 |
| GO:0009941 | chloroplast envelope | 2/22 | 258/13359 | 0.0666267298641929 | 0.140487475962427 | 0.0897361749670382 | OG0000127/OG0000721 | 2 |
| GO:0006402 | mRNA catabolic process | 1/22 | 42/13359 | 0.0669816448861107 | 0.140487475962427 | 0.0897361749670382 | OG0000167 | 1 |
| GO:0009832 | plant-type cell wall biogenesis | 1/22 | 42/13359 | 0.0669816448861107 | 0.140487475962427 | 0.0897361749670382 | OG0000124 | 1 |
| GO:0010054 | trichoblast differentiation | 1/22 | 42/13359 | 0.0669816448861107 | 0.140487475962427 | 0.0897361749670382 | OG0000035 | 1 |
| GO:0015291 | secondary active transmembrane transporter activity | 1/22 | 42/13359 | 0.0669816448861107 | 0.140487475962427 | 0.0897361749670382 | OG0000127 | 1 |
| GO:0015399 | primary active transmembrane transporter activity | 1/22 | 42/13359 | 0.0669816448861107 | 0.140487475962427 | 0.0897361749670382 | OG0000035 | 1 |
| GO:0071395 | cellular response to jasmonic acid stimulus | 1/22 | 42/13359 | 0.0669816448861107 | 0.140487475962427 | 0.0897361749670382 | OG0000167 | 1 |
| GO:0032269 | negative regulation of cellular protein metabolic process | 1/22 | 43/13359 | 0.068523013348415 | 0.141877777638065 | 0.0906242281800901 | OG0000167 | 1 |
| GO:0046395 | carboxylic acid catabolic process | 1/22 | 43/13359 | 0.068523013348415 | 0.141877777638065 | 0.0906242281800901 | OG0000124 | 1 |
| GO:0048527 | lateral root development | 1/22 | 43/13359 | 0.068523013348415 | 0.141877777638065 | 0.0906242281800901 | OG0000035 | 1 |
| GO:0051248 | negative regulation of protein metabolic process | 1/22 | 43/13359 | 0.068523013348415 | 0.141877777638065 | 0.0906242281800901 | OG0000167 | 1 |
| GO:0050793 | regulation of developmental process | 2/22 | 263/13359 | 0.0689011187323856 | 0.142204864859812 | 0.0908331546766274 | OG0000047/OG0000167 | 2 |
| GO:0015297 | antiporter activity | 1/22 | 44/13359 | 0.0700619509953314 | 0.143227912477798 | 0.0914866248839771 | OG0000127 | 1 |
| GO:0051240 | positive regulation of multicellular organismal process | 1/22 | 44/13359 | 0.0700619509953314 | 0.143227912477798 | 0.0914866248839771 | OG0000047 | 1 |
| GO:0071398 | cellular response to fatty acid | 1/22 | 44/13359 | 0.0700619509953314 | 0.143227912477798 | 0.0914866248839771 | OG0000167 | 1 |
| GO:0031329 | regulation of cellular catabolic process | 1/22 | 45/13359 | 0.0715984614781027 | 0.145907274810266 | 0.0931980637536651 | OG0000167 | 1 |
| GO:0042170 | plastid membrane | 2/22 | 271/13359 | 0.072594188484233 | 0.147471213084322 | 0.0941970270963897 | OG0000047/OG0000127 | 2 |
| GO:0016627 | oxidoreductase activity, acting on the CH-CH group of donors | 1/22 | 47/13359 | 0.0746642155301401 | 0.149872587397948 | 0.0957309015153912 | OG0000721 | 1 |
| GO:0048581 | negative regulation of post-embryonic development | 1/22 | 47/13359 | 0.0746642155301401 | 0.149872587397948 | 0.0957309015153912 | OG0000167 | 1 |
| GO:0016020 | membrane | 8/22 | 2831/13359 | 0.074969258425108 | 0.149872587397948 | 0.0957309015153912 | OG0000035/OG0000047/OG0000124/OG0000127/OG0000167/OG0000189/OG0000721/OG0000785 | 8 |
| GO:0046872 | metal ion binding | 4/22 | 992/13359 | 0.075737506213207 | 0.149872587397948 | 0.0957309015153912 | OG0000108/OG0000124/OG0000167/OG0000006 | 4 |
| GO:0009416 | response to light stimulus | 2/22 | 278/13359 | 0.0758786984584043 | 0.149872587397948 | 0.0957309015153912 | OG0000035/OG0000047 | 2 |
| GO:0005996 | monosaccharide metabolic process | 1/22 | 48/13359 | 0.0761934663758687 | 0.149872587397948 | 0.0957309015153912 | OG0000124 | 1 |
| GO:0010053 | root epidermal cell differentiation | 1/22 | 48/13359 | 0.0761934663758687 | 0.149872587397948 | 0.0957309015153912 | OG0000035 | 1 |
| GO:0018130 | heterocycle biosynthetic process | 3/22 | 609/13359 | 0.0763100086119999 | 0.149872587397948 | 0.0957309015153912 | OG0000047/OG0000124/OG0000167 | 3 |
| GO:0071310 | cellular response to organic substance | 2/22 | 279/13359 | 0.0763518680054615 | 0.149872587397948 | 0.0957309015153912 | OG0000035/OG0000167 | 2 |
| GO:0000302 | response to reactive oxygen species | 1/22 | 49/13359 | 0.0777203046103909 | 0.149872587397948 | 0.0957309015153912 | OG0000124 | 1 |
| GO:0006066 | alcohol metabolic process | 1/22 | 49/13359 | 0.0777203046103909 | 0.149872587397948 | 0.0957309015153912 | OG0000721 | 1 |
| GO:0009267 | cellular response to starvation | 1/22 | 49/13359 | 0.0777203046103909 | 0.149872587397948 | 0.0957309015153912 | OG0000127 | 1 |
| GO:0009735 | response to cytokinin | 1/22 | 49/13359 | 0.0777203046103909 | 0.149872587397948 | 0.0957309015153912 | OG0000035 | 1 |
| GO:0016054 | organic acid catabolic process | 1/22 | 49/13359 | 0.0777203046103909 | 0.149872587397948 | 0.0957309015153912 | OG0000124 | 1 |
| GO:0046148 | pigment biosynthetic process | 1/22 | 49/13359 | 0.0777203046103909 | 0.149872587397948 | 0.0957309015153912 | OG0000047 | 1 |
| GO:0098660 | inorganic ion transmembrane transport | 1/22 | 49/13359 | 0.0777203046103909 | 0.149872587397948 | 0.0957309015153912 | OG0000127 | 1 |
| GO:1901607 | alpha-amino acid biosynthetic process | 1/22 | 49/13359 | 0.0777203046103909 | 0.149872587397948 | 0.0957309015153912 | OG0000721 | 1 |
| GO:0006401 | RNA catabolic process | 1/22 | 50/13359 | 0.0792447338589691 | 0.151455911458266 | 0.0967422475014509 | OG0000167 | 1 |
| GO:0048528 | post-embryonic root development | 1/22 | 50/13359 | 0.0792447338589691 | 0.151455911458266 | 0.0967422475014509 | OG0000035 | 1 |
| GO:0140359 | ABC-type transporter activity | 1/22 | 50/13359 | 0.0792447338589691 | 0.151455911458266 | 0.0967422475014509 | OG0000035 | 1 |
| GO:0009314 | response to radiation | 2/22 | 287/13359 | 0.0801720113565946 | 0.152559431289854 | 0.0974471192205034 | OG0000035/OG0000047 | 2 |
| GO:0065007 | biological regulation | 5/22 | 1439/13359 | 0.0802961934543794 | 0.152559431289854 | 0.0974471192205034 | OG0000035/OG0000047/OG0000124/OG0000167/OG0001956 | 5 |
| GO:0044272 | sulfur compound biosynthetic process | 1/22 | 51/13359 | 0.0807667577416876 | 0.152559431289854 | 0.0974471192205034 | OG0000721 | 1 |
| GO:0051241 | negative regulation of multicellular organismal process | 1/22 | 51/13359 | 0.0807667577416876 | 0.152559431289854 | 0.0974471192205034 | OG0000167 | 1 |
| GO:0019752 | carboxylic acid metabolic process | 2/22 | 289/13359 | 0.0811365595600463 | 0.152811129667026 | 0.0976078912000557 | OG0000124/OG0000721 | 2 |
| GO:0071840 | cellular component organization or biogenesis | 4/22 | 1019/13359 | 0.0819011577695365 | 0.153802755578839 | 0.0982412908373877 | OG0000035/OG0000124/OG0000721/OG0001956 | 4 |
| GO:0065008 | regulation of biological quality | 2/22 | 292/13359 | 0.0825903836928051 | 0.154647501059571 | 0.098780870808786 | OG0000035/OG0000167 | 2 |
| GO:0008509 | anion transmembrane transporter activity | 1/22 | 53/13359 | 0.0838036038640543 | 0.155566460046492 | 0.0993678545514497 | OG0000127 | 1 |
| GO:0031669 | cellular response to nutrient levels | 1/22 | 53/13359 | 0.0838036038640543 | 0.155566460046492 | 0.0993678545514497 | OG0000127 | 1 |
| GO:0042546 | cell wall biogenesis | 1/22 | 53/13359 | 0.0838036038640543 | 0.155566460046492 | 0.0993678545514497 | OG0000124 | 1 |
| GO:0098542 | defense response to other organism | 2/22 | 295/13359 | 0.0840525131452163 | 0.155581442669942 | 0.0993774246808167 | OG0000124/OG0000167 | 2 |
| GO:0009894 | regulation of catabolic process | 1/22 | 54/13359 | 0.0853184333180597 | 0.157024808898765 | 0.100299372801558 | OG0000167 | 1 |
| GO:0080167 | response to karrikin | 1/22 | 54/13359 | 0.0853184333180597 | 0.157024808898765 | 0.100299372801558 | OG0000047 | 1 |
| GO:0043436 | oxoacid metabolic process | 2/22 | 299/13359 | 0.0860147514535385 | 0.157856617724392 | 0.100830689502952 | OG0000124/OG0000721 | 2 |
| GO:0071482 | cellular response to light stimulus | 1/22 | 55/13359 | 0.086830871834933 | 0.158902955255996 | 0.101499036109419 | OG0000047 | 1 |
| GO:0002376 | immune system process | 2/22 | 302/13359 | 0.0874958402939676 | 0.15921778261944 | 0.101700131638948 | OG0000124/OG0000167 | 2 |
| GO:0006955 | immune response | 2/22 | 302/13359 | 0.0874958402939676 | 0.15921778261944 | 0.101700131638948 | OG0000124/OG0000167 | 2 |
| GO:0009150 | purine ribonucleotide metabolic process | 1/22 | 56/13359 | 0.0883409230089887 | 0.159408481183818 | 0.101821940075047 | OG0000721 | 1 |
| GO:0010150 | leaf senescence | 1/22 | 56/13359 | 0.0883409230089887 | 0.159408481183818 | 0.101821940075047 | OG0000167 | 1 |
| GO:0071478 | cellular response to radiation | 1/22 | 56/13359 | 0.0883409230089887 | 0.159408481183818 | 0.101821940075047 | OG0000047 | 1 |
| GO:0010243 | response to organonitrogen compound | 1/22 | 57/13359 | 0.089848590429405 | 0.16122830393721 | 0.102984349264113 | OG0000167 | 1 |
| GO:0090693 | plant organ senescence | 1/22 | 57/13359 | 0.089848590429405 | 0.16122830393721 | 0.102984349264113 | OG0000167 | 1 |
| GO:0009734 | auxin-activated signaling pathway | 1/22 | 58/13359 | 0.091353877680236 | 0.161242090113203 | 0.102993155164373 | OG0000035 | 1 |
| GO:0010118 | stomatal movement | 1/22 | 58/13359 | 0.091353877680236 | 0.161242090113203 | 0.102993155164373 | OG0001956 | 1 |
| GO:0031225 | anchored component of membrane | 1/22 | 58/13359 | 0.091353877680236 | 0.161242090113203 | 0.102993155164373 | OG0000124 | 1 |
| GO:0051093 | negative regulation of developmental process | 1/22 | 58/13359 | 0.091353877680236 | 0.161242090113203 | 0.102993155164373 | OG0000167 | 1 |
| GO:0051094 | positive regulation of developmental process | 1/22 | 58/13359 | 0.091353877680236 | 0.161242090113203 | 0.102993155164373 | OG0000047 | 1 |
| GO:0090627 | plant epidermal cell differentiation | 1/22 | 58/13359 | 0.091353877680236 | 0.161242090113203 | 0.102993155164373 | OG0000035 | 1 |
| GO:0008652 | cellular amino acid biosynthetic process | 1/22 | 59/13359 | 0.0928567883404143 | 0.162801966354959 | 0.103989523889757 | OG0000721 | 1 |
| GO:0009642 | response to light intensity | 1/22 | 59/13359 | 0.0928567883404143 | 0.162801966354959 | 0.103989523889757 | OG0000047 | 1 |
| GO:0006082 | organic acid metabolic process | 2/22 | 313/13359 | 0.092993692856006 | 0.162801966354959 | 0.103989523889757 | OG0000124/OG0000721 | 2 |
| GO:0034248 | regulation of cellular amide metabolic process | 1/22 | 60/13359 | 0.094357325983761 | 0.163816532379969 | 0.104637576491686 | OG0000167 | 1 |
| GO:0034655 | nucleobase-containing compound catabolic process | 1/22 | 60/13359 | 0.094357325983761 | 0.163816532379969 | 0.104637576491686 | OG0000167 | 1 |
| GO:0048367 | shoot system development | 2/22 | 317/13359 | 0.095018393818426 | 0.163816532379969 | 0.104637576491686 | OG0000035/OG0000167 | 2 |
| GO:0048731 | system development | 3/22 | 671/13359 | 0.0956652762501082 | 0.163816532379969 | 0.104637576491686 | OG0000035/OG0000124/OG0000167 | 3 |
| GO:0006163 | purine nucleotide metabolic process | 1/22 | 61/13359 | 0.0958554941789909 | 0.163816532379969 | 0.104637576491686 | OG0000721 | 1 |
| GO:0006417 | regulation of translation | 1/22 | 61/13359 | 0.0958554941789909 | 0.163816532379969 | 0.104637576491686 | OG0000167 | 1 |
| GO:0009523 | photosystem II | 1/22 | 61/13359 | 0.0958554941789909 | 0.163816532379969 | 0.104637576491686 | OG0000047 | 1 |
| GO:0042440 | pigment metabolic process | 1/22 | 61/13359 | 0.0958554941789909 | 0.163816532379969 | 0.104637576491686 | OG0000047 | 1 |
| GO:0042594 | response to starvation | 1/22 | 61/13359 | 0.0958554941789909 | 0.163816532379969 | 0.104637576491686 | OG0000127 | 1 |
| GO:0009259 | ribonucleotide metabolic process | 1/22 | 62/13359 | 0.0973512964897187 | 0.165062828168919 | 0.105433646149937 | OG0000721 | 1 |
| GO:0009553 | embryo sac development | 1/22 | 62/13359 | 0.0973512964897187 | 0.165062828168919 | 0.105433646149937 | OG0001956 | 1 |
| GO:0015318 | inorganic molecular entity transmembrane transporter activity | 1/22 | 62/13359 | 0.0973512964897187 | 0.165062828168919 | 0.105433646149937 | OG0000127 | 1 |
| GO:0033993 | response to lipid | 2/22 | 323/13359 | 0.0980801251768188 | 0.165863248335667 | 0.105944913390226 | OG0000167/OG0001956 | 2 |
| GO:0048438 | floral whorl development | 1/22 | 63/13359 | 0.098844736474469 | 0.166285676464862 | 0.10621473875546 | OG0000035 | 1 |
| GO:0071365 | cellular response to auxin stimulus | 1/22 | 63/13359 | 0.098844736474469 | 0.166285676464862 | 0.10621473875546 | OG0000035 | 1 |
| GO:0044248 | cellular catabolic process | 2/22 | 327/13359 | 0.100137381650628 | 0.167919529081854 | 0.107258359785052 | OG0000124/OG0000167 | 2 |
| GO:0044765 | | 1/22 | 64/13359 | 0.10033581768668 | 0.167919529081854 | 0.107258359785052 | OG0000127 | 1 |
| GO:0046914 | transition metal ion binding | 2/22 | 329/13359 | 0.101170751165286 | 0.168879341738436 | 0.107871438750965 | OG0000124/OG0000167 | 2 |
| GO:0009653 | anatomical structure morphogenesis | 2/22 | 331/13359 | 0.102207242139654 | 0.17016978974798 | 0.108695710577168 | OG0000035/OG0000721 | 2 |
| GO:0005975 | carbohydrate metabolic process | 2/22 | 332/13359 | 0.102726648992324 | 0.170252176061936 | 0.108748334717743 | OG0000039/OG0000124 | 2 |
| GO:0019693 | ribose phosphate metabolic process | 1/22 | 66/13359 | 0.103310917981856 | 0.170252176061936 | 0.108748334717743 | OG0000721 | 1 |
| GO:0048588 | developmental cell growth | 1/22 | 66/13359 | 0.103310917981856 | 0.170252176061936 | 0.108748334717743 | OG0000035 | 1 |
| GO:1902578 | | 1/22 | 66/13359 | 0.103310917981856 | 0.170252176061936 | 0.108748334717743 | OG0000127 | 1 |
| GO:1901575 | organic substance catabolic process | 2/22 | 334/13359 | 0.103767767143245 | 0.170569917492459 | 0.108951291603461 | OG0000124/OG0000167 | 2 |
| GO:0006820 | anion transport | 1/22 | 67/13359 | 0.104794944146334 | 0.171386161819067 | 0.109472666503298 | OG0000127 | 1 |
| GO:0071669 | plant-type cell wall organization or biogenesis | 1/22 | 67/13359 | 0.104794944146334 | 0.171386161819067 | 0.109472666503298 | OG0000124 | 1 |
| GO:0031667 | response to nutrient levels | 1/22 | 68/13359 | 0.106276625701312 | 0.172499246741326 | 0.110183647910379 | OG0000127 | 1 |
| GO:0031668 | cellular response to extracellular stimulus | 1/22 | 68/13359 | 0.106276625701312 | 0.172499246741326 | 0.110183647910379 | OG0000127 | 1 |
| GO:0090696 | post-embryonic plant organ development | 1/22 | 68/13359 | 0.106276625701312 | 0.172499246741326 | 0.110183647910379 | OG0000035 | 1 |
| GO:0005507 | copper ion binding | 1/22 | 70/13359 | 0.109232969090195 | 0.176853378526982 | 0.112964843380441 | OG0000124 | 1 |
| GO:0009636 | response to toxic substance | 1/22 | 71/13359 | 0.110707637965206 | 0.177824239641588 | 0.113584979533163 | OG0000721 | 1 |
| GO:0009753 | response to jasmonic acid | 1/22 | 71/13359 | 0.110707637965206 | 0.177824239641588 | 0.113584979533163 | OG0000167 | 1 |
| GO:0072521 | purine-containing compound metabolic process | 1/22 | 71/13359 | 0.110707637965206 | 0.177824239641588 | 0.113584979533163 | OG0000721 | 1 |
| GO:0050794 | regulation of cellular process | 4/22 | 1134/13359 | 0.110933697485387 | 0.177824239641588 | 0.113584979533163 | OG0000035/OG0000047/OG0000167/OG0001956 | 4 |
| GO:0004497 | monooxygenase activity | 1/22 | 73/13359 | 0.113649987641395 | 0.179068029308149 | 0.114379448409312 | OG0000104 | 1 |
| GO:0007568 | aging | 1/22 | 73/13359 | 0.113649987641395 | 0.179068029308149 | 0.114379448409312 | OG0000167 | 1 |
| GO:0048437 | floral organ development | 1/22 | 73/13359 | 0.113649987641395 | 0.179068029308149 | 0.114379448409312 | OG0000035 | 1 |
| GO:0070542 | response to fatty acid | 1/22 | 73/13359 | 0.113649987641395 | 0.179068029308149 | 0.114379448409312 | OG0000167 | 1 |
| GO:0071496 | cellular response to external stimulus | 1/22 | 73/13359 | 0.113649987641395 | 0.179068029308149 | 0.114379448409312 | OG0000127 | 1 |
| GO:0071695 | anatomical structure maturation | 1/22 | 73/13359 | 0.113649987641395 | 0.179068029308149 | 0.114379448409312 | OG0000035 | 1 |
| GO:1901605 | alpha-amino acid metabolic process | 1/22 | 73/13359 | 0.113649987641395 | 0.179068029308149 | 0.114379448409312 | OG0000721 | 1 |
| GO:0044249 | cellular biosynthetic process | 4/22 | 1147/13359 | 0.114488644675065 | 0.17995052180071 | 0.114943139230696 | OG0000047/OG0000124/OG0000167/OG0000721 | 4 |
| GO:0044282 | small molecule catabolic process | 1/22 | 74/13359 | 0.11511767545352 | 0.180500044521781 | 0.115294146085283 | OG0000124 | 1 |
| GO:0002831 | regulation of response to biotic stimulus | 1/22 | 75/13359 | 0.116583043247274 | 0.181914603714345 | 0.116197693752686 | OG0000124 | 1 |
| GO:0046700 | heterocycle catabolic process | 1/22 | 75/13359 | 0.116583043247274 | 0.181914603714345 | 0.116197693752686 | OG0000167 | 1 |
| GO:0006952 | defense response | 2/22 | 359/13359 | 0.117029683475053 | 0.182171507288878 | 0.116361790544631 | OG0000124/OG0000167 | 2 |
| GO:0010817 | regulation of hormone levels | 1/22 | 76/13359 | 0.118046094515609 | 0.182872367043365 | 0.116809463713539 | OG0000035 | 1 |
| GO:0044270 | cellular nitrogen compound catabolic process | 1/22 | 76/13359 | 0.118046094515609 | 0.182872367043365 | 0.116809463713539 | OG0000167 | 1 |
| GO:1901576 | organic substance biosynthetic process | 4/22 | 1166/13359 | 0.119779955954593 | 0.185114477384371 | 0.118241608497105 | OG0000047/OG0000124/OG0000167/OG0000721 | 4 |
| GO:0032101 | regulation of response to external stimulus | 1/22 | 79/13359 | 0.122421384022709 | 0.188295747806357 | 0.120273640443363 | OG0000124 | 1 |
| GO:1901698 | response to nitrogen compound | 1/22 | 79/13359 | 0.122421384022709 | 0.188295747806357 | 0.120273640443363 | OG0000167 | 1 |
| GO:0048519 | negative regulation of biological process | 2/22 | 374/13359 | 0.125192535706041 | 0.190945906302917 | 0.121966425404503 | OG0000124/OG0000167 | 2 |
| GO:0002682 | regulation of immune system process | 1/22 | 81/13359 | 0.125326724879933 | 0.190945906302917 | 0.121966425404503 | OG0000124 | 1 |
| GO:0021700 | developmental maturation | 1/22 | 81/13359 | 0.125326724879933 | 0.190945906302917 | 0.121966425404503 | OG0000035 | 1 |
| GO:0050776 | regulation of immune response | 1/22 | 81/13359 | 0.125326724879933 | 0.190945906302917 | 0.121966425404503 | OG0000124 | 1 |
| GO:0009056 | catabolic process | 2/22 | 378/13359 | 0.127393457581452 | 0.193638055523807 | 0.123686031880939 | OG0000124/OG0000167 | 2 |
| GO:0009991 | response to extracellular stimulus | 1/22 | 84/13359 | 0.129667527255713 | 0.196171481515668 | 0.125304254121137 | OG0000127 | 1 |
| GO:0048585 | negative regulation of response to stimulus | 1/22 | 84/13359 | 0.129667527255713 | 0.196171481515668 | 0.125304254121137 | OG0000124 | 1 |
| GO:0009408 | response to heat | 1/22 | 87/13359 | 0.133987761785963 | 0.201293242124958 | 0.128575771407827 | OG0000047 | 1 |
| GO:0044425 | | 1/22 | 87/13359 | 0.133987761785963 | 0.201293242124958 | 0.128575771407827 | OG0000127 | 1 |
| GO:0099080 | supramolecular complex | 1/22 | 87/13359 | 0.133987761785963 | 0.201293242124958 | 0.128575771407827 | OG0000167 | 1 |
| GO:0046483 | heterocycle metabolic process | 4/22 | 1217/13359 | 0.134528757388977 | 0.201637070239627 | 0.12879539112585 | OG0000047/OG0000124/OG0000167/OG0000721 | 4 |
| GO:0071214 | cellular response to abiotic stimulus | 1/22 | 88/13359 | 0.135423285387584 | 0.20204028258748 | 0.129052942438149 | OG0000047 | 1 |
| GO:0104004 | cellular response to environmental stimulus | 1/22 | 88/13359 | 0.135423285387584 | 0.20204028258748 | 0.129052942438149 | OG0000047 | 1 |
| GO:0031090 | organelle membrane | 3/22 | 786/13359 | 0.136376204860292 | 0.202993152856564 | 0.129661586963945 | OG0000035/OG0000047/OG0000127 | 3 |
| GO:0005783 | endoplasmic reticulum | 2/22 | 397/13359 | 0.137977719569571 | 0.204425031496807 | 0.130576197403859 | OG0000035/OG0000721 | 2 |
| GO:0009611 | response to wounding | 1/22 | 90/13359 | 0.138287521306664 | 0.204425031496807 | 0.130576197403859 | OG0000721 | 1 |
| GO:0034220 | ion transmembrane transport | 1/22 | 90/13359 | 0.138287521306664 | 0.204425031496807 | 0.130576197403859 | OG0000127 | 1 |
| GO:0006725 | cellular aromatic compound metabolic process | 4/22 | 1231/13359 | 0.138712324478667 | 0.204584843865796 | 0.130678277326694 | OG0000047/OG0000124/OG0000167/OG0000721 | 4 |
| GO:0045229 | external encapsulating structure organization | 1/22 | 92/13359 | 0.141142698313133 | 0.207695177927754 | 0.132664998774124 | OG0000721 | 1 |
| GO:0006753 | nucleoside phosphate metabolic process | 1/22 | 94/13359 | 0.143988843697992 | 0.210922433172116 | 0.13472640346011 | OG0000721 | 1 |
| GO:0009117 | nucleotide metabolic process | 1/22 | 94/13359 | 0.143988843697992 | 0.210922433172116 | 0.13472640346011 | OG0000721 | 1 |
| GO:0009058 | biosynthetic process | 4/22 | 1256/13359 | 0.146322352493461 | 0.213855745951981 | 0.136600052815996 | OG0000047/OG0000124/OG0000167/OG0000721 | 4 |
| GO:0048584 | positive regulation of response to stimulus | 1/22 | 96/13359 | 0.146825984674126 | 0.214107417831795 | 0.136760807870399 | OG0000124 | 1 |
| GO:0090558 | plant epidermis development | 1/22 | 97/13359 | 0.148241186991639 | 0.215684249541889 | 0.137768006877009 | OG0000035 | 1 |
| GO:1901360 | organic cyclic compound metabolic process | 4/22 | 1264/13359 | 0.148794542430129 | 0.21600286384239 | 0.137971521307181 | OG0000047/OG0000124/OG0000167/OG0000721 | 4 |
| GO:0023052 | signaling | 2/22 | 422/13359 | 0.152203674729931 | 0.220456443667119 | 0.14081623907041 | OG0000035/OG0000167 | 2 |
| GO:0010608 | post-transcriptional regulation of gene expression | 1/22 | 101/13359 | 0.153879620708425 | 0.22144202928068 | 0.141445780475846 | OG0000167 | 1 |
| GO:0009505 | plant-type cell wall | 1/22 | 102/13359 | 0.155283652111684 | 0.22144202928068 | 0.141445780475846 | OG0000124 | 1 |
| GO:0010015 | root morphogenesis | 1/22 | 102/13359 | 0.155283652111684 | 0.22144202928068 | 0.141445780475846 | OG0000035 | 1 |
| GO:0016705 | oxidoreductase activity, acting on paired donors, with incorporation or reduction of molecular oxygen | 1/22 | 102/13359 | 0.155283652111684 | 0.22144202928068 | 0.141445780475846 | OG0000104 | 1 |
| GO:0023051 | regulation of signaling | 1/22 | 102/13359 | 0.155283652111684 | 0.22144202928068 | 0.141445780475846 | OG0000035 | 1 |
| GO:0031347 | regulation of defense response | 1/22 | 102/13359 | 0.155283652111684 | 0.22144202928068 | 0.141445780475846 | OG0000124 | 1 |
| GO:0031969 | chloroplast membrane | 1/22 | 102/13359 | 0.155283652111684 | 0.22144202928068 | 0.141445780475846 | OG0000127 | 1 |
| GO:0009966 | regulation of signal transduction | 1/22 | 105/13359 | 0.159482415245008 | 0.226928723013822 | 0.144950398275082 | OG0000035 | 1 |
| GO:0010646 | regulation of cell communication | 1/22 | 106/13359 | 0.160877570433223 | 0.22841079230739 | 0.145897067923247 | OG0000035 | 1 |
| GO:0048468 | cell development | 1/22 | 110/13359 | 0.166436117637809 | 0.235784499986895 | 0.150607013190261 | OG0000035 | 1 |
| GO:0071554 | cell wall organization or biogenesis | 1/22 | 112/13359 | 0.169202193703445 | 0.239178593287584 | 0.152774985446852 | OG0000124 | 1 |
| GO:0050832 | defense response to fungus | 1/22 | 113/13359 | 0.17058194396679 | 0.240423164475222 | 0.153569953518473 | OG0000167 | 1 |
| GO:0031224 | intrinsic component of membrane | 4/22 | 1333/13359 | 0.170826985285026 | 0.240423164475222 | 0.153569953518473 | OG0000035/OG0000047/OG0000124/OG0000127 | 4 |
| GO:0004553 | hydrolase activity, hydrolyzing O-glycosyl compounds | 1/22 | 114/13359 | 0.171959506795774 | 0.240805023717759 | 0.153813865565197 | OG0000039 | 1 |
| GO:0060560 | developmental growth involved in morphogenesis | 1/22 | 114/13359 | 0.171959506795774 | 0.240805023717759 | 0.153813865565197 | OG0000035 | 1 |
| GO:0044085 | cellular component biogenesis | 2/22 | 458/13359 | 0.17319933484283 | 0.240805023717759 | 0.153813865565197 | OG0000124/OG0000721 | 2 |
| GO:0006790 | sulfur compound metabolic process | 1/22 | 115/13359 | 0.173334885493434 | 0.240805023717759 | 0.153813865565197 | OG0000721 | 1 |
| GO:0015075 | ion transmembrane transporter activity | 1/22 | 115/13359 | 0.173334885493434 | 0.240805023717759 | 0.153813865565197 | OG0000127 | 1 |
| GO:0055086 | nucleobase-containing small molecule metabolic process | 1/22 | 115/13359 | 0.173334885493434 | 0.240805023717759 | 0.153813865565197 | OG0000721 | 1 |
| GO:0008270 | zinc ion binding | 2/22 | 461/13359 | 0.17497263661027 | 0.242558633584194 | 0.154933981367124 | OG0000167/OG0000006 | 2 |
| GO:0048608 | reproductive structure development | 2/22 | 465/13359 | 0.177342160026611 | 0.244792810635023 | 0.1563610587729 | OG0000035/OG0000167 | 2 |
| GO:0061458 | reproductive system development | 2/22 | 465/13359 | 0.177342160026611 | 0.244792810635023 | 0.1563610587729 | OG0000035/OG0000167 | 2 |
| GO:0016741 | transferase activity, transferring one-carbon groups | 1/22 | 120/13359 | 0.180179132286698 | 0.248178506305346 | 0.158523667055069 | OG0000721 | 1 |
| GO:0010038 | response to metal ion | 1/22 | 122/13359 | 0.182901657161316 | 0.251392490481298 | 0.16057659486503 | OG0000124 | 1 |
| GO:0051254 | positive regulation of RNA metabolic process | 1/22 | 123/13359 | 0.184259688150321 | 0.252721346121247 | 0.161425399510394 | OG0000167 | 1 |
| GO:0016043 | cellular component organization | 3/22 | 908/13359 | 0.185193726951424 | 0.253464295785211 | 0.16189995754897 | OG0000035/OG0000721/OG0001956 | 3 |
| GO:0005506 | iron ion binding | 1/22 | 128/13359 | 0.191017546896763 | 0.260331931002761 | 0.166286649752456 | OG0000104 | 1 |
| GO:0006520 | cellular amino acid metabolic process | 1/22 | 128/13359 | 0.191017546896763 | 0.260331931002761 | 0.166286649752456 | OG0000721 | 1 |
| GO:0010629 | negative regulation of gene expression | 1/22 | 129/13359 | 0.192362691932533 | 0.261613261028244 | 0.167105097479341 | OG0000167 | 1 |
| GO:0044281 | small molecule metabolic process | 2/22 | 492/13359 | 0.193477292501802 | 0.262335185270595 | 0.167566225559839 | OG0000124/OG0000721 | 2 |
| GO:2000113 | negative regulation of cellular macromolecule biosynthetic process | 1/22 | 130/13359 | 0.193705701817452 | 0.262335185270595 | 0.167566225559839 | OG0000167 | 1 |
| GO:0010558 | negative regulation of macromolecule biosynthetic process | 1/22 | 131/13359 | 0.195046579779506 | 0.263598515768956 | 0.168373176114439 | OG0000167 | 1 |
| GO:0045935 | positive regulation of nucleobase-containing compound metabolic process | 1/22 | 132/13359 | 0.196385329042044 | 0.26485370054522 | 0.169174923600662 | OG0000167 | 1 |
| GO:0032268 | regulation of cellular protein metabolic process | 1/22 | 133/13359 | 0.197721952823784 | 0.266100794842009 | 0.169971503304656 | OG0000167 | 1 |
| GO:0009555 | pollen development | 1/22 | 134/13359 | 0.199056454338821 | 0.267339853436337 | 0.170762950215161 | OG0000721 | 1 |
| GO:0140657 | ATP-dependent activity | 1/22 | 135/13359 | 0.200388836796632 | 0.268570930644448 | 0.171549299026599 | OG0000035 | 1 |
| GO:0031327 | negative regulation of cellular biosynthetic process | 1/22 | 137/13359 | 0.203047257355432 | 0.271570451866686 | 0.173465238930651 | OG0000167 | 1 |
| GO:0009620 | response to fungus | 1/22 | 138/13359 | 0.204373301852345 | 0.272216810302299 | 0.173878099459836 | OG0000167 | 1 |
| GO:0009733 | response to auxin | 1/22 | 138/13359 | 0.204373301852345 | 0.272216810302299 | 0.173878099459836 | OG0000035 | 1 |
| GO:0009890 | negative regulation of biosynthetic process | 1/22 | 139/13359 | 0.205697240083889 | 0.273416496078585 | 0.174644397038953 | OG0000167 | 1 |
| GO:0051246 | regulation of protein metabolic process | 1/22 | 142/13359 | 0.20965644902825 | 0.278106911852669 | 0.177640393427157 | OG0000167 | 1 |
| GO:0031326 | regulation of cellular biosynthetic process | 2/22 | 520/13359 | 0.210431680242008 | 0.278563248844953 | 0.177931878030343 | OG0000047/OG0000167 | 2 |
| GO:0048366 | leaf development | 1/22 | 144/13359 | 0.212285448627827 | 0.280442535406086 | 0.179132269641822 | OG0000167 | 1 |
| GO:0020037 | heme binding | 1/22 | 147/13359 | 0.21621330180073 | 0.284468010108496 | 0.181703535868552 | OG0000104 | 1 |
| GO:0051172 | negative regulation of nitrogen compound metabolic process | 1/22 | 147/13359 | 0.21621330180073 | 0.284468010108496 | 0.181703535868552 | OG0000167 | 1 |
| GO:0009889 | regulation of biosynthetic process | 2/22 | 532/13359 | 0.217754281055176 | 0.285913141385455 | 0.182626611411273 | OG0000047/OG0000167 | 2 |
| GO:0046394 | carboxylic acid biosynthetic process | 1/22 | 150/13359 | 0.220122455184053 | 0.288436320586001 | 0.184238288528128 | OG0000721 | 1 |
| GO:0015979 | photosynthesis | 1/22 | 153/13359 | 0.224012993583959 | 0.292348270414621 | 0.186737040903587 | OG0000047 | 1 |
| GO:0016049 | cell growth | 1/22 | 153/13359 | 0.224012993583959 | 0.292348270414621 | 0.186737040903587 | OG0000035 | 1 |
| GO:0003006 | developmental process involved in reproduction | 2/22 | 544/13359 | 0.22510515921877 | 0.29285209981492 | 0.187058861214679 | OG0000035/OG0000167 | 2 |
| GO:0009414 | response to water deprivation | 1/22 | 154/13359 | 0.225305717659467 | 0.29285209981492 | 0.187058861214679 | OG0000124 | 1 |
| GO:0009415 | response to water | 1/22 | 158/13359 | 0.230456086800955 | 0.298345956058952 | 0.190568054057535 | OG0000124 | 1 |
| GO:0080134 | regulation of response to stress | 1/22 | 158/13359 | 0.230456086800955 | 0.298345956058952 | 0.190568054057535 | OG0000124 | 1 |
| GO:0005618 | cell wall | 1/22 | 161/13359 | 0.234297397078395 | 0.30210802098332 | 0.192971067664105 | OG0000124 | 1 |
| GO:0016053 | organic acid biosynthetic process | 1/22 | 161/13359 | 0.234297397078395 | 0.30210802098332 | 0.192971067664105 | OG0000721 | 1 |
| GO:0001101 | response to acid chemical | 1/22 | 162/13359 | 0.235573761471809 | 0.3025460236795 | 0.193250841261796 | OG0000124 | 1 |
| GO:1905392 | plant organ morphogenesis | 1/22 | 162/13359 | 0.235573761471809 | 0.3025460236795 | 0.193250841261796 | OG0000035 | 1 |
| GO:0030312 | external encapsulating structure | 1/22 | 163/13359 | 0.236848094823906 | 0.302977958923254 | 0.193526739282899 | OG0000124 | 1 |
| GO:0055114 | | 1/22 | 163/13359 | 0.236848094823906 | 0.302977958923254 | 0.193526739282899 | OG0000124 | 1 |
| GO:0071396 | cellular response to lipid | 1/22 | 164/13359 | 0.238120400212953 | 0.304003514896379 | 0.194181811698518 | OG0000167 | 1 |
| GO:0051171 | regulation of nitrogen compound metabolic process | 2/22 | 566/13359 | 0.238643021492536 | 0.304069806477669 | 0.19422415535155 | OG0000047/OG0000167 | 2 |
| GO:0005774 | vacuolar membrane | 1/22 | 165/13359 | 0.239390680712788 | 0.304422007363112 | 0.194449123164966 | OG0000035 | 1 |
| GO:0048589 | developmental growth | 1/22 | 169/13359 | 0.244451615141905 | 0.310102802445666 | 0.198077722924395 | OG0000035 | 1 |
| GO:0006810 | transport | 2/22 | 576/13359 | 0.244818001930789 | 0.310102802445666 | 0.198077722924395 | OG0000035/OG0000127 | 2 |
| GO:0031324 | negative regulation of cellular metabolic process | 1/22 | 171/13359 | 0.246970012616133 | 0.31221649344427 | 0.199427840036099 | OG0000167 | 1 |
| GO:0051234 | establishment of localization | 2/22 | 584/13359 | 0.249765727258507 | 0.315134101189445 | 0.201291457823468 | OG0000035/OG0000127 | 2 |
| GO:0006355 | regulation of transcription, DNA-templated | 2/22 | 590/13359 | 0.253480395950645 | 0.319197535641553 | 0.203886970786196 | OG0000018/OG0000167 | 2 |
| GO:0009908 | flower development | 1/22 | 177/13359 | 0.254477218204287 | 0.319829344280097 | 0.204290537653247 | OG0000035 | 1 |
| GO:0090567 | reproductive shoot system development | 1/22 | 183/13359 | 0.261912951456966 | 0.328535469206213 | 0.209851562536802 | OG0000035 | 1 |
| GO:0051173 | positive regulation of nitrogen compound metabolic process | 1/22 | 186/13359 | 0.265604218783863 | 0.331876838170939 | 0.21198585719897 | OG0000167 | 1 |
| GO:1901135 | carbohydrate derivative metabolic process | 1/22 | 186/13359 | 0.265604218783863 | 0.331876838170939 | 0.21198585719897 | OG0000721 | 1 |
| GO:0008610 | lipid biosynthetic process | 1/22 | 188/13359 | 0.268055266827736 | 0.334292861719532 | 0.213529088795921 | OG0000721 | 1 |
| GO:0009535 | chloroplast thylakoid membrane | 1/22 | 189/13359 | 0.269277860718085 | 0.335170516423667 | 0.214089689486846 | OG0000047 | 1 |
| GO:0055035 | plastid thylakoid membrane | 1/22 | 190/13359 | 0.270498505142095 | 0.336042373695757 | 0.214646587076318 | OG0000047 | 1 |
| GO:0019637 | organophosphate metabolic process | 1/22 | 191/13359 | 0.271717203060454 | 0.336908470589354 | 0.215199805232242 | OG0000721 | 1 |
| GO:0016071 | mRNA metabolic process | 1/22 | 192/13359 | 0.272933957429582 | 0.337213239012562 | 0.215394475628034 | OG0000167 | 1 |
| GO:0022414 | reproductive process | 2/22 | 623/13359 | 0.273951978158775 | 0.337213239012562 | 0.215394475628034 | OG0000035/OG0000167 | 2 |
| GO:0006811 | ion transport | 1/22 | 193/13359 | 0.274148771201629 | 0.337213239012562 | 0.215394475628034 | OG0000127 | 1 |
| GO:0044265 | cellular macromolecule catabolic process | 1/22 | 193/13359 | 0.274148771201629 | 0.337213239012562 | 0.215394475628034 | OG0000167 | 1 |
| GO:0000003 | reproduction | 2/22 | 624/13359 | 0.27457300885543 | 0.337213239012562 | 0.215394475628034 | OG0000035/OG0000167 | 2 |
| GO:0051179 | localization | 2/22 | 633/13359 | 0.280162983838115 | 0.341519277075429 | 0.218144951301235 | OG0000035/OG0000127 | 2 |
| GO:0034357 | photosynthetic membrane | 1/22 | 198/13359 | 0.280193834133092 | 0.341519277075429 | 0.218144951301235 | OG0000047 | 1 |
| GO:0043043 | peptide biosynthetic process | 1/22 | 198/13359 | 0.280193834133092 | 0.341519277075429 | 0.218144951301235 | OG0000167 | 1 |
| GO:0098796 | membrane protein complex | 1/22 | 198/13359 | 0.280193834133092 | 0.341519277075429 | 0.218144951301235 | OG0000047 | 1 |
| GO:0042651 | thylakoid membrane | 1/22 | 200/13359 | 0.282598377975293 | 0.343801416519848 | 0.219602664617534 | OG0000047 | 1 |
| GO:0010604 | positive regulation of macromolecule metabolic process | 1/22 | 203/13359 | 0.286190822517516 | 0.347517427342698 | 0.221976261232422 | OG0000167 | 1 |
| GO:0009737 | response to abscisic acid | 1/22 | 204/13359 | 0.287384483349426 | 0.348312150551087 | 0.222483889548682 | OG0001956 | 1 |
| GO:0031325 | positive regulation of cellular metabolic process | 1/22 | 206/13359 | 0.289766091404773 | 0.35054100196158 | 0.223907565209287 | OG0000167 | 1 |
| GO:0005737 | cytoplasm | 8/22 | 3880/13359 | 0.292983957521382 | 0.353626804834703 | 0.225878617394824 | OG0000035/OG0000047/OG0000124/OG0000127/OG0000167/OG0000572/OG0000721/OG0001956 | 8 |
| GO:0009057 | macromolecule catabolic process | 1/22 | 210/13359 | 0.294506534057384 | 0.353626804834703 | 0.225878617394824 | OG0000167 | 1 |
| GO:0009793 | embryo development ending in seed dormancy | 1/22 | 210/13359 | 0.294506534057384 | 0.353626804834703 | 0.225878617394824 | OG0000167 | 1 |
| GO:1901701 | cellular response to oxygen-containing compound | 1/22 | 210/13359 | 0.294506534057384 | 0.353626804834703 | 0.225878617394824 | OG0000167 | 1 |
| GO:0097305 | response to alcohol | 1/22 | 211/13359 | 0.295686917071358 | 0.353731950761711 | 0.225945779205786 | OG0001956 | 1 |
| GO:0031323 | regulation of cellular metabolic process | 2/22 | 658/13359 | 0.295689246766756 | 0.353731950761711 | 0.225945779205786 | OG0000047/OG0000167 | 2 |
| GO:0140110 | transcription regulator activity | 1/22 | 212/13359 | 0.296865414776289 | 0.354482547034164 | 0.226425221504631 | OG0000167 | 1 |
| GO:0043604 | amide biosynthetic process | 1/22 | 213/13359 | 0.298042030040222 | 0.355230906653106 | 0.226903235144236 | OG0000167 | 1 |
| GO:0006518 | peptide metabolic process | 1/22 | 217/13359 | 0.302729723901896 | 0.36015359418163 | 0.230047594784421 | OG0000167 | 1 |
| GO:0009790 | embryo development | 1/22 | 218/13359 | 0.303896969836621 | 0.360877651680988 | 0.23051008547979 | OG0000167 | 1 |
| GO:0044283 | small molecule biosynthetic process | 1/22 | 221/13359 | 0.307387527075786 | 0.364352922001758 | 0.232729909442218 | OG0000721 | 1 |
| GO:0010605 | negative regulation of macromolecule metabolic process | 1/22 | 225/13359 | 0.312015621136551 | 0.369161339293428 | 0.235801279131536 | OG0000167 | 1 |
| GO:0009987 | cellular process | 7/22 | 3392/13359 | 0.315880086144632 | 0.373050339395671 | 0.238285372401993 | OG0000035/OG0000047/OG0000124/OG0000127/OG0000167/OG0000721/OG0001956 | 7 |
| GO:0009534 | chloroplast thylakoid | 1/22 | 230/13359 | 0.317759235193664 | 0.373902488042089 | 0.238829681134918 | OG0000047 | 1 |
| GO:0009893 | positive regulation of metabolic process | 1/22 | 230/13359 | 0.317759235193664 | 0.373902488042089 | 0.238829681134918 | OG0000167 | 1 |
| GO:0031976 | plastid thylakoid | 1/22 | 231/13359 | 0.318902452257091 | 0.374565425741966 | 0.239253131645512 | OG0000047 | 1 |
| GO:0040007 | growth | 1/22 | 234/13359 | 0.322321142212346 | 0.377893752938612 | 0.241379095896914 | OG0000035 | 1 |
| GO:0009892 | negative regulation of metabolic process | 1/22 | 241/13359 | 0.330234501175909 | 0.386470086521082 | 0.246857216744768 | OG0000167 | 1 |
| GO:0009888 | tissue development | 1/22 | 242/13359 | 0.331357754794916 | 0.387083380827334 | 0.247248957608465 | OG0000035 | 1 |
| GO:0048523 | negative regulation of cellular process | 1/22 | 251/13359 | 0.341386442005666 | 0.398078775335126 | 0.254272250173325 | OG0000167 | 1 |
| GO:1901564 | organonitrogen compound metabolic process | 3/22 | 1256/13359 | 0.342553041345827 | 0.3987193958728 | 0.254681445628382 | OG0000047/OG0000167/OG0000721 | 3 |
| GO:0000325 | plant-type vacuole | 1/22 | 254/13359 | 0.344697312096388 | 0.400493639594005 | 0.255814741275623 | OG0000035 | 1 |
| GO:0008168 | methyltransferase activity | 1/22 | 259/13359 | 0.350180151612296 | 0.406133533108695 | 0.259417215216895 | OG0000721 | 1 |
| GO:0019222 | regulation of metabolic process | 2/22 | 751/13359 | 0.353085705155078 | 0.408769472276308 | 0.261100917601944 | OG0000047/OG0000167 | 2 |
| GO:0012505 | endomembrane system | 2/22 | 757/13359 | 0.356752821489983 | 0.412276069199516 | 0.263340751387013 | OG0000035/OG0000721 | 2 |
| GO:0009579 | thylakoid | 1/22 | 268/13359 | 0.359939096429957 | 0.415215457667415 | 0.265218281579969 | OG0000047 | 1 |
| GO:0016021 | integral component of membrane | 3/22 | 1300/13359 | 0.363177301966866 | 0.41820416590124 | 0.26712731470309 | OG0000035/OG0000047/OG0000127 | 3 |
| GO:0044271 | cellular nitrogen compound biosynthetic process | 2/22 | 772/13359 | 0.365894473311908 | 0.420583326974186 | 0.268647000446278 | OG0000047/OG0000167 | 2 |
| GO:0048316 | seed development | 1/22 | 275/13359 | 0.367432503155965 | 0.421601060459597 | 0.269297076259023 | OG0000167 | 1 |
| GO:0043167 | ion binding | 3/22 | 1328/13359 | 0.376287695465407 | 0.430996190196193 | 0.275298201982903 | OG0000035/OG0000124/OG0000167 | 3 |
| GO:0010154 | fruit development | 1/22 | 285/13359 | 0.377992250435278 | 0.432182289878212 | 0.276055821463678 | OG0000167 | 1 |
| GO:0003723 | RNA binding | 2/22 | 800/13359 | 0.382848621980565 | 0.436961501412447 | 0.279108536017076 | OG0000088/OG0000167 | 2 |
| GO:0034641 | cellular nitrogen compound metabolic process | 3/22 | 1351/13359 | 0.387037430620267 | 0.440963280741962 | 0.281664666858154 | OG0000047/OG0000167/OG0000721 | 3 |
| GO:0044763 | | 1/22 | 297/13359 | 0.390441892811028 | 0.443278493419902 | 0.283143505655209 | OG0000127 | 1 |
| GO:0048522 | positive regulation of cellular process | 1/22 | 297/13359 | 0.390441892811028 | 0.443278493419902 | 0.283143505655209 | OG0000167 | 1 |
| GO:0010035 | response to inorganic substance | 1/22 | 302/13359 | 0.395558728357325 | 0.448299892138302 | 0.286350916927187 | OG0000124 | 1 |
| GO:0022607 | cellular component assembly | 1/22 | 307/13359 | 0.400634548189441 | 0.45325729970294 | 0.289517453940936 | OG0000721 | 1 |
| GO:0043169 | cation binding | 2/22 | 848/13359 | 0.411525160653044 | 0.464764429688577 | 0.296867616812648 | OG0000124/OG0000167 | 2 |
| GO:0016301 | kinase activity | 1/22 | 327/13359 | 0.420533901146069 | 0.474109773368866 | 0.302836941764047 | OG0000035 | 1 |
| GO:0005773 | vacuole | 1/22 | 330/13359 | 0.423463855184462 | 0.476581272559516 | 0.304415608348265 | OG0000035 | 1 |
| GO:0006412 | translation | 1/22 | 344/13359 | 0.436950999387919 | 0.490904948877558 | 0.313564836174031 | OG0000167 | 1 |
| GO:0098588 | bounding membrane of organelle | 1/22 | 347/13359 | 0.43980165629557 | 0.493249774248156 | 0.3150625900363 | OG0000035 | 1 |
| GO:0006629 | lipid metabolic process | 1/22 | 369/13359 | 0.460289495952659 | 0.515332780563982 | 0.329168078835068 | OG0000721 | 1 |
| GO:1903506 | regulation of nucleic acid-templated transcription | 1/22 | 394/13359 | 0.48270282971993 | 0.53855963384987 | 0.344004198255091 | OG0000167 | 1 |
| GO:2001141 | regulation of RNA biosynthetic process | 1/22 | 394/13359 | 0.48270282971993 | 0.53855963384987 | 0.344004198255091 | OG0000167 | 1 |
| GO:0005794 | Golgi apparatus | 1/22 | 396/13359 | 0.48445698633358 | 0.539584850295677 | 0.344659053798119 | OG0000035 | 1 |
| GO:0110165 | cellular anatomical entity | 9/22 | 5182/13359 | 0.499250914858403 | 0.555105148018121 | 0.35457262183983 | OG0000035/OG0000047/OG0000124/OG0000127/OG0000167/OG0000572/OG0000721/OG0000785/OG0001956 | 9 |
| GO:0051252 | regulation of RNA metabolic process | 1/22 | 422/13359 | 0.506750570396927 | 0.562475719031641 | 0.359280563565917 | OG0000167 | 1 |
| GO:0044699 | | 1/22 | 427/13359 | 0.510930948299364 | 0.566143040482657 | 0.361623059913967 | OG0000127 | 1 |
| GO:0043231 | intracellular membrane-bounded organelle | 8/22 | 4635/13359 | 0.514420887209359 | 0.568389950064092 | 0.363058270205514 | OG0000035/OG0000047/OG0000124/OG0000127/OG0000167/OG0000572/OG0000721/OG0001956 | 8 |
| GO:0043227 | membrane-bounded organelle | 8/22 | 4636/13359 | 0.514718453231414 | 0.568389950064092 | 0.363058270205514 | OG0000035/OG0000047/OG0000124/OG0000127/OG0000167/OG0000572/OG0000721/OG0001956 | 8 |
| GO:0005488 | binding | 6/22 | 3449/13359 | 0.518867521539471 | 0.571374860001362 | 0.364964877172126 | OG0000035/OG0000124/OG0000127/OG0000167/OG0000721/OG0001956 | 6 |
| GO:0097659 | nucleic acid-templated transcription | 1/22 | 437/13359 | 0.519190468762848 | 0.571374860001362 | 0.364964877172126 | OG0000167 | 1 |
| GO:0043229 | intracellular organelle | 8/22 | 4657/13359 | 0.520960568491084 | 0.572027356178703 | 0.365381658175088 | OG0000035/OG0000047/OG0000124/OG0000127/OG0000167/OG0000572/OG0000721/OG0001956 | 8 |
| GO:0043226 | organelle | 8/22 | 4659/13359 | 0.521554354162935 | 0.572027356178703 | 0.365381658175088 | OG0000035/OG0000047/OG0000124/OG0000127/OG0000167/OG0000572/OG0000721/OG0001956 | 8 |
| GO:0019219 | regulation of nucleobase-containing compound metabolic process | 1/22 | 443/13359 | 0.524082088808662 | 0.573825473509146 | 0.366530203056192 | OG0000167 | 1 |
| GO:0032774 | RNA biosynthetic process | 1/22 | 451/13359 | 0.530530395901197 | 0.579902936974912 | 0.370412174179836 | OG0000167 | 1 |
| GO:0005829 | cytosol | 2/22 | 1064/13359 | 0.532465903125574 | 0.58103542807284 | 0.371135551253957 | OG0000035/OG0000721 | 2 |
| GO:0005575 | cellular_component | 9/22 | 5329/13359 | 0.541816779266178 | 0.590242224967876 | 0.377016379643812 | OG0000035/OG0000047/OG0000124/OG0000127/OG0000167/OG0000572/OG0000721/OG0000785/OG0001956 | 9 |
| GO:2000112 | regulation of cellular macromolecule biosynthetic process | 1/22 | 481/13359 | 0.553976684258121 | 0.602472959647721 | 0.384828743982249 | OG0000167 | 1 |
| GO:0010556 | regulation of macromolecule biosynthetic process | 1/22 | 483/13359 | 0.555499360411481 | 0.603113591303893 | 0.385237946539231 | OG0000167 | 1 |
| GO:0006351 | transcription, DNA-templated | 1/22 | 486/13359 | 0.557774074319698 | 0.604567201359941 | 0.386166437890006 | OG0000167 | 1 |
| GO:0032559 | adenyl ribonucleotide binding | 1/22 | 496/13359 | 0.565276474824312 | 0.611232801914129 | 0.390424080740327 | OG0000035 | 1 |
| GO:0031984 | organelle subcompartment | 1/22 | 497/13359 | 0.566019996363133 | 0.611232801914129 | 0.390424080740327 | OG0000047 | 1 |
| GO:0030554 | adenyl nucleotide binding | 1/22 | 498/13359 | 0.566762303942049 | 0.611232801914129 | 0.390424080740327 | OG0000035 | 1 |
| GO:0006139 | nucleobase-containing compound metabolic process | 2/22 | 1137/13359 | 0.569744489483042 | 0.613424900343408 | 0.391824280486583 | OG0000167/OG0000721 | 2 |
| GO:0034654 | nucleobase-containing compound biosynthetic process | 1/22 | 528/13359 | 0.588476304002747 | 0.63253858966019 | 0.404033122285799 | OG0000167 | 1 |
| GO:0006796 | phosphate-containing compound metabolic process | 1/22 | 531/13359 | 0.590589633511865 | 0.633755653236984 | 0.404810519910213 | OG0000721 | 1 |
| GO:0035639 | purine ribonucleoside triphosphate binding | 1/22 | 536/13359 | 0.594088846918377 | 0.636144494964525 | 0.406336389157722 | OG0000035 | 1 |
| GO:0006793 | phosphorus metabolic process | 1/22 | 537/13359 | 0.594785255353828 | 0.636144494964525 | 0.406336389157722 | OG0000721 | 1 |
| GO:0005622 | intracellular anatomical structure | 8/22 | 4921/13359 | 0.597817533965114 | 0.637515885672735 | 0.407212363017292 | OG0000035/OG0000047/OG0000124/OG0000127/OG0000167/OG0000572/OG0000721/OG0001956 | 8 |
| GO:0010468 | regulation of gene expression | 1/22 | 544/13359 | 0.59962827748344 | 0.637515885672735 | 0.407212363017292 | OG0000167 | 1 |
| GO:0032555 | purine ribonucleotide binding | 1/22 | 546/13359 | 0.601001817917485 | 0.637515885672735 | 0.407212363017292 | OG0000035 | 1 |
| GO:0043228 | non-membrane-bounded organelle | 1/22 | 546/13359 | 0.601001817917485 | 0.637515885672735 | 0.407212363017292 | OG0000167 | 1 |
| GO:0043232 | intracellular non-membrane-bounded organelle | 1/22 | 546/13359 | 0.601001817917485 | 0.637515885672735 | 0.407212363017292 | OG0000167 | 1 |
| GO:0017076 | purine nucleotide binding | 1/22 | 549/13359 | 0.603053697955189 | 0.638643752260741 | 0.407932786192293 | OG0000035 | 1 |
| GO:0032553 | ribonucleotide binding | 1/22 | 552/13359 | 0.605095503314301 | 0.639757275189916 | 0.408644047375667 | OG0000035 | 1 |
| GO:0044237 | cellular metabolic process | 4/22 | 2489/13359 | 0.607555314389759 | 0.641308387411413 | 0.409634818095607 | OG0000047/OG0000124/OG0000167/OG0000721 | 4 |
| GO:0097367 | carbohydrate derivative binding | 1/22 | 563/13359 | 0.612496687751892 | 0.645469592639025 | 0.412292782001789 | OG0000035 | 1 |
| GO:0071704 | organic substance metabolic process | 4/22 | 2523/13359 | 0.619245048710218 | 0.651518406297721 | 0.416156453102015 | OG0000047/OG0000124/OG0000167/OG0000721 | 4 |
| GO:0080090 | regulation of primary metabolic process | 1/22 | 606/13359 | 0.640177947547401 | 0.672447079862799 | 0.429524613502065 | OG0000167 | 1 |
| GO:0008150 | biological_process | 8/22 | 5132/13359 | 0.656009555093014 | 0.687958072386505 | 0.439432237861349 | OG0000035/OG0000047/OG0000124/OG0000127/OG0000167/OG0000572/OG0000721/OG0001956 | 8 |
| GO:0006996 | organelle organization | 1/22 | 640/13359 | 0.660718568226393 | 0.690762171242367 | 0.441223352007508 | OG0001956 | 1 |
| GO:0003677 | DNA binding | 1/22 | 642/13359 | 0.661891306500039 | 0.690762171242367 | 0.441223352007508 | OG0000167 | 1 |
| GO:1901265 | nucleoside phosphate binding | 1/22 | 642/13359 | 0.661891306500039 | 0.690762171242367 | 0.441223352007508 | OG0000035 | 1 |
| GO:0060255 | regulation of macromolecule metabolic process | 1/22 | 652/13359 | 0.667697191145892 | 0.695697395935881 | 0.444375719743955 | OG0000167 | 1 |
| GO:0043168 | anion binding | 1/22 | 662/13359 | 0.673407836381628 | 0.700517652661082 | 0.447454651854561 | OG0000035 | 1 |
| GO:0000166 | nucleotide binding | 1/22 | 684/13359 | 0.68564331861555 | 0.712099009365989 | 0.454852226937376 | OG0000035 | 1 |
| GO:0005634 | nucleus | 4/22 | 2749/13359 | 0.69199262619974 | 0.717539705497644 | 0.458327463834245 | OG0000035/OG0000167/OG0000721/OG0001956 | 4 |
| GO:0006807 | nitrogen compound metabolic process | 3/22 | 2098/13359 | 0.694375747447585 | 0.71869047764165 | 0.459062517900484 | OG0000047/OG0000167/OG0000721 | 3 |
| GO:0036094 | small molecule binding | 1/22 | 702/13359 | 0.695327474498501 | 0.71869047764165 | 0.459062517900484 | OG0000035 | 1 |
| GO:0005524 | ATP binding | 2/22 | 1446/13359 | 0.705020426136826 | 0.727545040390398 | 0.464718357231605 | OG0000035/OG0000538 | 2 |
| GO:0005515 | protein binding | 5/22 | 3451/13359 | 0.707995154256186 | 0.72944955287001 | 0.465934861862545 | OG0000035/OG0000127/OG0000167/OG0000721/OG0001956 | 5 |
| GO:0034645 | cellular macromolecule biosynthetic process | 1/22 | 753/13359 | 0.721243702014726 | 0.741912209726779 | 0.473895366160823 | OG0000167 | 1 |
| GO:0008152 | metabolic process | 4/22 | 2853/13359 | 0.722388204207653 | 0.741912209726779 | 0.473895366160823 | OG0000047/OG0000124/OG0000167/OG0000721 | 4 |
| GO:1901363 | heterocyclic compound binding | 2/22 | 1509/13359 | 0.728140570818368 | 0.746633029759787 | 0.476910783226066 | OG0000035/OG0000167 | 2 |
| GO:0097159 | organic cyclic compound binding | 2/22 | 1516/13359 | 0.730619385560258 | 0.747971029330939 | 0.477765428544449 | OG0000035/OG0000167 | 2 |
| GO:0009059 | macromolecule biosynthetic process | 1/22 | 775/13359 | 0.731761130862467 | 0.747971029330939 | 0.477765428544449 | OG0000167 | 1 |
| GO:0003824 | catalytic activity | 3/22 | 2240/13359 | 0.739075720653349 | 0.754254210966925 | 0.481778801856012 | OG0000124/OG0000721/OG0001850 | 3 |
| GO:0003674 | molecular_function | 8/22 | 5498/13359 | 0.747009087602945 | 0.761148060869878 | 0.486182238652423 | OG0000035/OG0000124/OG0000127/OG0000167/OG0000572/OG0000721/OG0001850/OG0001956 | 8 |
| GO:0016070 | RNA metabolic process | 1/22 | 850/13359 | 0.764842258765455 | 0.778091494744069 | 0.497004832881986 | OG0000167 | 1 |
| GO:0044238 | primary metabolic process | 3/22 | 2347/13359 | 0.769521704299545 | 0.78162110216589 | 0.499259364590238 | OG0000124/OG0000167/OG0000721 | 3 |
| GO:0016740 | transferase activity | 1/22 | 876/13359 | 0.775372526522492 | 0.786327554369748 | 0.502265604225096 | OG0000721 | 1 |
| GO:0044267 | cellular protein metabolic process | 1/22 | 893/13359 | 0.782012673513149 | 0.791818475061903 | 0.505772921988375 | OG0000167 | 1 |
| GO:0032991 | protein-containing complex | 1/22 | 925/13359 | 0.79400683981739 | 0.802704880316172 | 0.512726597822942 | OG0000047 | 1 |
| GO:0010467 | gene expression | 1/22 | 978/13359 | 0.812498323999463 | 0.820115495786958 | 0.523847603631233 | OG0000167 | 1 |
| GO:0019538 | protein metabolic process | 1/22 | 1002/13359 | 0.820340367935355 | 0.826739278761684 | 0.528078535562295 | OG0000167 | 1 |
| GO:0090304 | nucleic acid metabolic process | 1/22 | 1010/13359 | 0.822884076767942 | 0.828011080361512 | 0.528890897020878 | OG0000167 | 1 |
| GO:0003676 | nucleic acid binding | 1/22 | 1137/13359 | 0.858940011183195 | 0.862947507347347 | 0.551206489946488 | OG0000167 | 1 |
| GO:0005739 | mitochondrion | 1/22 | 1326/13359 | 0.899914586667511 | 0.902709352464615 | 0.576604311823577 | OG0000035 | 1 |
| GO:0044260 | cellular macromolecule metabolic process | 1/22 | 1534/13359 | 0.931825324516141 | 0.933270014941748 | 0.596124891408123 | OG0000167 | 1 |
| GO:0043170 | macromolecule metabolic process | 1/22 | 1923/13359 | 0.967360756801155 | 0.967360756801155 | 0.617900304164987 | OG0000167 | 1 |

**Table S4b. Related to Figure 3. Enriched Pfam domains in orthogroups expanded in Draparnaldia with respect to Chlamydomonas.**

| ID | Description | GeneRatio | BgRatio | pvalue | p.adjust | qvalue | geneID | Count |
|---|---|---|---|---|---|---|---|---|
| **PF13191** | AAA_16 | 2/18 | 59/13933 | 0.00258210845119555 | 0.0250999478326064 | 0.00812953775954863 | OG0000035/OG0000691 | 2 |
| **PF00305** | Lipoxygenase | 1/18 | 2/13933 | 0.00258221748192833 | 0.0250999478326064 | 0.00812953775954863 | OG0000108 | 1 |
| **PF13668** | Ferritin_2 | 1/18 | 2/13933 | 0.00258221748192833 | 0.0250999478326064 | 0.00812953775954863 | OG0000572 | 1 |
| **PF00724** | Oxidored_FMN | 1/18 | 3/13933 | 0.00387096345029614 | 0.0250999478326064 | 0.00812953775954863 | OG0000253 | 1 |
| **PF09818** | ABC_ATPase | 1/18 | 3/13933 | 0.00387096345029614 | 0.0250999478326064 | 0.00812953775954863 | OG0000035 | 1 |
| **PF01740** | STAS | 1/18 | 4/13933 | 0.00515813664899611 | 0.0250999478326064 | 0.00812953775954863 | OG0000127 | 1 |
| **PF06414** | Zeta_toxin | 1/18 | 4/13933 | 0.00515813664899611 | 0.0250999478326064 | 0.00812953775954863 | OG0000035 | 1 |
| **PF00394** | Cu-oxidase | 1/18 | 5/13933 | 0.00644373888464267 | 0.0250999478326064 | 0.00812953775954863 | OG0000124 | 1 |
| **PF00759** | Glyco_hydro_9 | 1/18 | 5/13933 | 0.00644373888464267 | 0.0250999478326064 | 0.00812953775954863 | OG0000039 | 1 |
| **PF00916** | Sulfate_transp | 1/18 | 5/13933 | 0.00644373888464267 | 0.0250999478326064 | 0.00812953775954863 | OG0000127 | 1 |
| **PF01582** | TIR | 1/18 | 6/13933 | 0.00772777196190244 | 0.0250999478326064 | 0.00812953775954863 | OG0000691 | 1 |
| **PF13768** | VWA_3 | 1/18 | 6/13933 | 0.00772777196190244 | 0.0250999478326064 | 0.00812953775954863 | OG0000785 | 1 |
| **PF08417** | PaO | 1/18 | 7/13933 | 0.00901023768349973 | 0.0250999478326064 | 0.00812953775954863 | OG0000151 | 1 |
| **PF19112** | VanA_C | 1/18 | 7/13933 | 0.00901023768349973 | 0.0250999478326064 | 0.00812953775954863 | OG0000151 | 1 |
| **PF00092** | VWA | 1/18 | 8/13933 | 0.0102911378502162 | 0.0267569584105621 | 0.00866622134755047 | OG0000785 | 1 |
| **PF07732** | Cu-oxidase_3 | 1/18 | 9/13933 | 0.0115704742608947 | 0.0282030310109309 | 0.00913458494281164 | OG0000124 | 1 |
| **PF07731** | Cu-oxidase_2 | 1/18 | 10/13933 | 0.0128482487124391 | 0.0294753941050073 | 0.00954668634979992 | OG0000124 | 1 |
| **PF13676** | TIR_2 | 1/18 | 11/13933 | 0.014124462999818 | 0.0306030031662722 | 0.00991190385952137 | OG0000691 | 1 |
| **PF13519** | VWA_2 | 1/18 | 15/13933 | 0.0192137543394003 | 0.0394387589071901 | 0.0127736870954462 | OG0000785 | 1 |
| **PF00931** | NB-ARC | 1/18 | 16/13933 | 0.0204821946628585 | 0.0399402795925741 | 0.0129361229449633 | OG0000691 | 1 |
| **PF00664** | ABC_membrane | 1/18 | 17/13933 | 0.0217490855514171 | 0.0403911588812031 | 0.0130821567226569 | OG0000035 | 1 |
| **PF00355** | Rieske | 1/18 | 19/13933 | 0.0242782261478012 | 0.0411674269462716 | 0.0133335795777398 | OG0000151 | 1 |
| **PF00847** | AP2 | 1/18 | 19/13933 | 0.0242782261478012 | 0.0411674269462716 | 0.0133335795777398 | OG0000018 | 1 |
| **PF07993** | NAD_binding_4 | 1/18 | 24/13933 | 0.0305740869960234 | 0.049682891368538 | 0.0160916247347491 | OG0000721 | 1 |
| **PF13555** | AAA_29 | 1/18 | 26/13933 | 0.0330816722879347 | 0.0516074087691781 | 0.0167149502086407 | OG0000035 | 1 |
| **PF00504** | Chloroa_b-bind | 1/18 | 28/13933 | 0.0355831307524644 | 0.0533746961286967 | 0.0172873509728572 | OG0000047 | 1 |
| **PF00083** | Sugar_tr | 1/18 | 35/13933 | 0.0442901897808691 | 0.0639747185723664 | 0.0207205566226288 | OG0000189 | 1 |
| **PF13401** | AAA_22 | 1/18 | 48/13933 | 0.0602638249700786 | 0.0788791502030666 | 0.0255479028997787 | OG0000035 | 1 |
| **PF02463** | SMC_N | 1/18 | 49/13933 | 0.0614820641598907 | 0.0788791502030666 | 0.0255479028997787 | OG0000035 | 1 |
| **PF17123** | zf-RING_11 | 1/18 | 49/13933 | 0.0614820641598907 | 0.0788791502030666 | 0.0255479028997787 | OG0000006 | 1 |
| **PF00067** | p450 | 1/18 | 50/13933 | 0.0626988116998735 | 0.0788791502030666 | 0.0255479028997787 | OG0000104 | 1 |
| **PF12678** | zf-rbx1 | 1/18 | 59/13933 | 0.0735826983583495 | 0.0896789136242385 | 0.029045801983559 | OG0000006 | 1 |
| **PF00005** | ABC_tran | 1/18 | 82/13933 | 0.100857343393629 | 0.115689305657398 | 0.0374702204558374 | OG0000035 | 1 |
| **PF07690** | MFS_1 | 1/18 | 82/13933 | 0.100857343393629 | 0.115689305657398 | 0.0374702204558374 | OG0000189 | 1 |
| **PF13923** | zf-C3HC4_2 | 1/18 | 90/13933 | 0.110165089484375 | 0.122755385425446 | 0.039758829287594 | OG0000006 | 1 |
| **PF00097** | zf-C3HC4 | 1/18 | 97/13933 | 0.118234647921215 | 0.128087535247983 | 0.041485841375865 | OG0000006 | 1 |
| **PF00004** | AAA | 1/18 | 103/13933 | 0.125096354188291 | 0.13185831927955 | 0.0427071479447934 | OG0000538 | 1 |
| **PF13639** | zf-RING_2 | 1/18 | 116/13933 | 0.139790773028803 | 0.143469477582193 | 0.0464678469901839 | OG0000006 | 1 |
| **PF00076** | RRM_1 | 1/18 | 145/13933 | 0.171735053160299 | 0.171735053160299 | 0.055622689282688 | OG0000088 | 1 |

**Table S4c. Related to Figure 3. Enriched GO terms in orthogroups contracted in Draparnaldia with respect to Chlamydomonas. GO terms were not significant.**

| | GO | orthogroup | GO_term |
|---|---|---|---|
| | <chr> | <chr> | <chr> |
| 1 | GO:0005524 | OG0000029 | ATP binding |
| 2 | GO:0016887 | OG0000029 | ATP hydrolysis activity |
| 3 | GO:0140359 | OG0000029 | ABC-type transporter activity |
| 4 | GO:0016020 | OG0000029 | membrane |

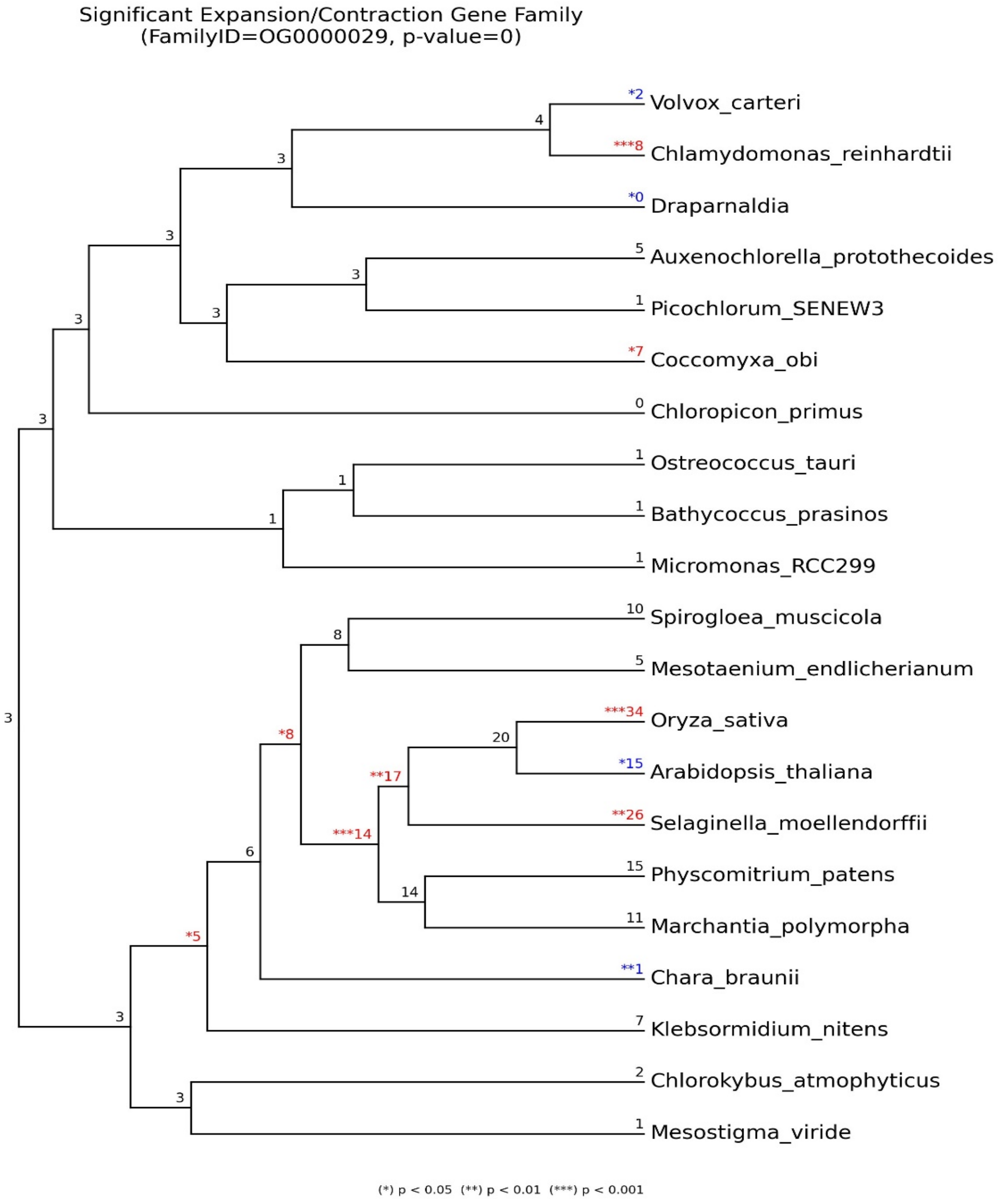

**Table S4d. Related to Figure 3. Enriched GO terms in orthogroups expanded in Chara with respect to Mesostigma.**

| ID | Description | GeneRatio | BgRatio | pvalue | p.adjust | qvalue | geneID | Count |
|---|---|---|---|---|---|---|---|---|
| GO:0048768 | root hair cell tip growth | 2/25 | 8/12891 | 0.000100384966387425 | 0.0564163511097331 | 0.0382519556128926 | OG0000169/OG0000257 | 2 |
| GO:0048767 | root hair elongation | 2/25 | 21/12891 | 0.000741338862391369 | 0.0576816819095214 | 0.0391098873407881 | OG0000169/OG0000257 | 2 |
| GO:0030036 | actin cytoskeleton organization | 2/25 | 25/12891 | 0.00105403431330086 | 0.0576816819095214 | 0.0391098873407881 | OG0000169/OG0000257 | 2 |
| GO:0005856 | cytoskeleton | 3/25 | 107/12891 | 0.00111959285630745 | 0.0576816819095214 | 0.0391098873407881 | OG0000169/OG0000219/OG0000257 | 3 |
| GO:0000325 | plant-type vacuole | 4/25 | 249/12891 | 0.00124879228818768 | 0.0576816819095214 | 0.0391098873407881 | OG0000169/OG0000219/OG0000257/OG0003793 | 4 |
| GO:0030029 | actin filament-based process | 2/25 | 30/12891 | 0.00151930110262697 | 0.0576816819095214 | 0.0391098873407881 | OG0000169/OG0000257 | 2 |
| GO:0080147 | root hair cell development | 2/25 | 31/12891 | 0.00162215336893547 | 0.0576816819095214 | 0.0391098873407881 | OG0000169/OG0000257 | 2 |
| GO:0000146 | microfilament motor activity | 1/25 | 1/12891 | 0.00193933752230202 | 0.0576816819095214 | 0.0391098873407881 | OG0000169 | 1 |
| GO:0004097 | catechol oxidase activity | 1/25 | 1/12891 | 0.00193933752230202 | 0.0576816819095214 | 0.0391098873407881 | OG0000555 | 1 |
| GO:0004630 | phospholipase D activity | 1/25 | 1/12891 | 0.00193933752230202 | 0.0576816819095214 | 0.0391098873407881 | OG0000219 | 1 |
| GO:0016034 | maleylacetoacetate isomerase activity | 1/25 | 1/12891 | 0.00193933752230202 | 0.0576816819095214 | 0.0391098873407881 | OG0002466 | 1 |
| GO:0016461 | unconventional myosin complex | 1/25 | 1/12891 | 0.00193933752230202 | 0.0576816819095214 | 0.0391098873407881 | OG0000169 | 1 |
| GO:0030050 | vesicle transport along actin filament | 1/25 | 1/12891 | 0.00193933752230202 | 0.0576816819095214 | 0.0391098873407881 | OG0000169 | 1 |
| GO:0030742 | GTP-dependent protein binding | 1/25 | 1/12891 | 0.00193933752230202 | 0.0576816819095214 | 0.0391098873407881 | OG0000169 | 1 |
| GO:0060151 | peroxisome localization | 1/25 | 1/12891 | 0.00193933752230202 | 0.0576816819095214 | 0.0391098873407881 | OG0000169 | 1 |
| GO:0070290 | N-acylphosphatidylethanolamine-specific phospholipase D activity | 1/25 | 1/12891 | 0.00193933752230202 | 0.0576816819095214 | 0.0391098873407881 | OG0000219 | 1 |
| GO:0090395 | plant cell papilla | 1/25 | 1/12891 | 0.00193933752230202 | 0.0576816819095214 | 0.0391098873407881 | OG0000219 | 1 |
| GO:0099518 | vesicle cytoskeletal trafficking | 1/25 | 1/12891 | 0.00193933752230202 | 0.0576816819095214 | 0.0391098873407881 | OG0000169 | 1 |
| GO:0042995 | cell projection | 2/25 | 34/12891 | 0.00195009244889841 | 0.0576816819095214 | 0.0391098873407881 | OG0000169/OG0000219 | 2 |
| GO:0048764 | trichoblast maturation | 2/25 | 39/12891 | 0.0025605532259487 | 0.0604940574091081 | 0.0410167611577021 | OG0000169/OG0000257 | 2 |
| GO:0048765 | root hair cell differentiation | 2/25 | 39/12891 | 0.0025605532259487 | 0.0604940574091081 | 0.0410167611577021 | OG0000169/OG0000257 | 2 |
| GO:0048469 | cell maturation | 2/25 | 40/12891 | 0.00269212362510951 | 0.0604940574091081 | 0.0410167611577021 | OG0000169/OG0000257 | 2 |
| GO:0010054 | trichoblast differentiation | 2/25 | 41/12891 | 0.00282682610903752 | 0.0604940574091081 | 0.0410167611577021 | OG0000169/OG0000257 | 2 |
| GO:0005773 | vacuole | 4/25 | 324/12891 | 0.00326397644273989 | 0.0604940574091081 | 0.0410167611577021 | OG0000169/OG0000219/OG0000257/OG0003793 | 4 |
| GO:1905392 | plant organ morphogenesis | 3/25 | 161/12891 | 0.00359404470649679 | 0.0604940574091081 | 0.0410167611577021 | OG0000169/OG0000219/OG0000257 | 3 |
| GO:0010053 | root epidermal cell differentiation | 2/25 | 47/12891 | 0.00370016440182755 | 0.0604940574091081 | 0.0410167611577021 | OG0000169/OG0000257 | 2 |
| GO:0009395 | phospholipid catabolic process | 1/25 | 2/12891 | 0.0038750641756724 | 0.0604940574091081 | 0.0410167611577021 | OG0000219 | 1 |
| GO:0010358 | leaf shaping | 1/25 | 2/12891 | 0.0038750641756724 | 0.0604940574091081 | 0.0410167611577021 | OG0000219 | 1 |
| GO:0016157 | sucrose synthase activity | 1/25 | 2/12891 | 0.0038750641756724 | 0.0604940574091081 | 0.0410167611577021 | OG0000433 | 1 |
| GO:0045471 | response to ethanol | 1/25 | 2/12891 | 0.0038750641756724 | 0.0604940574091081 | 0.0410167611577021 | OG0000017 | 1 |
| GO:0046473 | phosphatidic acid metabolic process | 1/25 | 2/12891 | 0.0038750641756724 | 0.0604940574091081 | 0.0410167611577021 | OG0000219 | 1 |
| GO:0051645 | Golgi localization | 1/25 | 2/12891 | 0.0038750641756724 | 0.0604940574091081 | 0.0410167611577021 | OG0000169 | 1 |
| GO:0071963 | establishment or maintenance of cell polarity regulating cell shape | 1/25 | 2/12891 | 0.0038750641756724 | 0.0604940574091081 | 0.0410167611577021 | OG0000169 | 1 |
| GO:1901999 | homogentisate metabolic process | 1/25 | 2/12891 | 0.0038750641756724 | 0.0604940574091081 | 0.0410167611577021 | OG0002466 | 1 |
| GO:1902000 | homogentisate catabolic process | 1/25 | 2/12891 | 0.0038750641756724 | 0.0604940574091081 | 0.0410167611577021 | OG0002466 | 1 |
| GO:2000769 | regulation of establishment or maintenance of cell polarity regulating cell shape | 1/25 | 2/12891 | 0.0038750641756724 | 0.0604940574091081 | 0.0410167611577021 | OG0000169 | 1 |
| GO:0009932 | cell tip growth | 2/25 | 55/12891 | 0.00503505956200764 | 0.0694391225280165 | 0.0470817800246151 | OG0000169/OG0000257 | 2 |
| GO:0090627 | plant epidermal cell differentiation | 2/25 | 57/12891 | 0.00539861355935922 | 0.0694391225280165 | 0.0470817800246151 | OG0000169/OG0000257 | 2 |
| GO:0005544 | calcium-dependent phospholipid binding | 1/25 | 3/12891 | 0.00580718640358857 | 0.0694391225280165 | 0.0470817800246151 | OG0000705 | 1 |
| GO:0005782 | peroxisomal matrix | 1/25 | 3/12891 | 0.00580718640358857 | 0.0694391225280165 | 0.0470817800246151 | OG0000017 | 1 |
| GO:0006542 | glutamine biosynthetic process | 1/25 | 3/12891 | 0.00580718640358857 | 0.0694391225280165 | 0.0470817800246151 | OG0000306 | 1 |
| GO:0030048 | actin filament-based movement | 1/25 | 3/12891 | 0.00580718640358857 | 0.0694391225280165 | 0.0470817800246151 | OG0000169 | 1 |
| GO:0031907 | microbody lumen | 1/25 | 3/12891 | 0.00580718640358857 | 0.0694391225280165 | 0.0470817800246151 | OG0000017 | 1 |
| GO:0051646 | mitochondrion localization | 1/25 | 3/12891 | 0.00580718640358857 | 0.0694391225280165 | 0.0470817800246151 | OG0000169 | 1 |
| GO:0090436 | leaf pavement cell development | 1/25 | 3/12891 | 0.00580718640358857 | 0.0694391225280165 | 0.0470817800246151 | OG0000169 | 1 |
| GO:0099515 | actin filament-based transport | 1/25 | 3/12891 | 0.00580718640358857 | 0.0694391225280165 | 0.0470817800246151 | OG0000169 | 1 |
| GO:0101031 | chaperone complex | 1/25 | 3/12891 | 0.00580718640358857 | 0.0694391225280165 | 0.0470817800246151 | OG0000017 | 1 |
| GO:0005509 | calcium ion binding | 3/25 | 198/12891 | 0.00639943001098898 | 0.073685921675475 | 0.0499612355244839 | OG0000153/OG0000219/OG0000705 | 3 |
| GO:0048588 | developmental cell growth | 2/25 | 65/12891 | 0.00696946479950357 | 0.073685921675475 | 0.0499612355244839 | OG0000169/OG0000257 | 2 |
| GO:0004252 | serine-type endopeptidase activity | 2/25 | 67/12891 | 0.00739089466133566 | 0.073685921675475 | 0.0499612355244839 | OG0000564/OG0001496 | 2 |
| GO:0006559 | L-phenylalanine catabolic process | 1/25 | 4/12891 | 0.00773571063852851 | 0.073685921675475 | 0.0499612355244839 | OG0002466 | 1 |
| GO:0007097 | nuclear migration | 1/25 | 4/12891 | 0.00773571063852851 | 0.073685921675475 | 0.0499612355244839 | OG0000169 | 1 |
| GO:0009407 | toxin catabolic process | 1/25 | 4/12891 | 0.00773571063852851 | 0.073685921675475 | 0.0499612355244839 | OG0002466 | 1 |
| GO:0016459 | myosin complex | 1/25 | 4/12891 | 0.00773571063852851 | 0.073685921675475 | 0.0499612355244839 | OG0000169 | 1 |
| GO:0019336 | phenol-containing compound catabolic process | 1/25 | 4/12891 | 0.00773571063852851 | 0.073685921675475 | 0.0499612355244839 | OG0002466 | 1 |
| GO:0030705 | cytoskeleton-dependent intracellular transport | 1/25 | 4/12891 | 0.00773571063852851 | 0.073685921675475 | 0.0499612355244839 | OG0000169 | 1 |
| GO:0046466 | membrane lipid catabolic process | 1/25 | 4/12891 | 0.00773571063852851 | 0.073685921675475 | 0.0499612355244839 | OG0000219 | 1 |
| GO:0046688 | response to copper ion | 1/25 | 4/12891 | 0.00773571063852851 | 0.073685921675475 | 0.0499612355244839 | OG0000017 | 1 |
| GO:1902222 | erythrose 4-phosphate/phosphoenolpyruvate family amino acid catabolic process | 1/25 | 4/12891 | 0.00773571063852851 | 0.073685921675475 | 0.0499612355244839 | OG0002466 | 1 |
| GO:0071695 | anatomical structure maturation | 2/25 | 72/12891 | 0.00849376853600453 | 0.0793790101479782 | 0.0538213179875784 | OG0000169/OG0000257 | 2 |
| GO:0070887 | cellular response to chemical stimulus | 4/25 | 431/12891 | 0.00892786192126134 | 0.0793790101479782 | 0.0538213179875784 | OG0000017/OG0000219/OG0002466/OG0003793 | 4 |
| GO:0009845 | seed germination | 2/25 | 74/12891 | 0.0089544109789856 | 0.0793790101479782 | 0.0538213179875784 | OG0000219/OG0000257 | 2 |
| GO:0008360 | regulation of cell shape | 1/25 | 5/12891 | 0.00966064330199046 | 0.0793790101479782 | 0.0538213179875784 | OG0000169 | 1 |
| GO:0009074 | aromatic amino acid family catabolic process | 1/25 | 5/12891 | 0.00966064330199046 | 0.0793790101479782 | 0.0538213179875784 | OG0002466 | 1 |
| GO:0035619 | root hair tip | 1/25 | 5/12891 | 0.00966064330199046 | 0.0793790101479782 | 0.0538213179875784 | OG0000169 | 1 |
| GO:0051647 | nucleus localization | 1/25 | 5/12891 | 0.00966064330199046 | 0.0793790101479782 | 0.0538213179875784 | OG0000169 | 1 |
| GO:0007010 | cytoskeleton organization | 2/25 | 80/12891 | 0.0104020589569325 | 0.0793790101479782 | 0.0538213179875784 | OG0000169/OG0000257 | 2 |
| GO:0021700 | developmental maturation | 2/25 | 80/12891 | 0.0104020589569325 | 0.0793790101479782 | 0.0538213179875784 | OG0000169/OG0000257 | 2 |
| GO:0000904 | cell morphogenesis involved in differentiation | 2/25 | 82/12891 | 0.0109062328165057 | 0.0793790101479782 | 0.0538213179875784 | OG0000169/OG0000257 | 2 |
| GO:0090351 | seedling development | 2/25 | 84/12891 | 0.0114210788843144 | 0.0793790101479782 | 0.0538213179875784 | OG0000219/OG0000257 | 2 |
| GO:0004867 | serine-type endopeptidase inhibitor activity | 1/25 | 6/12891 | 0.0115819908045093 | 0.0793790101479782 | 0.0538213179875784 | OG0000394 | 1 |
| GO:0006032 | chitin catabolic process | 1/25 | 6/12891 | 0.0115819908045093 | 0.0793790101479782 | 0.0538213179875784 | OG0000473 | 1 |
| GO:0006558 | L-phenylalanine metabolic process | 1/25 | 6/12891 | 0.0115819908045093 | 0.0793790101479782 | 0.0538213179875784 | OG0002466 | 1 |
| GO:0010466 | negative regulation of peptidase activity | 1/25 | 6/12891 | 0.0115819908045093 | 0.0793790101479782 | 0.0538213179875784 | OG0000394 | 1 |
| GO:0010951 | negative regulation of endopeptidase activity | 1/25 | 6/12891 | 0.0115819908045093 | 0.0793790101479782 | 0.0538213179875784 | OG0000394 | 1 |
| GO:0016859 | cis-trans isomerase activity | 1/25 | 6/12891 | 0.0115819908045093 | 0.0793790101479782 | 0.0538213179875784 | OG0002466 | 1 |
| GO:0030414 | peptidase inhibitor activity | 1/25 | 6/12891 | 0.0115819908045093 | 0.0793790101479782 | 0.0538213179875784 | OG0000394 | 1 |
| GO:0032878 | regulation of establishment or maintenance of cell polarity | 1/25 | 6/12891 | 0.0115819908045093 | 0.0793790101479782 | 0.0538213179875784 | OG0000169 | 1 |
| GO:0035618 | root hair | 1/25 | 6/12891 | 0.0115819908045093 | 0.0793790101479782 | 0.0538213179875784 | OG0000169 | 1 |
| GO:0045861 | negative regulation of proteolysis | 1/25 | 6/12891 | 0.0115819908045093 | 0.0793790101479782 | 0.0538213179875784 | OG0000394 | 1 |
| GO:0061135 | endopeptidase regulator activity | 1/25 | 6/12891 | 0.0115819908045093 | 0.0793790101479782 | 0.0538213179875784 | OG0000394 | 1 |
| GO:1902221 | erythrose 4-phosphate/phosphoenolpyruvate family amino acid metabolic process | 1/25 | 6/12891 | 0.0115819908045093 | 0.0793790101479782 | 0.0538213179875784 | OG0002466 | 1 |
| GO:0046686 | response to cadmium ion | 2/25 | 88/12891 | 0.0124824546072197 | 0.081628890581017 | 0.0553468035031432 | OG0000017/OG0000219 | 2 |
| GO:0003774 | cytoskeletal motor activity | 1/25 | 7/12891 | 0.0134997595456723 | 0.081628890581017 | 0.0553468035031432 | OG0000169 | 1 |
| GO:0004356 | glutamate-ammonia ligase activity | 1/25 | 7/12891 | 0.0134997595456723 | 0.081628890581017 | 0.0553468035031432 | OG0000306 | 1 |
| GO:0004866 | endopeptidase inhibitor activity | 1/25 | 7/12891 | 0.0134997595456723 | 0.081628890581017 | 0.0553468035031432 | OG0000394 | 1 |
| GO:0004869 | cysteine-type endopeptidase inhibitor activity | 1/25 | 7/12891 | 0.0134997595456723 | 0.081628890581017 | 0.0553468035031432 | OG0000394 | 1 |
| GO:0005546 | phosphatidylinositol-4,5-bisphosphate binding | 1/25 | 7/12891 | 0.0134997595456723 | 0.081628890581017 | 0.0553468035031432 | OG0000219 | 1 |
| GO:0006893 | Golgi to plasma membrane transport | 1/25 | 7/12891 | 0.0134997595456723 | 0.081628890581017 | 0.0553468035031432 | OG0000257 | 1 |
| GO:0046470 | phosphatidylcholine metabolic process | 1/25 | 7/12891 | 0.0134997595456723 | 0.081628890581017 | 0.0553468035031432 | OG0000219 | 1 |
| GO:0052548 | regulation of endopeptidase activity | 1/25 | 7/12891 | 0.0134997595456723 | 0.081628890581017 | 0.0553468035031432 | OG0000394 | 1 |
| GO:1902936 | phosphatidylinositol bisphosphate binding | 1/25 | 7/12891 | 0.0134997595456723 | 0.081628890581017 | 0.0553468035031432 | OG0000219 | 1 |
| GO:0005829 | cytosol | 6/25 | 1054/12891 | 0.0135079836726594 | 0.081628890581017 | 0.0553468035031432 | OG0000017/OG0000169/OG0000219/OG0000257/OG0000394/OG0002466 | 6 |
| GO:0071456 | cellular response to hypoxia | 2/25 | 93/12891 | 0.0138677297413682 | 0.0817230492806371 | 0.0554106458879765 | OG0000017/OG0003793 | 2 |
| GO:0036294 | cellular response to decreased oxygen levels | 2/25 | 94/12891 | 0.0141524898034211 | 0.0817230492806371 | 0.0554106458879765 | OG0000017/OG0003793 | 2 |
| GO:0071453 | cellular response to oxygen levels | 2/25 | 94/12891 | 0.0141524898034211 | 0.0817230492806371 | 0.0554106458879765 | OG0000017/OG0003793 | 2 |
| GO:0090558 | plant epidermis development | 2/25 | 96/12891 | 0.0147296337242322 | 0.0817230492806371 | 0.0554106458879765 | OG0000169/OG0000257 | 2 |
| GO:0004364 | glutathione transferase activity | 1/25 | 8/12891 | 0.0154139559141415 | 0.0817230492806371 | 0.0554106458879765 | OG0002466 | 1 |
| GO:0004568 | chitinase activity | 1/25 | 8/12891 | 0.0154139559141415 | 0.0817230492806371 | 0.0554106458879765 | OG0000473 | 1 |
| GO:0007163 | establishment or maintenance of cell polarity | 1/25 | 8/12891 | 0.0154139559141415 | 0.0817230492806371 | 0.0554106458879765 | OG0000169 | 1 |
| GO:0010044 | response to aluminum ion | 1/25 | 8/12891 | 0.0154139559141415 | 0.0817230492806371 | 0.0554106458879765 | OG0000219 | 1 |
| GO:0016998 | cell wall macromolecule catabolic process | 1/25 | 8/12891 | 0.0154139559141415 | 0.0817230492806371 | 0.0554106458879765 | OG0000473 | 1 |
| GO:0051648 | vesicle localization | 1/25 | 8/12891 | 0.0154139559141415 | 0.0817230492806371 | 0.0554106458879765 | OG0000169 | 1 |
| GO:0051650 | establishment of vesicle localization | 1/25 | 8/12891 | 0.0154139559141415 | 0.0817230492806371 | 0.0554106458879765 | OG0000169 | 1 |
| GO:0061134 | peptidase regulator activity | 1/25 | 8/12891 | 0.0154139559141415 | 0.0817230492806371 | 0.0554106458879765 | OG0000394 | 1 |
| GO:0090333 | regulation of stomatal closure | 1/25 | 8/12891 | 0.0154139559141415 | 0.0817230492806371 | 0.0554106458879765 | OG0000219 | 1 |
| GO:0051716 | cellular response to stimulus | 5/25 | 789/12891 | 0.0161830435438573 | 0.0836116575881802 | 0.056691178211128 | OG0000017/OG0000219/OG0000394/OG0002466/OG0003793 | 5 |
| GO:0001666 | response to hypoxia | 2/25 | 101/12891 | 0.0162164958667467 | 0.0836116575881802 | 0.056691178211128 | OG0000017/OG0003793 | 2 |
| GO:0010015 | root morphogenesis | 2/25 | 101/12891 | 0.0162164958667467 | 0.0836116575881802 | 0.056691178211128 | OG0000169/OG0000257 | 2 |
| GO:0009826 | unidimensional cell growth | 2/25 | 102/12891 | 0.0165213314970128 | 0.0844089845574653 | 0.0572317849420443 | OG0000169/OG0000257 | 2 |
| GO:0008519 | ammonium transmembrane transporter activity | 1/25 | 9/12891 | 0.0173245862876686 | 0.0854071709971033 | 0.057908589438006 | OG0000132 | 1 |
| GO:0051346 | negative regulation of hydrolase activity | 1/25 | 9/12891 | 0.0173245862876686 | 0.0854071709971033 | 0.057908589438006 | OG0000394 | 1 |
| GO:0052547 | regulation of peptidase activity | 1/25 | 9/12891 | 0.0173245862876686 | 0.0854071709971033 | 0.057908589438006 | OG0000394 | 1 |
| GO:0072488 | ammonium transmembrane transport | 1/25 | 9/12891 | 0.0173245862876686 | 0.0854071709971033 | 0.057908589438006 | OG0000132 | 1 |
| GO:0036293 | response to decreased oxygen levels | 2/25 | 106/12891 | 0.0177652204489334 | 0.0868178599330482 | 0.0588650782838799 | OG0000017/OG0003793 | 2 |
| GO:0070482 | response to oxygen levels | 2/25 | 107/12891 | 0.0180822800221262 | 0.0871628326823318 | 0.0590989800168648 | OG0000017/OG0003793 | 2 |
| GO:0043228 | non-membrane-bounded organelle | 4/25 | 536/12891 | 0.0186679244403389 | 0.0871628326823318 | 0.0590989800168648 | OG0000017/OG0000169/OG0000219/OG0000257 | 4 |
| GO:0043232 | intracellular non-membrane-bounded organelle | 4/25 | 536/12891 | 0.0186679244403389 | 0.0871628326823318 | 0.0590989800168648 | OG0000017/OG0000169/OG0000219/OG0000257 | 4 |
| GO:0030234 | enzyme regulator activity | 2/25 | 109/12891 | 0.0187236353574896 | 0.0871628326823318 | 0.0590989800168648 | OG0000219/OG0000394 | 2 |
| GO:0048468 | cell development | 2/25 | 109/12891 | 0.0187236353574896 | 0.0871628326823318 | 0.0590989800168648 | OG0000169/OG0000257 | 2 |
| GO:0005200 | structural constituent of cytoskeleton | 1/25 | 10/12891 | 0.0192316570331124 | 0.0871628326823318 | 0.0590989800168648 | OG0000257 | 1 |
| GO:0005985 | sucrose metabolic process | 1/25 | 10/12891 | 0.0192316570331124 | 0.0871628326823318 | 0.0590989800168648 | OG0000433 | 1 |
| GO:0006928 | movement of cell or subcellular component | 1/25 | 10/12891 | 0.0192316570331124 | 0.0871628326823318 | 0.0590989800168648 | OG0000169 | 1 |
| GO:0046685 | response to arsenic-containing substance | 1/25 | 10/12891 | 0.0192316570331124 | 0.0871628326823318 | 0.0590989800168648 | OG0000017 | 1 |
| GO:0060560 | developmental growth involved in morphogenesis | 2/25 | 112/12891 | 0.0197036127860136 | 0.0885874430859171 | 0.0600649080297846 | OG0000169/OG0000257 | 2 |
| GO:0000427 | plastid-encoded plastid RNA polymerase complex | 1/25 | 11/12891 | 0.0211351745064584 | 0.0920772718808498 | 0.0624311152291958 | OG0000017 | 1 |
| GO:0009404 | toxin metabolic process | 1/25 | 11/12891 | 0.0211351745064584 | 0.0920772718808498 | 0.0624311152291958 | OG0002466 | 1 |
| GO:0051286 | cell tip | 1/25 | 11/12891 | 0.0211351745064584 | 0.0920772718808498 | 0.0624311152291958 | OG0000169 | 1 |
| GO:0060187 | cell pole | 1/25 | 11/12891 | 0.0211351745064584 | 0.0920772718808498 | 0.0624311152291958 | OG0000169 | 1 |
| GO:0042221 | response to chemical | 5/25 | 847/12891 | 0.0213927928670317 | 0.0924826891636293 | 0.0627060001446597 | OG0000017/OG0000219/OG0000257/OG0002466/OG0003793 | 5 |
| GO:0098772 | molecular function regulator activity | 2/25 | 119/12891 | 0.0220726161518586 | 0.0934965291625974 | 0.0633934136670917 | OG0000219/OG0000394 | 2 |
| GO:0009789 | positive regulation of abscisic acid-activated signaling pathway | 1/25 | 12/12891 | 0.0230351450528362 | 0.0934965291625974 | 0.0633934136670917 | OG0000219 | 1 |
| GO:0010091 | trichome branching | 1/25 | 12/12891 | 0.0230351450528362 | 0.0934965291625974 | 0.0633934136670917 | OG0000169 | 1 |
| GO:0015074 | DNA integration | 1/25 | 12/12891 | 0.0230351450528362 | 0.0934965291625974 | 0.0633934136670917 | OG0003409 | 1 |
| GO:0051020 | GTPase binding | 1/25 | 12/12891 | 0.0230351450528362 | 0.0934965291625974 | 0.0633934136670917 | OG0000169 | 1 |
| GO:0098876 | vesicle-mediated transport to the plasma membrane | 1/25 | 12/12891 | 0.0230351450528362 | 0.0934965291625974 | 0.0633934136670917 | OG0000257 | 1 |
| GO:0010038 | response to metal ion | 2/25 | 122/12891 | 0.0231225919403864 | 0.0934965291625974 | 0.0633934136670917 | OG0000017/OG0000219 | 2 |
| GO:0005911 | cell-cell junction | 3/25 | 321/12891 | 0.0234573142560965 | 0.0934965291625974 | 0.0633934136670917 | OG0000169/OG0000219/OG0000257 | 3 |
| GO:0009506 | plasmodesma | 3/25 | 321/12891 | 0.0234573142560965 | 0.0934965291625974 | 0.0633934136670917 | OG0000169/OG0000219/OG0000257 | 3 |
| GO:0055044 | symplast | 3/25 | 321/12891 | 0.0234573142560965 | 0.0934965291625974 | 0.0633934136670917 | OG0000169/OG0000219/OG0000257 | 3 |
| GO:0070161 | anchoring junction | 3/25 | 321/12891 | 0.0234573142560965 | 0.0934965291625974 | 0.0633934136670917 | OG0000169/OG0000219/OG0000257 | 3 |
| GO:0030054 | cell junction | 3/25 | 322/12891 | 0.023647896283457 | 0.0935923782486117 | 0.0634584021839248 | OG0000169/OG0000219/OG0000257 | 3 |
| GO:0009653 | anatomical structure morphogenesis | 3/25 | 326/12891 | 0.0244187799644261 | 0.0959675128671849 | 0.0650688137439988 | OG0000169/OG0000219/OG0000257 | 3 |
| GO:0051015 | actin filament binding | 1/25 | 13/12891 | 0.024931575006534 | 0.0959694873539185 | 0.0650701525044362 | OG0000169 | 1 |
| GO:1901616 | organic hydroxy compound catabolic process | 1/25 | 13/12891 | 0.024931575006534 | 0.0959694873539185 | 0.0650701525044362 | OG0002466 | 1 |
| GO:1901981 | phosphatidylinositol phosphate binding | 1/25 | 13/12891 | 0.024931575006534 | 0.0959694873539185 | 0.0650701525044362 | OG0000219 | 1 |
| GO:0042579 | microbody | 2/25 | 129/12891 | 0.0256516057721236 | 0.0980694043804997 | 0.0664939583924722 | OG0000017/OG0000169 | 2 |
| GO:0004620 | phospholipase activity | 1/25 | 14/12891 | 0.026824470691022 | 0.101176862606405 | 0.0686009070303777 | OG0000219 | 1 |
| GO:0090332 | stomatal closure | 1/25 | 14/12891 | 0.026824470691022 | 0.101176862606405 | 0.0686009070303777 | OG0000219 | 1 |
| GO:0050790 | regulation of catalytic activity | 2/25 | 134/12891 | 0.0275244254759335 | 0.103124847449831 | 0.0699216984020206 | OG0000219/OG0000394 | 2 |
| GO:0005777 | peroxisome | 2/25 | 135/12891 | 0.0279055080879727 | 0.103860235400269 | 0.0704203131951628 | OG0000017/OG0000169 | 2 |
| GO:0006749 | glutathione metabolic process | 1/25 | 15/12891 | 0.0287138384189651 | 0.106165639417489 | 0.0719834453439431 | OG0002466 | 1 |
| GO:0000902 | cell morphogenesis | 2/25 | 138/12891 | 0.029061629653146 | 0.106749254020053 | 0.0723791533157128 | OG0000169/OG0000257 | 2 |
| GO:0099402 | plant organ development | 3/25 | 351/12891 | 0.0295470148101244 | 0.107481391779004 | 0.072875564382842 | OG0000169/OG0000219/OG0000257 | 3 |
| GO:0051301 | cell division | 2/25 | 141/12891 | 0.0302368724774883 | 0.107481391779004 | 0.072875564382842 | OG0000169/OG0000257 | 2 |
| GO:0010218 | response to far red light | 1/25 | 16/12891 | 0.030599684492243 | 0.107481391779004 | 0.072875564382842 | OG0000257 | 1 |
| GO:0015629 | actin cytoskeleton | 1/25 | 16/12891 | 0.030599684492243 | 0.107481391779004 | 0.072875564382842 | OG0000169 | 1 |
| GO:0030136 | clathrin-coated vesicle | 1/25 | 16/12891 | 0.030599684492243 | 0.107481391779004 | 0.072875564382842 | OG0000219 | 1 |
| GO:0051259 | protein complex oligomerization | 1/25 | 16/12891 | 0.030599684492243 | 0.107481391779004 | 0.072875564382842 | OG0000017 | 1 |
| GO:1901606 | alpha-amino acid catabolic process | 1/25 | 16/12891 | 0.030599684492243 | 0.107481391779004 | 0.072875564382842 | OG0002466 | 1 |
| GO:0048366 | leaf development | 2/25 | 143/12891 | 0.0310308711356668 | 0.108318941479781 | 0.0734434478660436 | OG0000169/OG0000219 | 2 |
| GO:0065009 | regulation of molecular function | 2/25 | 146/12891 | 0.0322374464018613 | 0.110635712384881 | 0.0750142871011926 | OG0000219/OG0000394 | 2 |
| GO:0006892 | post-Golgi vesicle-mediated transport | 1/25 | 17/12891 | 0.0324820152019667 | 0.110635712384881 | 0.0750142871011926 | OG0000257 | 1 |
| GO:0016702 | oxidoreductase activity, acting on single donors with incorporation of molecular oxygen, incorporation of two atoms of oxygen | 1/25 | 17/12891 | 0.0324820152019667 | 0.110635712384881 | 0.0750142871011926 | OG0000108 | 1 |
| GO:0042542 | response to hydrogen peroxide | 1/25 | 17/12891 | 0.0324820152019667 | 0.110635712384881 | 0.0750142871011926 | OG0000017 | 1 |
| GO:0016049 | cell growth | 2/25 | 151/12891 | 0.034289356866815 | 0.114265031346836 | 0.0774750727618553 | OG0000169/OG0000257 | 2 |
| GO:0007015 | actin filament organization | 1/25 | 18/12891 | 0.0343608368284969 | 0.114265031346836 | 0.0774750727618553 | OG0000169 | 1 |
| GO:0009072 | aromatic amino acid family metabolic process | 1/25 | 18/12891 | 0.0343608368284969 | 0.114265031346836 | 0.0774750727618553 | OG0002466 | 1 |
| GO:0046434 | organophosphate catabolic process | 1/25 | 18/12891 | 0.0343608368284969 | 0.114265031346836 | 0.0774750727618553 | OG0000219 | 1 |
| GO:0032787 | monocarboxylic acid metabolic process | 2/25 | 153/12891 | 0.0351242501171395 | 0.116116638622544 | 0.0787305172904304 | OG0000219/OG0002466 | 2 |
| GO:0016298 | lipase activity | 1/25 | 19/12891 | 0.0362361556414612 | 0.117715141447984 | 0.0798143495114631 | OG0000219 | 1 |
| GO:0030133 | transport vesicle | 1/25 | 19/12891 | 0.0362361556414612 | 0.117715141447984 | 0.0798143495114631 | OG0000169 | 1 |
| GO:0098590 | plasma membrane region | 1/25 | 19/12891 | 0.0362361556414612 | 0.117715141447984 | 0.0798143495114631 | OG0000219 | 1 |
| GO:0009651 | response to salt stress | 2/25 | 158/12891 | 0.0372462000631044 | 0.120300945031406 | 0.0815676008641488 | OG0000017/OG0000219 | 2 |
| GO:0000723 | telomere maintenance | 1/25 | 20/12891 | 0.0381079778997708 | 0.121685702157223 | 0.0825065071753412 | OG0000381 | 1 |
| GO:0072329 | monocarboxylic acid catabolic process | 1/25 | 20/12891 | 0.0381079778997708 | 0.121685702157223 | 0.0825065071753412 | OG0002466 | 1 |
| GO:0005783 | endoplasmic reticulum | 3/25 | 393/12891 | 0.039361745688589 | 0.12497910212987 | 0.0847395298202153 | OG0000017/OG0000219/OG0000257 | 3 |
| GO:0006508 | proteolysis | 3/25 | 395/12891 | 0.0398663181409755 | 0.125512213053753 | 0.0851009948032565 | OG0000394/OG0000650/OG0001496 | 3 |
| GO:0006972 | hyperosmotic response | 1/25 | 21/12891 | 0.0399763098516402 | 0.125512213053753 | 0.0851009948032565 | OG0000017 | 1 |
| GO:0006950 | response to stress | 5/25 | 1004/12891 | 0.0407683053618308 | 0.127287708963049 | 0.0863048335729984 | OG0000017/OG0000219/OG0000257/OG0000394/OG0003793 | 5 |
| GO:0048589 | developmental growth | 2/25 | 167/12891 | 0.0411878239551321 | 0.127887055595493 | 0.0867112083265938 | OG0000169/OG0000257 | 2 |
| GO:0006979 | response to oxidative stress | 2/25 | 168/12891 | 0.0416352629390979 | 0.128495795884404 | 0.0871239522572656 | OG0000017/OG0000219 | 2 |
| GO:0009063 | cellular amino acid catabolic process | 1/25 | 22/12891 | 0.0418411577346012 | 0.128495795884404 | 0.0871239522572656 | OG0002466 | 1 |
| GO:0006575 | cellular modified amino acid metabolic process | 1/25 | 23/12891 | 0.0437025277755237 | 0.13134128668366 | 0.0890532792273549 | OG0002466 | 1 |
| GO:0010114 | response to red light | 1/25 | 23/12891 | 0.0437025277755237 | 0.13134128668366 | 0.0890532792273549 | OG0000257 | 1 |
| GO:0030162 | regulation of proteolysis | 1/25 | 23/12891 | 0.0437025277755237 | 0.13134128668366 | 0.0890532792273549 | OG0000394 | 1 |
| GO:0031965 | nuclear membrane | 1/25 | 23/12891 | 0.0437025277755237 | 0.13134128668366 | 0.0890532792273549 | OG0000169 | 1 |
| GO:0005615 | extracellular space | 1/25 | 24/12891 | 0.0455604261906322 | 0.132206442089239 | 0.0896398801953636 | OG0000394 | 1 |
| GO:0008081 | phosphoric diester hydrolase activity | 1/25 | 24/12891 | 0.0455604261906322 | 0.132206442089239 | 0.0896398801953636 | OG0000219 | 1 |
| GO:0016684 | oxidoreductase activity, acting on peroxide as acceptor | 1/25 | 24/12891 | 0.0455604261906322 | 0.132206442089239 | 0.0896398801953636 | OG0002466 | 1 |
| GO:0018958 | phenol-containing compound metabolic process | 1/25 | 24/12891 | 0.0455604261906322 | 0.132206442089239 | 0.0896398801953636 | OG0002466 | 1 |
| GO:0031267 | small GTPase binding | 1/25 | 24/12891 | 0.0455604261906322 | 0.132206442089239 | 0.0896398801953636 | OG0000169 | 1 |
| GO:0042537 | benzene-containing compound metabolic process | 1/25 | 24/12891 | 0.0455604261906322 | 0.132206442089239 | 0.0896398801953636 | OG0002466 | 1 |
| GO:0033554 | cellular response to stress | 3/25 | 417/12891 | 0.0456370992265346 | 0.132206442089239 | 0.0896398801953636 | OG0000017/OG0000394/OG0003793 | 3 |
| GO:0006281 | DNA repair | 2/25 | 179/12891 | 0.0466784111803319 | 0.134529574786393 | 0.091215032913793 | OG0000381/OG0000394 | 2 |
| GO:0009628 | response to abiotic stimulus | 4/25 | 717/12891 | 0.0473270617757199 | 0.13526472518915 | 0.0917134866425781 | OG0000017/OG0000219/OG0000257/OG0003793 | 4 |
| GO:0003678 | DNA helicase activity | 1/25 | 25/12891 | 0.0474148591855207 | 0.13526472518915 | 0.0917134866425781 | OG0000381 | 1 |
| GO:0006970 | response to osmotic stress | 2/25 | 183/12891 | 0.048566125834891 | 0.13784930666267 | 0.0934659093685833 | OG0000017/OG0000219 | 2 |
| GO:0044242 | cellular lipid catabolic process | 1/25 | 26/12891 | 0.0492658329551742 | 0.138436990604039 | 0.0938643764724897 | OG0000219 | 1 |
| GO:0051656 | establishment of organelle localization | 1/25 | 26/12891 | 0.0492658329551742 | 0.138436990604039 | 0.0938643764724897 | OG0000169 | 1 |
| GO:0022604 | regulation of cell morphogenesis | 1/25 | 27/12891 | 0.0511133536839826 | 0.142913954081583 | 0.0968989801995376 | OG0000169 | 1 |
| GO:0048827 | phyllome development | 2/25 | 191/12891 | 0.0524247565382396 | 0.1458550157153 | 0.0988940170236723 | OG0000169/OG0000219 | 2 |
| GO:0009967 | positive regulation of signal transduction | 1/25 | 28/12891 | 0.0529574275457597 | 0.145892520983907 | 0.0989194467057019 | OG0000219 | 1 |
| GO:0010286 | heat acclimation | 1/25 | 28/12891 | 0.0529574275457597 | 0.145892520983907 | 0.0989194467057019 | OG0000017 | 1 |
| GO:0022622 | root system development | 2/25 | 193/12891 | 0.0534063805989121 | 0.146411638519944 | 0.0992714237576697 | OG0000169/OG0000257 | 2 |
| GO:0048364 | root development | 2/25 | 195/12891 | 0.0543946666984901 | 0.147351723040733 | 0.0999088288832091 | OG0000169/OG0000257 | 2 |
| GO:0012505 | endomembrane system | 4/25 | 751/12891 | 0.0545127766873291 | 0.147351723040733 | 0.0999088288832091 | OG0000017/OG0000169/OG0000219/OG0000257 | 4 |
| GO:0010090 | trichome morphogenesis | 1/25 | 29/12891 | 0.0547980607037601 | 0.147351723040733 | 0.0999088288832091 | OG0000169 | 1 |
| GO:0023056 | positive regulation of signaling | 1/25 | 29/12891 | 0.0547980607037601 | 0.147351723040733 | 0.0999088288832091 | OG0000219 | 1 |
| GO:0003779 | actin binding | 1/25 | 30/12891 | 0.0566352593106958 | 0.15013686666326 | 0.101797238681588 | OG0000169 | 1 |
| GO:0010119 | regulation of stomatal movement | 1/25 | 30/12891 | 0.0566352593106958 | 0.15013686666326 | 0.101797238681588 | OG0000219 | 1 |
| GO:0048467 | gynoecium development | 1/25 | 30/12891 | 0.0566352593106958 | 0.15013686666326 | 0.101797238681588 | OG0000169 | 1 |
| GO:0016757 | glycosyltransferase activity | 2/25 | 203/12891 | 0.0584132035891031 | 0.154123100549652 | 0.104500023223402 | OG0000433/OG0000512 | 2 |
| GO:0009873 | ethylene-activated signaling pathway | 1/25 | 32/12891 | 0.0602993774296163 | 0.155450688602956 | 0.105400167211594 | OG0000219 | 1 |
| GO:0010647 | positive regulation of cell communication | 1/25 | 32/12891 | 0.0602993774296163 | 0.155450688602956 | 0.105400167211594 | OG0000219 | 1 |
| GO:0042644 | chloroplast nucleoid | 1/25 | 32/12891 | 0.0602993774296163 | 0.155450688602956 | 0.105400167211594 | OG0000017 | 1 |
| GO:0048532 | anatomical structure arrangement | 1/25 | 32/12891 | 0.0602993774296163 | 0.155450688602956 | 0.105400167211594 | OG0000219 | 1 |
| GO:0120025 | plasma membrane bounded cell projection | 1/25 | 32/12891 | 0.0602993774296163 | 0.155450688602956 | 0.105400167211594 | OG0000169 | 1 |
| GO:0097305 | response to alcohol | 2/25 | 208/12891 | 0.0609766881886444 | 0.156478989780905 | 0.106097385841333 | OG0000017/OG0000219 | 2 |
| GO:0004857 | enzyme inhibitor activity | 1/25 | 33/12891 | 0.0621263091944702 | 0.156569442902656 | 0.106158715734709 | OG0000394 | 1 |
| GO:0030695 | GTPase regulator activity | 1/25 | 33/12891 | 0.0621263091944702 | 0.156569442902656 | 0.106158715734709 | OG0000219 | 1 |
| GO:0042646 | plastid nucleoid | 1/25 | 33/12891 | 0.0621263091944702 | 0.156569442902656 | 0.106158715734709 | OG0000017 | 1 |
| GO:0060589 | nucleoside-triphosphatase regulator activity | 1/25 | 33/12891 | 0.0621263091944702 | 0.156569442902656 | 0.106158715734709 | OG0000219 | 1 |
| GO:0050896 | response to stimulus | 6/25 | 1508/12891 | 0.0638712124035704 | 0.158325132042672 | 0.107349124928727 | OG0000017/OG0000219/OG0000257/OG0000394/OG0002466/OG0003793 | 6 |
| GO:0009644 | response to high light intensity | 1/25 | 34/12891 | 0.0639498309440329 | 0.158325132042672 | 0.107349124928727 | OG0000257 | 1 |
| GO:0043086 | negative regulation of catalytic activity | 1/25 | 34/12891 | 0.0639498309440329 | 0.158325132042672 | 0.107349124928727 | OG0000394 | 1 |
| GO:0071369 | cellular response to ethylene stimulus | 1/25 | 34/12891 | 0.0639498309440329 | 0.158325132042672 | 0.107349124928727 | OG0000219 | 1 |
| GO:0000428 | DNA-directed RNA polymerase complex | 1/25 | 35/12891 | 0.0657699486885645 | 0.158638245334649 | 0.107561425006823 | OG0000017 | 1 |
| GO:0009295 | nucleoid | 1/25 | 35/12891 | 0.0657699486885645 | 0.158638245334649 | 0.107561425006823 | OG0000017 | 1 |
| GO:0009787 | regulation of abscisic acid-activated signaling pathway | 1/25 | 35/12891 | 0.0657699486885645 | 0.158638245334649 | 0.107561425006823 | OG0000219 | 1 |
| GO:0098869 | cellular oxidant detoxification | 1/25 | 35/12891 | 0.0657699486885645 | 0.158638245334649 | 0.107561425006823 | OG0002466 | 1 |
| GO:1901419 | regulation of response to alcohol | 1/25 | 35/12891 | 0.0657699486885645 | 0.158638245334649 | 0.107561425006823 | OG0000219 | 1 |
| GO:1905957 | regulation of cellular response to alcohol | 1/25 | 35/12891 | 0.0657699486885645 | 0.158638245334649 | 0.107561425006823 | OG0000219 | 1 |
| GO:0009266 | response to temperature stimulus | 2/25 | 218/12891 | 0.0662191394486289 | 0.159039129786878 | 0.107833236529031 | OG0000017/OG0000219 | 2 |
| GO:0006643 | membrane lipid metabolic process | 1/25 | 36/12891 | 0.0675866686078848 | 0.160268809103929 | 0.108666995496577 | OG0000219 | 1 |
| GO:0010026 | trichome differentiation | 1/25 | 36/12891 | 0.0675866686078848 | 0.160268809103929 | 0.108666995496577 | OG0000169 | 1 |
| GO:0051336 | regulation of hydrolase activity | 1/25 | 36/12891 | 0.0675866686078848 | 0.160268809103929 | 0.108666995496577 | OG0000394 | 1 |
| GO:0030135 | coated vesicle | 1/25 | 37/12891 | 0.0693999967513937 | 0.163191624160181 | 0.110648750601209 | OG0000219 | 1 |
| GO:0030880 | RNA polymerase complex | 1/25 | 37/12891 | 0.0693999967513937 | 0.163191624160181 | 0.110648750601209 | OG0000017 | 1 |
| GO:0005096 | GTPase activator activity | 1/25 | 38/12891 | 0.0712099391880838 | 0.163346880913074 | 0.11075401927427 | OG0000219 | 1 |
| GO:0005516 | calmodulin binding | 1/25 | 38/12891 | 0.0712099391880838 | 0.163346880913074 | 0.11075401927427 | OG0000169 | 1 |
| GO:0035091 | phosphatidylinositol binding | 1/25 | 38/12891 | 0.0712099391880838 | 0.163346880913074 | 0.11075401927427 | OG0000219 | 1 |
| GO:0043621 | protein self-association | 1/25 | 38/12891 | 0.0712099391880838 | 0.163346880913074 | 0.11075401927427 | OG0000017 | 1 |
| GO:0097435 | supramolecular fiber organization | 1/25 | 38/12891 | 0.0712099391880838 | 0.163346880913074 | 0.11075401927427 | OG0000169 | 1 |
| GO:1990748 | cellular detoxification | 1/25 | 38/12891 | 0.0712099391880838 | 0.163346880913074 | 0.11075401927427 | OG0002466 | 1 |
| GO:0090626 | plant epidermis morphogenesis | 1/25 | 39/12891 | 0.0730165019765608 | 0.166003881967838 | 0.112555544619512 | OG0000169 | 1 |
| GO:0097237 | cellular response to toxic substance | 1/25 | 39/12891 | 0.0730165019765608 | 0.166003881967838 | 0.112555544619512 | OG0002466 | 1 |
| GO:0030154 | cell differentiation | 2/25 | 231/12891 | 0.0732543820783341 | 0.166003881967838 | 0.112555544619512 | OG0000169/OG0000257 | 2 |
| GO:0040007 | growth | 2/25 | 232/12891 | 0.0738054595887652 | 0.166580997144121 | 0.112946845787922 | OG0000169/OG0000257 | 2 |
| GO:0016765 | transferase activity, transferring alkyl or aryl (other than methyl) groups | 1/25 | 40/12891 | 0.0748196911650594 | 0.167524567469177 | 0.113586614391912 | OG0002466 | 1 |
| GO:0051082 | unfolded protein binding | 1/25 | 40/12891 | 0.0748196911650594 | 0.167524567469177 | 0.113586614391912 | OG0000017 | 1 |
| GO:0046395 | carboxylic acid catabolic process | 1/25 | 41/12891 | 0.0766195127914604 | 0.170873675352384 | 0.115857408648741 | OG0002466 | 1 |
| GO:0009888 | tissue development | 2/25 | 240/12891 | 0.0782631826959549 | 0.171477730585283 | 0.11626697597279 | OG0000169/OG0000257 | 2 |
| GO:0016042 | lipid catabolic process | 1/25 | 42/12891 | 0.0784159728833057 | 0.171477730585283 | 0.11626697597279 | OG0000219 | 1 |
| GO:0032269 | negative regulation of cellular protein metabolic process | 1/25 | 42/12891 | 0.0784159728833057 | 0.171477730585283 | 0.11626697597279 | OG0000394 | 1 |
| GO:0044092 | negative regulation of molecular function | 1/25 | 42/12891 | 0.0784159728833057 | 0.171477730585283 | 0.11626697597279 | OG0000394 | 1 |
| GO:0051248 | negative regulation of protein metabolic process | 1/25 | 42/12891 | 0.0784159728833057 | 0.171477730585283 | 0.11626697597279 | OG0000394 | 1 |
| GO:0006650 | glycerophospholipid metabolic process | 1/25 | 44/12891 | 0.0819988325219184 | 0.17861761192759 | 0.121108026817358 | OG0000219 | 1 |
| GO:0009965 | leaf morphogenesis | 1/25 | 45/12891 | 0.0837852440722378 | 0.18180427478223 | 0.123268678537493 | OG0000219 | 1 |
| GO:0016209 | antioxidant activity | 1/25 | 46/12891 | 0.0855683180951397 | 0.183800537248558 | 0.12462220356617 | OG0002466 | 1 |
| GO:0051640 | organelle localization | 1/25 | 46/12891 | 0.0855683180951397 | 0.183800537248558 | 0.12462220356617 | OG0000169 | 1 |
| GO:0009941 | chloroplast envelope | 2/25 | 253/12891 | 0.0856863714575128 | 0.183800537248558 | 0.12462220356617 | OG0000219/OG0000257 | 2 |
| GO:0016054 | organic acid catabolic process | 1/25 | 47/12891 | 0.0873480605667331 | 0.186355658988476 | 0.126354651720975 | OG0002466 | 1 |
| GO:0046872 | metal ion binding | 4/25 | 882/12891 | 0.0875407366066864 | 0.186355658988476 | 0.126354651720975 | OG0000108/OG0000219/OG0000257/OG0003793 | 4 |
| GO:0031967 | organelle envelope | 3/25 | 550/12891 | 0.0887124857283908 | 0.186545104307991 | 0.126483101253966 | OG0000169/OG0000219/OG0000257 | 3 |
| GO:0031975 | envelope | 3/25 | 550/12891 | 0.0887124857283908 | 0.186545104307991 | 0.126483101253966 | OG0000169/OG0000219/OG0000257 | 3 |
| GO:0022626 | cytosolic ribosome | 1/25 | 48/12891 | 0.0891244774528921 | 0.186545104307991 | 0.126483101253966 | OG0000219 | 1 |
| GO:0098754 | detoxification | 1/25 | 48/12891 | 0.0891244774528921 | 0.186545104307991 | 0.126483101253966 | OG0002466 | 1 |
| GO:0031410 | cytoplasmic vesicle | 2/25 | 261/12891 | 0.0903594867742419 | 0.186545104307991 | 0.126483101253966 | OG0000169/OG0000219 | 2 |
| GO:0048869 | cellular developmental process | 2/25 | 261/12891 | 0.0903594867742419 | 0.186545104307991 | 0.126483101253966 | OG0000169/OG0000257 | 2 |
| GO:0000302 | response to reactive oxygen species | 1/25 | 49/12891 | 0.0908975747092716 | 0.186545104307991 | 0.126483101253966 | OG0000017 | 1 |
| GO:0009735 | response to cytokinin | 1/25 | 49/12891 | 0.0908975747092716 | 0.186545104307991 | 0.126483101253966 | OG0000257 | 1 |
| GO:0009860 | pollen tube growth | 1/25 | 49/12891 | 0.0908975747092716 | 0.186545104307991 | 0.126483101253966 | OG0000169 | 1 |
| GO:0097708 | intracellular vesicle | 2/25 | 262/12891 | 0.0909490366199101 | 0.186545104307991 | 0.126483101253966 | OG0000169/OG0000219 | 2 |
| GO:0010033 | response to organic substance | 3/25 | 564/12891 | 0.0940069658784071 | 0.189542195595076 | 0.128515217841202 | OG0000017/OG0000219/OG0000257 | 3 |
| GO:0004601 | peroxidase activity | 1/25 | 51/12891 | 0.0944338341043084 | 0.189542195595076 | 0.128515217841202 | OG0002466 | 1 |
| GO:0005543 | phospholipid binding | 1/25 | 51/12891 | 0.0944338341043084 | 0.189542195595076 | 0.128515217841202 | OG0000219 | 1 |
| GO:0022603 | regulation of anatomical structure morphogenesis | 1/25 | 51/12891 | 0.0944338341043084 | 0.189542195595076 | 0.128515217841202 | OG0000169 | 1 |
| GO:0046486 | glycerolipid metabolic process | 1/25 | 51/12891 | 0.0944338341043084 | 0.189542195595076 | 0.128515217841202 | OG0000219 | 1 |
| GO:0051668 | localization within membrane | 1/25 | 51/12891 | 0.0944338341043084 | 0.189542195595076 | 0.128515217841202 | OG0000257 | 1 |
| GO:0031982 | vesicle | 2/25 | 269/12891 | 0.0951085998211948 | 0.19021719964239 | 0.128972890561051 | OG0000169/OG0000219 | 2 |
| GO:0019752 | carboxylic acid metabolic process | 2/25 | 271/12891 | 0.0963073732582419 | 0.191272254623672 | 0.12968824906119 | OG0000219/OG0002466 | 2 |
| GO:0009791 | post-embryonic development | 3/25 | 570/12891 | 0.0963168114919913 | 0.191272254623672 | 0.12968824906119 | OG0000169/OG0000219/OG0000257 | 3 |
| GO:0009416 | response to light stimulus | 2/25 | 276/12891 | 0.099323879184063 | 0.196549366554378 | 0.133266287118721 | OG0000017/OG0000257 | 2 |
| GO:0043168 | anion binding | 3/25 | 581/12891 | 0.100613842332466 | 0.198403436459108 | 0.13452340138265 | OG0000169/OG0000219/OG0000257 | 3 |
| GO:0043436 | oxoacid metabolic process | 2/25 | 279/12891 | 0.101146958221321 | 0.198691043134141 | 0.134718407219627 | OG0000219/OG0002466 | 2 |
| GO:0009723 | response to ethylene | 1/25 | 55/12891 | 0.101466778255335 | 0.198691043134141 | 0.134718407219627 | OG0000219 | 1 |
| GO:0005635 | nuclear envelope | 1/25 | 56/12891 | 0.10321680400663 | 0.200065745373034 | 0.135650496019925 | OG0000169 | 1 |
| GO:0008047 | enzyme activator activity | 1/25 | 56/12891 | 0.10321680400663 | 0.200065745373034 | 0.135650496019925 | OG0000219 | 1 |
| GO:0010154 | fruit development | 2/25 | 283/12891 | 0.103592761394222 | 0.200065745373034 | 0.135650496019925 | OG0000169/OG0000219 | 2 |
| GO:0051641 | cellular localization | 2/25 | 283/12891 | 0.103592761394222 | 0.200065745373034 | 0.135650496019925 | OG0000169/OG0000257 | 2 |
| GO:0009314 | response to radiation | 2/25 | 285/12891 | 0.104821996631886 | 0.201746445572328 | 0.136790060492944 | OG0000017/OG0000257 | 2 |
| GO:0010118 | stomatal movement | 1/25 | 58/12891 | 0.106707044322295 | 0.203977411255544 | 0.138302721248374 | OG0000219 | 1 |
| GO:0061695 | transferase complex, transferring phosphorus-containing groups | 1/25 | 58/12891 | 0.106707044322295 | 0.203977411255544 | 0.138302721248374 | OG0000017 | 1 |
| GO:0008236 | serine-type peptidase activity | 1/25 | 59/12891 | 0.108447270605467 | 0.205903263784704 | 0.139608506255971 | OG0000650 | 1 |
| GO:0009642 | response to light intensity | 1/25 | 59/12891 | 0.108447270605467 | 0.205903263784704 | 0.139608506255971 | OG0000257 | 1 |
| GO:0006082 | organic acid metabolic process | 2/25 | 293/12891 | 0.109780053130971 | 0.207731952389245 | 0.14084841124725 | OG0000219/OG0002466 | 2 |
| GO:0009570 | chloroplast stroma | 2/25 | 294/12891 | 0.110404338757428 | 0.208212209334479 | 0.141174039668629 | OG0000017/OG0000257 | 2 |
| GO:0048193 | Golgi vesicle transport | 1/25 | 61/12891 | 0.111917964644188 | 0.210360856622186 | 0.142630886115811 | OG0000257 | 1 |
| GO:0009532 | plastid stroma | 2/25 | 298/12891 | 0.112911292958108 | 0.211491475366891 | 0.143397479083751 | OG0000017/OG0000257 | 2 |
| GO:0010035 | response to inorganic substance | 2/25 | 299/12891 | 0.113540455679232 | 0.211491475366891 | 0.143397479083751 | OG0000017/OG0000219 | 2 |
| GO:0006644 | phospholipid metabolic process | 1/25 | 62/12891 | 0.113648444058365 | 0.211491475366891 | 0.143397479083751 | OG0000219 | 1 |
| GO:0048438 | floral whorl development | 1/25 | 63/12891 | 0.115375686158182 | 0.21399714726369 | 0.145096398781524 | OG0000169 | 1 |
| GO:1901605 | alpha-amino acid metabolic process | 1/25 | 65/12891 | 0.118820481622184 | 0.219661548262063 | 0.148937030288194 | OG0002466 | 1 |
| GO:0006996 | organelle organization | 3/25 | 629/12891 | 0.120267591318759 | 0.220659225308327 | 0.149613484850374 | OG0000017/OG0000169/OG0000257 | 3 |
| GO:0012501 | programmed cell death | 1/25 | 66/12891 | 0.120538046565225 | 0.220659225308327 | 0.149613484850374 | OG0000219 | 1 |
| GO:0016853 | isomerase activity | 1/25 | 66/12891 | 0.120538046565225 | 0.220659225308327 | 0.149613484850374 | OG0002466 | 1 |
| GO:0048367 | shoot system development | 2/25 | 314/12891 | 0.123089727020971 | 0.224598787616187 | 0.152284624680764 | OG0000169/OG0000219 | 2 |
| GO:0005507 | copper ion binding | 1/25 | 68/12891 | 0.12396353974608 | 0.224733901088055 | 0.152376235613179 | OG0000257 | 1 |
| GO:0008092 | cytoskeletal protein binding | 1/25 | 68/12891 | 0.12396353974608 | 0.224733901088055 | 0.152376235613179 | OG0000169 | 1 |
| GO:0048868 | pollen tube development | 1/25 | 69/12891 | 0.125671479503263 | 0.227097657494642 | 0.153978932408804 | OG0000169 | 1 |
| GO:0009636 | response to toxic substance | 1/25 | 70/12891 | 0.127376222368059 | 0.229135363694737 | 0.155360557515442 | OG0002466 | 1 |
| GO:0044248 | cellular catabolic process | 2/25 | 321/12891 | 0.127614535296179 | 0.229135363694737 | 0.155360557515442 | OG0000219/OG0002466 | 2 |
| GO:0006457 | protein folding | 1/25 | 71/12891 | 0.129077774075476 | 0.230291139779104 | 0.156144207904168 | OG0000017 | 1 |
| GO:0010016 | shoot system morphogenesis | 1/25 | 71/12891 | 0.129077774075476 | 0.230291139779104 | 0.156144207904168 | OG0000219 | 1 |
| GO:0044282 | small molecule catabolic process | 1/25 | 72/12891 | 0.13077614035068 | 0.232582882522413 | 0.157698077304951 | OG0002466 | 1 |
| GO:1901575 | organic substance catabolic process | 2/25 | 328/12891 | 0.132180352145471 | 0.234338668472413 | 0.158888552139003 | OG0000219/OG0002466 | 2 |
| GO:0008219 | cell death | 1/25 | 74/12891 | 0.134163339455997 | 0.236362999292383 | 0.160261108342092 | OG0000219 | 1 |
| GO:0015630 | microtubule cytoskeleton | 1/25 | 74/12891 | 0.134163339455997 | 0.236362999292383 | 0.160261108342092 | OG0000219 | 1 |
| GO:0019748 | secondary metabolic process | 1/25 | 75/12891 | 0.135852183687377 | 0.238590397600956 | 0.161771350311942 | OG0002466 | 1 |
| GO:0048731 | system development | 3/25 | 666/12891 | 0.136351397122921 | 0.238721137641998 | 0.161859995928832 | OG0000169/OG0000219/OG0000257 | 3 |
| GO:0000160 | phosphorelay signal transduction system | 1/25 | 77/12891 | 0.139220389937395 | 0.242987140201293 | 0.164752471910223 | OG0000219 | 1 |
| GO:0009738 | abscisic acid-activated signaling pathway | 1/25 | 78/12891 | 0.140899763298685 | 0.245156863696164 | 0.166223608649581 | OG0000219 | 1 |
| GO:0019439 | aromatic compound catabolic process | 1/25 | 79/12891 | 0.142575991029703 | 0.247307737526831 | 0.16768196475878 | OG0002466 | 1 |
| GO:0044877 | protein-containing complex binding | 1/25 | 80/12891 | 0.14424907877746 | 0.249439945455177 | 0.169127664833815 | OG0000169 | 1 |
| GO:0009639 | response to red or far red light | 1/25 | 81/12891 | 0.145919032179268 | 0.250784391696478 | 0.170039239172364 | OG0000257 | 1 |
| GO:1901361 | organic cyclic compound catabolic process | 1/25 | 81/12891 | 0.145919032179268 | 0.250784391696478 | 0.170039239172364 | OG0002466 | 1 |
| GO:0045087 | innate immune response | 1/25 | 83/12891 | 0.149249558445898 | 0.255726377581081 | 0.173390051853065 | OG0000219 | 1 |
| GO:0050794 | regulation of cellular process | 4/25 | 1077/12891 | 0.151266104086008 | 0.258393770505582 | 0.17519862319352 | OG0000017/OG0000169/OG0000219/OG0000394 | 4 |
| GO:0009408 | response to heat | 1/25 | 86/12891 | 0.154221980628209 | 0.262644706403193 | 0.178080883532414 | OG0000017 | 1 |
| GO:0071215 | cellular response to abscisic acid stimulus | 1/25 | 88/12891 | 0.157521415810512 | 0.266647697847915 | 0.18079503019469 | OG0000219 | 1 |
| GO:0097306 | cellular response to alcohol | 1/25 | 88/12891 | 0.157521415810512 | 0.266647697847915 | 0.18079503019469 | OG0000219 | 1 |
| GO:0009056 | catabolic process | 2/25 | 370/12891 | 0.160337581548862 | 0.270132687943427 | 0.1831579566127 | OG0000219/OG0002466 | 2 |
| GO:0009611 | response to wounding | 1/25 | 90/12891 | 0.160808492604652 | 0.270132687943427 | 0.1831579566127 | OG0000257 | 1 |
| GO:0009526 | plastid envelope | 2/25 | 371/12891 | 0.161022153845281 | 0.270132687943427 | 0.1831579566127 | OG0000219/OG0000257 | 2 |
| GO:0043169 | cation binding | 3/25 | 732/12891 | 0.166818313229746 | 0.278249439560028 | 0.188661354412306 | OG0000219/OG0000257/OG0003793 | 3 |
| GO:0009856 | pollination | 1/25 | 94/12891 | 0.16734574834749 | 0.278249439560028 | 0.188661354412306 | OG0000169 | 1 |
| GO:0044706 | multi-multicellular organism process | 1/25 | 94/12891 | 0.16734574834749 | 0.278249439560028 | 0.188661354412306 | OG0000169 | 1 |
| GO:0048584 | positive regulation of response to stimulus | 1/25 | 95/12891 | 0.168972407430972 | 0.280125348012408 | 0.189933275857823 | OG0000219 | 1 |
| GO:0006631 | fatty acid metabolic process | 1/25 | 96/12891 | 0.170596015575253 | 0.281158242678276 | 0.190633609008309 | OG0000219 | 1 |
| GO:0008289 | lipid binding | 1/25 | 96/12891 | 0.170596015575253 | 0.281158242678276 | 0.190633609008309 | OG0000219 | 1 |
| GO:0019899 | enzyme binding | 1/25 | 98/12891 | 0.17383410097398 | 0.285657206863675 | 0.193684039866362 | OG0000169 | 1 |
| GO:0023051 | regulation of signaling | 1/25 | 101/12891 | 0.178668483829369 | 0.292745445807888 | 0.19849007563674 | OG0000219 | 1 |
| GO:0009505 | plant-type cell wall | 1/25 | 102/12891 | 0.180273901179195 | 0.293663572355674 | 0.199112592606769 | OG0000257 | 1 |
| GO:1901615 | organic hydroxy compound metabolic process | 1/25 | 102/12891 | 0.180273901179195 | 0.293663572355674 | 0.199112592606769 | OG0002466 | 1 |
| GO:0043167 | ion binding | 4/25 | 1156/12891 | 0.181327360870168 | 0.294525944534782 | 0.199697306464864 | OG0000169/OG0000219/OG0000257/OG0003793 | 4 |
| GO:0042578 | phosphoric ester hydrolase activity | 1/25 | 103/12891 | 0.18187630578241 | 0.294566235878139 | 0.199724625188025 | OG0000219 | 1 |
| GO:0009966 | regulation of signal transduction | 1/25 | 104/12891 | 0.183475703057624 | 0.296302715857428 | 0.200902010002601 | OG0000219 | 1 |
| GO:0031090 | organelle membrane | 3/25 | 768/12891 | 0.184273705588918 | 0.296326265836859 | 0.200917977585583 | OG0000169/OG0000219/OG0003793 | 3 |
| GO:0010646 | regulation of cell communication | 1/25 | 105/12891 | 0.185072098414124 | 0.296326265836859 | 0.200917977585583 | OG0000219 | 1 |
| GO:0048046 | apoplast | 1/25 | 105/12891 | 0.185072098414124 | 0.296326265836859 | 0.200917977585583 | OG0000394 | 1 |
| GO:0005198 | structural molecule activity | 1/25 | 108/12891 | 0.189843326924678 | 0.30138968850754 | 0.204351128000991 | OG0000257 | 1 |
| GO:0006790 | sulfur compound metabolic process | 1/25 | 108/12891 | 0.189843326924678 | 0.30138968850754 | 0.204351128000991 | OG0002466 | 1 |
| GO:0009658 | chloroplast organization | 1/25 | 108/12891 | 0.189843326924678 | 0.30138968850754 | 0.204351128000991 | OG0000017 | 1 |
| GO:0016787 | hydrolase activity | 3/25 | 787/12891 | 0.193694020140799 | 0.30663673047642 | 0.207908777734527 | OG0000169/OG0000219/OG0000394 | 3 |
| GO:0004553 | hydrolase activity, hydrolyzing O-glycosyl compounds | 1/25 | 111/12891 | 0.194587732009338 | 0.307186251093393 | 0.208281368975105 | OG0000162 | 1 |
| GO:0005488 | binding | 8/25 | 2958/12891 | 0.19683748973953 | 0.309867420822453 | 0.210099228169756 | OG0000017/OG0000169/OG0000219/OG0000257/OG0000394/OG0000905/OG0002466/OG0003793 | 8 |
| GO:0006520 | cellular amino acid metabolic process | 1/25 | 116/12891 | 0.202435858470167 | 0.317790370000654 | 0.215471275407823 | OG0002466 | 1 |
| GO:0031974 | membrane-enclosed lumen | 2/25 | 433/12891 | 0.204442835097095 | 0.318273887325671 | 0.215799114463182 | OG0000017/OG0000257 | 2 |
| GO:0043233 | organelle lumen | 2/25 | 433/12891 | 0.204442835097095 | 0.318273887325671 | 0.215799114463182 | OG0000017/OG0000257 | 2 |
| GO:0070013 | intracellular organelle lumen | 2/25 | 433/12891 | 0.204442835097095 | 0.318273887325671 | 0.215799114463182 | OG0000017/OG0000257 | 2 |
| GO:0003924 | GTPase activity | 1/25 | 125/12891 | 0.216377851010007 | 0.335923625048686 | 0.227766158957903 | OG0000345 | 1 |
| GO:0032559 | adenyl ribonucleotide binding | 2/25 | 452/12891 | 0.218047887130608 | 0.337583781177415 | 0.228891793943106 | OG0000169/OG0000257 | 2 |
| GO:0009719 | response to endogenous stimulus | 2/25 | 454/12891 | 0.219485921340745 | 0.337948185735613 | 0.229138871017591 | OG0000219/OG0000257 | 2 |
| GO:0030554 | adenyl nucleotide binding | 2/25 | 454/12891 | 0.219485921340745 | 0.337948185735613 | 0.229138871017591 | OG0000169/OG0000257 | 2 |
| GO:0009725 | response to hormone | 2/25 | 457/12891 | 0.221644873799863 | 0.339583934140757 | 0.230247956843892 | OG0000219/OG0000257 | 2 |
| GO:0050789 | regulation of biological process | 4/25 | 1255/12891 | 0.221756768380174 | 0.339583934140757 | 0.230247956843892 | OG0000017/OG0000169/OG0000219/OG0000394 | 4 |
| GO:0044281 | small molecule metabolic process | 2/25 | 459/12891 | 0.223085402070377 | 0.340690206422695 | 0.230998042189578 | OG0000219/OG0002466 | 2 |
| GO:0048608 | reproductive structure development | 2/25 | 462/12891 | 0.225247965059574 | 0.341656926276742 | 0.231653506859301 | OG0000169/OG0000219 | 2 |
| GO:0061458 | reproductive system development | 2/25 | 462/12891 | 0.225247965059574 | 0.341656926276742 | 0.231653506859301 | OG0000169/OG0000219 | 2 |
| GO:0032268 | regulation of cellular protein metabolic process | 1/25 | 131/12891 | 0.225542205780554 | 0.341656926276742 | 0.231653506859301 | OG0000394 | 1 |
| GO:0140657 | ATP-dependent activity | 1/25 | 133/12891 | 0.228574060823351 | 0.345318876835278 | 0.23413641770813 | OG0000169 | 1 |
| GO:0008233 | peptidase activity | 1/25 | 134/12891 | 0.230085712216941 | 0.346670697763863 | 0.2350529923029 | OG0000394 | 1 |
| GO:0009657 | plastid organization | 1/25 | 135/12891 | 0.231594519710441 | 0.348011016249379 | 0.235961767900871 | OG0000017 | 1 |
| GO:0009733 | response to auxin | 1/25 | 137/12891 | 0.234603623499365 | 0.351592630417715 | 0.238390208299705 | OG0000257 | 1 |
| GO:0007275 | multicellular organism development | 3/25 | 869/12891 | 0.23572697330996 | 0.351963948735729 | 0.238641973107949 | OG0000169/OG0000219/OG0000257 | 3 |
| GO:0035556 | intracellular signal transduction | 1/25 | 138/12891 | 0.23610393002379 | 0.351963948735729 | 0.238641973107949 | OG0000219 | 1 |
| GO:0009507 | chloroplast | 5/25 | 1731/12891 | 0.238485894309211 | 0.354543524394403 | 0.240391001743349 | OG0000017/OG0000169/OG0000219/OG0000257/OG0003793 | 5 |
| GO:0051246 | regulation of protein metabolic process | 1/25 | 140/12891 | 0.23909607783893 | 0.354543524394403 | 0.240391001743349 | OG0000394 | 1 |
| GO:0042802 | identical protein binding | 1/25 | 141/12891 | 0.240587929305797 | 0.355816884920679 | 0.241254377863431 | OG0000169 | 1 |
| GO:0009409 | response to cold | 1/25 | 142/12891 | 0.242076972581669 | 0.357079418873748 | 0.242110413246482 | OG0000219 | 1 |
| GO:0042742 | defense response to bacterium | 1/25 | 144/12891 | 0.245046654816293 | 0.360264411372245 | 0.24426993241572 | OG0000219 | 1 |
| GO:0035639 | purine ribonucleoside triphosphate binding | 2/25 | 490/12891 | 0.245518273230552 | 0.360264411372245 | 0.24426993241572 | OG0000169/OG0000257 | 2 |
| GO:0031966 | mitochondrial membrane | 1/25 | 145/12891 | 0.246527303881139 | 0.360802981201042 | 0.244635098697841 | OG0000219 | 1 |
| GO:0016043 | cellular component organization | 3/25 | 892/12891 | 0.247848581805113 | 0.36108524018541 | 0.244826478642289 | OG0000017/OG0000169/OG0000257 | 3 |
| GO:0051172 | negative regulation of nitrogen compound metabolic process | 1/25 | 146/12891 | 0.248005164967203 | 0.36108524018541 | 0.244826478642289 | OG0000394 | 1 |
| GO:0032555 | purine ribonucleotide binding | 2/25 | 500/12891 | 0.252787055690408 | 0.367096447798473 | 0.248902255296961 | OG0000169/OG0000257 | 2 |
| GO:0017076 | purine nucleotide binding | 2/25 | 503/12891 | 0.25496987267851 | 0.367969970504215 | 0.249494529542097 | OG0000169/OG0000257 | 2 |
| GO:0032553 | ribonucleotide binding | 2/25 | 503/12891 | 0.25496987267851 | 0.367969970504215 | 0.249494529542097 | OG0000169/OG0000257 | 2 |
| GO:0009414 | response to water deprivation | 1/25 | 151/12891 | 0.255352826506483 | 0.367969970504215 | 0.249494529542097 | OG0000219 | 1 |
| GO:0005740 | mitochondrial envelope | 1/25 | 152/12891 | 0.256814065072994 | 0.368187511660771 | 0.249642028884059 | OG0000219 | 1 |
| GO:0006974 | cellular response to DNA damage stimulus | 1/25 | 152/12891 | 0.256814065072994 | 0.368187511660771 | 0.249642028884059 | OG0000394 | 1 |
| GO:1901700 | response to oxygen-containing compound | 2/25 | 509/12891 | 0.259338014434719 | 0.370475375770245 | 0.251193268456319 | OG0000017/OG0000219 | 2 |
| GO:0009415 | response to water | 1/25 | 154/12891 | 0.259728288351382 | 0.370475375770245 | 0.251193268456319 | OG0000219 | 1 |
| GO:0005886 | plasma membrane | 3/25 | 919/12891 | 0.262223547092571 | 0.372185509427738 | 0.252352789685037 | OG0000219/OG0000257/OG0003793 | 3 |
| GO:0097367 | carbohydrate derivative binding | 2/25 | 513/12891 | 0.262251711269367 | 0.372185509427738 | 0.252352789685037 | OG0000169/OG0000257 | 2 |
| GO:0016491 | oxidoreductase activity | 2/25 | 514/12891 | 0.26298030909376 | 0.372279430001746 | 0.252416470613658 | OG0000555/OG0002466 | 2 |
| GO:0001101 | response to acid chemical | 1/25 | 159/12891 | 0.266965930431029 | 0.376027200256236 | 0.254957569756055 | OG0000219 | 1 |
| GO:0005618 | cell wall | 1/25 | 159/12891 | 0.266965930431029 | 0.376027200256236 | 0.254957569756055 | OG0000257 | 1 |
| GO:0005774 | vacuolar membrane | 1/25 | 161/12891 | 0.26984192448345 | 0.378182447779797 | 0.256418891358469 | OG0003793 | 1 |
| GO:0030312 | external encapsulating structure | 1/25 | 161/12891 | 0.26984192448345 | 0.378182447779797 | 0.256418891358469 | OG0000257 | 1 |
| GO:0071396 | cellular response to lipid | 1/25 | 162/12891 | 0.271275856289256 | 0.379246346354631 | 0.257140246076749 | OG0000219 | 1 |
| GO:0032501 | multicellular organismal process | 3/25 | 940/12891 | 0.273496913192099 | 0.381402643210818 | 0.258602279157737 | OG0000169/OG0000219/OG0000257 | 3 |
| GO:0005515 | protein binding | 8/25 | 3228/12891 | 0.274959248223807 | 0.382492815598464 | 0.25934144829864 | OG0000017/OG0000153/OG0000169/OG0000219/OG0000257/OG0000394/OG0000905/OG0002466 | 8 |
| GO:0006810 | transport | 2/25 | 532/12891 | 0.276103225083032 | 0.383135833325097 | 0.259777433346479 | OG0000169/OG0000257 | 2 |
| GO:0065007 | biological regulation | 4/25 | 1382/12891 | 0.277074218499976 | 0.383536233490115 | 0.260048916507626 | OG0000017/OG0000169/OG0000219/OG0000394 | 4 |
| GO:0051704 | | 1/25 | 167/12891 | 0.278405058621047 | 0.38443155514749 | 0.260655971087079 | OG0000169 | 1 |
| GO:0009617 | response to bacterium | 1/25 | 169/12891 | 0.281237950604507 | 0.387391490783659 | 0.262662895043425 | OG0000219 | 1 |
| GO:0003006 | developmental process involved in reproduction | 2/25 | 540/12891 | 0.281938007106312 | 0.387406259153416 | 0.262672908435169 | OG0000169/OG0000219 | 2 |
| GO:0031324 | negative regulation of cellular metabolic process | 1/25 | 170/12891 | 0.282650389783479 | 0.387437851361744 | 0.262694328887341 | OG0000394 | 1 |
| GO:0048856 | anatomical structure development | 3/25 | 960/12891 | 0.284297100138729 | 0.388564755631221 | 0.263458403331152 | OG0000169/OG0000219/OG0000257 | 3 |
| GO:0051234 | establishment of localization | 2/25 | 544/12891 | 0.284855301281251 | 0.388564755631221 | 0.263458403331152 | OG0000169/OG0000257 | 2 |
| GO:0019637 | organophosphate metabolic process | 1/25 | 174/12891 | 0.288273546940562 | 0.392275383488126 | 0.26597431882881 | OG0000219 | 1 |
| GO:0009908 | flower development | 1/25 | 175/12891 | 0.289672710369554 | 0.393227205863984 | 0.266619682567451 | OG0000169 | 1 |
| GO:0009536 | plastid | 5/25 | 1869/12891 | 0.291874889543928 | 0.395261898611295 | 0.267999264463924 | OG0000017/OG0000169/OG0000219/OG0000257/OG0003793 | 5 |
| GO:0051171 | regulation of nitrogen compound metabolic process | 2/25 | 556/12891 | 0.293604808766931 | 0.396648804151479 | 0.268939627463636 | OG0000017/OG0000394 | 2 |
| GO:1901265 | nucleoside phosphate binding | 2/25 | 561/12891 | 0.297248635300705 | 0.399720691746782 | 0.271022458161332 | OG0000169/OG0000257 | 2 |
| GO:0005525 | GTP binding | 1/25 | 181/12891 | 0.298012401853917 | 0.399720691746782 | 0.271022458161332 | OG0000345 | 1 |
| GO:0090567 | reproductive shoot system development | 1/25 | 181/12891 | 0.298012401853917 | 0.399720691746782 | 0.271022458161332 | OG0000169 | 1 |
| GO:0005730 | nucleolus | 1/25 | 182/12891 | 0.299393179977729 | 0.40061658844639 | 0.271629902636436 | OG0000257 | 1 |
| GO:0032502 | developmental process | 3/25 | 999/12891 | 0.305498079301741 | 0.407814538165269 | 0.27651032555877 | OG0000169/OG0000219/OG0000257 | 3 |
| GO:0071840 | cellular component organization or biogenesis | 3/25 | 1001/12891 | 0.306589314313652 | 0.408301409109651 | 0.276840438467304 | OG0000017/OG0000169/OG0000257 | 3 |
| GO:0016192 | vesicle-mediated transport | 1/25 | 190/12891 | 0.31034593047918 | 0.412327217326949 | 0.279570055576616 | OG0000257 | 1 |
| GO:0071944 | cell periphery | 3/25 | 1011/12891 | 0.312050149291566 | 0.413613641277972 | 0.280442289085271 | OG0000219/OG0000257/OG0003793 | 3 |
| GO:1901565 | organonitrogen compound catabolic process | 1/25 | 194/12891 | 0.315760476609718 | 0.417546794952145 | 0.283109083672367 | OG0002466 | 1 |
| GO:0051179 | localization | 2/25 | 593/12891 | 0.320520791279904 | 0.422846677697902 | 0.286702561016371 | OG0000169/OG0000257 | 2 |
| GO:0080090 | regulation of primary metabolic process | 2/25 | 595/12891 | 0.321971511648665 | 0.423765783481381 | 0.287325741937179 | OG0000017/OG0000394 | 2 |
| GO:0009737 | response to abscisic acid | 1/25 | 201/12891 | 0.325137880550349 | 0.426449716763147 | 0.28914430767608 | OG0000219 | 1 |
| GO:0000166 | nucleotide binding | 2/25 | 600/12891 | 0.325595861588739 | 0.426449716763147 | 0.28914430767608 | OG0000169/OG0000257 | 2 |
| GO:0065003 | protein-containing complex assembly | 1/25 | 202/12891 | 0.326467396152102 | 0.426449716763147 | 0.28914430767608 | OG0000017 | 1 |
| GO:0036094 | small molecule binding | 2/25 | 602/12891 | 0.327044576734727 | 0.426449716763147 | 0.28914430767608 | OG0000169/OG0000257 | 2 |
| GO:0006259 | DNA metabolic process | 1/25 | 205/12891 | 0.330440873292212 | 0.429879099051442 | 0.291470750808432 | OG0000394 | 1 |
| GO:1901701 | cellular response to oxygen-containing compound | 1/25 | 208/12891 | 0.334391841412176 | 0.433713939442982 | 0.294070886080463 | OG0000219 | 1 |
| GO:0006518 | peptide metabolic process | 1/25 | 209/12891 | 0.335703849924728 | 0.433713939442982 | 0.294070886080463 | OG0002466 | 1 |
| GO:0009793 | embryo development ending in seed dormancy | 1/25 | 209/12891 | 0.335703849924728 | 0.433713939442982 | 0.294070886080463 | OG0000219 | 1 |
| GO:0022414 | reproductive process | 2/25 | 616/12891 | 0.337167562473574 | 0.434539608916441 | 0.294630714417965 | OG0000169/OG0000219 | 2 |
| GO:0000003 | reproduction | 2/25 | 617/12891 | 0.337889340029332 | 0.434539608916441 | 0.294630714417965 | OG0000169/OG0000219 | 2 |
| GO:0044255 | cellular lipid metabolic process | 1/25 | 214/12891 | 0.342226738858454 | 0.43811259052039 | 0.297053301682677 | OG0000219 | 1 |
| GO:0046907 | intracellular transport | 1/25 | 214/12891 | 0.342226738858454 | 0.43811259052039 | 0.297053301682677 | OG0000169 | 1 |
| GO:1990234 | transferase complex | 1/25 | 215/12891 | 0.343523917333531 | 0.43877373077601 | 0.297501574341479 | OG0000017 | 1 |
| GO:0009790 | embryo development | 1/25 | 217/12891 | 0.346110910733848 | 0.441075582386446 | 0.299062297853331 | OG0000219 | 1 |
| GO:0045184 | establishment of protein localization | 1/25 | 219/12891 | 0.348688115740059 | 0.443354572502066 | 0.300607520595145 | OG0000257 | 1 |
| GO:0006807 | nitrogen compound metabolic process | 5/25 | 2014/12891 | 0.350608389428834 | 0.444789875528227 | 0.301580698522604 | OG0000017/OG0000219/OG0000306/OG0000394/OG0002466 | 5 |
| GO:0010605 | negative regulation of macromolecule metabolic process | 1/25 | 223/12891 | 0.353813302441865 | 0.447844765703442 | 0.303652004466466 | OG0000394 | 1 |
| GO:0060255 | regulation of macromolecule metabolic process | 2/25 | 642/12891 | 0.35586932246679 | 0.449047267664548 | 0.304467336382406 | OG0000017/OG0000394 | 2 |
| GO:0051649 | establishment of localization in cell | 1/25 | 225/12891 | 0.356361354765816 | 0.449047267664548 | 0.304467336382406 | OG0000169 | 1 |
| GO:0043933 | protein-containing complex organization | 1/25 | 226/12891 | 0.357631761060689 | 0.449639932250799 | 0.304869180510937 | OG0000017 | 1 |
| GO:0031323 | regulation of cellular metabolic process | 2/25 | 646/12891 | 0.358733535831789 | 0.450018408788985 | 0.305125798804294 | OG0000017/OG0000394 | 2 |
| GO:0140535 | intracellular protein-containing complex | 1/25 | 228/12891 | 0.360165355815565 | 0.450808307279171 | 0.305661373356546 | OG0000017 | 1 |
| GO:0009755 | hormone-mediated signaling pathway | 1/25 | 229/12891 | 0.361428553012486 | 0.450954464638001 | 0.305760472371149 | OG0000219 | 1 |
| GO:0015031 | protein transport | 1/25 | 230/12891 | 0.362689355901026 | 0.450954464638001 | 0.305760472371149 | OG0000257 | 1 |
| GO:0046914 | transition metal ion binding | 1/25 | 230/12891 | 0.362689355901026 | 0.450954464638001 | 0.305760472371149 | OG0000257 | 1 |
| GO:0009892 | negative regulation of metabolic process | 1/25 | 239/12891 | 0.373929553200688 | 0.463903772403503 | 0.314540486252235 | OG0000394 | 1 |
| GO:0043603 | cellular amide metabolic process | 1/25 | 240/12891 | 0.375166650985227 | 0.464413343290083 | 0.314885990393351 | OG0002466 | 1 |
| GO:0032870 | cellular response to hormone stimulus | 1/25 | 243/12891 | 0.378863880134822 | 0.46693311542932 | 0.316594470472774 | OG0000219 | 1 |
| GO:0071495 | cellular response to endogenous stimulus | 1/25 | 243/12891 | 0.378863880134822 | 0.46693311542932 | 0.316594470472774 | OG0000219 | 1 |
| GO:0048523 | negative regulation of cellular process | 1/25 | 249/12891 | 0.386195432050651 | 0.4749274240973 | 0.322014848329692 | OG0000394 | 1 |
| GO:0008104 | protein localization | 1/25 | 250/12891 | 0.387409252189769 | 0.475379912075656 | 0.322321648597146 | OG0000257 | 1 |
| GO:0005840 | ribosome | 1/25 | 254/12891 | 0.392241529249006 | 0.478604879060787 | 0.324508271623909 | OG0000219 | 1 |
| GO:0048583 | regulation of response to stimulus | 1/25 | 254/12891 | 0.392241529249006 | 0.478604879060787 | 0.324508271623909 | OG0000219 | 1 |
| GO:0016788 | hydrolase activity, acting on ester bonds | 1/25 | 255/12891 | 0.39344386855175 | 0.478604879060787 | 0.324508271623909 | OG0000219 | 1 |
| GO:0065008 | regulation of biological quality | 1/25 | 255/12891 | 0.39344386855175 | 0.478604879060787 | 0.324508271623909 | OG0000169 | 1 |
| GO:0003729 | mRNA binding | 1/25 | 257/12891 | 0.395841700378051 | 0.480481718385453 | 0.325780824228384 | OG0000257 | 1 |
| GO:0006725 | cellular aromatic compound metabolic process | 3/25 | 1174/12891 | 0.401262505307815 | 0.485612154933669 | 0.32925941203594 | OG0000017/OG0000394/OG0002466 | 3 |
| GO:0050793 | regulation of developmental process | 1/25 | 262/12891 | 0.401796533886399 | 0.485612154933669 | 0.32925941203594 | OG0000169 | 1 |
| GO:0048316 | seed development | 1/25 | 274/12891 | 0.415859126785594 | 0.500511830848296 | 0.339361833240463 | OG0000219 | 1 |
| GO:1901360 | organic cyclic compound metabolic process | 3/25 | 1201/12891 | 0.415905738445114 | 0.500511830848296 | 0.339361833240463 | OG0000017/OG0000394/OG0002466 | 3 |
| GO:0071310 | cellular response to organic substance | 1/25 | 279/12891 | 0.421624388390839 | 0.506309628794127 | 0.3432929118252 | OG0000219 | 1 |
| GO:0019222 | regulation of metabolic process | 2/25 | 737/12891 | 0.422670954102463 | 0.506484171014039 | 0.343411256615625 | OG0000017/OG0000394 | 2 |
| GO:1901564 | organonitrogen compound metabolic process | 3/25 | 1216/12891 | 0.424004937967774 | 0.507001649229551 | 0.343762122159763 | OG0000219/OG0000394/OG0002466 | 3 |
| GO:0009987 | cellular process | 7/25 | 3205/12891 | 0.432519080899625 | 0.51608433856813 | 0.349920454320403 | OG0000017/OG0000169/OG0000219/OG0000257/OG0000394/OG0002466/OG0003793 | 7 |
| GO:0005634 | nucleus | 6/25 | 2719/12891 | 0.43705057771918 | 0.520386492962244 | 0.352837442315663 | OG0000169/OG0000219/OG0000257/OG0000394/OG0000905/OG0003793 | 6 |
| GO:0048522 | positive regulation of cellular process | 1/25 | 295/12891 | 0.4397084410451 | 0.522444278789316 | 0.354232682003619 | OG0000219 | 1 |
| GO:0098542 | defense response to other organism | 1/25 | 296/12891 | 0.440820483675608 | 0.522660573471923 | 0.354379336199356 | OG0000219 | 1 |
| GO:0071705 | nitrogen compound transport | 1/25 | 298/12891 | 0.443038215754556 | 0.524184162640127 | 0.355412374743821 | OG0000257 | 1 |
| GO:0002376 | immune system process | 1/25 | 301/12891 | 0.446348988864824 | 0.524786886489605 | 0.355821039350509 | OG0000219 | 1 |
| GO:0006955 | immune response | 1/25 | 301/12891 | 0.446348988864824 | 0.524786886489605 | 0.355821039350509 | OG0000219 | 1 |
| GO:0022607 | cellular component assembly | 1/25 | 301/12891 | 0.446348988864824 | 0.524786886489605 | 0.355821039350509 | OG0000017 | 1 |
| GO:0003723 | RNA binding | 2/25 | 784/12891 | 0.454563079943297 | 0.533328707574391 | 0.361612646828862 | OG0000257/OG0000711 | 2 |
| GO:0016020 | membrane | 6/25 | 2782/12891 | 0.461098303683383 | 0.539869263895961 | 0.366047337573212 | OG0000132/OG0000169/OG0000219/OG0000257/OG0000564/OG0003793 | 6 |
| GO:0034641 | cellular nitrogen compound metabolic process | 3/25 | 1288/12891 | 0.462431033912212 | 0.540304035464996 | 0.366342125563455 | OG0000017/OG0000394/OG0002466 | 3 |
| GO:0005975 | carbohydrate metabolic process | 1/25 | 317/12891 | 0.463689796387648 | 0.540650758443689 | 0.366577214003775 | OG0000162 | 1 |
| GO:0033036 | macromolecule localization | 1/25 | 319/12891 | 0.465820376692757 | 0.542010459008964 | 0.367499131225407 | OG0000257 | 1 |
| GO:0033993 | response to lipid | 1/25 | 320/12891 | 0.46688261743271 | 0.542124030985915 | 0.367576136386779 | OG0000219 | 1 |
| GO:0005737 | cytoplasm | 8/25 | 3820/12891 | 0.471994088626388 | 0.546929232593876 | 0.370834205279984 | OG0000017/OG0000169/OG0000219/OG0000257/OG0000394/OG0000905/OG0002466/OG0003793 | 8 |
| GO:0005739 | mitochondrion | 3/25 | 1312/12891 | 0.475043525069058 | 0.549330166849405 | 0.372462109757416 | OG0000219/OG0000257/OG0000905 | 3 |
| GO:0071702 | organic substance transport | 1/25 | 334/12891 | 0.481542729087639 | 0.555702286955346 | 0.376782595762943 | OG0000257 | 1 |
| GO:0098588 | bounding membrane of organelle | 1/25 | 339/12891 | 0.486684073577307 | 0.560484527357473 | 0.380025096279088 | OG0003793 | 1 |
| GO:0031981 | nuclear lumen | 1/25 | 350/12891 | 0.497823304512456 | 0.572140484940696 | 0.387928180461756 | OG0000257 | 1 |
| GO:0006629 | lipid metabolic process | 1/25 | 352/12891 | 0.499823528352763 | 0.573266985580108 | 0.388691981232439 | OG0000219 | 1 |
| GO:0006952 | defense response | 1/25 | 361/12891 | 0.508730275269756 | 0.579873036024237 | 0.393171078930088 | OG0000219 | 1 |
| GO:0043207 | response to external biotic stimulus | 1/25 | 361/12891 | 0.508730275269756 | 0.579873036024237 | 0.393171078930088 | OG0000219 | 1 |
| GO:0051707 | response to other organism | 1/25 | 361/12891 | 0.508730275269756 | 0.579873036024237 | 0.393171078930088 | OG0000219 | 1 |
| GO:0009607 | response to biotic stimulus | 1/25 | 362/12891 | 0.509710462270414 | 0.579873036024237 | 0.393171078930088 | OG0000219 | 1 |
| GO:1902494 | catalytic complex | 1/25 | 364/12891 | 0.511665206909793 | 0.580920901582432 | 0.393881562788613 | OG0000017 | 1 |
| GO:0044419 | biological process involved in interspecies interaction between organisms | 1/25 | 366/12891 | 0.513612468683449 | 0.58195606330665 | 0.39458343307175 | OG0000219 | 1 |
| GO:0048519 | negative regulation of biological process | 1/25 | 372/12891 | 0.519409630885052 | 0.587340467922332 | 0.398234218744866 | OG0000394 | 1 |
| GO:0032991 | protein-containing complex | 2/25 | 894/12891 | 0.525452845382983 | 0.592980921898065 | 0.402058613461509 | OG0000017/OG0000169 | 2 |
| GO:1903506 | regulation of nucleic acid-templated transcription | 1/25 | 384/12891 | 0.530805594216722 | 0.596625487899596 | 0.404529737066218 | OG0000017 | 1 |
| GO:2001141 | regulation of RNA biosynthetic process | 1/25 | 384/12891 | 0.530805594216722 | 0.596625487899596 | 0.404529737066218 | OG0000017 | 1 |
| GO:0005794 | Golgi apparatus | 1/25 | 391/12891 | 0.537332958613331 | 0.6015560213958 | 0.407872784688668 | OG0000169 | 1 |
| GO:0048518 | positive regulation of biological process | 1/25 | 391/12891 | 0.537332958613331 | 0.6015560213958 | 0.407872784688668 | OG0000219 | 1 |
| GO:0003824 | catalytic activity | 4/25 | 1967/12891 | 0.543486324153191 | 0.607235217045912 | 0.411723447407043 | OG0000169/OG0000219/OG0000394/OG0002466 | 4 |
| GO:0051252 | regulation of RNA metabolic process | 1/25 | 413/12891 | 0.557284865073339 | 0.621173432324322 | 0.421173969847171 | OG0000017 | 1 |
| GO:0023052 | signaling | 1/25 | 414/12891 | 0.558171856448012 | 0.621173432324322 | 0.421173969847171 | OG0000219 | 1 |
| GO:0097659 | nucleic acid-templated transcription | 1/25 | 423/12891 | 0.566078377777942 | 0.628727368203959 | 0.426295761921396 | OG0000017 | 1 |
| GO:0090304 | nucleic acid metabolic process | 2/25 | 970/12891 | 0.570994055014461 | 0.63293621088388 | 0.429149481813008 | OG0000017/OG0000394 | 2 |
| GO:0019219 | regulation of nucleobase-containing compound metabolic process | 1/25 | 434/12891 | 0.57555754010215 | 0.636738853420095 | 0.431727785986279 | OG0000017 | 1 |
| GO:0032774 | RNA biosynthetic process | 1/25 | 437/12891 | 0.578108064837788 | 0.638303993003609 | 0.432788996941949 | OG0000017 | 1 |
| GO:0044085 | cellular component biogenesis | 1/25 | 449/12891 | 0.588163806762148 | 0.648133449804563 | 0.439453659541584 | OG0000017 | 1 |
| GO:0007165 | signal transduction | 1/25 | 451/12891 | 0.589817234488921 | 0.648683533821475 | 0.439826632783993 | OG0000219 | 1 |
| GO:0006351 | transcription, DNA-templated | 1/25 | 468/12891 | 0.603616398200046 | 0.66256331208677 | 0.44923753319987 | OG0000017 | 1 |
| GO:2000112 | regulation of cellular macromolecule biosynthetic process | 1/25 | 470/12891 | 0.605210218871929 | 0.663017822623829 | 0.44954570479458 | OG0000017 | 1 |
| GO:0010556 | regulation of macromolecule biosynthetic process | 1/25 | 472/12891 | 0.60679788629696 | 0.663463836768271 | 0.449848115583656 | OG0000017 | 1 |
| GO:0009605 | response to external stimulus | 1/25 | 481/12891 | 0.613866851902198 | 0.669438627884255 | 0.453899200775614 | OG0000219 | 1 |
| GO:0007154 | cell communication | 1/25 | 482/12891 | 0.614644718840348 | 0.669438627884255 | 0.453899200775614 | OG0000219 | 1 |
| GO:0006796 | phosphate-containing compound metabolic process | 1/25 | 499/12891 | 0.627640535413019 | 0.682270756096937 | 0.462599763452128 | OG0000219 | 1 |
| GO:0006793 | phosphorus metabolic process | 1/25 | 505/12891 | 0.632126020484703 | 0.684269265547232 | 0.463954812002431 | OG0000219 | 1 |
| GO:0006139 | nucleobase-containing compound metabolic process | 2/25 | 1084/12891 | 0.633647962689854 | 0.684269265547232 | 0.463954812002431 | OG0000017/OG0000394 | 2 |
| GO:0031326 | regulation of cellular biosynthetic process | 1/25 | 508/12891 | 0.634349265747523 | 0.684269265547232 | 0.463954812002431 | OG0000017 | 1 |
| GO:0034654 | nucleobase-containing compound biosynthetic process | 1/25 | 508/12891 | 0.634349265747523 | 0.684269265547232 | 0.463954812002431 | OG0000017 | 1 |
| GO:0044238 | primary metabolic process | 4/25 | 2198/12891 | 0.637921889573133 | 0.686804792988699 | 0.465673974643021 | OG0000017/OG0000219/OG0000394/OG0002466 | 4 |
| GO:0140096 | catalytic activity, acting on a protein | 1/25 | 517/12891 | 0.64094187713563 | 0.688736778107503 | 0.466983917727882 | OG0000394 | 1 |
| GO:0009889 | regulation of biosynthetic process | 1/25 | 520/12891 | 0.643113944090998 | 0.689751978204467 | 0.467672253437005 | OG0000017 | 1 |
| GO:0010468 | regulation of gene expression | 1/25 | 533/12891 | 0.65238133634908 | 0.698358687672729 | 0.473507857159633 | OG0000017 | 1 |
| GO:0046483 | heterocycle metabolic process | 2/25 | 1154/12891 | 0.66869401804926 | 0.714460148562137 | 0.48442512414215 | OG0000017/OG0000394 | 2 |
| GO:0006355 | regulation of transcription, DNA-templated | 1/25 | 576/12891 | 0.681417543172566 | 0.72667212655923 | 0.492705206620044 | OG0000017 | 1 |
| GO:0018130 | heterocycle biosynthetic process | 1/25 | 578/12891 | 0.682709755913298 | 0.72667212655923 | 0.492705206620044 | OG0000017 | 1 |
| GO:0019438 | aromatic compound biosynthetic process | 1/25 | 581/12891 | 0.684638644870634 | 0.727347671866344 | 0.493163246330056 | OG0000017 | 1 |
| GO:0044237 | cellular metabolic process | 4/25 | 2364/12891 | 0.698394128864297 | 0.740561321550444 | 0.502122491854768 | OG0000017/OG0000219/OG0000394/OG0002466 | 4 |
| GO:0071704 | organic substance metabolic process | 4/25 | 2368/12891 | 0.699767968602346 | 0.740620712532049 | 0.502162760697887 | OG0000017/OG0000219/OG0000394/OG0002466 | 4 |
| GO:0005576 | extracellular region | 1/25 | 613/12891 | 0.704525331244113 | 0.744254203306751 | 0.504626374971051 | OG0000394 | 1 |
| GO:1901362 | organic cyclic compound biosynthetic process | 1/25 | 617/12891 | 0.706924820789858 | 0.745387897343152 | 0.50539505307777 | OG0000017 | 1 |
| GO:1901363 | heterocyclic compound binding | 2/25 | 1374/12891 | 0.762342683987357 | 0.802035240537355 | 0.54380362815981 | OG0000169/OG0000257 | 2 |
| GO:0008152 | metabolic process | 4/25 | 2569/12891 | 0.763591964397592 | 0.802035240537355 | 0.54380362815981 | OG0000017/OG0000219/OG0000394/OG0002466 | 4 |
| GO:0097159 | organic cyclic compound binding | 2/25 | 1381/12891 | 0.764930407345235 | 0.802035240537355 | 0.54380362815981 | OG0000169/OG0000257 | 2 |
| GO:0034645 | cellular macromolecule biosynthetic process | 1/25 | 730/12891 | 0.767480430134531 | 0.803210431537442 | 0.544600442435951 | OG0000017 | 1 |
| GO:0044271 | cellular nitrogen compound biosynthetic process | 1/25 | 741/12891 | 0.772686863355153 | 0.807156165809658 | 0.547275767040825 | OG0000017 | 1 |
| GO:0009059 | macromolecule biosynthetic process | 1/25 | 752/12891 | 0.777781280016377 | 0.810413140292104 | 0.549484092125382 | OG0000017 | 1 |
| GO:0005622 | intracellular anatomical structure | 8/25 | 4835/12891 | 0.778688782487075 | 0.810413140292104 | 0.549484092125382 | OG0000017/OG0000169/OG0000219/OG0000257/OG0000394/OG0000905/OG0002466/OG0003793 | 8 |
| GO:0005524 | ATP binding | 2/25 | 1424/12891 | 0.780326120264451 | 0.810616043601888 | 0.549621666574047 | OG0000169/OG0000257 | 2 |
| GO:0003674 | molecular_function | 8/25 | 4855/12891 | 0.783398974437017 | 0.811210313300036 | 0.550024599015945 | OG0000017/OG0000169/OG0000219/OG0000257/OG0000394/OG0000905/OG0002466/OG0003793 | 8 |
| GO:0044260 | cellular macromolecule metabolic process | 2/25 | 1434/12891 | 0.783785053597722 | 0.811210313300036 | 0.550024599015945 | OG0000017/OG0000394 | 2 |
| GO:0016740 | transferase activity | 1/25 | 812/12891 | 0.803696066681107 | 0.830288951240408 | 0.562960480144274 | OG0002466 | 1 |
| GO:0016070 | RNA metabolic process | 1/25 | 820/12891 | 0.80692388137125 | 0.832093984092922 | 0.56418434583562 | OG0000017 | 1 |
| GO:0044267 | cellular protein metabolic process | 1/25 | 854/12891 | 0.820082546392073 | 0.844114269363269 | 0.572334454971912 | OG0000394 | 1 |
| GO:0043231 | intracellular membrane-bounded organelle | 7/25 | 4573/12891 | 0.839116688816271 | 0.860019961000839 | 0.583118984608173 | OG0000017/OG0000169/OG0000219/OG0000257/OG0000394/OG0000905/OG0003793 | 7 |
| GO:0043227 | membrane-bounded organelle | 7/25 | 4574/12891 | 0.839312171410287 | 0.860019961000839 | 0.583118984608173 | OG0000017/OG0000169/OG0000219/OG0000257/OG0000394/OG0000905/OG0003793 | 7 |
| GO:0110165 | cellular anatomical entity | 8/25 | 5126/12891 | 0.840960039431864 | 0.860019961000839 | 0.583118984608173 | OG0000017/OG0000169/OG0000219/OG0000257/OG0000394/OG0000905/OG0002466/OG0003793 | 8 |
| GO:0043229 | intracellular organelle | 7/25 | 4592/12891 | 0.842802359429398 | 0.860019961000839 | 0.583118984608173 | OG0000017/OG0000169/OG0000219/OG0000257/OG0000394/OG0000905/OG0003793 | 7 |
| GO:0043226 | organelle | 7/25 | 4594/12891 | 0.84318683009157 | 0.860019961000839 | 0.583118984608173 | OG0000017/OG0000169/OG0000219/OG0000257/OG0000394/OG0000905/OG0003793 | 7 |
| GO:0010467 | gene expression | 1/25 | 943/12891 | 0.850577363977162 | 0.864420395217296 | 0.586102609231432 | OG0000017 | 1 |
| GO:0019538 | protein metabolic process | 1/25 | 943/12891 | 0.850577363977162 | 0.864420395217296 | 0.586102609231432 | OG0000394 | 1 |
| GO:0005575 | cellular_component | 8/25 | 5202/12891 | 0.855014884122692 | 0.867361669452983 | 0.588096880205993 | OG0000017/OG0000169/OG0000219/OG0000257/OG0000394/OG0000905/OG0002466/OG0003793 | 8 |
| GO:0008150 | biological_process | 7/25 | 4787/12891 | 0.877217248816099 | 0.888281250152519 | 0.602280975005079 | OG0000017/OG0000169/OG0000219/OG0000257/OG0000394/OG0002466/OG0003793 | 7 |
| GO:0003676 | nucleic acid binding | 1/25 | 1061/12891 | 0.883441019057156 | 0.892974555234032 | 0.605463174741936 | OG0000257 | 1 |
| GO:0043170 | macromolecule metabolic process | 2/25 | 1830/12891 | 0.888463238021936 | 0.896438671038291 | 0.607811947772732 | OG0000017/OG0000394 | 2 |
| GO:0044249 | cellular biosynthetic process | 1/25 | 1102/12891 | 0.893140543009053 | 0.899542984177577 | 0.609916763948835 | OG0000017 | 1 |
| GO:1901576 | organic substance biosynthetic process | 1/25 | 1120/12891 | 0.897149634459519 | 0.90196439099508 | 0.611558549429142 | OG0000017 | 1 |
| GO:0009058 | biosynthetic process | 1/25 | 1203/12891 | 0.913841484174901 | 0.91710520376124 | 0.621824468555102 | OG0000017 | 1 |
| GO:0016021 | integral component of membrane | 1/25 | 1241/12891 | 0.920584575963148 | 0.92222554668679 | 0.625296212587784 | OG0000169 | 1 |
| GO:0031224 | intrinsic component of membrane | 1/25 | 1285/12891 | 0.927759885987 | 0.927759885987 | 0.629048658414111 | OG0000169 | 1 |

**Table S4e. Related to Figure 3. Enriched Pfam domains in orthogroups expanded in Chara with respect to Mesostigma.**

| ID | Description | GeneRatio | BgRatio | pvalue | p.adjust | qvalue | geneID | Count |
|---|---|---|---|---|---|---|---|---|
| **PF00078** | RVT_1 | 6/33 | 40/13933 | 0.0000000000395805506374271 | 0.00000000237483303824562 | 0.000000000499964850156973 | OG0000815/OG0002659/OG0005467/OG0006091/OG0000005/OG0000056 | 6 |
| **PF17917** | RT_RNaseH | 5/33 | 26/13933 | 0.00000000344693848007659 | 0.0000000844648920096167 | 0.0000000177820825283404 | OG0000815/OG0002659/OG0005467/OG0000005/OG0000056 | 5 |
| **PF17919** | RT_RNaseH_2 | 5/33 | 27/13933 | 0.00000000422324460048083 | 0.0000000844648920096167 | 0.0000000177820825283404 | OG0000815/OG0002659/OG0005467/OG0000005/OG0000056 | 5 |
| **PF17921** | Integrase_H2C2 | 4/33 | 18/13933 | 0.0000000779366974551079 | 0.00000116905046182662 | 0.000000246115886700341 | OG0000815/OG0003409/OG0005467/OG0000056 | 4 |
| **PF13692** | Glyco_trans_1_4 | 2/33 | 26/13933 | 0.00170623431417397 | 0.0129189693533345 | 0.00271978302175463 | OG0000433/OG0000512 | 2 |
| **PF00534** | Glycos_transf_1 | 2/33 | 30/13933 | 0.00227025325656954 | 0.0129189693533345 | 0.00271978302175463 | OG0000433/OG0000512 | 2 |
| **PF00079** | Serpin | 1/33 | 1/13933 | 0.00236847771477799 | 0.0129189693533345 | 0.00271978302175463 | OG0000394 | 1 |
| **PF12142** | PPO1_DWL | 1/33 | 1/13933 | 0.00236847771477799 | 0.0129189693533345 | 0.00271978302175463 | OG0000555 | 1 |
| **PF15787** | DUF4704 | 1/33 | 1/13933 | 0.00236847771477799 | 0.0129189693533345 | 0.00271978302175463 | OG0000905 | 1 |
| **PF16057** | DUF4800 | 1/33 | 1/13933 | 0.00236847771477799 | 0.0129189693533345 | 0.00271978302175463 | OG0000905 | 1 |
| **PF20425** | Neurobeachin | 1/33 | 1/13933 | 0.00236847771477799 | 0.0129189693533345 | 0.00271978302175463 | OG0000905 | 1 |
| **PF00191** | Annexin | 1/33 | 2/13933 | 0.00473151534293026 | 0.0177431825359885 | 0.00373540684968178 | OG0000705 | 1 |
| **PF00305** | Lipoxygenase | 1/33 | 2/13933 | 0.00473151534293026 | 0.0177431825359885 | 0.00373540684968178 | OG0000108 | 1 |
| **PF00862** | Sucrose_synth | 1/33 | 2/13933 | 0.00473151534293026 | 0.0177431825359885 | 0.00373540684968178 | OG0000433 | 1 |
| **PF03951** | Gln-synt_N | 1/33 | 2/13933 | 0.00473151534293026 | 0.0177431825359885 | 0.00373540684968178 | OG0000306 | 1 |
| **PF12357** | PLD_C | 1/33 | 2/13933 | 0.00473151534293026 | 0.0177431825359885 | 0.00373540684968178 | OG0000219 | 1 |
| **PF00063** | Myosin_head | 1/33 | 3/13933 | 0.00708912499002523 | 0.0193339772455234 | 0.00407031099905755 | OG0000169 | 1 |
| **PF00182** | Glyco_hydro_19 | 1/33 | 3/13933 | 0.00708912499002523 | 0.0193339772455234 | 0.00407031099905755 | OG0000473 | 1 |
| **PF00264** | Tyrosinase | 1/33 | 3/13933 | 0.00708912499002523 | 0.0193339772455234 | 0.00407031099905755 | OG0000555 | 1 |
| **PF01843** | DIL | 1/33 | 3/13933 | 0.00708912499002523 | 0.0193339772455234 | 0.00407031099905755 | OG0000169 | 1 |
| **PF14844** | PH_BEACH | 1/33 | 3/13933 | 0.00708912499002523 | 0.0193339772455234 | 0.00407031099905755 | OG0000905 | 1 |
| **PF17886** | ArsA_HSP20 | 1/33 | 3/13933 | 0.00708912499002523 | 0.0193339772455234 | 0.00407031099905755 | OG0000017 | 1 |
| **PF00614** | PLDc | 1/33 | 4/13933 | 0.00944131873556198 | 0.0226591649653487 | 0.00477035051902079 | OG0000219 | 1 |
| **PF02138** | Beach | 1/33 | 4/13933 | 0.00944131873556198 | 0.0226591649653487 | 0.00477035051902079 | OG0000905 | 1 |
| **PF10162** | G8 | 1/33 | 4/13933 | 0.00944131873556198 | 0.0226591649653487 | 0.00477035051902079 | OG0000932 | 1 |
| **PF05970** | PIF1 | 1/33 | 5/13933 | 0.0117881086330228 | 0.027203327614668 | 0.00572701633993012 | OG0000381 | 1 |
| **PF03143** | GTP_EFTU_D3 | 1/33 | 6/13933 | 0.0141295067099259 | 0.0308728593147811 | 0.00649954932942759 | OG0000345 | 1 |
| **PF00120** | Gln-synt_C | 1/33 | 7/13933 | 0.0164655249678832 | 0.0308728593147811 | 0.00649954932942759 | OG0000306 | 1 |
| **PF00909** | Ammonium_transp | 1/33 | 7/13933 | 0.0164655249678832 | 0.0308728593147811 | 0.00649954932942759 | OG0000132 | 1 |
| **PF02225** | PA | 1/33 | 7/13933 | 0.0164655249678832 | 0.0308728593147811 | 0.00649954932942759 | OG0000650 | 1 |
| **PF04937** | DUF659 | 1/33 | 7/13933 | 0.0164655249678832 | 0.0308728593147811 | 0.00649954932942759 | OG0000730 | 1 |
| **PF13359** | DDE_Tnp_4 | 1/33 | 7/13933 | 0.0164655249678832 | 0.0308728593147811 | 0.00649954932942759 | OG0003793 | 1 |
| **PF00011** | HSP20 | 1/33 | 8/13933 | 0.0187961753826511 | 0.0341748643320929 | 0.00719470828044061 | OG0000017 | 1 |
| **PF08323** | Glyco_transf_5 | 1/33 | 9/13933 | 0.0211214699041861 | 0.0342510322770585 | 0.00721074363727547 | OG0000512 | 1 |
| **PF12763** | EF-hand_4 | 1/33 | 9/13933 | 0.0211214699041861 | 0.0342510322770585 | 0.00721074363727547 | OG0000153 | 1 |
| **PF13091** | PLDc_2 | 1/33 | 9/13933 | 0.0211214699041861 | 0.0342510322770585 | 0.00721074363727547 | OG0000219 | 1 |
| **PF14658** | EF-hand_9 | 1/33 | 9/13933 | 0.0211214699041861 | 0.0342510322770585 | 0.00721074363727547 | OG0000153 | 1 |
| **PF00022** | Actin | 1/33 | 10/13933 | 0.0234414204566972 | 0.0370127691421534 | 0.00779216192466388 | OG0000257 | 1 |
| **PF00332** | Glyco_hydro_17 | 1/33 | 11/13933 | 0.0257560389386999 | 0.0386340584080499 | 0.00813348598064208 | OG0000162 | 1 |
| **PF00665** | rve | 1/33 | 11/13933 | 0.0257560389386999 | 0.0386340584080499 | 0.00813348598064208 | OG0003409 | 1 |
| **PF00082** | Peptidase_S8 | 1/33 | 12/13933 | 0.028065337223072 | 0.0410712252044957 | 0.00864657372726224 | OG0000650 | 1 |
| **PF00089** | Trypsin | 1/33 | 17/13933 | 0.0395324394430789 | 0.0527099192574386 | 0.0110968251068292 | OG0001496 | 1 |
| **PF01694** | Rhomboid | 1/33 | 17/13933 | 0.0395324394430789 | 0.0527099192574386 | 0.0110968251068292 | OG0000564 | 1 |
| **PF02798** | GST_N | 1/33 | 17/13933 | 0.0395324394430789 | 0.0527099192574386 | 0.0110968251068292 | OG0002466 | 1 |
| **PF13385** | Laminin_G_3 | 1/33 | 17/13933 | 0.0395324394430789 | 0.0527099192574386 | 0.0110968251068292 | OG0000905 | 1 |
| **PF03144** | GTP_EFTU_D2 | 1/33 | 18/13933 | 0.0418100644429623 | 0.0545348666647335 | 0.0114810245609965 | OG0000345 | 1 |
| **PF01833** | TIG | 1/33 | 21/13933 | 0.0486115592905373 | 0.0607644491131716 | 0.012792515602773 | OG0000932 | 1 |
| **PF13409** | GST_N_2 | 1/33 | 21/13933 | 0.0486115592905373 | 0.0607644491131716 | 0.012792515602773 | OG0002466 | 1 |
| **PF13417** | GST_N_3 | 1/33 | 23/13933 | 0.0531198554447009 | 0.065044720952695 | 0.0136936254637253 | OG0002466 | 1 |
| **PF00612** | IQ | 1/33 | 25/13933 | 0.0576074327592159 | 0.069128919311059 | 0.0145534566970651 | OG0000169 | 1 |
| **PF00009** | GTP_EFTU | 1/33 | 35/13933 | 0.0797377518004423 | 0.0938091197652262 | 0.0197492883716266 | OG0000345 | 1 |
| **PF00168** | C2 | 1/33 | 40/13933 | 0.0906130967325195 | 0.104553573152907 | 0.0220112785585068 | OG0000219 | 1 |
| **PF20426** | NBCH_WD40 | 1/33 | 81/13933 | 0.175211642295079 | 0.19755576182083 | 0.0415906866991222 | OG0000905 | 1 |
| **PF13202** | EF-hand_5 | 1/33 | 83/13933 | 0.179136934654191 | 0.19755576182083 | 0.0415906866991222 | OG0000153 | 1 |
| **PF13833** | EF-hand_8 | 1/33 | 84/13933 | 0.181092781669094 | 0.19755576182083 | 0.0415906866991222 | OG0000153 | 1 |
| **PF13405** | EF-hand_6 | 1/33 | 107/13933 | 0.224850195128283 | 0.240910923351732 | 0.0507180891266803 | OG0000153 | 1 |
| **PF00036** | EF-hand_1 | 1/33 | 111/13933 | 0.232225072443079 | 0.244447444676925 | 0.0514626199319843 | OG0000153 | 1 |
| **PF13499** | EF-hand_7 | 1/33 | 122/13933 | 0.252156883706955 | 0.260851948662367 | 0.0549161997183931 | OG0000153 | 1 |
| **PF00076** | RRM_1 | 1/33 | 145/13933 | 0.292224226275827 | 0.297177179263552 | 0.0625636166870637 | OG0000711 | 1 |
| **PF00400** | WD40 | 1/33 | 197/13933 | 0.375287955380493 | 0.375287955380493 | 0.0790079906064196 | OG0000905 | 1 |

**Table S4f. Related to Figure 3. Enriched GO terms in orthogroups contracted in Chara with respect to Mesostigma.**

| ID | Description | GeneRatio | BgRatio | pvalue | p.adjust | qvalue | geneID | Count |
|---|---|---|---|---|---|---|---|---|
| GO:0010037 | response to carbon dioxide | 1/6 | 2/12891 | 0.000930701467258599 | 0.0383283937870334 | 0.0196019094342822 | OG0000180 | 1 |
| GO:0071490 | cellular response to far red light | 1/6 | 2/12891 | 0.000930701467258599 | 0.0383283937870334 | 0.0196019094342822 | OG0000047 | 1 |
| GO:0070141 | response to UV-A | 1/6 | 3/12891 | 0.00139578144174823 | 0.0383283937870334 | 0.0196019094342822 | OG0000047 | 1 |
| GO:0071491 | cellular response to red light | 1/6 | 3/12891 | 0.00139578144174823 | 0.0383283937870334 | 0.0196019094342822 | OG0000047 | 1 |
| GO:0071492 | cellular response to UV-A | 1/6 | 3/12891 | 0.00139578144174823 | 0.0383283937870334 | 0.0196019094342822 | OG0000047 | 1 |
| GO:0071486 | cellular response to high light intensity | 1/6 | 4/12891 | 0.00186068098483871 | 0.0383283937870334 | 0.0196019094342822 | OG0000047 | 1 |
| GO:0016020 | membrane | 5/6 | 2782/12891 | 0.00229779364726571 | 0.0383283937870334 | 0.0196019094342822 | OG0000029/OG0000047/OG0000071/OG0000180/OG0000363 | 5 |
| GO:0005388 | P-type calcium transporter activity | 1/6 | 5/12891 | 0.00232540015253402 | 0.0383283937870334 | 0.0196019094342822 | OG0000180 | 1 |
| GO:0006730 | one-carbon metabolic process | 1/6 | 5/12891 | 0.00232540015253402 | 0.0383283937870334 | 0.0196019094342822 | OG0000180 | 1 |
| GO:0071484 | cellular response to light intensity | 1/6 | 5/12891 | 0.00232540015253402 | 0.0383283937870334 | 0.0196019094342822 | OG0000047 | 1 |
| GO:0009535 | chloroplast thylakoid membrane | 2/6 | 179/12891 | 0.002772515244408 | 0.0383283937870334 | 0.0196019094342822 | OG0000047/OG0000180 | 2 |
| GO:0034644 | cellular response to UV | 1/6 | 6/12891 | 0.00278993900082558 | 0.0383283937870334 | 0.0196019094342822 | OG0000047 | 1 |
| GO:0055035 | plastid thylakoid membrane | 2/6 | 180/12891 | 0.00280308372892831 | 0.0383283937870334 | 0.0196019094342822 | OG0000047/OG0000180 | 2 |
| GO:0034357 | photosynthetic membrane | 2/6 | 187/12891 | 0.00302156804345896 | 0.0383283937870334 | 0.0196019094342822 | OG0000047/OG0000180 | 2 |
| GO:0042651 | thylakoid membrane | 2/6 | 189/12891 | 0.00308543699237832 | 0.0383283937870334 | 0.0196019094342822 | OG0000047/OG0000180 | 2 |
| GO:0004089 | carbonate dehydratase activity | 1/6 | 7/12891 | 0.00325429758569151 | 0.0383283937870334 | 0.0196019094342822 | OG0000180 | 1 |
| GO:0010117 | photoprotection | 1/6 | 7/12891 | 0.00325429758569151 | 0.0383283937870334 | 0.0196019094342822 | OG0000047 | 1 |
| GO:0010380 | regulation of chlorophyll biosynthetic process | 1/6 | 7/12891 | 0.00325429758569151 | 0.0383283937870334 | 0.0196019094342822 | OG0000047 | 1 |
| GO:1901463 | regulation of tetrapyrrole biosynthetic process | 1/6 | 8/12891 | 0.00371847596309627 | 0.0407728063070174 | 0.0208520310308977 | OG0000047 | 1 |
| GO:0090056 | regulation of chlorophyll metabolic process | 1/6 | 9/12891 | 0.00418247418899265 | 0.0407728063070174 | 0.0208520310308977 | OG0000047 | 1 |
| GO:0009534 | chloroplast thylakoid | 2/6 | 221/12891 | 0.00419389570915631 | 0.0407728063070174 | 0.0208520310308977 | OG0000047/OG0000180 | 2 |
| GO:0031976 | plastid thylakoid | 2/6 | 222/12891 | 0.00423114027714332 | 0.0407728063070174 | 0.0208520310308977 | OG0000047/OG0000180 | 2 |
| GO:1901401 | regulation of tetrapyrrole metabolic process | 1/6 | 10/12891 | 0.00464629231931912 | 0.0426591113189272 | 0.0218167252524802 | OG0000047 | 1 |
| GO:0015662 | P-type ion transporter activity | 1/6 | 11/12891 | 0.00510993041000196 | 0.0426591113189272 | 0.0218167252524802 | OG0000180 | 1 |
| GO:0010030 | positive regulation of seed germination | 1/6 | 12/12891 | 0.00557338851695355 | 0.0426591113189272 | 0.0218167252524802 | OG0000047 | 1 |
| GO:0009579 | thylakoid | 2/6 | 257/12891 | 0.00563269301932748 | 0.0426591113189272 | 0.0218167252524802 | OG0000047/OG0000180 | 2 |
| GO:0042170 | plastid membrane | 2/6 | 261/12891 | 0.00580489586019223 | 0.0426591113189272 | 0.0218167252524802 | OG0000047/OG0000180 | 2 |
| GO:0015085 | calcium ion transmembrane transporter activity | 1/6 | 13/12891 | 0.00603666669607461 | 0.0426591113189272 | 0.0218167252524802 | OG0000180 | 1 |
| GO:0071483 | cellular response to blue light | 1/6 | 13/12891 | 0.00603666669607461 | 0.0426591113189272 | 0.0218167252524802 | OG0000047 | 1 |
| GO:0140358 | P-type transmembrane transporter activity | 1/6 | 13/12891 | 0.00603666669607461 | 0.0426591113189272 | 0.0218167252524802 | OG0000180 | 1 |
| GO:0019829 | ATPase-coupled cation transmembrane transporter activity | 1/6 | 14/12891 | 0.00649976500325111 | 0.044450005828685 | 0.0227326246293672 | OG0000180 | 1 |
| GO:0010218 | response to far red light | 1/6 | 16/12891 | 0.00742542222525544 | 0.0491934222423173 | 0.0251585029342536 | OG0000047 | 1 |
| GO:0016702 | oxidoreductase activity, acting on single donors with incorporation of molecular oxygen, incorporation of two atoms of oxygen | 1/6 | 17/12891 | 0.00788798125179113 | 0.050579327243362 | 0.0258672825524642 | OG0000214 | 1 |
| GO:0022857 | transmembrane transporter activity | 2/6 | 311/12891 | 0.00816147343367897 | 0.050579327243362 | 0.0258672825524642 | OG0000180/OG0000363 | 2 |
| GO:0034605 | cellular response to heat | 1/6 | 18/12891 | 0.00835036062980032 | 0.050579327243362 | 0.0258672825524642 | OG0000047 | 1 |
| GO:0034062 | 5'-3' RNA polymerase activity | 1/6 | 20/12891 | 0.00927458066351261 | 0.0531408405585047 | 0.0271772918447169 | OG0000071 | 1 |
| GO:0097747 | RNA polymerase activity | 1/6 | 20/12891 | 0.00927458066351261 | 0.0531408405585047 | 0.0271772918447169 | OG0000071 | 1 |
| GO:0005887 | integral component of plasma membrane | 1/6 | 22/12891 | 0.0101980827728108 | 0.0535823611279056 | 0.027403094320627 | OG0000180 | 1 |
| GO:0015995 | chlorophyll biosynthetic process | 1/6 | 22/12891 | 0.0101980827728108 | 0.0535823611279056 | 0.027403094320627 | OG0000047 | 1 |
| GO:0010114 | response to red light | 1/6 | 23/12891 | 0.0106595647452532 | 0.0535823611279056 | 0.027403094320627 | OG0000047 | 1 |
| GO:0009521 | photosystem | 1/6 | 24/12891 | 0.0111208674039049 | 0.0535823611279056 | 0.027403094320627 | OG0000047 | 1 |
| GO:0009522 | photosystem I | 1/6 | 24/12891 | 0.0111208674039049 | 0.0535823611279056 | 0.027403094320627 | OG0000047 | 1 |
| GO:0016836 | hydro-lyase activity | 1/6 | 24/12891 | 0.0111208674039049 | 0.0535823611279056 | 0.027403094320627 | OG0000180 | 1 |
| GO:0042626 | ATPase-coupled transmembrane transporter activity | 1/6 | 24/12891 | 0.0111208674039049 | 0.0535823611279056 | 0.027403094320627 | OG0000180 | 1 |
| GO:0009526 | plastid envelope | 2/6 | 371/12891 | 0.0114755033724941 | 0.0540623714437498 | 0.0276485812249564 | OG0000047/OG0000180 | 2 |
| GO:0006779 | porphyrin-containing compound biosynthetic process | 1/6 | 26/12891 | 0.0120429350027973 | 0.0552246752406493 | 0.0282430067020203 | OG0000047 | 1 |
| GO:0022853 | active ion transmembrane transporter activity | 1/6 | 27/12891 | 0.0125037000544866 | 0.0552246752406493 | 0.0282430067020203 | OG0000180 | 1 |
| GO:0033014 | tetrapyrrole biosynthetic process | 1/6 | 27/12891 | 0.0125037000544866 | 0.0552246752406493 | 0.0282430067020203 | OG0000047 | 1 |
| GO:0016021 | integral component of membrane | 3/6 | 1241/12891 | 0.0142442203728236 | 0.0574740918222852 | 0.0293934034642273 | OG0000047/OG0000071/OG0000180 | 3 |
| GO:0015994 | chlorophyll metabolic process | 1/6 | 31/12891 | 0.0143449699091588 | 0.0574740918222852 | 0.0293934034642273 | OG0000047 | 1 |
| GO:0016835 | carbon-oxygen lyase activity | 1/6 | 31/12891 | 0.0143449699091588 | 0.0574740918222852 | 0.0293934034642273 | OG0000180 | 1 |
| GO:0033554 | cellular response to stress | 2/6 | 417/12891 | 0.0143628258455704 | 0.0574740918222852 | 0.0293934034642273 | OG0000047/OG0000071 | 2 |
| GO:0071489 | cellular response to red or far red light | 1/6 | 32/12891 | 0.0148048400631665 | 0.0574740918222852 | 0.0293934034642273 | OG0000047 | 1 |
| GO:0006778 | porphyrin-containing compound metabolic process | 1/6 | 33/12891 | 0.01526453140461 | 0.0574740918222852 | 0.0293934034642273 | OG0000047 | 1 |
| GO:0033013 | tetrapyrrole metabolic process | 1/6 | 33/12891 | 0.01526453140461 | 0.0574740918222852 | 0.0293934034642273 | OG0000047 | 1 |
| GO:0031224 | intrinsic component of membrane | 3/6 | 1285/12891 | 0.0156859114442444 | 0.0574740918222852 | 0.0293934034642273 | OG0000047/OG0000071/OG0000180 | 3 |
| GO:0009644 | response to high light intensity | 1/6 | 34/12891 | 0.0157240439891158 | 0.0574740918222852 | 0.0293934034642273 | OG0000047 | 1 |
| GO:0010224 | response to UV-B | 1/6 | 34/12891 | 0.0157240439891158 | 0.0574740918222852 | 0.0293934034642273 | OG0000047 | 1 |
| GO:0010029 | regulation of seed germination | 1/6 | 35/12891 | 0.0161833778722972 | 0.0578396232666912 | 0.0295803435773049 | OG0000047 | 1 |
| GO:0009637 | response to blue light | 1/6 | 36/12891 | 0.0166425331097555 | 0.0578396232666912 | 0.0295803435773049 | OG0000047 | 1 |
| GO:0048582 | positive regulation of post-embryonic development | 1/6 | 36/12891 | 0.0166425331097555 | 0.0578396232666912 | 0.0295803435773049 | OG0000047 | 1 |
| GO:1900140 | regulation of seedling development | 1/6 | 39/12891 | 0.0180189275036071 | 0.0616131069478179 | 0.0315101788263418 | OG0000047 | 1 |
| GO:0015399 | primary active transmembrane transporter activity | 1/6 | 41/12891 | 0.0189356315563254 | 0.062971578983354 | 0.0322049286756974 | OG0000180 | 1 |
| GO:0031984 | organelle subcompartment | 2/6 | 483/12891 | 0.0190102879949748 | 0.062971578983354 | 0.0322049286756974 | OG0000047/OG0000180 | 2 |
| GO:0022890 | inorganic cation transmembrane transporter activity | 1/6 | 42/12891 | 0.0193937160863382 | 0.0632533509277493 | 0.0323490325002889 | OG0000180 | 1 |
| GO:0003899 | DNA-directed 5'-3' RNA polymerase activity | 1/6 | 44/12891 | 0.0203093504312207 | 0.064262422259982 | 0.0328650918211427 | OG0000071 | 1 |
| GO:0051240 | positive regulation of multicellular organismal process | 1/6 | 44/12891 | 0.0203093504312207 | 0.064262422259982 | 0.0328650918211427 | OG0000047 | 1 |
| GO:0046148 | pigment biosynthetic process | 1/6 | 48/12891 | 0.0221384818138827 | 0.0685313346246835 | 0.0350482992370526 | OG0000047 | 1 |
| GO:0046873 | metal ion transmembrane transporter activity | 1/6 | 50/12891 | 0.0230519797388906 | 0.0685313346246835 | 0.0350482992370526 | OG0000180 | 1 |
| GO:0140359 | ABC-type transporter activity | 1/6 | 50/12891 | 0.0230519797388906 | 0.0685313346246835 | 0.0350482992370526 | OG0000029 | 1 |
| GO:0031967 | organelle envelope | 2/6 | 550/12891 | 0.0243113875152729 | 0.0685313346246835 | 0.0350482992370526 | OG0000047/OG0000180 | 2 |
| GO:0031975 | envelope | 2/6 | 550/12891 | 0.0243113875152729 | 0.0685313346246835 | 0.0350482992370526 | OG0000047/OG0000180 | 2 |
| GO:0050896 | response to stimulus | 3/6 | 1508/12891 | 0.024321531462933 | 0.0685313346246835 | 0.0350482992370526 | OG0000047/OG0000071/OG0000180 | 3 |
| GO:0080167 | response to karrikin | 1/6 | 53/12891 | 0.0244208931653644 | 0.0685313346246835 | 0.0350482992370526 | OG0000047 | 1 |
| GO:0071482 | cellular response to light stimulus | 1/6 | 54/12891 | 0.0248768422727803 | 0.0685313346246835 | 0.0350482992370526 | OG0000047 | 1 |
| GO:0009411 | response to UV | 1/6 | 55/12891 | 0.0253326137884274 | 0.0685313346246835 | 0.0350482992370526 | OG0000047 | 1 |
| GO:0031226 | intrinsic component of plasma membrane | 1/6 | 55/12891 | 0.0253326137884274 | 0.0685313346246835 | 0.0350482992370526 | OG0000180 | 1 |
| GO:0071478 | cellular response to radiation | 1/6 | 55/12891 | 0.0253326137884274 | 0.0685313346246835 | 0.0350482992370526 | OG0000047 | 1 |
| GO:0009523 | photosystem II | 1/6 | 57/12891 | 0.0262436242657696 | 0.0685313346246835 | 0.0350482992370526 | OG0000047 | 1 |
| GO:0018130 | heterocycle biosynthetic process | 2/6 | 578/12891 | 0.0266942088329813 | 0.0685313346246835 | 0.0350482992370526 | OG0000047/OG0000071 | 2 |
| GO:0051094 | positive regulation of developmental process | 1/6 | 58/12891 | 0.0266988633381091 | 0.0685313346246835 | 0.0350482992370526 | OG0000047 | 1 |
| GO:0019438 | aromatic compound biosynthetic process | 2/6 | 581/12891 | 0.0269552255561898 | 0.0685313346246835 | 0.0350482992370526 | OG0000047/OG0000071 | 2 |
| GO:0009642 | response to light intensity | 1/6 | 59/12891 | 0.0271539250399689 | 0.0685313346246835 | 0.0350482992370526 | OG0000047 | 1 |
| GO:0015318 | inorganic molecular entity transmembrane transporter activity | 1/6 | 59/12891 | 0.0271539250399689 | 0.0685313346246835 | 0.0350482992370526 | OG0000180 | 1 |
| GO:0042440 | pigment metabolic process | 1/6 | 60/12891 | 0.0276088094266398 | 0.0688596188052663 | 0.0352161903522464 | OG0000047 | 1 |
| GO:1901362 | organic cyclic compound biosynthetic process | 2/6 | 617/12891 | 0.0301722738194436 | 0.074378163368861 | 0.038038484741771 | OG0000047/OG0000071 | 2 |
| GO:0008324 | cation transmembrane transporter activity | 1/6 | 69/12891 | 0.031694798839619 | 0.0772333029195314 | 0.0394986603808924 | OG0000180 | 1 |
| GO:0016779 | nucleotidyltransferase activity | 1/6 | 70/12891 | 0.0321479131125068 | 0.0774472452255845 | 0.0396080747678014 | OG0000071 | 1 |
| GO:0009845 | seed germination | 1/6 | 74/12891 | 0.0339586036768268 | 0.0808901570728908 | 0.0413688489498895 | OG0000047 | 1 |
| GO:0009507 | chloroplast | 3/6 | 1731/12891 | 0.0352705861991522 | 0.083081825269114 | 0.042489712029388 | OG0000047/OG0000071/OG0000180 | 3 |
| GO:0022804 | active transmembrane transporter activity | 1/6 | 79/12891 | 0.0362179963286718 | 0.0843759914470157 | 0.0431515745732007 | OG0000180 | 1 |
| GO:0009639 | response to red or far red light | 1/6 | 81/12891 | 0.0371205195193937 | 0.0855385884577332 | 0.0437461500056928 | OG0000047 | 1 |
| GO:0090351 | seedling development | 1/6 | 84/12891 | 0.0384729836773632 | 0.0869503572340831 | 0.0444681568774109 | OG0000047 | 1 |
| GO:0009408 | response to heat | 1/6 | 86/12891 | 0.0393737466720376 | 0.0869503572340831 | 0.0444681568774109 | OG0000047 | 1 |
| GO:0071214 | cellular response to abiotic stimulus | 1/6 | 86/12891 | 0.0393737466720376 | 0.0869503572340831 | 0.0444681568774109 | OG0000047 | 1 |
| GO:0104004 | cellular response to environmental stimulus | 1/6 | 86/12891 | 0.0393737466720376 | 0.0869503572340831 | 0.0444681568774109 | OG0000047 | 1 |
| GO:0009628 | response to abiotic stimulus | 2/6 | 717/12891 | 0.0399022794928425 | 0.0872091056956969 | 0.0446004860310664 | OG0000047/OG0000071 | 2 |
| GO:0044271 | cellular nitrogen compound biosynthetic process | 2/6 | 741/12891 | 0.0424042003081347 | 0.089879399549858 | 0.0459661278730654 | OG0000047/OG0000071 | 2 |
| GO:0071456 | cellular response to hypoxia | 1/6 | 93/12891 | 0.0425208814782235 | 0.089879399549858 | 0.0459661278730654 | OG0000071 | 1 |
| GO:0036294 | cellular response to decreased oxygen levels | 1/6 | 94/12891 | 0.0429697699538549 | 0.089879399549858 | 0.0459661278730654 | OG0000071 | 1 |
| GO:0071453 | cellular response to oxygen levels | 1/6 | 94/12891 | 0.0429697699538549 | 0.089879399549858 | 0.0459661278730654 | OG0000071 | 1 |
| GO:0009536 | plastid | 3/6 | 1869/12891 | 0.0432438620475732 | 0.089879399549858 | 0.0459661278730654 | OG0000047/OG0000071/OG0000180 | 3 |
| GO:0016829 | lyase activity | 1/6 | 97/12891 | 0.0443153832710355 | 0.0912122451792186 | 0.0466477718642479 | OG0000180 | 1 |
| GO:0031090 | organelle membrane | 2/6 | 768/12891 | 0.0452930119381444 | 0.0921885423791638 | 0.0471470698364145 | OG0000047/OG0000180 | 2 |
| GO:0015075 | ion transmembrane transporter activity | 1/6 | 100/12891 | 0.0456594195745859 | 0.0921885423791638 | 0.0471470698364145 | OG0000180 | 1 |
| GO:0001666 | response to hypoxia | 1/6 | 101/12891 | 0.0461070814839404 | 0.0922141629678808 | 0.0471601727194226 | OG0000071 | 1 |
| GO:0051716 | cellular response to stimulus | 2/6 | 789/12891 | 0.0475931192240293 | 0.0942966474345254 | 0.0482251970494345 | OG0000047/OG0000071 | 2 |
| GO:0036293 | response to decreased oxygen levels | 1/6 | 106/12891 | 0.0483427670554039 | 0.0948931055836024 | 0.0485302377115742 | OG0000071 | 1 |
| GO:0070482 | response to oxygen levels | 1/6 | 107/12891 | 0.0487893797576069 | 0.0948931055836024 | 0.0485302377115742 | OG0000071 | 1 |
| GO:0042221 | response to chemical | 2/6 | 847/12891 | 0.0541810866541583 | 0.10442173064256 | 0.0534033677069694 | OG0000071/OG0000180 | 2 |
| GO:0048580 | regulation of post-embryonic development | 1/6 | 125/12891 | 0.056798591642184 | 0.108223231562508 | 0.0553475315339542 | OG0000047 | 1 |
| GO:0140098 | catalytic activity, acting on RNA | 1/6 | 126/12891 | 0.057241894826435 | 0.108223231562508 | 0.0553475315339542 | OG0000071 | 1 |
| GO:2000026 | regulation of multicellular organismal development | 1/6 | 127/12891 | 0.0576850243705823 | 0.108223231562508 | 0.0553475315339542 | OG0000047 | 1 |
| GO:0140657 | ATP-dependent activity | 1/6 | 133/12891 | 0.06034015823967 | 0.112211522340439 | 0.0573872234412374 | OG0000180 | 1 |
| GO:0009409 | response to cold | 1/6 | 142/12891 | 0.0643111676925352 | 0.118556239572326 | 0.0606320391060057 | OG0000047 | 1 |
| GO:0015979 | photosynthesis | 1/6 | 148/12891 | 0.0669507301905594 | 0.122358231037919 | 0.0625764538078731 | OG0000047 | 1 |
| GO:0051239 | regulation of multicellular organismal process | 1/6 | 153/12891 | 0.0691456228458203 | 0.125289504643709 | 0.0640755659299999 | OG0000047 | 1 |
| GO:0006950 | response to stress | 2/6 | 1004/12891 | 0.0736390003349626 | 0.132300576872984 | 0.0676611689072359 | OG0000047/OG0000071 | 2 |
| GO:0044237 | cellular metabolic process | 3/6 | 2364/12891 | 0.0794895149880263 | 0.141611572919845 | 0.0724229990602981 | OG0000047/OG0000071/OG0000180 | 3 |
| GO:0005215 | transporter activity | 1/6 | 187/12891 | 0.0839570865048757 | 0.147856380173778 | 0.0756167187581883 | OG0000180 | 1 |
| GO:0098796 | membrane protein complex | 1/6 | 188/12891 | 0.0843897264199392 | 0.147856380173778 | 0.0756167187581883 | OG0000047 | 1 |
| GO:0140640 | catalytic activity, acting on a nucleic acid | 1/6 | 192/12891 | 0.0861185837119034 | 0.149648686450193 | 0.0765333401408633 | OG0000071 | 1 |
| GO:0044249 | cellular biosynthetic process | 2/6 | 1102/12891 | 0.0868848035710738 | 0.149752669569656 | 0.0765865191940146 | OG0000047/OG0000071 | 2 |
| GO:0016887 | ATP hydrolysis activity | 1/6 | 198/12891 | 0.0887067684400429 | 0.151626422216664 | 0.0775447938843914 | OG0000029 | 1 |
| GO:1901576 | organic substance biosynthetic process | 2/6 | 1120/12891 | 0.0894023715900143 | 0.151626422216664 | 0.0775447938843914 | OG0000047/OG0000071 | 2 |
| GO:0046483 | heterocycle metabolic process | 2/6 | 1154/12891 | 0.0942261994306318 | 0.157769220226521 | 0.080686344008598 | OG0000047/OG0000071 | 2 |
| GO:0043167 | ion binding | 2/6 | 1156/12891 | 0.0945126932489063 | 0.157769220226521 | 0.080686344008598 | OG0000071/OG0000180 | 2 |
| GO:0006725 | cellular aromatic compound metabolic process | 2/6 | 1174/12891 | 0.0971045997847481 | 0.159745848133653 | 0.0816972311706366 | OG0000047/OG0000071 | 2 |
| GO:0009266 | response to temperature stimulus | 1/6 | 218/12891 | 0.0972899794651981 | 0.159745848133653 | 0.0816972311706366 | OG0000047 | 1 |
| GO:0008152 | metabolic process | 3/6 | 2569/12891 | 0.0979573597045983 | 0.159745848133653 | 0.0816972311706366 | OG0000047/OG0000071/OG0000180 | 3 |
| GO:1901360 | organic cyclic compound metabolic process | 2/6 | 1201/12891 | 0.101037212800409 | 0.162743119323044 | 0.0832300957809013 | OG0000047/OG0000071 | 2 |
| GO:0009058 | biosynthetic process | 2/6 | 1203/12891 | 0.101330621465291 | 0.162743119323044 | 0.0832300957809013 | OG0000047/OG0000071 | 2 |
| GO:0046914 | transition metal ion binding | 1/6 | 230/12891 | 0.10240749081847 | 0.163236000402373 | 0.0834821650518592 | OG0000180 | 1 |
| GO:0000325 | plant-type vacuole | 1/6 | 249/12891 | 0.110460763094655 | 0.17475881922438 | 0.0893751657403732 | OG0000180 | 1 |
| GO:0034641 | cellular nitrogen compound metabolic process | 2/6 | 1288/12891 | 0.114058877584096 | 0.179114681835766 | 0.0916028412566229 | OG0000047/OG0000071 | 2 |
| GO:0050793 | regulation of developmental process | 1/6 | 262/12891 | 0.115936129895794 | 0.180723967190502 | 0.0924258620686281 | OG0000047 | 1 |
| GO:0009416 | response to light stimulus | 1/6 | 276/12891 | 0.121801240724165 | 0.188480752069511 | 0.0963928374536229 | OG0000047 | 1 |
| GO:0009314 | response to radiation | 1/6 | 285/12891 | 0.125554518408366 | 0.192880854366475 | 0.0986431380325069 | OG0000047 | 1 |
| GO:0009570 | chloroplast stroma | 1/6 | 294/12891 | 0.129294419148351 | 0.197197243593168 | 0.100850626068005 | OG0000180 | 1 |
| GO:0009532 | plastid stroma | 1/6 | 298/12891 | 0.130952313406785 | 0.19829921744456 | 0.101414197600743 | OG0000180 | 1 |
| GO:0005524 | ATP binding | 2/6 | 1424/12891 | 0.135397850523499 | 0.203576910006963 | 0.104113315445468 | OG0000029/OG0000071 | 2 |
| GO:0005773 | vacuole | 1/6 | 324/12891 | 0.141664633134816 | 0.211499311440712 | 0.108164990458755 | OG0000180 | 1 |
| GO:0016772 | transferase activity, transferring phosphorus-containing groups | 1/6 | 341/12891 | 0.148609167348299 | 0.220315688656219 | 0.112673862619615 | OG0000071 | 1 |
| GO:0008270 | zinc ion binding | 1/6 | 371/12891 | 0.160750011851758 | 0.236659739670644 | 0.121032538163239 | OG0000180 | 1 |
| GO:0009987 | cellular process | 3/6 | 3205/12891 | 0.167215058544914 | 0.244479947665668 | 0.125031949402005 | OG0000047/OG0000071/OG0000180 | 3 |
| GO:0048518 | positive regulation of biological process | 1/6 | 391/12891 | 0.168763479364973 | 0.244537394861875 | 0.125061329050512 | OG0000047 | 1 |
| GO:0005783 | endoplasmic reticulum | 1/6 | 393/12891 | 0.169561306814602 | 0.244537394861875 | 0.125061329050512 | OG0000180 | 1 |
| GO:1901566 | organonitrogen compound biosynthetic process | 1/6 | 414/12891 | 0.177900039537282 | 0.254829786364214 | 0.130325064525889 | OG0000047 | 1 |
| GO:0070887 | cellular response to chemical stimulus | 1/6 | 431/12891 | 0.184599228205039 | 0.262651250868914 | 0.134325118368908 | OG0000071 | 1 |
| GO:0032774 | RNA biosynthetic process | 1/6 | 437/12891 | 0.186952758651533 | 0.264226565560834 | 0.135130765902512 | OG0000071 | 1 |
| GO:0032559 | adenyl ribonucleotide binding | 1/6 | 452/12891 | 0.192811830163212 | 0.270007627381809 | 0.138087316883447 | OG0000071 | 1 |
| GO:0030554 | adenyl nucleotide binding | 1/6 | 454/12891 | 0.193590374349222 | 0.270007627381809 | 0.138087316883447 | OG0000071 | 1 |
| GO:0044281 | small molecule metabolic process | 1/6 | 459/12891 | 0.195533997537596 | 0.270935996588042 | 0.138562103518214 | OG0000180 | 1 |
| GO:0035639 | purine ribonucleoside triphosphate binding | 1/6 | 490/12891 | 0.207497536201489 | 0.282784914437586 | 0.144621877790821 | OG0000071 | 1 |
| GO:0055085 | transmembrane transport | 1/6 | 499/12891 | 0.210942916459224 | 0.282784914437586 | 0.144621877790821 | OG0000363 | 1 |
| GO:0032555 | purine ribonucleotide binding | 1/6 | 500/12891 | 0.211324964756613 | 0.282784914437586 | 0.144621877790821 | OG0000071 | 1 |
| GO:0017076 | purine nucleotide binding | 1/6 | 503/12891 | 0.212470184866095 | 0.282784914437586 | 0.144621877790821 | OG0000071 | 1 |
| GO:0032553 | ribonucleotide binding | 1/6 | 503/12891 | 0.212470184866095 | 0.282784914437586 | 0.144621877790821 | OG0000071 | 1 |
| GO:0031326 | regulation of cellular biosynthetic process | 1/6 | 508/12891 | 0.214375805424387 | 0.282784914437586 | 0.144621877790821 | OG0000047 | 1 |
| GO:0034654 | nucleobase-containing compound biosynthetic process | 1/6 | 508/12891 | 0.214375805424387 | 0.282784914437586 | 0.144621877790821 | OG0000071 | 1 |
| GO:1901700 | response to oxygen-containing compound | 1/6 | 509/12891 | 0.214756468039865 | 0.282784914437586 | 0.144621877790821 | OG0000180 | 1 |
| GO:0097367 | carbohydrate derivative binding | 1/6 | 513/12891 | 0.216277581837068 | 0.283029921910237 | 0.144747179527083 | OG0000071 | 1 |
| GO:0009889 | regulation of biosynthetic process | 1/6 | 520/12891 | 0.218933622465441 | 0.284748024310881 | 0.145625851559189 | OG0000047 | 1 |
| GO:0003824 | catalytic activity | 2/6 | 1967/12891 | 0.22958620242062 | 0.296782164104704 | 0.151780352049576 | OG0000071/OG0000180 | 2 |
| GO:0051171 | regulation of nitrogen compound metabolic process | 1/6 | 556/12891 | 0.232475075791311 | 0.298695248895502 | 0.152758741987273 | OG0000047 | 1 |
| GO:1901265 | nucleoside phosphate binding | 1/6 | 561/12891 | 0.234340263084725 | 0.298834990223159 | 0.152830208505389 | OG0000071 | 1 |
| GO:0010033 | response to organic substance | 1/6 | 564/12891 | 0.23545756127276 | 0.298834990223159 | 0.152830208505389 | OG0000180 | 1 |
| GO:0009791 | post-embryonic development | 1/6 | 570/12891 | 0.237688081902799 | 0.298834990223159 | 0.152830208505389 | OG0000047 | 1 |
| GO:0006807 | nitrogen compound metabolic process | 2/6 | 2014/12891 | 0.238222232772236 | 0.298834990223159 | 0.152830208505389 | OG0000047/OG0000071 | 2 |
| GO:0043168 | anion binding | 1/6 | 581/12891 | 0.24176328607871 | 0.301493039109921 | 0.154189588025431 | OG0000071 | 1 |
| GO:0000166 | nucleotide binding | 1/6 | 600/12891 | 0.248759513359304 | 0.305736864146508 | 0.156359965278502 | OG0000071 | 1 |
| GO:0005737 | cytoplasm | 3/6 | 3820/12891 | 0.248886968716679 | 0.305736864146508 | 0.156359965278502 | OG0000047/OG0000071/OG0000180 | 3 |
| GO:0036094 | small molecule binding | 1/6 | 602/12891 | 0.249492818383707 | 0.305736864146508 | 0.156359965278502 | OG0000071 | 1 |
| GO:0005576 | extracellular region | 1/6 | 613/12891 | 0.253515342287197 | 0.30888076186716 | 0.157967817638121 | OG0000071 | 1 |
| GO:0031323 | regulation of cellular metabolic process | 1/6 | 646/12891 | 0.26547527644492 | 0.321604334893275 | 0.164474908113244 | OG0000047 | 1 |
| GO:0043169 | cation binding | 1/6 | 732/12891 | 0.295895583536339 | 0.356419680168772 | 0.182280174068438 | OG0000180 | 1 |
| GO:0019222 | regulation of metabolic process | 1/6 | 737/12891 | 0.29763139769499 | 0.356485063905864 | 0.182313612623158 | OG0000047 | 1 |
| GO:0012505 | endomembrane system | 1/6 | 751/12891 | 0.302472716848612 | 0.358644140798277 | 0.18341780785612 | OG0000180 | 1 |
| GO:0009059 | macromolecule biosynthetic process | 1/6 | 752/12891 | 0.302817458504205 | 0.358644140798277 | 0.18341780785612 | OG0000071 | 1 |
| GO:0071704 | organic substance metabolic process | 2/6 | 2368/12891 | 0.304615964656142 | 0.358769913928345 | 0.183482130757793 | OG0000047/OG0000071 | 2 |
| GO:0016740 | transferase activity | 1/6 | 812/12891 | 0.323243780425612 | 0.378605974863148 | 0.193626690222961 | OG0000071 | 1 |
| GO:0016070 | RNA metabolic process | 1/6 | 820/12891 | 0.325929206595656 | 0.379653801089446 | 0.194162569574046 | OG0000071 | 1 |
| GO:0007275 | multicellular organism development | 1/6 | 869/12891 | 0.342184424852943 | 0.396410371960786 | 0.202732216047472 | OG0000047 | 1 |
| GO:0046872 | metal ion binding | 1/6 | 882/12891 | 0.346441763154335 | 0.399161161895213 | 0.204139025199637 | OG0000180 | 1 |
| GO:0032991 | protein-containing complex | 1/6 | 894/12891 | 0.350351210018307 | 0.401483548777736 | 0.205326740437471 | OG0000047 | 1 |
| GO:0005886 | plasma membrane | 1/6 | 919/12891 | 0.358433327532682 | 0.406023212694225 | 0.207648415628129 | OG0000180 | 1 |
| GO:0043231 | intracellular membrane-bounded organelle | 3/6 | 4573/12891 | 0.362496157123656 | 0.406023212694225 | 0.207648415628129 | OG0000047/OG0000071/OG0000180 | 3 |
| GO:0043227 | membrane-bounded organelle | 3/6 | 4574/12891 | 0.362653551016438 | 0.406023212694225 | 0.207648415628129 | OG0000047/OG0000071/OG0000180 | 3 |
| GO:0032501 | multicellular organismal process | 1/6 | 940/12891 | 0.365157390015262 | 0.406023212694225 | 0.207648415628129 | OG0000047 | 1 |
| GO:0043229 | intracellular organelle | 3/6 | 4592/12891 | 0.365488681893086 | 0.406023212694225 | 0.207648415628129 | OG0000047/OG0000071/OG0000180 | 3 |
| GO:0043226 | organelle | 3/6 | 4594/12891 | 0.365803932191495 | 0.406023212694225 | 0.207648415628129 | OG0000047/OG0000071/OG0000180 | 3 |
| GO:0005634 | nucleus | 2/6 | 2719/12891 | 0.371471619513451 | 0.408079742670041 | 0.208700166310895 | OG0000071/OG0000180 | 2 |
| GO:0048856 | anatomical structure development | 1/6 | 960/12891 | 0.371506558185462 | 0.408079742670041 | 0.208700166310895 | OG0000047 | 1 |
| GO:0090304 | nucleic acid metabolic process | 1/6 | 970/12891 | 0.374661243433486 | 0.40942362684484 | 0.209387455635643 | OG0000071 | 1 |
| GO:0032502 | developmental process | 1/6 | 999/12891 | 0.383735305320484 | 0.417189152450988 | 0.213358901203832 | OG0000047 | 1 |
| GO:0071944 | cell periphery | 1/6 | 1011/12891 | 0.387457854702935 | 0.419087067331747 | 0.214329532945233 | OG0000180 | 1 |
| GO:0008150 | biological_process | 3/6 | 4787/12891 | 0.396419417698809 | 0.426603637320546 | 0.218173657616764 | OG0000047/OG0000071/OG0000180 | 3 |
| GO:0005622 | intracellular anatomical structure | 3/6 | 4835/12891 | 0.404083411570792 | 0.432654965924282 | 0.221268428451842 | OG0000047/OG0000071/OG0000180 | 3 |
| GO:0050794 | regulation of cellular process | 1/6 | 1077/12891 | 0.407598613754158 | 0.434225658873776 | 0.222071712333659 | OG0000047 | 1 |
| GO:0006139 | nucleobase-containing compound metabolic process | 1/6 | 1084/12891 | 0.409701989634108 | 0.434284109012154 | 0.222101604906911 | OG0000071 | 1 |
| GO:0005488 | binding | 2/6 | 2958/12891 | 0.41675103474642 | 0.439558305304682 | 0.224798934689088 | OG0000071/OG0000180 | 2 |
| GO:1901564 | organonitrogen compound metabolic process | 1/6 | 1216/12891 | 0.448216086157445 | 0.470405001313754 | 0.240574553799984 | OG0000047 | 1 |
| GO:0110165 | cellular anatomical entity | 3/6 | 5126/12891 | 0.450789109942744 | 0.470774834028876 | 0.240763693669187 | OG0000047/OG0000071/OG0000180 | 3 |
| GO:0050789 | regulation of biological process | 1/6 | 1255/12891 | 0.459185777108393 | 0.477193062485193 | 0.244046104448733 | OG0000047 | 1 |
| GO:0005575 | cellular_component | 3/6 | 5202/12891 | 0.463016155946998 | 0.478826463711041 | 0.244881458600979 | OG0000047/OG0000071/OG0000180 | 3 |
| GO:0005739 | mitochondrion | 1/6 | 1312/12891 | 0.474891079751276 | 0.488722858773157 | 0.249942673553303 | OG0000071 | 1 |
| GO:1901363 | heterocyclic compound binding | 1/6 | 1374/12891 | 0.491540643288642 | 0.500742537046952 | 0.256089778132254 | OG0000071 | 1 |
| GO:0097159 | organic cyclic compound binding | 1/6 | 1381/12891 | 0.493392470418319 | 0.500742537046952 | 0.256089778132254 | OG0000071 | 1 |
| GO:0065007 | biological regulation | 1/6 | 1382/12891 | 0.493656557749117 | 0.500742537046952 | 0.256089778132254 | OG0000047 | 1 |
| GO:0043170 | macromolecule metabolic process | 1/6 | 1830/12891 | 0.601009026886241 | 0.606732922380396 | 0.310295387314701 | OG0000071 | 1 |
| GO:0044238 | primary metabolic process | 1/6 | 2198/12891 | 0.67433293403938 | 0.677528824722031 | 0.346501831908487 | OG0000071 | 1 |
| GO:0003674 | molecular_function | 2/6 | 4855/12891 | 0.728648477124954 | 0.728648477124954 | 0.372645447586248 | OG0000071/OG0000180 | 2 |

**Table S4g. Related to Figure 3. Enriched Pfam domains in orthogroups contracted in Chara with respect to Mesostigma.**

| ID | Description | GeneRatio | BgRatio | pvalue | p.adjust | qvalue | geneID | Count |
|---|---|---|---|---|---|---|---|---|
| **PF00194** | Carb_anhydrase | 1/6 | 2/13933 | 0.000861110075639937 | 0.00559721549165959 | | OG0000180 | 1 |
| **PF08370** | PDR_assoc | 1/6 | 2/13933 | 0.000861110075639937 | 0.00559721549165959 | | OG0000029 | 1 |
| **PF03055** | RPE65 | 1/6 | 7/13933 | 0.00301118222901831 | 0.0111823843200325 | | OG0000214 | 1 |
| **PF04720** | PDDEXK_6 | 1/6 | 8/13933 | 0.00344073363693309 | 0.0111823843200325 | | OG0000071 | 1 |
| **PF00324** | AA_permease | 1/6 | 13/13933 | 0.00558617800019301 | 0.0121033856670849 | | OG0000363 | 1 |
| **PF13520** | AA_permease_2 | 1/6 | 13/13933 | 0.00558617800019301 | 0.0121033856670849 | | OG0000363 | 1 |
| **PF19055** | ABC2_membrane_7 | 1/6 | 19/13933 | 0.00815562919282087 | 0.0146426839763669 | | OG0000029 | 1 |
| **PF01061** | ABC2_membrane | 1/6 | 21/13933 | 0.00901088244699499 | 0.0146426839763669 | | OG0000029 | 1 |
| **PF13555** | AAA_29 | 1/6 | 26/13933 | 0.0111463272665585 | 0.0155992600341593 | | OG0000029 | 1 |
| **PF00504** | Chloroa_b-bind | 1/6 | 28/13933 | 0.0119994307955071 | 0.0155992600341593 | | OG0000047 | 1 |
| **PF13304** | AAA_21 | 1/6 | 44/13933 | 0.0188022058001617 | 0.0222207886729183 | | OG0000029 | 1 |
| **PF13191** | AAA_16 | 1/6 | 59/13933 | 0.0251443131639131 | 0.0272396725942392 | | OG0000029 | 1 |
| **PF00005** | ABC_tran | 1/6 | 82/13933 | 0.0348025091329028 | 0.0348025091329028 | | OG0000029 | 1 |
