## Supplemental Table S5 for "The *Draparnaldia* genome: alternative mechanisms for multicellularity and terrestrialization in green plants"

**Table S5d. Related to Figure 4. GO enrichment of downregulated genes in agar_culture_20_days_vs_liquid_culture**

| GOBPID | Pvalue | OddsRatio | ExpCount | Count | Size | Term | gene_id |
|---|---|---|---|---|---|---|---|
| GO:0009190 | 2.99065077626434E-07 | 5.29040404040404 | 4.63374125874126 | 18 | 62 | cyclic nucleotide biosynthetic process | DRAP1A001130,DRAP1A002751,DRAP1A003147,DRAP1A003816,DRAP1A005483,DRAP1A005540,DRAP1A005937,DRAP1A008618,DRAP1A009488,DRAP1A009818,DRAP1A010650,DRAP1A012845,DRAP1A012906,DRAP1A015127,DRAP1A016776,[illegible] |
| GO:0009187 | 2.99065077626434E-07 | 5.29040404040404 | 4.63374125874126 | 18 | 62 | cyclic nucleotide metabolic process | DRAP1A001130,DRAP1A002751,DRAP1A003147,DRAP1A003816,DRAP1A005483,DRAP1A005540,DRAP1A005937,DRAP1A008618,DRAP1A009488,DRAP1A009818,DRAP1A010650,DRAP1A012845,DRAP1A012906,DRAP1A015127,DRAP1A016776,[illegible] |
| GO:0006793 | 4.58448819840297E-07 | 1.97182638293674 | 54.5585664335664 | 89 | 730 | phosphorus metabolic process | DRAP1A000041,DRAP1A000862,DRAP1A001130,DRAP1A001440,DRAP1A001868,DRAP1A001925,DRAP1A002166,DRAP1A002405,DRAP1A002722,DRAP1A002751,DRAP1A002971,DRAP1A003147,DRAP1A003575,DRAP1A003816,DRAP1A003968,[illegible] |
| GO:0006796 | 4.58448819840297E-07 | 1.97182638293674 | 54.5585664335664 | 89 | 730 | phosphate-containing compound metabolic process | DRAP1A000041,DRAP1A000862,DRAP1A001130,DRAP1A001440,DRAP1A001868,DRAP1A001925,DRAP1A002166,DRAP1A002405,DRAP1A002722,DRAP1A002751,DRAP1A002971,DRAP1A003147,DRAP1A003575,DRAP1A003816,DRAP1A003968,[illegible] |
| GO:0006928 | 3.34105580256566E-06 | 4.30041152263374 | 5.38111888111888 | 18 | 72 | movement of cell or subcellular component | DRAP1A000146,DRAP1A002648,DRAP1A003653,DRAP1A004424,DRAP1A005983,DRAP1A009351,DRAP1A011746,DRAP1A014583,DRAP1A018632,DRAP1A018713,DRAP1A019768,DRAP1A020259,DRAP1A021397,DRAP1A021578,DRAP1A023123,[illegible] |
| GO:0009165 | 5.95286690828153E-06 | 2.93126114649682 | 11.4348776223776 | 28 | 153 | nucleotide biosynthetic process | DRAP1A001130,DRAP1A001868,DRAP1A002166,DRAP1A002751,DRAP1A003147,DRAP1A003816,DRAP1A005483,DRAP1A005540,DRAP1A005937,DRAP1A007840,DRAP1A008618,DRAP1A009488,DRAP1A009818,DRAP1A010650,DRAP1A012845,[illegible] |
| GO:1901293 | 5.95286690828153E-06 | 2.93126114649682 | 11.4348776223776 | 28 | 153 | nucleoside phosphate biosynthetic process | DRAP1A001130,DRAP1A001868,DRAP1A002166,DRAP1A002751,DRAP1A003147,DRAP1A003816,DRAP1A005483,DRAP1A005540,DRAP1A005937,DRAP1A007840,DRAP1A008618,DRAP1A009488,DRAP1A009818,DRAP1A010650,DRAP1A012845,[illegible] |
| GO:0035556 | 2.59735477524069E-05 | 3.09524456521739 | 8.52010489510489 | 22 | 114 | intracellular signal transduction | DRAP1A001130,DRAP1A002751,DRAP1A003147,DRAP1A003816,DRAP1A005483,DRAP1A005540,DRAP1A005937,DRAP1A007926,DRAP1A008618,DRAP1A009488,DRAP1A009818,DRAP1A010650,DRAP1A011200,DRAP1A012845,DRAP1A012906,[illegible] |
| GO:0006468 | 8.50945653233955E-05 | 1.94246974719438 | 29.3719405594406 | 50 | 393 | protein phosphorylation | DRAP1A000041,DRAP1A000862,DRAP1A001440,DRAP1A002405,DRAP1A002722,DRAP1A002971,DRAP1A003968,DRAP1A003993,DRAP1A004431,DRAP1A005626,DRAP1A005943,DRAP1A006630,DRAP1A006860,DRAP1A006932,DRAP1A007099,[illegible] |
| GO:0007018 | 0.000106812062851658 | 3.61866020425827 | 5.08216783216783 | 15 | 68 | microtubule-based movement | DRAP1A000146,DRAP1A002648,DRAP1A003653,DRAP1A004424,DRAP1A005983,DRAP1A009351,DRAP1A011746,DRAP1A014583,DRAP1A018632,DRAP1A018713,DRAP1A020259,DRAP1A021397,DRAP1A021578,DRAP1A023123,DRAP1A025294 |
| GO:0090407 | 0.000191253525046528 | 2.33104687244815 | 13.7517482517483 | 28 | 184 | organophosphate biosynthetic process | DRAP1A001130,DRAP1A001868,DRAP1A002166,DRAP1A002751,DRAP1A003147,DRAP1A003816,DRAP1A005483,DRAP1A005540,DRAP1A005937,DRAP1A007840,DRAP1A008618,DRAP1A009488,DRAP1A009818,DRAP1A010650,DRAP1A012845,[illegible] |
| GO:0009117 | 0.0002650317108359 | 2.14965081476555 | 16.8907342657343 | 32 | 226 | nucleotide metabolic process | DRAP1A001130,DRAP1A001868,DRAP1A002166,DRAP1A002751,DRAP1A003147,DRAP1A003816,DRAP1A005483,DRAP1A005540,DRAP1A005937,DRAP1A007840,DRAP1A008618,DRAP1A009488,DRAP1A009772,DRAP1A009818,DRAP1A010544,[illegible] |
| GO:0006753 | 0.000312088916163888 | 2.12666227781435 | 17.0402097902098 | 32 | 228 | nucleoside phosphate metabolic process | DRAP1A001130,DRAP1A001868,DRAP1A002166,DRAP1A002751,DRAP1A003147,DRAP1A003816,DRAP1A005483,DRAP1A005540,DRAP1A005937,DRAP1A007840,DRAP1A008618,DRAP1A009488,DRAP1A009772,DRAP1A009818,DRAP1A010544,[illegible] |
| GO:0016310 | 0.000567816757124657 | 1.71441461764042 | 36.6215034965035 | 56 | 490 | phosphorylation | DRAP1A000041,DRAP1A000862,DRAP1A001440,DRAP1A001868,DRAP1A002166,DRAP1A002405,DRAP1A002722,DRAP1A002971,DRAP1A003968,DRAP1A003993,DRAP1A004431,DRAP1A005626,DRAP1A005943,DRAP1A006630,DRAP1A006860,[illegible] |
| GO:0007017 | 0.000799902367996201 | 2.68189933523267 | 7.3243006993007 | 17 | 98 | microtubule-based process | DRAP1A000146,DRAP1A001512,DRAP1A002648,DRAP1A003653,DRAP1A004424,DRAP1A005983,DRAP1A009351,DRAP1A011746,DRAP1A013838,DRAP1A014583,DRAP1A018632,DRAP1A018713,DRAP1A020259,DRAP1A021397,DRAP1A021578,[illegible] |
| GO:0051674 | 0.00089520988436183 | 16.6903353057199 | 0.523164335664336 | 4 | 7 | localization of cell | DRAP1A019768,DRAP1A021397,DRAP1A024094,DRAP1A024842 |
| GO:0040011 | 0.00089520988436183 | 16.6903353057199 | 0.523164335664336 | 4 | 7 | locomotion | DRAP1A019768,DRAP1A021397,DRAP1A024094,DRAP1A024842 |
| GO:0048870 | 0.00089520988436183 | 16.6903353057199 | 0.523164335664336 | 4 | 7 | cell motility | DRAP1A019768,DRAP1A021397,DRAP1A024094,DRAP1A024842 |
| GO:0019637 | 0.00110015014172477 | 1.87482352941176 | 21.375 | 36 | 286 | organophosphate metabolic process | DRAP1A001130,DRAP1A001868,DRAP1A002166,DRAP1A002751,DRAP1A003147,DRAP1A003575,DRAP1A003816,DRAP1A005483,DRAP1A005540,DRAP1A005937,DRAP1A007840,DRAP1A008618,DRAP1A009488,DRAP1A009772,DRAP1A009818,[illegible] |
| GO:0055086 | 0.00196762377950427 | 1.78810408921933 | 22.8697552447552 | 37 | 306 | nucleobase-containing small molecule metabolic process | DRAP1A001130,DRAP1A001868,DRAP1A002166,DRAP1A002751,DRAP1A003147,DRAP1A003816,DRAP1A004743,DRAP1A005483,DRAP1A005540,DRAP1A005937,DRAP1A007840,DRAP1A008618,DRAP1A009488,DRAP1A009772,DRAP1A009818,[illegible] |
| GO:0030030 | 0.00262753502546412 | 5.38265306122449 | 1.49475524475524 | 6 | 20 | cell projection organization | DRAP1A005983,DRAP1A009351,DRAP1A011746,DRAP1A018713,DRAP1A021397,DRAP1A021578 |
| GO:0036211 | 0.00573640538236943 | 1.48145161290323 | 46.5616258741259 | 63 | 623 | protein modification process | DRAP1A000041,DRAP1A000862,DRAP1A001440,DRAP1A001925,DRAP1A002405,DRAP1A002689,DRAP1A002722,DRAP1A002971,DRAP1A003968,DRAP1A003993,DRAP1A004431,DRAP1A005217,DRAP1A005626,DRAP1A005943,DRAP1A006630,[illegible] |
| GO:0006464 | 0.00573640538236943 | 1.48145161290323 | 46.5616258741259 | 63 | 623 | cellular protein modification process | DRAP1A000041,DRAP1A000862,DRAP1A001440,DRAP1A001925,DRAP1A002405,DRAP1A002689,DRAP1A002722,DRAP1A002971,DRAP1A003968,DRAP1A003993,DRAP1A004431,DRAP1A005217,DRAP1A005626,DRAP1A005943,DRAP1A006630,[illegible] |
| GO:0001539 | 0.00698032901998358 | 12.4808259587021 | 0.448426573426573 | 3 | 6 | cilium or flagellum-dependent cell motility | DRAP1A019768,DRAP1A021397,DRAP1A024094 |
| GO:0072330 | 0.0125117198507759 | 2.06521739130435 | 7.99694055944056 | 15 | 107 | monocarboxylic acid biosynthetic process | DRAP1A001868,DRAP1A002166,DRAP1A004184,DRAP1A008106,DRAP1A009982,DRAP1A011537,DRAP1A012810,DRAP1A015921,DRAP1A018106,DRAP1A019891,DRAP1A021520,DRAP1A023576,DRAP1A024440,DRAP1A025410,DRAP1A025880 |
| GO:0044267 | 0.0138064516145845 | 1.35672048527426 | 66.6660839160839 | 83 | 892 | cellular protein metabolic process | DRAP1A000041,DRAP1A000862,DRAP1A001027,DRAP1A001440,DRAP1A001925,DRAP1A002405,DRAP1A002689,DRAP1A002722,DRAP1A002971,DRAP1A003584,DRAP1A003911,DRAP1A003968,DRAP1A003993,DRAP1A004431,DRAP1A005217,[illegible] |
| GO:0010207 | 0.0158836901484403 | 24.9 | 0.224213286713287 | 2 | 3 | photosystem II assembly | DRAP1A014165,DRAP1A014220 |
| GO:0042026 | 0.0158836901484403 | 24.9 | 0.224213286713287 | 2 | 3 | protein refolding | DRAP1A015883,DRAP1A022645 |
| GO:0006457 | 0.0211010648142782 | 2.1020202020202 | 6.27797202797203 | 12 | 84 | protein folding | DRAP1A001189,DRAP1A002689,DRAP1A003499,DRAP1A010884,DRAP1A014402,DRAP1A014481,DRAP1A015883,DRAP1A019030,DRAP1A022411,DRAP1A022645,DRAP1A023725,DRAP1A025802 |
| GO:0006633 | 0.0215167465370674 | 2.57638569015815 | 3.51267482517482 | 8 | 47 | fatty acid biosynthetic process | DRAP1A008106,DRAP1A009982,DRAP1A012810,DRAP1A021520,DRAP1A023576,DRAP1A024440,DRAP1A025410,DRAP1A025880 |
| GO:0120036 | 0.0216176711905691 | 4.54330285099516 | 1.12106643356643 | 4 | 15 | plasma membrane bounded cell projection organization | DRAP1A009351,DRAP1A018713,DRAP1A021397,DRAP1A021578 |
| GO:0044782 | 0.0216176711905691 | 4.54330285099516 | 1.12106643356643 | 4 | 15 | cilium organization | DRAP1A009351,DRAP1A018713,DRAP1A021397,DRAP1A021578 |
| GO:0006979 | 0.0234898281708787 | 2.74384328358209 | 2.91477272727273 | 7 | 39 | response to oxidative stress | DRAP1A008214,DRAP1A008303,DRAP1A015110,DRAP1A015111,DRAP1A024311,DRAP1A026260,DRAP1A027338 |
| GO:0006487 | 0.0247612456768398 | 6.23598820058997 | 0.67263986013986 | 3 | 9 | protein N-linked glycosylation | DRAP1A013663,DRAP1A020204,DRAP1A021523 |
| GO:0023052 | 0.0257643005971604 | 1.4788838612368 | 26.0087412587413 | 36 | 348 | signaling | DRAP1A001130,DRAP1A002147,DRAP1A002751,DRAP1A003147,DRAP1A003816,DRAP1A005483,DRAP1A005540,DRAP1A005937,DRAP1A005943,DRAP1A007926,DRAP1A008618,DRAP1A009488,DRAP1A009818,DRAP1A010650,DRAP1A011026,[illegible] |
| GO:0007165 | 0.0257643005971604 | 1.4788838612368 | 26.0087412587413 | 36 | 348 | signal transduction | DRAP1A001130,DRAP1A002147,DRAP1A002751,DRAP1A003147,DRAP1A003816,DRAP1A005483,DRAP1A005540,DRAP1A005937,DRAP1A005943,DRAP1A007926,DRAP1A008618,DRAP1A009488,DRAP1A009818,DRAP1A010650,DRAP1A011026,[illegible] |
| GO:0007154 | 0.0257643005971604 | 1.4788838612368 | 26.0087412587413 | 36 | 348 | cell communication | DRAP1A001130,DRAP1A002147,DRAP1A002751,DRAP1A003147,DRAP1A003816,DRAP1A005483,DRAP1A005540,DRAP1A005937,DRAP1A005943,DRAP1A007926,DRAP1A008618,DRAP1A009488,DRAP1A009818,DRAP1A010650,DRAP1A011026,[illegible] |
| GO:0042073 | 0.0302031417554893 | 12.4470588235294 | 0.298951048951049 | 2 | 4 | intraciliary transport | DRAP1A009351,DRAP1A018713 |
| GO:0030705 | 0.0302031417554893 | 12.4470588235294 | 0.298951048951049 | 2 | 4 | cytoskeleton-dependent intracellular transport | DRAP1A009351,DRAP1A018713 |
| GO:0099111 | 0.0302031417554893 | 12.4470588235294 | 0.298951048951049 | 2 | 4 | microtubule-based transport | DRAP1A009351,DRAP1A018713 |
| GO:0010970 | 0.0302031417554893 | 12.4470588235294 | 0.298951048951049 | 2 | 4 | transport along microtubule | DRAP1A009351,DRAP1A018713 |
| GO:0008610 | 0.0315493457965307 | 1.90584576235244 | 7.39903846153846 | 13 | 99 | lipid biosynthetic process | DRAP1A001796,DRAP1A008106,DRAP1A009982,DRAP1A012727,DRAP1A012810,DRAP1A016943,DRAP1A018969,DRAP1A019286,DRAP1A021520,DRAP1A023576,DRAP1A024440,DRAP1A025410,DRAP1A025880 |
| GO:0043412 | 0.0428632828801225 | 1.30238450748835 | 52.3164335664336 | 64 | 700 | macromolecule modification | DRAP1A000041,DRAP1A000862,DRAP1A001440,DRAP1A001925,DRAP1A002405,DRAP1A002689,DRAP1A002722,DRAP1A002971,DRAP1A003968,DRAP1A003993,DRAP1A004431,DRAP1A005217,DRAP1A005626,DRAP1A005943,DRAP1A006630,[illegible] |
| GO:0034654 | 0.0459670292508428 | 1.3682951653944 | 32.5109265734266 | 42 | 435 | nucleobase-containing compound biosynthetic process | DRAP1A001130,DRAP1A001412,DRAP1A001868,DRAP1A002166,DRAP1A002373,DRAP1A002751,DRAP1A003147,DRAP1A003816,DRAP1A005051,DRAP1A005483,DRAP1A005540,DRAP1A005937,DRAP1A006500,DRAP1A007234,DRAP1A007840,[illegible] |
| GO:0006813 | 0.0467047660073125 | 2.840571891017 | 2.01791958041958 | 5 | 27 | potassium ion transport | DRAP1A012361,DRAP1A013272,DRAP1A015169,DRAP1A016488,DRAP1A019677 |
| GO:0060285 | 0.0478759090932062 | 8.29607843137255 | 0.373688811188811 | 2 | 5 | cilium-dependent cell motility | DRAP1A021397,DRAP1A024094 |
| GO:0006631 | 0.049747561119564 | 2.13377500318512 | 4.11057692307692 | 8 | 55 | fatty acid metabolic process | DRAP1A008106,DRAP1A009982,DRAP1A012810,DRAP1A021520,DRAP1A023576,DRAP1A024440,DRAP1A025410,DRAP1A025880 |

**Table S5e. Related to Figure 4. GO enrichment of upregulated genes in agar_culture_20_days_vs_liquid_culture**

| GOBPID | Pvalue | OddsRatio | ExpCount | Count | Size | Term | gene_id |
|---|---|---|---|---|---|---|---|
| **GO:0055085** | 3.78541851148926E-07 | 3.27043031977892 | 11.1483828671329 | 30 | 285 | transmembrane transport | DRAP1A000051,DRAP1A002066,DRAP1A003685,DRAP1A005178,DRAP1A006013,DRAP1A006650,DRAP1A007493,DRAP1A007590,DRAP1A008450,DRAP1A008968,DRAP1A009355,DRAP1A009584,DRAP1A010216,DRAP1A012007,DRAP1A012102,DRAP1A014767,DRAP1A014955,DRAP1A015480,DRAP1A015762,DRAP1A016049,DRAP1A017258,DRAP1A017570,DRAP1A017820,DRAP1A017841,DRAP1A018486,DRAP1A020079,DRAP1A021005,DRAP1A024478,DRAP1A025638,DRAP1A026322 |
| **GO:0006811** | 3.45621108441602E-05 | 3.16636717580114 | 7.35402097902098 | 20 | 188 | ion transport | DRAP1A000051,DRAP1A001628,DRAP1A002066,DRAP1A005178,DRAP1A006013,DRAP1A007493,DRAP1A008450,DRAP1A008968,DRAP1A009584,DRAP1A010216,DRAP1A011859,DRAP1A012007,DRAP1A013589,DRAP1A014767,DRAP1A015738,DRAP1A016049,DRAP1A017258,DRAP1A017841,DRAP1A020079,DRAP1A024478 |
| **GO:0015696** | 8.63377237512236E-05 | 15.7650862068966 | 0.508522727272727 | 5 | 13 | ammonium transport | DRAP1A002066,DRAP1A007493,DRAP1A014767,DRAP1A015738,DRAP1A020079 |
| **GO:0072488** | 0.000244556833289985 | 20.0777142857143 | 0.352054195804196 | 4 | 9 | ammonium transmembrane transport | DRAP1A002066,DRAP1A007493,DRAP1A014767,DRAP1A020079 |
| **GO:0006812** | 0.000270646505751041 | 3.3378926883514 | 4.81140734265734 | 14 | 123 | cation transport | DRAP1A001628,DRAP1A002066,DRAP1A006013,DRAP1A007493,DRAP1A008968,DRAP1A009584,DRAP1A010216,DRAP1A013589,DRAP1A014767,DRAP1A015738,DRAP1A016049,DRAP1A017841,DRAP1A020079,DRAP1A024478 |
| **GO:0006810** | 0.000392596537665871 | 1.99196037031022 | 21.9838286713287 | 38 | 562 | transport | DRAP1A000051,DRAP1A001628,DRAP1A002066,DRAP1A003685,DRAP1A005178,DRAP1A006013,DRAP1A006259,DRAP1A006650,DRAP1A007493,DRAP1A007590,DRAP1A008450,DRAP1A008968,DRAP1A009355,DRAP1A009584,DRAP1A010216,DRAP1A011859,DRAP1A012007,DRAP1A012023,DRAP1A012102,DRAP1A013589,DRAP1A014767,DRAP1A014955,DRAP1A015480,DRAP1A015738,DRAP1A015762,DRAP1A016049,DRAP1A016558,DRAP1A017258,DRAP1A017570,DRAP1A017820,DRAP1A017841,DRAP1A018486,DRAP1A020079,DRAP1A021005,DRAP1A021431,DRAP1A024478,DRAP1A025638,DRAP1A026322 |
| **GO:0051234** | 0.00046938647376411 | 1.97058547507005 | 22.1794143356643 | 38 | 567 | establishment of localization | DRAP1A000051,DRAP1A001628,DRAP1A002066,DRAP1A003685,DRAP1A005178,DRAP1A006013,DRAP1A006259,DRAP1A006650,DRAP1A007493,DRAP1A007590,DRAP1A008450,DRAP1A008968,DRAP1A009355,DRAP1A009584,DRAP1A010216,DRAP1A011859,DRAP1A012007,DRAP1A012023,DRAP1A012102,DRAP1A013589,DRAP1A014767,DRAP1A014955,DRAP1A015480,DRAP1A015738,DRAP1A015762,DRAP1A016049,DRAP1A016558,DRAP1A017258,DRAP1A017570,DRAP1A017820,DRAP1A017841,DRAP1A018486,DRAP1A020079,DRAP1A021005,DRAP1A021431,DRAP1A024478,DRAP1A025638,DRAP1A026322 |
| **GO:0051179** | 0.000687415280738596 | 1.92495403204623 | 22.6097027972028 | 38 | 578 | localization | DRAP1A000051,DRAP1A001628,DRAP1A002066,DRAP1A003685,DRAP1A005178,DRAP1A006013,DRAP1A006259,DRAP1A006650,DRAP1A007493,DRAP1A007590,DRAP1A008450,DRAP1A008968,DRAP1A009355,DRAP1A009584,DRAP1A010216,DRAP1A011859,DRAP1A012007,DRAP1A012023,DRAP1A012102,DRAP1A013589,DRAP1A014767,DRAP1A014955,DRAP1A015480,DRAP1A015738,DRAP1A015762,DRAP1A016049,DRAP1A016558,DRAP1A017258,DRAP1A017570,DRAP1A017820,DRAP1A017841,DRAP1A018486,DRAP1A020079,DRAP1A021005,DRAP1A021431,DRAP1A024478,DRAP1A025638,DRAP1A026322 |
| **GO:0034220** | 0.000705339364709224 | 4.73711410308718 | 1.99497377622378 | 8 | 51 | ion transmembrane transport | DRAP1A000051,DRAP1A002066,DRAP1A007493,DRAP1A008450,DRAP1A010216,DRAP1A014767,DRAP1A016049,DRAP1A020079 |
| **GO:0006885** | 0.00122660445902013 | 11.144126984127 | 0.508522727272727 | 4 | 13 | regulation of pH | DRAP1A008968,DRAP1A009584,DRAP1A017841,DRAP1A024478 |
| **GO:0055067** | 0.00122660445902013 | 11.144126984127 | 0.508522727272727 | 4 | 13 | monovalent inorganic cation homeostasis | DRAP1A008968,DRAP1A009584,DRAP1A017841,DRAP1A024478 |
| **GO:0098655** | 0.00278565638714684 | 4.88457952638449 | 1.44733391608392 | 6 | 37 | cation transmembrane transport | DRAP1A002066,DRAP1A007493,DRAP1A010216,DRAP1A014767,DRAP1A016049,DRAP1A020079 |
| **GO:0050801** | 0.00361103467062959 | 7.70813186813187 | 0.664991258741259 | 4 | 17 | ion homeostasis | DRAP1A008968,DRAP1A009584,DRAP1A017841,DRAP1A024478 |
| **GO:0055080** | 0.00361103467062959 | 7.70813186813187 | 0.664991258741259 | 4 | 17 | cation homeostasis | DRAP1A008968,DRAP1A009584,DRAP1A017841,DRAP1A024478 |
| **GO:0098771** | 0.00361103467062959 | 7.70813186813187 | 0.664991258741259 | 4 | 17 | inorganic ion homeostasis | DRAP1A008968,DRAP1A009584,DRAP1A017841,DRAP1A024478 |
| **GO:0071804** | 0.00444801451424775 | 49.6723163841808 | 0.117351398601399 | 2 | 3 | cellular potassium ion transport | DRAP1A010216,DRAP1A016049 |
| **GO:0071805** | 0.00444801451424775 | 49.6723163841808 | 0.117351398601399 | 2 | 3 | potassium ion transmembrane transport | DRAP1A010216,DRAP1A016049 |
| **GO:0048878** | 0.004502803099129 | 7.15591836734694 | 0.704108391608392 | 4 | 18 | chemical homeostasis | DRAP1A008968,DRAP1A009584,DRAP1A017841,DRAP1A024478 |
| **GO:0006814** | 0.0125981736280201 | 7.47784090909091 | 0.508522727272727 | 3 | 13 | sodium ion transport | DRAP1A008968,DRAP1A009584,DRAP1A024478 |
| **GO:0006769** | 0.0140751108611035 | 16.5499058380414 | 0.195585664335664 | 2 | 5 | nicotinamide metabolic process | DRAP1A008965,DRAP1A015971 |
| **GO:0009820** | 0.0140751108611035 | 16.5499058380414 | 0.195585664335664 | 2 | 5 | alkaloid metabolic process | DRAP1A008965,DRAP1A015971 |
| **GO:0015995** | 0.0155798886447198 | 6.79648760330578 | 0.54763986013986 | 3 | 14 | chlorophyll biosynthetic process | DRAP1A005312,DRAP1A006494,DRAP1A013755 |
| **GO:0055114** | 0.0157307356183231 | 1.58755374462554 | 22.7270541958042 | 33 | 581 | oxidation-reduction process | DRAP1A000078,DRAP1A000614,DRAP1A001208,DRAP1A002501,DRAP1A002509,DRAP1A004187,DRAP1A004778,DRAP1A006494,DRAP1A006790,DRAP1A008029,DRAP1A008898,DRAP1A009432,DRAP1A009675,DRAP1A009735,DRAP1A009999,DRAP1A011449,DRAP1A012351,DRAP1A013016,DRAP1A013755,DRAP1A015623,DRAP1A015913,DRAP1A016462,DRAP1A016513,DRAP1A017677,DRAP1A017873,DRAP1A018105,DRAP1A019420,DRAP1A020051,DRAP1A020809,DRAP1A024003,DRAP1A024615,DRAP1A026339,DRAP1A026559 |
| **GO:0030001** | 0.0200761365567996 | 2.75535973837209 | 2.77731643356643 | 7 | 71 | metal ion transport | DRAP1A006013,DRAP1A008968,DRAP1A009584,DRAP1A010216,DRAP1A013589,DRAP1A016049,DRAP1A024478 |
| **GO:0015672** | 0.0330295890442069 | 2.68848059454996 | 2.42526223776224 | 6 | 62 | monovalent inorganic cation transport | DRAP1A006013,DRAP1A008968,DRAP1A009584,DRAP1A010216,DRAP1A016049,DRAP1A024478 |
| **GO:0006741** | 0.0391171328671333 | Inf | 0.0391171328671329 | 1 | 1 | NADP biosynthetic process | DRAP1A015971 |
| **GO:0015707** | 0.0391171328671333 | Inf | 0.0391171328671329 | 1 | 1 | nitrite transport | DRAP1A012007 |
| **GO:0046497** | 0.0457098270545265 | 7.0863599677159 | 0.352054195804196 | 2 | 9 | nicotinate nucleotide metabolic process | DRAP1A008965,DRAP1A015971 |
| **GO:0042546** | 0.0457098270545265 | 7.0863599677159 | 0.352054195804196 | 2 | 9 | cell wall biogenesis | DRAP1A009717,DRAP1A014902 |

**Table S5f. Related to Figure 4. GO enrichment of downregulated genes in agar_culture_40_days_vs_agar_culture_20_days**

| GOBPID | Pvalue | OddsRatio | ExpCount | Count | Size | Term | gene_id |
|---|---|---|---|---|---|---|---|
| GO:0006260 | 2.2774540935794E-1 | 55.4188129899216 | 0.380244755244755 | 11 | 58 | DNA replication | DRAP1A003552,DRAP1A007686,DRAP1A008066,DRAP1A008624,DRAP1A009468,DRAP1A012267,DRAP1A017508,DRAP1A018425,DRAP1A020630,DRAP1A024055,DRAP1A027211 |
| GO:0006261 | 2.697246074626E-1 | 153.425120772947 | 0.104895104895105 | 7 | 16 | DNA-dependent DNA replication | DRAP1A003552,DRAP1A007686,DRAP1A008066,DRAP1A008624,DRAP1A009468,DRAP1A017508,DRAP1A020630 |
| GO:0006270 | 1.77739947426908E | 454.4 | 0.045891608391608 | 5 | 7 | DNA replication initiation | DRAP1A003552,DRAP1A007686,DRAP1A009468,DRAP1A017508,DRAP1A020630 |
| GO:0006259 | 1.74311493883852E | 9.71232876712329 | 1.99300699300699 | 12 | 304 | DNA metabolic process | DRAP1A003552,DRAP1A007686,DRAP1A008066,DRAP1A008624,DRAP1A009468,DRAP1A012267,DRAP1A017508,DRAP1A018425,DRAP1A020630,DRAP1A024055,DRAP1A026863,DRAP1A027211 |
| GO:0015696 | 6.95996027466159E | 50.4 | 0.085227272727272 | 3 | 13 | ammonium transport | DRAP1A007493,DRAP1A015738,DRAP1A020079 |
| GO:0034645 | 0.000247045717442 | 4.40133779264214 | 3.99912587412587 | 12 | 610 | cellular macromolecule biosynthetic process | DRAP1A000027,DRAP1A003552,DRAP1A007686,DRAP1A008066,DRAP1A008624,DRAP1A009468,DRAP1A012267,DRAP1A017508,DRAP1A018425,DRAP1A020630,DRAP1A024055,DRAP1A027211 |
| GO:0009059 | 0.000250970310374 | 4.39287701725097 | 4.00568181818182 | 12 | 611 | macromolecule biosynthetic process | DRAP1A000027,DRAP1A003552,DRAP1A007686,DRAP1A008066,DRAP1A008624,DRAP1A009468,DRAP1A012267,DRAP1A017508,DRAP1A018425,DRAP1A020630,DRAP1A024055,DRAP1A027211 |
| GO:0072488 | 0.001453870493842 | 46.3163265306122 | 0.059003496503496 | 2 | 9 | ammonium transmembrane transport | DRAP1A007493,DRAP1A020079 |
| GO:0090304 | 0.003620457708686 | 3.09346567411084 | 5.36276223776224 | 12 | 818 | nucleic acid metabolic process | DRAP1A003552,DRAP1A007686,DRAP1A008066,DRAP1A008624,DRAP1A009468,DRAP1A012267,DRAP1A017508,DRAP1A018425,DRAP1A020630,DRAP1A024055,DRAP1A026863,DRAP1A027211 |
| GO:0016567 | 0.003823716299255 | 10.8695652173913 | 0.321241258741259 | 3 | 49 | protein ubiquitination | DRAP1A002095,DRAP1A015729,DRAP1A018905 |
| GO:0032446 | 0.004776824010521 | 9.99111111111111 | 0.347465034965035 | 3 | 53 | protein modification by small protein conjugation | DRAP1A002095,DRAP1A015729,DRAP1A018905 |
| GO:0048478 | 0.006555944055944 | Inf | 0.006555944055944 | 1 | 1 | replication fork protection | DRAP1A008066 |
| GO:0090329 | 0.006555944055944 | Inf | 0.006555944055944 | 1 | 1 | regulation of DNA-dependent DNA replication | DRAP1A008066 |
| GO:2000104 | 0.006555944055944 | Inf | 0.006555944055944 | 1 | 1 | negative regulation of DNA-dependent DNA replication | DRAP1A008066 |
| GO:0008156 | 0.006555944055944 | Inf | 0.006555944055944 | 1 | 1 | negative regulation of DNA replication | DRAP1A008066 |
| GO:0044260 | 0.007249191390045 | 2.62403337403337 | 10.0240384615385 | 17 | 1529 | cellular macromolecule metabolic process | DRAP1A000027,DRAP1A002095,DRAP1A003159,DRAP1A003552,DRAP1A007686,DRAP1A008066,DRAP1A008624,DRAP1A009468,DRAP1A012267,DRAP1A015729,DRAP1A017508,DRAP1A018425,DRAP1A018905,DRAP1A020630,DRAP1A024055,DRAP1A026863,DRAP1A027211 |
| GO:0055085 | 0.009225406779078 | 3.82347670250896 | 1.86844405594406 | 6 | 285 | transmembrane transport | DRAP1A003685,DRAP1A007493,DRAP1A009355,DRAP1A015480,DRAP1A020079,DRAP1A025638 |
| GO:0044249 | 0.010519169533096 | 2.58115835362056 | 6.89685314685315 | 13 | 1052 | cellular biosynthetic process | DRAP1A000027,DRAP1A003552,DRAP1A006494,DRAP1A007686,DRAP1A008066,DRAP1A008624,DRAP1A009468,DRAP1A012267,DRAP1A017508,DRAP1A018425,DRAP1A020630,DRAP1A024055,DRAP1A027211 |
| GO:1901576 | 0.011409544813661 | 2.54926260303931 | 6.96241258741259 | 13 | 1062 | organic substance biosynthetic process | DRAP1A000027,DRAP1A003552,DRAP1A006494,DRAP1A007686,DRAP1A008066,DRAP1A008624,DRAP1A009468,DRAP1A012267,DRAP1A017508,DRAP1A018425,DRAP1A020630,DRAP1A024055,DRAP1A027211 |
| GO:0009186 | 0.013070331308036 | 156.724137931034 | 0.013111888111888 | 1 | 2 | deoxyribonucleoside diphosphate metabolic process | DRAP1A018425 |
| GO:0006269 | 0.013070331308036 | 156.724137931034 | 0.013111888111888 | 1 | 2 | DNA replication, synthesis of RNA primer | DRAP1A008624 |
| GO:0070647 | 0.014876949795244 | 6.44877344877345 | 0.524475524475524 | 3 | 80 | protein modification by small protein conjugation or removal | DRAP1A002095,DRAP1A015729,DRAP1A018905 |
| GO:0009058 | 0.015598689041122 | 2.42753740614703 | 7.22465034965035 | 13 | 1102 | biosynthetic process | DRAP1A000027,DRAP1A003552,DRAP1A006494,DRAP1A007686,DRAP1A008066,DRAP1A008624,DRAP1A009468,DRAP1A012267,DRAP1A017508,DRAP1A018425,DRAP1A020630,DRAP1A024055,DRAP1A027211 |
| GO:0000724 | 0.019543416148604 | 78.3448275862069 | 0.019667832167832 | 1 | 3 | double-strand break repair via homologous recombination | DRAP1A026863 |
| GO:0000725 | 0.019543416148604 | 78.3448275862069 | 0.019667832167832 | 1 | 3 | recombinational repair | DRAP1A026863 |
| GO:0006275 | 0.019543416148604 | 78.3448275862069 | 0.019667832167832 | 1 | 3 | regulation of DNA replication | DRAP1A008066 |
| GO:0045005 | 0.019543416148604 | 78.3448275862069 | 0.019667832167832 | 1 | 3 | DNA-dependent DNA replication maintenance of fidelity | DRAP1A008066 |
| GO:0051053 | 0.019543416148604 | 78.3448275862069 | 0.019667832167832 | 1 | 3 | negative regulation of DNA metabolic process | DRAP1A008066 |
| GO:0006974 | 0.023467850601467 | 5.3792270531401 | 0.622814685314685 | 3 | 95 | cellular response to DNA damage stimulus | DRAP1A008066,DRAP1A024055,DRAP1A026863 |
| GO:0098655 | 0.024006806407443 | 9.20612244897959 | 0.24256993006993 | 2 | 37 | cation transmembrane transport | DRAP1A007493,DRAP1A020079 |
| GO:0033554 | 0.024780068544384 | 5.26241134751773 | 0.635926573426573 | 3 | 97 | cellular response to stress | DRAP1A008066,DRAP1A024055,DRAP1A026863 |
| GO:0031327 | 0.032366688663080 | 39.1551724137931 | 0.032779720279720 | 1 | 5 | negative regulation of cellular biosynthetic process | DRAP1A008066 |
| GO:2000113 | 0.032366688663080 | 39.1551724137931 | 0.032779720279720 | 1 | 5 | negative regulation of cellular macromolecule biosynthetic process | DRAP1A008066 |
| GO:0010558 | 0.032366688663080 | 39.1551724137931 | 0.032779720279720 | 1 | 5 | negative regulation of macromolecule biosynthetic process | DRAP1A008066 |
| GO:0009890 | 0.038717377645821 | 31.3172413793103 | 0.039335664335664 | 1 | 6 | negative regulation of biosynthetic process | DRAP1A008066 |
| GO:0051052 | 0.038717377645821 | 31.3172413793103 | 0.039335664335664 | 1 | 6 | regulation of DNA metabolic process | DRAP1A008066 |
| GO:0006725 | 0.039321736923859 | 2.07545174932718 | 8.1097027972028 | 13 | 1237 | cellular aromatic compound metabolic process | DRAP1A003552,DRAP1A006494,DRAP1A007686,DRAP1A008066,DRAP1A008624,DRAP1A009468,DRAP1A012267,DRAP1A017508,DRAP1A018425,DRAP1A020630,DRAP1A024055,DRAP1A026863,DRAP1A027211 |
| GO:0006139 | 0.041047347179992 | 2.08848484848485 | 7.29020979020979 | 12 | 1112 | nucleobase-containing compound metabolic process | DRAP1A003552,DRAP1A007686,DRAP1A008066,DRAP1A008624,DRAP1A009468,DRAP1A012267,DRAP1A017508,DRAP1A018425,DRAP1A020630,DRAP1A024055,DRAP1A026863,DRAP1A027211 |
| GO:0034220 | 0.043439308800303 | 6.55539358600583 | 0.334353146853147 | 2 | 51 | ion transmembrane transport | DRAP1A007493,DRAP1A020079 |
| GO:0046483 | 0.044136550264770 | 2.03203823766031 | 8.23426573426574 | 13 | 1256 | heterocycle metabolic process | DRAP1A003552,DRAP1A006494,DRAP1A007686,DRAP1A008066,DRAP1A008624,DRAP1A009468,DRAP1A012267,DRAP1A017508,DRAP1A018425,DRAP1A020630,DRAP1A024055,DRAP1A026863,DRAP1A027211 |
| GO:0045934 | 0.045027766851646 | 26.0919540229885 | 0.045891608391608 | 1 | 7 | negative regulation of nucleobase-containing compound metabolic process | DRAP1A008066 |
| GO:0051172 | 0.045027766851646 | 26.0919540229885 | 0.045891608391608 | 1 | 7 | negative regulation of nitrogen compound metabolic process | DRAP1A008066 |
| GO:0006812 | 0.045401482312220 | 4.09814814814815 | 0.806381118881119 | 3 | 123 | cation transport | DRAP1A007493,DRAP1A015738,DRAP1A020079 |
| GO:0006310 | 0.048170638203583 | 6.17307692307692 | 0.354020979020979 | 2 | 54 | DNA recombination | DRAP1A024055,DRAP1A026863 |
| GO:0071705 | 0.048199967547909 | 3.99548328816622 | 0.826048951048951 | 3 | 126 | nitrogen compound transport | DRAP1A007493,DRAP1A015738,DRAP1A020079 |

**Table S5g. Related to Figure 4. GO enrichment of upregulated genes in agar_culture_40_days_vs_agar_culture_20_days**

| GOBPID | Pvalue | OddsRatio | ExpCount | Count | Size | Term | gene_id |
|---|---|---|---|---|---|---|---|
| GO:0006468 | 0.002173583382731 | 3.74729567307692 | 3.00590034965035 | 9 | 393 | protein phosphorylation | DRAP1A003993,DRAP1A005626,DRAP1A008142,DRAP1A011053,DRAP1A014442,DRAP1A016135,DRAP1A018377,DRAP1A021082,DRAP1A022862 |
| GO:0016310 | 0.002788865693463 | 3.38416666666667 | 3.74781468531469 | 10 | 490 | phosphorylation | DRAP1A003993,DRAP1A005626,DRAP1A008142,DRAP1A011053,DRAP1A014442,DRAP1A016135,DRAP1A018377,DRAP1A021082,DRAP1A021790,DRAP1A022862 |
| GO:0006979 | 0.003103010222282 | 11.7317708333333 | 0.298295454545455 | 3 | 39 | response to oxidative stress | DRAP1A000015,DRAP1A008214,DRAP1A015110 |
| GO:0036211 | 0.005128832228070 | 2.94247004357298 | 4.76507867132867 | 11 | 623 | protein modification process | DRAP1A003993,DRAP1A005626,DRAP1A006141,DRAP1A008142,DRAP1A011053,DRAP1A014442,DRAP1A016135,DRAP1A018377,DRAP1A021082,DRAP1A022862,DRAP1A023725 |
| GO:0006464 | 0.005128832228070 | 2.94247004357298 | 4.76507867132867 | 11 | 623 | cellular protein modification process | DRAP1A003993,DRAP1A005626,DRAP1A006141,DRAP1A008142,DRAP1A011053,DRAP1A014442,DRAP1A016135,DRAP1A018377,DRAP1A021082,DRAP1A022862,DRAP1A023725 |
| GO:0055114 | 0.009511942188311 | 2.78108581436077 | 4.44383741258741 | 10 | 581 | oxidation-reduction process | DRAP1A000015,DRAP1A000726,DRAP1A005489,DRAP1A007729,DRAP1A008214,DRAP1A015110,DRAP1A021481,DRAP1A022611,DRAP1A024361,DRAP1A027413 |
| GO:0043412 | 0.012348412144192 | 2.56240928882438 | 5.35402097902098 | 11 | 700 | macromolecule modification | DRAP1A003993,DRAP1A005626,DRAP1A006141,DRAP1A008142,DRAP1A011053,DRAP1A014442,DRAP1A016135,DRAP1A018377,DRAP1A021082,DRAP1A022862,DRAP1A023725 |
| GO:0009765 | 0.015240360732163 | 133.529411764706 | 0.015297202797202 | 1 | 2 | photosynthesis, light harvesting | DRAP1A006141 |
| GO:0018298 | 0.022775688098664 | 66.75 | 0.022945804195804 | 1 | 3 | protein-chromophore linkage | DRAP1A006141 |
| GO:0019253 | 0.022775688098664 | 66.75 | 0.022945804195804 | 1 | 3 | reductive pentose-phosphate cycle | DRAP1A021790 |
| GO:0019685 | 0.030254990726380 | 44.4901960784314 | 0.030594405594405 | 1 | 4 | photosynthesis, dark reaction | DRAP1A021790 |
| GO:0006793 | 0.041893392439709 | 2.12277777777778 | 5.58347902097902 | 10 | 730 | phosphorus metabolic process | DRAP1A003993,DRAP1A005626,DRAP1A008142,DRAP1A011053,DRAP1A014442,DRAP1A016135,DRAP1A018377,DRAP1A021082,DRAP1A021790,DRAP1A022862 |
| GO:0006796 | 0.041893392439709 | 2.12277777777778 | 5.58347902097902 | 10 | 730 | phosphate-containing compound metabolic process | DRAP1A003993,DRAP1A005626,DRAP1A008142,DRAP1A011053,DRAP1A014442,DRAP1A016135,DRAP1A018377,DRAP1A021082,DRAP1A021790,DRAP1A022862 |
| GO:0009086 | 0.045047136446692 | 26.6823529411765 | 0.045891608391608 | 1 | 6 | methionine biosynthetic process | DRAP1A014374 |
| GO:0015979 | 0.04956952984455 | 6.05521885521885 | 0.359484265734266 | 2 | 47 | photosynthesis | DRAP1A006141,DRAP1A021790 |
