## Supplemental Table S6 for "The *Draparnaldia* genome: alternative mechanisms for multicellularity and terrestrialization in green plants"

**Table S6. Related to Figure 5. Correspondence between the gene name used of the protein-protein interaction network and the respective Gene IDs in Draparnaldia and Arabidopsis.**

| Draparnaldia ID | Arabidopsis ID | Gene name |
|---|---|---|
| DRAP1A000051 | AT4G21120.1, AT1G17120.1, AT2G34960.1, AT3G10600.1, AT5G04770.1 | AAT1, CAT8, CAT5, CAT7, CAT6 |
| DRAP1A002066, DRAP1A014767, DRAP1A020079 | AT3G24300.1, AT3G24290.1, AT4G13510.1, AT4G28700.1, AT1G64780.1 | AMT1;3, AMT1;5, AMT1-1, AMT1;4, AMT1;2 |
| DRAP1A005178, DRAP1A017258 | AT3G02850.1, AT5G37500.2, AT4G32650.1, AT4G22200.1, AT5G46240.1, AT4G18290.2, AT4G32500.1, AT2G25600.1, AT2G26650.1 | SKOR, GORK, KAT3, KT2/3, KAT1, KAT2, KT5, SPIK, KT1 |
| DRAP1A007590, DRAP1A012102 | AT5G45380.1 | DUR3 |
| DRAP1A008450 | AT2G01170.1 | BAT1 |
| DRAP1A008968, DRAP1A024478 | AT1G54370.1, AT1G79610.1 | NHX5, NHX6 |
| DRAP1A009355 | AT5G49990.2, AT5G62890.2, AT1G10540.1, AT1G60030.1, AT5G25420.1, AT1G65550.1, AT1G49960.1, AT2G26510.1, AT2G05760.1, AT2G34190.1, AT4G38050.1, AT2G27810.1 | AT5G49990, AT5G62890, NAT8, NAT7, AT5G25420, AT1G65550, AT1G49960, PDE135, AT2G05760, AT2G34190, AT4G38050, NAT12 |
| DRAP1A009584 | AT5G55470.1, AT3G05030.1, AT5G27150.1, AT3G06370.2 | NHX3, NHX2, NHX1, NHX4 |
| DRAP1A012007 | AT5G14570.1, AT1G12940.1, AT1G08100.1, AT1G08090.1, AT5G60770.1, AT5G60780.1, AT3G45060.1, AT4G14358.1 | NRT2.7, NRT2.5, NRT2.2, NRT2:1, NRT2.4, NRT2.3, NRT2.6, AT4G14358 |
| DRAP1A017570 | AT4G03115.2 | AT4G03115 |
| DRAP1A017820 | AT3G28390.1, AT3G28360.1, AT3G28380.1, AT3G28415.1, AT3G28345.1, AT3G28860.1, AT2G36910.1, AT4G25960.1, AT1G10680.1, AT1G27940.1, AT1G28010.1, AT3G55320.1, AT2G39480.1, AT5G46540.1, AT4G18050.2, AT2G47000.2, AT3G62150.3, AT4G01830.1, AT4G01820.1, AT1G02530.2, AT1G02520.1 | ABCB18, ABCB16, ABCB17, ABCB22, ABCB15, ABCB19, ABCB1, ABCB2, PGP10, ABCB13, ABCB14, ABCB20, ABCB6, ABCB7, ABCB9, ABCB4, ABCB21, ABCB5, ABCB3, ABCB12, ABCB11 |
| DRAP1A017841 | AT5G51710.1, AT5G11800.1, AT2G19600.1 | KEA5, KEA6, KEA4 |
| DRAP1A025638 | AT5G50300.2, AT3G10960.1 | AZG2, AZG1 |
