## Supplemental Table S7 for "The *Draparnaldia* genome: alternative mechanisms for multicellularity and terrestrialization in green plants"

**Table S7. Related to Figure 6 and Figure 7. Absolute count numbers of conserved transcription factors among land plants and algae, including Draparnaldia**

| query_name | Arabidopsis_thaliana | Auxenochlorella_protothecoides | Bathycoccus_prasinos | Chara_braunii | Chlamydomonas_reinhardtii | Chlorokybus_atmophyticus | Chloropicon_primus | Chondrus_crispus | Coccomyxa_obi | Cyanidioschyzon_merolae | Cyanophora_paradoxa | Draparnaldia | Galdieria_sulphuraria | Klebsormidium_nitens | Marchantia_polymorpha | Mesostigma_viride | Mesotaenium_endlicherianum | Micromonas_RCC299 | Oryza_sativa | Ostreococcus_tauri | Physcomitrium_patens | Picochlorum_SENEW3 | Selaginella_moellendorffii | Spirogloea_muscicola | Volvox_carteri |
|---|---|---|---|---|---|---|---|---|---|---|---|---|---|---|---|---|---|---|---|---|---|---|---|---|---|
| ABI3/VP1 | 58 | 1 | 1 | 3 | 2 | 2 | 1 | 1 | 1 | 0 | 1 | 2 | 0 | 3 | 4 | 5 | 1 | 1 | 74 | 1 | 25 | 1 | 11 | 2 | 1 |
| AP2 | 43 | 3 | 4 | 1 | 6 | 3 | 8 | 0 | 5 | 0 | 0 | 5 | 0 | 2 | 5 | 1 | 1 | 5 | 80 | 2 | 29 | 4 | 12 | 12 | 5 |
| ARF | 23 | 0 | 0 | 1 | 0 | 1 | 0 | 1 | 0 | 0 | 0 | 0 | 0 | 0 | 4 | 1 | 1 | 0 | 43 | 0 | 15 | 0 | 8 | 3 | 0 |
| ARID | 9 | 3 | 1 | 3 | 5 | 2 | 3 | 0 | 7 | 3 | 1 | 6 | 5 | 1 | 5 | 5 | 3 | 2 | 15 | 1 | 8 | 2 | 5 | 8 | 2 |
| AS2/LOB | 43 | 0 | 0 | 2 | 0 | 0 | 0 | 0 | 0 | 0 | 0 | 0 | 0 | 5 | 23 | 0 | 4 | 0 | 44 | 0 | 33 | 0 | 15 | 18 | 0 |
| Argonaute | 10 | 1 | 0 | 3 | 4 | 3 | 0 | 3 | 2 | 0 | 2 | 6 | 0 | 6 | 4 | 9 | 3 | 1 | 26 | 0 | 7 | 0 | 7 | 6 | 1 |
| Aux/IAA | 28 | 0 | 0 | 2 | 0 | 0 | 0 | 0 | 0 | 0 | 0 | 0 | 0 | 0 | 3 | 0 | 0 | 0 | 51 | 0 | 4 | 0 | 9 | 0 | 0 |
| BBR/BPC | 7 | 0 | 0 | 0 | 0 | 0 | 0 | 0 | 0 | 0 | 0 | 0 | 0 | 0 | 2 | 0 | 0 | 0 | 9 | 0 | 1 | 0 | 1 | 3 | 0 |
| BES1 | 8 | 0 | 0 | 0 | 0 | 0 | 0 | 0 | 0 | 0 | 0 | 0 | 0 | 2 | 3 | 0 | 2 | 0 | 8 | 0 | 6 | 0 | 5 | 5 | 0 |
| C2C2_CO-like | 11 | 0 | 0 | 1 | 0 | 2 | 0 | 1 | 1 | 0 | 0 | 1 | 0 | 2 | 3 | 1 | 2 | 2 | 5 | 2 | 13 | 1 | 4 | 0 | 0 |
| C2C2_Dof | 36 | 0 | 1 | 1 | 1 | 1 | 1 | 0 | 1 | 0 | 0 | 1 | 0 | 1 | 2 | 1 | 1 | 1 | 37 | 2 | 21 | 0 | 30 | 0 | 1 |
| C2C2_GATA | 19 | 0 | 2 | 2 | 4 | 1 | 0 | 2 | 4 | 3 | 1 | 2 | 2 | 2 | 1 | 3 | 0 | 2 | 19 | 2 | 7 | 1 | 6 | 0 | 2 |
| C2C2_YABBY | 6 | 0 | 0 | 0 | 0 | 0 | 1 | 0 | 0 | 0 | 0 | 1 | 0 | 0 | 0 | 0 | 1 | 1 | 13 | 0 | 0 | 0 | 0 | 0 | 0 |
| C2H2 | 1 | 0 | 0 | 1 | 0 | 0 | 0 | 0 | 0 | 0 | 0 | 0 | 0 | 0 | 1 | 0 | 0 | 0 | 2 | 0 | 4 | 0 | 2 | 0 | 1 |
| C3H | 1 | 0 | 0 | 0 | 0 | 1 | 0 | 0 | 0 | 0 | 1 | 0 | 1 | 2 | 0 | 0 | 1 | 0 | 3 | 0 | 1 | 0 | 2 | 2 | 0 |
| CCAAT_HAP2(NF-YA) | 10 | 0 | 1 | 3 | 0 | 1 | 1 | 0 | 1 | 0 | 1 | 1 | 1 | 1 | 1 | 1 | 1 | 1 | 28 | 0 | 2 | 0 | 1 | 3 | 0 |
| CCAAT_HAP3(NF-YB) | 12 | 3 | 3 | 2 | 3 | 3 | 2 | 0 | 2 | 3 | 2 | 2 | 4 | 3 | 3 | 2 | 2 | 2 | 18 | 2 | 8 | 3 | 6 | 9 | 3 |
| CCAAT_HAP5(NF-YC) | 7 | 3 | 1 | 4 | 1 | 1 | 1 | 1 | 2 | 2 | 1 | 2 | 4 | 3 | 2 | 2 | 1 | 1 | 18 | 2 | 5 | 1 | 2 | 8 | 1 |
| CPP | 8 | 3 | 2 | 2 | 3 | 2 | 3 | 2 | 3 | 2 | 2 | 3 | 3 | 3 | 2 | 4 | 2 | 1 | 21 | 2 | 7 | 3 | 4 | 3 | 1 |
| CSD | 4 | 1 | 1 | 1 | 1 | 1 | 0 | 0 | 2 | 0 | 2 | 1 | 0 | 2 | 1 | 1 | 1 | 3 | 2 | 4 | 6 | 2 | 4 | 4 | 1 |
| Coactivator_p15 | 10 | 5 | 4 | 5 | 8 | 4 | 3 | 2 | 6 | 2 | 3 | 9 | 7 | 6 | 6 | 7 | 3 | 5 | 9 | 2 | 15 | 6 | 10 | 11 | 6 |
| DBP | 1 | 0 | 0 | 0 | 0 | 0 | 0 | 0 | 0 | 0 | 0 | 0 | 0 | 0 | 0 | 0 | 0 | 0 | 5 | 0 | 0 | 0 | 2 | 0 | 0 |
| DDT | 2 | 0 | 0 | 2 | 1 | 0 | 1 | 0 | 0 | 1 | 0 | 0 | 0 | 1 | 1 | 0 | 2 | 0 | 9 | 0 | 8 | 0 | 3 | 6 | 0 |
| Dicer | 2 | 0 | 0 | 0 | 0 | 0 | 0 | 0 | 0 | 0 | 0 | 0 | 0 | 0 | 1 | 0 | 0 | 0 | 1 | 0 | 1 | 0 | 1 | 3 | 0 |
| E2F/DP | 8 | 2 | 1 | 2 | 2 | 2 | 3 | 2 | 3 | 4 | 2 | 1 | 5 | 3 | 4 | 5 | 2 | 3 | 18 | 2 | 9 | 2 | 4 | 6 | 1 |
| EIL | 6 | 0 | 0 | 4 | 0 | 0 | 0 | 0 | 0 | 0 | 0 | 0 | 0 | 3 | 1 | 0 | 1 | 0 | 12 | 0 | 2 | 0 | 6 | 3 | 0 |
| ERF | 94 | 3 | 0 | 1 | 4 | 4 | 6 | 0 | 7 | 0 | 1 | 11 | 0 | 1 | 16 | 2 | 13 | 1 | 85 | 2 | 82 | 3 | 40 | 27 | 5 |
| FHA | 9 | 6 | 5 | 6 | 10 | 9 | 5 | 0 | 9 | 0 | 8 | 9 | 4 | 6 | 9 | 10 | 4 | 7 | 16 | 6 | 11 | 7 | 3 | 10 | 9 |
| GIF | 3 | 0 | 1 | 0 | 1 | 0 | 1 | 0 | 1 | 1 | 1 | 1 | 1 | 1 | 1 | 1 | 0 | 1 | 4 | 1 | 4 | 0 | 3 | 3 | 1 |
| GNAT | 21 | 13 | 18 | 16 | 31 | 15 | 13 | 9 | 22 | 9 | 19 | 19 | 9 | 17 | 20 | 16 | 10 | 19 | 29 | 17 | 32 | 15 | 19 | 38 | 26 |
| GRAS | 34 | 0 | 0 | 0 | 0 | 0 | 0 | 0 | 0 | 0 | 0 | 0 | 0 | 0 | 10 | 0 | 6 | 0 | 74 | 0 | 39 | 0 | 46 | 23 | 0 |
| GRF | 9 | 0 | 0 | 0 | 0 | 2 | 0 | 0 | 0 | 0 | 0 | 0 | 0 | 1 | 1 | 0 | 1 | 0 | 19 | 0 | 2 | 0 | 4 | 3 | 0 |
| GeBP | 19 | 0 | 0 | 0 | 0 | 0 | 0 | 0 | 0 | 0 | 0 | 0 | 0 | 0 | 0 | 0 | 0 | 0 | 18 | 0 | 0 | 0 | 1 | 0 | 0 |
| HD-Zip_III | 5 | 0 | 0 | 0 | 0 | 1 | 0 | 0 | 0 | 0 | 0 | 0 | 0 | 1 | 1 | 0 | 0 | 0 | 5 | 0 | 4 | 0 | 3 | 1 | 0 |
| HD-Zip_IV | 16 | 0 | 0 | 1 | 0 | 0 | 0 | 0 | 0 | 0 | 0 | 0 | 0 | 1 | 1 | 0 | 0 | 0 | 17 | 0 | 5 | 0 | 0 | 0 | 0 |
| HD-Zip_I_II | 22 | 0 | 0 | 2 | 0 | 1 | 0 | 0 | 0 | 0 | 0 | 0 | 0 | 2 | 2 | 0 | 0 | 0 | 38 | 0 | 21 | 0 | 6 | 3 | 0 |
| HD_BEL | 10 | 0 | 0 | 0 | 0 | 0 | 0 | 0 | 0 | 0 | 0 | 0 | 0 | 0 | 0 | 0 | 0 | 0 | 22 | 0 | 4 | 0 | 1 | 0 | 0 |
| HD_DDT | 2 | 0 | 0 | 2 | 0 | 0 | 0 | 0 | 0 | 0 | 0 | 0 | 0 | 0 | 1 | 0 | 0 | 0 | 3 | 0 | 2 | 0 | 0 | 0 | 0 |
| HD_KNOX1 | 4 | 0 | 0 | 0 | 0 | 0 | 0 | 0 | 0 | 0 | 0 | 0 | 0 | 1 | 0 | 0 | 1 | 0 | 10 | 0 | 2 | 0 | 2 | 6 | 0 |
| HD_KNOX2 | 4 | 0 | 0 | 0 | 0 | 0 | 0 | 0 | 0 | 0 | 0 | 0 | 0 | 0 | 0 | 0 | 0 | 0 | 6 | 0 | 1 | 0 | 2 | 0 | 0 |
| HD_PHD | 2 | 0 | 0 | 0 | 0 | 1 | 1 | 0 | 0 | 0 | 0 | 0 | 0 | 1 | 1 | 0 | 1 | 1 | 1 | 1 | 2 | 0 | 1 | 0 | 0 |
| HD_PINTOX | 1 | 0 | 1 | 0 | 0 | 1 | 0 | 0 | 0 | 0 | 0 | 0 | 0 | 0 | 0 | 1 | 0 | 0 | 1 | 1 | 0 | 0 | 0 | 1 | 0 |
| HD_PLINC | 17 | 0 | 0 | 3 | 0 | 2 | 0 | 0 | 0 | 0 | 0 | 0 | 0 | 2 | 1 | 0 | 1 | 0 | 17 | 0 | 10 | 0 | 7 | 2 | 0 |
| HD_WOX | 18 | 0 | 2 | 1 | 0 | 0 | 0 | 0 | 0 | 0 | 0 | 0 | 0 | 1 | 1 | 0 | 1 | 1 | 16 | 1 | 5 | 0 | 9 | 4 | 0 |
| HMG | 9 | 4 | 7 | 5 | 6 | 2 | 6 | 0 | 5 | 0 | 9 | 7 | 0 | 5 | 4 | 6 | 3 | 11 | 10 | 6 | 10 | 5 | 6 | 12 | 5 |
| HRT | 2 | 0 | 0 | 0 | 0 | 0 | 0 | 0 | 0 | 0 | 0 | 0 | 0 | 0 | 1 | 0 | 0 | 0 | 1 | 0 | 0 | 0 | 0 | 0 | 0 |
| HSF | 23 | 1 | 1 | 2 | 2 | 0 | 1 | 1 | 1 | 3 | 1 | 2 | 6 | 2 | 3 | 0 | 1 | 6 | 35 | 1 | 8 | 1 | 7 | 8 | 1 |
| IWS1 | 6 | 0 | 0 | 1 | 0 | 0 | 0 | 0 | 0 | 0 | 0 | 0 | 0 | 0 | 1 | 0 | 1 | 1 | 6 | 0 | 3 | 1 | 1 | 0 | 0 |
| Jumonji_Other | 13 | 5 | 6 | 3 | 6 | 5 | 5 | 3 | 9 | 2 | 2 | 7 | 2 | 8 | 8 | 9 | 4 | 6 | 26 | 6 | 7 | 4 | 12 | 17 | 2 |
| Jumonji_PKDM7 | 4 | 0 | 0 | 0 | 0 | 1 | 0 | 0 | 0 | 0 | 0 | 0 | 0 | 0 | 1 | 0 | 1 | 0 | 2 | 0 | 1 | 0 | 2 | 0 | 0 |
| LFY | 1 | 0 | 0 | 1 | 0 | 0 | 0 | 0 | 0 | 0 | 0 | 0 | 0 | 1 | 1 | 0 | 0 | 0 | 1 | 0 | 4 | 0 | 1 | 0 | 0 |
| LIM | 2 | 0 | 0 | 0 | 0 | 0 | 1 | 0 | 0 | 0 | 1 | 0 | 1 | 0 | 0 | 2 | 0 | 0 | 0 | 0 | 0 | 0 | 0 | 0 | 0 |
| MADS | 52 | 0 | 1 | 1 | 1 | 1 | 0 | 0 | 1 | 1 | 1 | 1 | 1 | 0 | 1 | 1 | 0 | 1 | 30 | 1 | 15 | 0 | 13 | 4 | 1 |
| MADS_MIKC | 38 | 0 | 0 | 2 | 0 | 1 | 0 | 0 | 0 | 0 | 0 | 0 | 0 | 1 | 1 | 0 | 0 | 0 | 79 | 0 | 6 | 0 | 3 | 2 | 0 |
| MBF1 | 3 | 0 | 0 | 1 | 1 | 1 | 1 | 0 | 1 | 0 | 1 | 1 | 0 | 1 | 2 | 1 | 1 | 1 | 2 | 0 | 3 | 1 | 1 | 3 | 1 |
| MYB | 141 | 7 | 8 | 16 | 14 | 9 | 8 | 6 | 7 | 10 | 9 | 14 | 11 | 25 | 32 | 7 | 11 | 12 | 178 | 8 | 55 | 6 | 36 | 18 | 8 |
| MYB-related | 21 | 2 | 0 | 4 | 6 | 3 | 4 | 3 | 4 | 5 | 6 | 4 | 8 | 5 | 4 | 3 | 4 | 4 | 32 | 1 | 16 | 1 | 11 | 18 | 1 |
| Med6 | 1 | 1 | 0 | 1 | 1 | 1 | 0 | 0 | 1 | 1 | 1 | 1 | 1 | 1 | 1 | 1 | 1 | 0 | 2 | 0 | 1 | 1 | 1 | 3 | 1 |
| Med7 | 2 | 1 | 1 | 1 | 1 | 1 | 0 | 0 | 1 | 0 | 1 | 0 | 1 | 1 | 1 | 0 | 1 | 1 | 3 | 1 | 1 | 1 | 2 | 3 | 1 |
| NAC | 98 | 0 | 0 | 0 | 0 | 0 | 0 | 0 | 0 | 0 | 0 | 0 | 0 | 0 | 9 | 0 | 2 | 0 | 164 | 0 | 29 | 0 | 20 | 13 | 0 |
| NZZ | 1 | 0 | 0 | 0 | 0 | 0 | 0 | 0 | 0 | 0 | 0 | 0 | 0 | 0 | 0 | 0 | 0 | 0 | 0 | 0 | 0 | 0 | 0 | 0 | 0 |
| OFP | 18 | 0 | 0 | 0 | 0 | 0 | 0 | 0 | 0 | 0 | 0 | 0 | 0 | 0 | 2 | 0 | 0 | 0 | 35 | 0 | 11 | 0 | 6 | 0 | 0 |
| PHD | 28 | 5 | 10 | 14 | 20 | 10 | 7 | 3 | 12 | 4 | 7 | 14 | 5 | 30 | 49 | 25 | 14 | 14 | 41 | 6 | 31 | 6 | 16 | 44 | 6 |
| PLATZ | 11 | 1 | 1 | 4 | 3 | 2 | 2 | 1 | 0 | 1 | 0 | 2 | 0 | 5 | 2 | 3 | 1 | 1 | 18 | 1 | 14 | 1 | 11 | 6 | 2 |
| PcG_EZ | 3 | 0 | 1 | 0 | 0 | 1 | 0 | 1 | 1 | 1 | 0 | 2 | 0 | 1 | 3 | 1 | 1 | 1 | 3 | 0 | 0 | 0 | 2 | 3 | 0 |
| PcG_MSI | 2 | 2 | 0 | 0 | 1 | 2 | 1 | 2 | 0 | 1 | 1 | 1 | 1 | 1 | 1 | 1 | 1 | 0 | 3 | 0 | 2 | 1 | 2 | 3 | 0 |
| PcG_VEFS | 4 | 1 | 1 | 1 | 1 | 0 | 0 | 1 | 1 | 1 | 0 | 1 | 1 | 1 | 1 | 2 | 0 | 1 | 9 | 1 | 3 | 0 | 1 | 3 | 0 |
| Pseudo_ARR-B | 4 | 0 | 2 | 0 | 2 | 1 | 2 | 0 | 2 | 0 | 0 | 2 | 0 | 2 | 2 | 3 | 2 | 0 | 16 | 1 | 4 | 1 | 1 | 3 | 0 |
| RB | 1 | 0 | 1 | 1 | 1 | 1 | 1 | 1 | 1 | 1 | 0 | 0 | 1 | 1 | 1 | 2 | 1 | 1 | 3 | 1 | 4 | 0 | 2 | 3 | 1 |
| RRN3 | 3 | 1 | 1 | 2 | 1 | 1 | 1 | 0 | 1 | 1 | 2 | 1 | 1 | 1 | 1 | 1 | 3 | 1 | 3 | 1 | 3 | 1 | 1 | 3 | 1 |
| RWP-RK | 14 | 4 | 4 | 6 | 15 | 8 | 8 | 1 | 4 | 1 | 42 | 5 | 9 | 6 | 4 | 10 | 6 | 6 | 23 | 4 | 14 | 2 | 16 | 8 | 8 |
| Rcd1-like | 3 | 1 | 1 | 1 | 2 | 1 | 1 | 1 | 1 | 1 | 1 | 1 | 1 | 1 | 2 | 2 | 2 | 1 | 14 | 1 | 3 | 1 | 1 | 7 | 2 |
| S1Fa-like | 3 | 0 | 0 | 1 | 0 | 1 | 0 | 0 | 0 | 0 | 0 | 0 | 0 | 1 | 1 | 0 | 1 | 0 | 2 | 0 | 2 | 0 | 0 | 3 | 0 |
| SAP | 1 | 0 | 0 | 0 | 0 | 0 | 0 | 0 | 0 | 0 | 0 | 0 | 0 | 0 | 0 | 0 | 0 | 0 | 0 | 0 | 0 | 0 | 1 | 0 | 0 |
| SBP | 17 | 8 | 3 | 2 | 27 | 3 | 0 | 0 | 4 | 0 | 0 | 13 | 0 | 2 | 4 | 0 | 3 | 6 | 51 | 0 | 13 | 5 | 11 | 4 | 9 |
| SET | 33 | 8 | 16 | 15 | 26 | 19 | 12 | 10 | 23 | 7 | 34 | 29 | 9 | 23 | 25 | 31 | 13 | 25 | 75 | 21 | 34 | 10 | 31 | 42 | 17 |
| SOH1 | 1 | 0 | 1 | 1 | 1 | 1 | 1 | 0 | 1 | 1 | 1 | 1 | 2 | 1 | 1 | 0 | 1 | 1 | 2 | 1 | 2 | 1 | 1 | 3 | 1 |
| SRS | 10 | 0 | 0 | 3 | 0 | 1 | 0 | 0 | 0 | 0 | 0 | 0 | 0 | 0 | 1 | 0 | 1 | 0 | 6 | 0 | 2 | 0 | 4 | 3 | 0 |
| SWI/SNF_BAF60b | 15 | 1 | 1 | 5 | 2 | 4 | 1 | 3 | 2 | 1 | 3 | 2 | 2 | 7 | 4 | 3 | 4 | 2 | 14 | 1 | 8 | 1 | 6 | 11 | 1 |
| SWI/SNF_SNF2 | 39 | 20 | 20 | 27 | 31 | 21 | 18 | 21 | 29 | 13 | 49 | 48 | 12 | 40 | 28 | 67 | 31 | 28 | 81 | 21 | 43 | 18 | 35 | 66 | 22 |
| SWI/SNF_SWI3 | 4 | 2 | 1 | 1 | 1 | 1 | 0 | 0 | 2 | 1 | 0 | 1 | 1 | 2 | 2 | 0 | 1 | 1 | 4 | 2 | 6 | 2 | 2 | 3 | 1 |
| Sigma70-like | 6 | 1 | 1 | 1 | 1 | 3 | 0 | 3 | 2 | 4 | 3 | 1 | 4 | 2 | 3 | 3 | 1 | 1 | 6 | 1 | 7 | 1 | 3 | 6 | 1 |
| Sin3 | 22 | 1 | 2 | 1 | 1 | 1 | 1 | 3 | 1 | 1 | 2 | 4 | 1 | 1 | 9 | 2 | 1 | 3 | 5 | 2 | 6 | 1 | 2 | 4 | 1 |
| Sir2 | 2 | 3 | 3 | 2 | 4 | 3 | 5 | 4 | 3 | 3 | 8 | 4 | 4 | 7 | 3 | 3 | 3 | 4 | 9 | 1 | 6 | 3 | 4 | 5 | 3 |
| TAZ | 8 | 1 | 1 | 1 | 1 | 0 | 1 | 0 | 1 | 0 | 0 | 2 | 0 | 1 | 2 | 1 | 1 | 2 | 5 | 1 | 5 | 1 | 3 | 6 | 1 |
| TFb2 | 1 | 1 | 6 | 2 | 1 | 1 | 1 | 1 | 1 | 1 | 1 | 1 | 1 | 1 | 1 | 1 | 2 | 1 | 2 | 1 | 1 | 1 | 1 | 3 | 1 |
| TRAF | 41 | 3 | 1 | 36 | 50 | 24 | 4 | 5 | 14 | 2 | 29 | 17 | 8 | 37 | 38 | 9 | 8 | 3 | 131 | 2 | 44 | 4 | 19 | 27 | 13 |
| TUB | 10 | 1 | 2 | 4 | 3 | 2 | 5 | 0 | 2 | 0 | 5 | 1 | 0 | 6 | 5 | 3 | 2 | 1 | 23 | 1 | 8 | 1 | 6 | 6 | 2 |
| Trihelix | 22 | 0 | 0 | 2 | 0 | 2 | 0 | 0 | 0 | 0 | 0 | 0 | 0 | 3 | 14 | 0 | 5 | 0 | 25 | 0 | 26 | 0 | 18 | 4 | 0 |
| WRKY | 72 | 1 | 3 | 1 | 2 | 0 | 1 | 0 | 1 | 0 | 0 | 1 | 0 | 2 | 14 | 2 | 8 | 1 | 131 | 3 | 37 | 1 | 18 | 30 | 1 |
| Whirly | 3 | 1 | 1 | 1 | 1 | 1 | 0 | 0 | 1 | 0 | 0 | 1 | 0 | 1 | 1 | 1 | 1 | 1 | 2 | 1 | 0 | 1 | 1 | 3 | 1 |
| Zinc_finger,AN1_and_A20_type | 10 | 1 | 1 | 1 | 1 | 1 | 1 | 0 | 1 | 1 | 1 | 1 | 2 | 1 | 2 | 0 | 1 | 0 | 19 | 0 | 6 | 1 | 5 | 3 | 1 |
| Zinc_finger,MIZ_type | 3 | 0 | 1 | 2 | 1 | 2 | 1 | 1 | 1 | 1 | 6 | 0 | 0 | 3 | 3 | 3 | 6 | 1 | 6 | 1 | 6 | 0 | 2 | 4 | 1 |
| Zinc_finger,ZPR1 | 2 | 1 | 0 | 4 | 1 | 0 | 1 | 0 | 1 | 1 | 0 | 0 | 1 | 1 | 1 | 1 | 1 | 0 | 2 | 0 | 1 | 0 | 1 | 4 | 1 |
| bHLH | 20 | 0 | 0 | 1 | 0 | 3 | 0 | 0 | 0 | 0 | 7 | 0 | 0 | 3 | 3 | 2 | 0 | 0 | 45 | 0 | 10 | 0 | 5 | 9 | 0 |
| bHLH_TCP | 24 | 0 | 0 | 2 | 0 | 0 | 0 | 0 | 0 | 0 | 0 | 0 | 0 | 0 | 2 | 0 | 1 | 0 | 26 | 0 | 6 | 0 | 6 | 4 | 0 |
| bZIP | 54 | 1 | 1 | 12 | 1 | 12 | 3 | 0 | 9 | 0 | 3 | 6 | 2 | 10 | 10 | 13 | 4 | 5 | 117 | 3 | 39 | 4 | 16 | 26 | 4 |
| mTERF | 35 | 2 | 2 | 8 | 3 | 2 | 2 | 1 | 2 | 1 | 23 | 5 | 0 | 9 | 14 | 4 | 19 | 4 | 54 | 2 | 15 | 5 | 12 | 22 | 4 |
| tify | 16 | 0 | 0 | 2 | 0 | 1 | 0 | 0 | 0 | 0 | 0 | 0 | 0 | 1 | 4 | 0 | 2 | 0 | 21 | 0 | 15 | 0 | 9 | 6 | 0 |
| VARL | 0 | 0 | 1 | 0 | 12 | 0 | 0 | 0 | 2 | 0 | 0 | 1 | 0 | 0 | 0 | 0 | 0 | 0 | 0 | 0 | 0 | 0 | 0 | 1 | 11 |
| Alfin-like | 0 | 0 | 0 | 0 | 0 | 0 | 0 | 0 | 0 | 0 | 0 | 0 | 0 | 0 | 1 | 0 | 0 | 0 | 0 | 0 | 0 | 0 | 0 | 0 | 0 |
