## Supplemental Table S8 for "The *Draparnaldia* genome: alternative mechanisms for multicellularity and terrestrialization in green plants"

**Table S8a. Related to Figure 7. Detection statistics of the cytokinins both in the alga and in the media from Draparnaldia and Klebsormidium nitens**

**Phytohormonal levels: pmol/g FW (algae); pmol/ml (media); mean ± SD)**

| | | | tZ | | | | tZR | | | | tZOG | |
|---|---|---|---|---|---|---|---|---|---|---|---|---|
| **Algae** | Draparnaldia | 211211_21086_H_Caisova_1 | < LOD | | | | 0.09 | **0.10** | ± | 0.01 | < LOD | |
| | | 211211_21086_H_Caisova_2 | < LOD | < LOD | | | 0.09 | RSD | | 6 % | < LOD | < LOD |
| | | 211211_21086_H_Caisova_3 | < LOD | | | | 0.10 | | | | < LOD | |
| | Klebsormidium | 211211_21086_H_Caisova_4 | < LOD | | | | 0.02 | | | | < LOD | |
| | | 211211_21086_H_Caisova_5 | < LOD | < LOD | | | < LOD | < LOD | | | < LOD | < LOD |
| | | 211211_21086_H_Caisova_6 | < LOD | | | | < LOD | | | | < LOD | |
| **Media** | Draparnaldia media (with algae) | 211211_21086_H_Caisova_7 | < LOD | | | | 7.30 | **6.77** | ± | 0.53 | < LOD | |
| | | 211211_21086_H_Caisova_8 | < LOD | < LOD | | | 6.78 | RSD | | 8 % | < LOD | < LOD |
| | | 211211_21086_H_Caisova_9 | < LOD | | | | 6.24 | | | | < LOD | |
| | Klebsormidium media (with algae) | 211211_21086_H_Caisova_10 | < LOD | | | | < LOD | | | | < LOD | |
| | | 211211_21086_H_Caisova_11 | < LOD | < LOD | | | < LOD | < LOD | | | < LOD | < LOD |
| | | 211211_21086_H_Caisova_12 | < LOD | | | | < LOD | | | | < LOD | |

| | | | tZROG | | | | tZ7G | | | | tZ9G | |
|---|---|---|---|---|---|---|---|---|---|---|---|---|
| **Algae** | Draparnaldia | 211211_21086_H_Caisova_1 | < LOD | | | | < LOD | | | | < LOD | |
| | | 211211_21086_H_Caisova_2 | < LOD | < LOD | | | < LOD | < LOD | | | < LOD | < LOD |
| | | 211211_21086_H_Caisova_3 | < LOD | | | | < LOD | | | | < LOD | |
| | Klebsormidium | 211211_21086_H_Caisova_4 | < LOD | | | | < LOD | | | | < LOD | |
| | | 211211_21086_H_Caisova_5 | < LOD | < LOD | | | < LOD | < LOD | | | < LOD | < LOD |
| | | 211211_21086_H_Caisova_6 | < LOD | | | | < LOD | | | | < LOD | |
| **Media** | Draparnaldia media (with algae) | 211211_21086_H_Caisova_7 | < LOD | | | | < LOD | | | | < LOD | |
| | | 211211_21086_H_Caisova_8 | < LOD | < LOD | | | < LOD | < LOD | | | < LOD | < LOD |
| | | 211211_21086_H_Caisova_9 | < LOD | | | | < LOD | | | | < LOD | |
| | Klebsormidium media (with algae) | 211211_21086_H_Caisova_10 | < LOD | | | | < LOD | | | | < LOD | |
| | | 211211_21086_H_Caisova_11 | < LOD | < LOD | | | < LOD | < LOD | | | < LOD | < LOD |
| | | 211211_21086_H_Caisova_12 | < LOD | | | | < LOD | | | | < LOD | |

| | | | cZ | | | | cZR | | | | cZOG | |
|---|---|---|---|---|---|---|---|---|---|---|---|---|
| **Algae** | Draparnaldia | 211211_21086_H_Caisova_1 | < LOD | | | | 3.67 | **4.38** | ± | 0.64 | < LOD | |
| | | 211211_21086_H_Caisova_2 | < LOD | < LOD | | | 4.56 | RSD | | 15 % | < LOD | < LOD |
| | | 211211_21086_H_Caisova_3 | < LOD | | | | 4.90 | | | | < LOD | |
| | Klebsormidium | 211211_21086_H_Caisova_4 | < LOD | | | | 0.10 | **0.11** | ± | 0.01 | < LOD | |
| | | 211211_21086_H_Caisova_5 | < LOD | < LOD | | | 0.12 | RSD | | 11 % | < LOD | < LOD |
| | | 211211_21086_H_Caisova_6 | < LOD | | | | 0.10 | | | | < LOD | |
| **Media** | Draparnaldia media (with algae) | 211211_21086_H_Caisova_7 | 135.18 | **132.93** | ± | 2.19 | 1705.59 | **1648.88** | ± | 74.33 | < LOD | |
| | | 211211_21086_H_Caisova_8 | 130.81 | RSD | | 2 % | 1564.74 | RSD | | 5 % | < LOD | < LOD |
| | | 211211_21086_H_Caisova_9 | 132.82 | | | | 1676.33 | | | | < LOD | |
| | Klebsormidium media (with algae) | 211211_21086_H_Caisova_10 | 14.95 | **14.29** | ± | 0.95 | 9.11 | **7.32** | ± | 1.55 | < LOD | |
| | | 211211_21086_H_Caisova_11 | 14.71 | RSD | | 7 % | 6.36 | RSD | | 21 % | < LOD | < LOD |
| | | 211211_21086_H_Caisova_12 | 13.20 | | | | 6.50 | | | | < LOD | |

| | | | cZROG | | | | cZ7G | | | | cZ9G | |
|---|---|---|---|---|---|---|---|---|---|---|---|---|
| **Algae** | Draparnaldia | 211211_21086_H_Caisova_1 | < LOD | | | | < LOD | | | | < LOD | |
| | | 211211_21086_H_Caisova_2 | < LOD | < LOD | | | < LOD | < LOD | | | < LOD | < LOD |
| | | 211211_21086_H_Caisova_3 | < LOD | | | | < LOD | | | | < LOD | |
| | Klebsormidium | 211211_21086_H_Caisova_4 | < LOD | | | | < LOD | | | | < LOD | |
| | | 211211_21086_H_Caisova_5 | < LOD | < LOD | | | < LOD | < LOD | | | < LOD | < LOD |
| | | 211211_21086_H_Caisova_6 | < LOD | | | | < LOD | | | | < LOD | |
| **Media** | Draparnaldia media (with algae) | 211211_21086_H_Caisova_7 | < LOD | | | | < LOD | | | | < LOD | |
| | | 211211_21086_H_Caisova_8 | < LOD | < LOD | | | < LOD | < LOD | | | < LOD | < LOD |
| | | 211211_21086_H_Caisova_9 | < LOD | | | | < LOD | | | | < LOD | |
| | Klebsormidium media (with algae) | 211211_21086_H_Caisova_10 | < LOD | | | | < LOD | | | | < LOD | |
| | | 211211_21086_H_Caisova_11 | < LOD | < LOD | | | < LOD | < LOD | | | < LOD | < LOD |
| | | 211211_21086_H_Caisova_12 | < LOD | | | | < LOD | | | | < LOD | |

| | | | DHZ | | | | DHZR | | | | DHZOG | |
|---|---|---|---|---|---|---|---|---|---|---|---|---|
| **Algae** | Draparnaldia | 211211_21086_H_Caisova_1 | < LOD | | | | < LOD | | | | < LOD | |
| | | 211211_21086_H_Caisova_2 | < LOD | < LOD | | | < LOD | < LOD | | | < LOD | < LOD |
| | | 211211_21086_H_Caisova_3 | < LOD | | | | < LOD | | | | < LOD | |
| | Klebsormidium | 211211_21086_H_Caisova_4 | < LOD | | | | < LOD | | | | < LOD | |
| | | 211211_21086_H_Caisova_5 | < LOD | < LOD | | | < LOD | < LOD | | | < LOD | < LOD |
| | | 211211_21086_H_Caisova_6 | < LOD | | | | < LOD | | | | < LOD | |
| **Media** | Draparnaldia media (with algae) | 211211_21086_H_Caisova_7 | 76.41 | **73.13** | ± | 3.75 | < LOD | | | | < LOD | |
| | | 211211_21086_H_Caisova_8 | 73.95 | RSD | | 5 % | < LOD | < LOD | | | < LOD | < LOD |
| | | 211211_21086_H_Caisova_9 | 69.04 | | | | < LOD | | | | < LOD | |
| | Klebsormidium media (with algae) | 211211_21086_H_Caisova_10 | < LOD | | | | < LOD | | | | < LOD | |
| | | 211211_21086_H_Caisova_11 | < LOD | < LOD | | | < LOD | < LOD | | | < LOD | < LOD |
| | | 211211_21086_H_Caisova_12 | < LOD | | | | < LOD | | | | < LOD | |

| | | | DHZROG | | | | DHZ7G | | | | DHZ9G | |
|---|---|---|---|---|---|---|---|---|---|---|---|---|
| **Algae** | Draparnaldia | 211211_21086_H_Caisova_1 | < LOD | | | | < LOD | | | | < LOD | |
| | | 211211_21086_H_Caisova_2 | < LOD | < LOD | | | < LOD | < LOD | | | < LOD | < LOD |
| | | 211211_21086_H_Caisova_3 | < LOD | | | | < LOD | | | | < LOD | |
| | Klebsormidium | 211211_21086_H_Caisova_4 | < LOD | | | | < LOD | | | | < LOD | |
| | | 211211_21086_H_Caisova_5 | < LOD | < LOD | | | < LOD | < LOD | | | < LOD | < LOD |
| | | 211211_21086_H_Caisova_6 | < LOD | | | | < LOD | | | | < LOD | |
| **Media** | Draparnaldia media (with algae) | 211211_21086_H_Caisova_7 | < LOD | | | | < LOD | | | | < LOD | |
| | | 211211_21086_H_Caisova_8 | < LOD | < LOD | | | < LOD | < LOD | | | < LOD | < LOD |
| | | 211211_21086_H_Caisova_9 | < LOD | | | | < LOD | | | | < LOD | |
| | Klebsormidium media (with algae) | 211211_21086_H_Caisova_10 | < LOD | | | | < LOD | | | | < LOD | |
| | | 211211_21086_H_Caisova_11 | < LOD | < LOD | | | < LOD | < LOD | | | < LOD | < LOD |
| | | 211211_21086_H_Caisova_12 | < LOD | | | | < LOD | | | | < LOD | |

| | | | iP | | | | iPR | | | | iP7G | | | | iP9G | |
|---|---|---|---|---|---|---|---|---|---|---|---|---|---|---|---|---|
| **Algae** | Draparnaldia | 211211_21086_H_Caisova_1 | 0.20 | **0.15** | ± | 0.04 | < LOD | | | | < LOD | | | | < LOD | |
| | | 211211_21086_H_Caisova_2 | 0.13 | RSD | | 30 % | < LOD | < LOD | | | < LOD | < LOD | | | < LOD | < LOD |
| | | 211211_21086_H_Caisova_3 | 0.12 | | | | < LOD | | | | < LOD | | | | < LOD | |
| | Klebsormidium | 211211_21086_H_Caisova_4 | 0.02 | **0.03** | ± | 0.01 | < LOD | | | | < LOD | | | | < LOD | |
| | | 211211_21086_H_Caisova_5 | 0.03 | RSD | | 30 % | < LOD | < LOD | | | < LOD | < LOD | | | < LOD | < LOD |
| | | 211211_21086_H_Caisova_6 | 0.04 | | | | < LOD | | | | < LOD | | | | < LOD | |
| **Media** | Draparnaldia media (with algae) | 211211_21086_H_Caisova_7 | 30.69 | **30.54** | ± | 0.45 | 141.90 | **146.26** | ± | 12.09 | < LOD | | | | < LOD | |
| | | 211211_21086_H_Caisova_8 | 30.90 | RSD | | 1 % | 136.96 | RSD | | 8 % | < LOD | < LOD | | | < LOD | < LOD |
| | | 211211_21086_H_Caisova_9 | 30.04 | | | | 159.93 | | | | < LOD | | | | < LOD | |
| | Klebsormidium media (with algae) | 211211_21086_H_Caisova_10 | 4.56 | **3.89** | ± | 1.19 | 7.25 | **7.87** | ± | 2.35 | < LOD | | | | < LOD | |
| | | 211211_21086_H_Caisova_11 | 4.60 | RSD | | 31 % | 10.47 | RSD | | 30 % | < LOD | < LOD | | | < LOD | < LOD |
| | | 211211_21086_H_Caisova_12 | 2.52 | | | | 5.89 | | | | < LOD | | | | < LOD | |

**Table S8b. Related to Figure 7. Detection statistics of the 2MeS cytokinins both in the alga and in the media from Draparnaldia and Klebsormidium nitens**

| Phytohormonal levels: pmol/g FW (algae); pmol/ml (media); mean ± SD) | | | | | | | | | | | | | | | | | | |
|---|---|---|---|---|---|---|---|---|---|---|---|---|---|---|---|---|---|---|
| | | | **2MeScZ** | | | | **2MeScZR** | | | | **2MeSiP** | | | | **2MeSiPR** | | | |
| **Algae** | Draparnaldia | 211211_21086_H_Caisova_1 | < LOD | **< LOD** | | | 0.77 | **0.76** | ± | 0.01 | 1.31 | **1.02** | ± | 0.25 | 1.30 | **1.30** | ± | 0.02 |
| | | 211211_21086_H_Caisova_2 | < LOD | | | | 0.75 | ***RSD*** | | ***1 %*** | 0.85 | ***RSD*** | | ***25 %*** | 1.28 | ***RSD*** | | ***1 %*** |
| | | 211211_21086_H_Caisova_3 | < LOD | | | | 0.76 | | | | 0.90 | | | | 1.31 | | | |
| | Klebsormidium | 211211_21086_H_Caisova_4 | < LOD | **< LOD** | | | 0.09 | **0.10** | ± | 0.01 | 1.39 | **1.61** | ± | 0.19 | 0.61 | **0.43** | ± | 0.16 |
| | | 211211_21086_H_Caisova_5 | < LOD | | | | 0.11 | ***RSD*** | | ***13 %*** | 1.74 | ***RSD*** | | ***12 %*** | 0.35 | ***RSD*** | | ***37 %*** |
| | | 211211_21086_H_Caisova_6 | < LOD | | | | 0.10 | | | | 1.69 | | | | 0.33 | | | |
| **Media** | Draparnaldia media (with algae) | 211211_21086_H_Caisova_7 | 194.29 | **216.28** | ± | 23.79 | 529.61 | **541.64** | ± | 10.49 | 250.58 | **279.80** | ± | 25.45 | 304.06 | **305.92** | ± | 7.25 |
| | | 211211_21086_H_Caisova_8 | 241.53 | ***RSD*** | | ***11 %*** | 548.86 | ***RSD*** | | ***2 %*** | 297.09 | ***RSD*** | | ***9 %*** | 299.78 | ***RSD*** | | ***2 %*** |
| | | 211211_21086_H_Caisova_9 | 213.02 | | | | 546.47 | | | | 291.73 | | | | 313.91 | | | |
| | Klebsormidium media (with algae) | 211211_21086_H_Caisova_10 | 56.57 | **50.72** | ± | 7.04 | 33.65 | **31.50** | ± | 2.78 | 45.44 | **40.59** | ± | 4.26 | 14.91 | **14.12** | ± | 0.80 |
| | | 211211_21086_H_Caisova_11 | 42.90 | ***RSD*** | | ***14 %*** | 28.36 | ***RSD*** | | ***9 %*** | 37.49 | ***RSD*** | | ***10 %*** | 14.16 | ***RSD*** | | ***6 %*** |
| | | 211211_21086_H_Caisova_12 | 52.70 | | | | 32.48 | | | | 38.84 | | | | 13.30 | | | |

**Table S8c. Related to Figure 7. Detection statistics of the jasmonates both in the alga and in the media from Draparnaldia and Klebsormidium nitens**

**Phytohormonal levels: pmol/g FW (algae); pmol/ml (media); mean ± SD)**

| | | | JA | | | | 9,10-DHJA | | | | 12-OH-JA | |
|---|---|---|---|---|---|---|---|---|---|---|---|---|
| **Algae** | Draparnaldia | 211211_21086_H_Caisova_1 | < LOD | | | | < LOD | | | | < LOD | |
| | | 211211_21086_H_Caisova_2 | < LOD | **< LOD** | | | < LOD | **< LOD** | | | < LOD | **< LOD** |
| | | 211211_21086_H_Caisova_3 | < LOD | | | | < LOD | | | | < LOD | |
| | Klebsormidium | 211211_21086_H_Caisova_4 | < LOD | | | | < LOD | | | | < LOD | |
| | | 211211_21086_H_Caisova_5 | < LOD | **< LOD** | | | < LOD | **< LOD** | | | < LOD | **< LOD** |
| | | 211211_21086_H_Caisova_6 | < LOD | | | | < LOD | | | | < LOD | |
| **Media** | Draparnaldia media (with algae) | 211211_21086_H_Caisova_7 | < LOD | | | | < LOD | | | | < LOD | |
| | | 211211_21086_H_Caisova_8 | < LOD | **< LOD** | | | < LOD | **< LOD** | | | < LOD | **< LOD** |
| | | 211211_21086_H_Caisova_9 | < LOD | | | | < LOD | | | | < LOD | |
| | Klebsormidium media (with algae) | 211211_21086_H_Caisova_10 | 694.04 | **873.15** | ± | 156.35 | < LOD | | | | < LOD | |
| | | 211211_21086_H_Caisova_11 | 943.10 | ***RSD*** | | ***18 %*** | < LOD | **< LOD** | | | < LOD | **< LOD** |
| | | 211211_21086_H_Caisova_12 | 982.32 | | | | < LOD | | | | < LOD | |

| | | | OPC-4 | | | | OPC-6 | | | | cisOPDA | | | |
|---|---|---|---|---|---|---|---|---|---|---|---|---|---|---|
| **Algae** | Draparnaldia | 211211_21086_H_Caisova_1 | < LOD | | | | < LOD | | | | < LOD | | | |
| | | 211211_21086_H_Caisova_2 | < LOD | **< LOD** | | | < LOD | **< LOD** | | | < LOD | **< LOD** | | |
| | | 211211_21086_H_Caisova_3 | < LOD | | | | < LOD | | | | < LOD | | | |
| | Klebsormidium | 211211_21086_H_Caisova_4 | 61.27 | **91.38** | ± | 28.56 | 341.14 | **308.56** | ± | 28.65 | 164859.39 | **157889.46** | ± | 6706.68 |
| | | 211211_21086_H_Caisova_5 | 118.08 | ***RSD*** | | ***31 %*** | 287.30 | ***RSD*** | | ***9 %*** | 151481.38 | ***RSD*** | | ***4 %*** |
| | | 211211_21086_H_Caisova_6 | 94.80 | | | | 297.23 | | | | 157327.61 | | | |
| **Media** | Draparnaldia media (with algae) | 211211_21086_H_Caisova_7 | < LOD | | | | < LOD | | | | 8661.21 | **4709.73** | ± | 3545.67 |
| | | 211211_21086_H_Caisova_8 | < LOD | **< LOD** | | | < LOD | **< LOD** | | | 3661.96 | ***RSD*** | | ***75 %*** |
| | | 211211_21086_H_Caisova_9 | < LOD | | | | < LOD | | | | 1806.02 | | | |
| | Klebsormidium media (with algae) | 211211_21086_H_Caisova_10 | 10622.38 | **10348.16** | ± | 1309.33 | 751.77 | **504.01** | ± | 221.63 | 463223.03 | **453489.78** | ± | 9877.70 |
| | | 211211_21086_H_Caisova_11 | 11498.66 | ***RSD*** | | ***13 %*** | 324.65 | ***RSD*** | | ***44 %*** | 453772.61 | ***RSD*** | | ***2 %*** |
| | | 211211_21086_H_Caisova_12 | 8923.43 | | | | 435.60 | | | | 443473.71 | | | |

| | | | dnOPDA | | JA-Ile | | JA-Val | | JA-Phe | | JA-Trp | | SA | |
|---|---|---|---|---|---|---|---|---|---|---|---|---|---|---|
| **Algae** | Draparnaldia | 211211_21086_H_Caisova_1 | < LOD | | < LOD | | < LOD | | < LOD | | < LOD | | < LOD | |
| | | 211211_21086_H_Caisova_2 | < LOD | **< LOD** | < LOD | **< LOD** | < LOD | **< LOD** | < LOD | **< LOD** | < LOD | **< LOD** | < LOD | **< LOD** |
| | | 211211_21086_H_Caisova_3 | < LOD | | < LOD | | < LOD | | < LOD | | < LOD | | < LOD | |
| | Klebsormidium | 211211_21086_H_Caisova_4 | < LOD | | < LOD | | < LOD | | < LOD | | < LOD | | < LOD | |
| | | 211211_21086_H_Caisova_5 | < LOD | **< LOD** | < LOD | **< LOD** | < LOD | **< LOD** | < LOD | **< LOD** | < LOD | **< LOD** | < LOD | **< LOD** |
| | | 211211_21086_H_Caisova_6 | < LOD | | < LOD | | < LOD | | < LOD | | < LOD | | < LOD | |
| **Media** | Draparnaldia media (with algae) | 211211_21086_H_Caisova_7 | < LOD | | < LOD | | < LOD | | < LOD | | < LOD | | < LOD | |
| | | 211211_21086_H_Caisova_8 | < LOD | **< LOD** | < LOD | **< LOD** | < LOD | **< LOD** | < LOD | **< LOD** | < LOD | **< LOD** | < LOD | **< LOD** |
| | | 211211_21086_H_Caisova_9 | < LOD | | < LOD | | < LOD | | < LOD | | < LOD | | < LOD | |
| | Klebsormidium media (with algae) | 211211_21086_H_Caisova_10 | < LOD | | < LOD | | < LOD | | < LOD | | < LOD | | < LOD | |
| | | 211211_21086_H_Caisova_11 | < LOD | **< LOD** | < LOD | **< LOD** | < LOD | **< LOD** | < LOD | **< LOD** | < LOD | **< LOD** | < LOD | **< LOD** |
| | | 211211_21086_H_Caisova_12 | < LOD | | < LOD | | < LOD | | < LOD | | < LOD | | < LOD | |

**Table S8d. Related to Figure 7. Detection statistics of the abscisates both in the alga and in the media from Draparnaldia and Klebsormidium nitens**

| Phytohormonal levels: pmol/g FW (algae); pmol/ml (media); mean ± SD) | | | | | | | | | | | | | | |
|---|---|---|---|---|---|---|---|---|---|---|---|---|---|---|
| | | | ABA | | | | 7-OH-ABA | | PA | | DPA | | neoPA | |
| **Algae** | Draparnaldia | 211211_21086_H_Caisova_1 | < LOD | | | | < LOD | | < LOD | | < LOD | | < LOD | |
| | | 211211_21086_H_Caisova_2 | < LOD | **< LOD** | | | < LOD | **< LOD** | < LOD | **< LOD** | < LOD | **< LOD** | < LOD | **< LOD** |
| | | 211211_21086_H_Caisova_3 | < LOD | | | | < LOD | | < LOD | | < LOD | | < LOD | |
| | Klebsormidium | 211211_21086_H_Caisova_4 | < LOD | | | | < LOD | | < LOD | | < LOD | | < LOD | |
| | | 211211_21086_H_Caisova_5 | < LOD | **< LOD** | | | < LOD | **< LOD** | < LOD | **< LOD** | < LOD | **< LOD** | < LOD | **< LOD** |
| | | 211211_21086_H_Caisova_6 | < LOD | | | | < LOD | | < LOD | | < LOD | | < LOD | |
| **Media** | Draparnaldia media (with algae) | 211211_21086_H_Caisova_7 | 7.99 | **17.21** | ± | 8.65 | < LOD | | < LOD | | < LOD | | < LOD | |
| | | 211211_21086_H_Caisova_8 | 18.52 | ***RSD*** | | ***50 %*** | < LOD | **< LOD** | < LOD | **< LOD** | < LOD | **< LOD** | < LOD | **< LOD** |
| | | 211211_21086_H_Caisova_9 | 25.13 | | | | < LOD | | < LOD | | < LOD | | < LOD | |
| | Klebsormidium media (with algae) | 211211_21086_H_Caisova_10 | 159.94 | **150.12** | ± | 10.40 | < LOD | | < LOD | | < LOD | | < LOD | |
| | | 211211_21086_H_Caisova_11 | 151.19 | ***RSD*** | | ***7 %*** | < LOD | **< LOD** | < LOD | **< LOD** | < LOD | **< LOD** | < LOD | **< LOD** |
| | | 211211_21086_H_Caisova_12 | 139.23 | | | | < LOD | | < LOD | | < LOD | | < LOD | |

**Table S8e. Related to Figure 7. Detection statistics of the gibberellins both in the alga and in the media from Draparnaldia and Klebsormidium nitens**

**Phytohormonal levels: pmol/g FW (algae); pmol/ml (media); mean ± SD)**

| | | | GA1 | | GA3 | | GA4 | | GA5 | | GA6 | | GA8 | |
|---|---|---|---|---|---|---|---|---|---|---|---|---|---|---|
| Algae | Draparnaldia | 211211_21086_H_Caisova_1 | < LOD | < LOD | < LOD | < LOD | < LOD | < LOD | < LOD | < LOD | < LOD | < LOD | < LOD | < LOD |
| | | 211211_21086_H_Caisova_2 | < LOD | | < LOD | | < LOD | | < LOD | | < LOD | | < LOD | |
| | | 211211_21086_H_Caisova_3 | < LOD | | < LOD | | < LOD | | < LOD | | < LOD | | < LOD | |
| | Klebsormidium | 211211_21086_H_Caisova_4 | < LOD | < LOD | < LOD | < LOD | < LOD | < LOD | < LOD | < LOD | < LOD | < LOD | < LOD | < LOD |
| | | 211211_21086_H_Caisova_5 | < LOD | | < LOD | | < LOD | | < LOD | | < LOD | | < LOD | |
| | | 211211_21086_H_Caisova_6 | < LOD | | < LOD | | < LOD | | < LOD | | < LOD | | < LOD | |
| Media | Draparnaldia media (with algae) | 211211_21086_H_Caisova_7 | < LOD | < LOD | < LOD | < LOD | < LOD | < LOD | < LOD | < LOD | < LOD | < LOD | < LOD | < LOD |
| | | 211211_21086_H_Caisova_8 | < LOD | | < LOD | | < LOD | | < LOD | | < LOD | | < LOD | |
| | | 211211_21086_H_Caisova_9 | < LOD | | < LOD | | < LOD | | < LOD | | < LOD | | < LOD | |
| | Klebsormidium media (with algae) | 211211_21086_H_Caisova_10 | < LOD | < LOD | < LOD | < LOD | < LOD | < LOD | < LOD | < LOD | < LOD | < LOD | < LOD | < LOD |
| | | 211211_21086_H_Caisova_11 | < LOD | | < LOD | | < LOD | | < LOD | | < LOD | | < LOD | |
| | | 211211_21086_H_Caisova_12 | < LOD | | < LOD | | < LOD | | < LOD | | < LOD | | n.d. | |

| | | | GA15 | | GA19 | | GA24 | | GA29 | | GA34 | | GA44 | |
|---|---|---|---|---|---|---|---|---|---|---|---|---|---|---|
| Algae | Draparnaldia | 211211_21086_H_Caisova_1 | < LOD | < LOD | < LOD | < LOD | < LOD | < LOD | < LOD | < LOD | n.d. | n.d. | < LOD | < LOD |
| | | 211211_21086_H_Caisova_2 | < LOD | | < LOD | | < LOD | | < LOD | | n.d. | | < LOD | |
| | | 211211_21086_H_Caisova_3 | < LOD | | < LOD | | < LOD | | < LOD | | n.d. | | < LOD | |
| | Klebsormidium | 211211_21086_H_Caisova_4 | < LOD | < LOD | < LOD | < LOD | < LOD | < LOD | < LOD | < LOD | n.d. | n.d. | < LOD | < LOD |
| | | 211211_21086_H_Caisova_5 | < LOD | | < LOD | | < LOD | | < LOD | | n.d. | | < LOD | |
| | | 211211_21086_H_Caisova_6 | < LOD | | < LOD | | < LOD | | < LOD | | n.d. | | < LOD | |
| Media | Draparnaldia media (with algae) | 211211_21086_H_Caisova_7 | < LOD | < LOD | < LOD | < LOD | < LOD | < LOD | < LOD | < LOD | n.d. | n.d. | < LOD | < LOD |
| | | 211211_21086_H_Caisova_8 | < LOD | | < LOD | | < LOD | | < LOD | | n.d. | | < LOD | |
| | | 211211_21086_H_Caisova_9 | < LOD | | < LOD | | < LOD | | < LOD | | n.d. | | < LOD | |
| | Klebsormidium media (with algae) | 211211_21086_H_Caisova_10 | < LOD | < LOD | < LOD | < LOD | < LOD | < LOD | < LOD | < LOD | n.d. | n.d. | < LOD | < LOD |
| | | 211211_21086_H_Caisova_11 | < LOD | | < LOD | | < LOD | | < LOD | | n.d. | | < LOD | |
| | | 211211_21086_H_Caisova_12 | < LOD | | < LOD | | < LOD | | n.d. | | n.d. | | < LOD | |

| | | | GA51 | | GA53 | |
|---|---|---|---|---|---|---|
| Algae | Draparnaldia | 211211_21086_H_Caisova_1 | n.d. | n.d. | n.d. | n.d. |
| | | 211211_21086_H_Caisova_2 | n.d. | | n.d. | |
| | | 211211_21086_H_Caisova_3 | n.d. | | n.d. | |
| | Klebsormidium | 211211_21086_H_Caisova_4 | n.d. | n.d. | n.d. | n.d. |
| | | 211211_21086_H_Caisova_5 | n.d. | | n.d. | |
| | | 211211_21086_H_Caisova_6 | n.d. | | n.d. | |
| Media | Draparnaldia media (with algae) | 211211_21086_H_Caisova_7 | n.d. | n.d. | n.d. | n.d. |
| | | 211211_21086_H_Caisova_8 | n.d. | | n.d. | |
| | | 211211_21086_H_Caisova_9 | n.d. | | n.d. | |
| | Klebsormidium media (with algae) | 211211_21086_H_Caisova_10 | n.d. | n.d. | n.d. | n.d. |
| | | 211211_21086_H_Caisova_11 | n.d. | | n.d. | |
| | | 211211_21086_H_Caisova_12 | n.d. | | n.d. | |

**Table S8f. Related to Figure 7. Detection statistics of the brassinosteroids both in the alga and in the media from Draparnaldia and Klebsormidium nitens**

**Phytohormonal levels: pmol/g FW (algae); pmol/ml (media); mean ± SD)**

| | | | **BL** | | | | **epiBL** | | | | **homoBL** | | | | **CS** | | | | **epiCS** | | | | **homoCS** | | | |
|---|---|---|---|---|---|---|---|---|---|---|---|---|---|---|---|---|---|---|---|---|---|---|---|---|---|---|
| **Algae** | Draparnaldia | 211211_21086_H_Caisova_1 | < LOD | | | | < LOD | | | | 43.32 | **58.67** | ± | 16.16 | < LOD | | | | < LOD | | | | < LOD | | | |
| | | 211211_21086_H_Caisova_2 | < LOD | **< LOD** | | | < LOD | **< LOD** | | | 75.53 | ***RSD*** | | ***28 %*** | < LOD | **< LOD** | | | < LOD | **< LOD** | | | < LOD | **< LOD** | | |
| | | 211211_21086_H_Caisova_3 | < LOD | | | | < LOD | | | | 57.17 | | | | < LOD | | | | < LOD | | | | < LOD | | | |
| | Klebsormidium | 211211_21086_H_Caisova_4 | 28.56 | **20.22** | ± | 10.85 | < LOD | | | | < LOD | | | | < LOD | | | | < LOD | | | | 594.00 | **457.66** | ± | 123.05 |
| | | 211211_21086_H_Caisova_5 | 24.15 | ***RSD*** | | ***54 %*** | < LOD | **< LOD** | | | < LOD | **< LOD** | | | < LOD | **< LOD** | | | < LOD | **< LOD** | | | 424.14 | ***RSD*** | | ***27 %*** |
| | | 211211_21086_H_Caisova_6 | 7.95 | | | | < LOD | | | | < LOD | | | | < LOD | | | | < LOD | | | | 354.85 | | | |
| **Media** | Draparnaldia media (with algae) | 211211_21086_H_Caisova_7 | < LOD | | | | < LOD | | | | < LOD | | | | < LOD | | | | < LOD | | | | < LOD | | | |
| | | 211211_21086_H_Caisova_8 | < LOD | **< LOD** | | | < LOD | **< LOD** | | | < LOD | **< LOD** | | | < LOD | **< LOD** | | | < LOD | **< LOD** | | | < LOD | **< LOD** | | |
| | | 211211_21086_H_Caisova_9 | < LOD | | | | < LOD | | | | < LOD | | | | < LOD | | | | < LOD | | | | < LOD | | | |
| | Klebsormidium media (with algae) | 211211_21086_H_Caisova_10 | < LOD | | | | < LOD | | | | < LOD | | | | < LOD | | | | < LOD | | | | < LOD | | | |
| | | 211211_21086_H_Caisova_11 | < LOD | **< LOD** | | | < LOD | **< LOD** | | | < LOD | **< LOD** | | | < LOD | **< LOD** | | | < LOD | **< LOD** | | | < LOD | **< LOD** | | |
| | | 211211_21086_H_Caisova_12 | < LOD | | | | < LOD | | | | < LOD | | | | < LOD | | | | < LOD | | | | < LOD | | | |

| | | | **DL** | | | | **homoDL** | | | | **DS** | | | | **homoDS** | | | | **TY** | | | | **TE** | |
|---|---|---|---|---|---|---|---|---|---|---|---|---|---|---|---|---|---|---|---|---|---|---|---|---|
| **Algae** | Draparnaldia | 211211_21086_H_Caisova_1 | 2.98 | **4.95** | ± | 1.76 | < LOD | | | | < LOD | | | | < LOD | | | | < LOD | | | | < LOD | |
| | | 211211_21086_H_Caisova_2 | 5.54 | ***RSD*** | | ***35 %*** | < LOD | **< LOD** | | | < LOD | **< LOD** | | | < LOD | **< LOD** | | | < LOD | **< LOD** | | | < LOD | **< LOD** |
| | | 211211_21086_H_Caisova_3 | 6.34 | | | | < LOD | | | | < LOD | | | | < LOD | | | | < LOD | | | | < LOD | |
| | Klebsormidium | 211211_21086_H_Caisova_4 | < LOD | | | | 0.20 | **0.16** | ± | 0.09 | < LOD | | | | < LOD | | | | < LOD | | | | < LOD | |
| | | 211211_21086_H_Caisova_5 | < LOD | **< LOD** | | | 0.22 | ***RSD*** | | ***59 %*** | < LOD | **< LOD** | | | < LOD | **< LOD** | | | < LOD | **< LOD** | | | < LOD | **< LOD** |
| | | 211211_21086_H_Caisova_6 | < LOD | | | | 0.05 | | | | < LOD | | | | < LOD | | | | < LOD | | | | < LOD | |
| **Media** | Draparnaldia media (with algae) | 211211_21086_H_Caisova_7 | < LOD | | | | < LOD | | | | < LOD | | | | 41.79 | **20.91** | ± | 18.20 | < LOD | | | | < LOD | |
| | | 211211_21086_H_Caisova_8 | < LOD | **< LOD** | | | < LOD | **< LOD** | | | < LOD | **< LOD** | | | 12.50 | ***RSD*** | | ***87 %*** | *n.d.* | **< LOD** | | | *n.d.* | **< LOD** |
| | | 211211_21086_H_Caisova_9 | < LOD | | | | < LOD | | | | < LOD | | | | 8.43 | | | | < LOD | | | | < LOD | |
| | Klebsormidium media (with algae) | 211211_21086_H_Caisova_10 | < LOD | | | | < LOD | | | | < LOD | | | | < LOD | | | | *n.d.* | | | | *n.d.* | |
| | | 211211_21086_H_Caisova_11 | < LOD | **< LOD** | | | < LOD | **< LOD** | | | < LOD | **< LOD** | | | < LOD | **< LOD** | | | *n.d.* | ***n.d.*** | | | *n.d.* | ***n.d.*** |
| | | 211211_21086_H_Caisova_12 | < LOD | | | | < LOD | | | | < LOD | | | | < LOD | | | | *n.d.* | | | | *n.d.* | |

**Table S8g. Related to Figure 7. Four biological replicates for the detection of auxin in two independent samples of Klebsormidium nitens in medium with or without the algae.**

| | | pmol*g$^{-1}$ FW | | | | | | | | | | | | | | | | | | | | | | | |
|---|---|---|---|---|---|---|---|---|---|---|---|---|---|---|---|---|---|---|---|---|---|---|---|---|---|
| **sample** | **replicate** | **ANT** | | | | **TRP** | | | | **TRA** | | | | **IPyA** | | | | **IAM** | | | | **IAN** | | | |
| **Klebsormidium I.** | klebsormidium_1,5 ml tube_a | 0.61 | **2.91** | ± | 2.30 | 52656.1 | **49,844.7** | ± | 1,630.8 | 67.23 | **59.0** | ± | 5.4 | 558.12 | **543.6** | ± | 25.1 | 4.01 | **3.0** | ± | 0.8 | 51.6 | **50.4** | ± | 12.8 |
| | klebsormidium_1,5 ml tube_b | <LOD | **RSD** | | **79 %** | 49070.9 | **RSD** | | **3 %** | 59.86 | **RSD** | | **9 %** | 507.61 | **RSD** | | **5 %** | <LOD | **RSD** | | **25 %** | 70.5 | **RSD** | | **25 %** |
| | klebsormidium_1,5 ml tube_c | <LOD | | | | 48653.5 | | | | 56.53 | | | | 534.66 | | | | 2.16 | | | | 36.2 | | | |
| | klebsormidium_1,5 ml tube_d | 5.20 | | | | 48998.3 | | | | 52.44 | | | | 574.12 | | | | 2.87 | | | | 43.3 | | | |
| **Klebsormidium II.** | klebsormidium_5 ml tube_a | <LOD | **2.76** | ± | 2.70 | 58820.5 | **58,707.7** | ± | 2,118.7 | 85.92 | **76.5** | ± | 11.6 | 811.88 | **830.5** | ± | 47.2 | 2.49 | ***<LOD*** | | | 43.0 | **41.3** | ± | 1.7 |
| | klebsormidium_5 ml tube_b | 0.23 | **RSD** | | **98 %** | 55329.6 | **RSD** | | **4 %** | 79.46 | **RSD** | | **15 %** | 782.88 | **RSD** | | **6 %** | <LOD | | | | 40.0 | **RSD** | | **4 %** |
| | klebsormidium_5 ml tube_c | 6.50 | | | | 61114.8 | | | | 83.88 | | | | 909.00 | | | | <LOD | | | | 42.8 | | | |
| | klebsormidium_5 ml tube_d | 1.56 | | | | 59566.0 | | | | 56.91 | | | | 818.17 | | | | <LOD | | | | 39.2 | | | |
| | | pmol*ml$^{-1}$ | | | | | | | | | | | | | | | | | | | | | | | |
| **medium with alga** | 3NBBM+3V + alga_media_a | 6.10 | **5.77** | ± | 0.21 | 170.1 | **143.2** | ± | 20.8 | <LOD | ***<LOD*** | | | 0.83 | **1.04** | ± | 0.13 | <LOD | ***<LOD*** | | | <LOD | ***<LOD*** | | |
| | 3NBBM+3V + alga_media_b | 5.52 | **RSD** | | **4 %** | 115.2 | **RSD** | | **15 %** | <LOD | | | | 1.16 | **RSD** | | **12 %** | <LOD | | | | <LOD | | | |
| | 3NBBM+3V + alga_media_c | 5.76 | | | | 133.0 | | | | <LOD | | | | 1.13 | | | | <LOD | | | | <LOD | | | |
| | 3NBBM+3V + alga_media_d | 5.71 | | | | 154.3 | | | | 0.05 | | | | 1.02 | | | | <LOD | | | | <LOD | | | |
| **medium control** | 3NBBM+3V control_media_a | 0.10 | **0.09** | ± | 0.01 | 35.5 | **28.4** | ± | 14.4 | <LOD | ***<LOD*** | | | 0.03 | **0.11** | ± | 0.06 | <LOD | ***<LOD*** | | | <LOD | ***<LOD*** | | |
| | 3NBBM+3V control_media_b | 0.08 | **RSD** | | **15 %** | 41.4 | **RSD** | | **51 %** | 0.28 | | | | 0.10 | **RSD** | | **56 %** | <LOD | | | | <LOD | | | |
| | 3NBBM+3V control_media_c | <LOD | | | | <LOD | | | | <LOD | | | | 0.21 | | | | <LOD | | | | <LOD | | | |
| | 3NBBM+3V control_media_d | <LOD | | | | 8.3 | | | | <LOD | | | | 0.12 | | | | <LOD | | | | <LOD | | | |

| | | pmol*g$^{-1}$ FW | | | | | | | | | | | | | | | | | | | | | |
|---|---|---|---|---|---|---|---|---|---|---|---|---|---|---|---|---|---|---|---|---|---|---|---|
| **sample** | **replicate** | **IAA** | | | | **oxIAA** | | | | **oxIAA-Glc** | | | | **IAA-Glc** | | | | **IAAsp** | | | | **IAGlu** | |
| **Klebsormidium I.** | klebsormidium_1,5 ml tube_a | 8.78 | **10.3** | ± | 1.1 | 0.83 | **1.16** | ± | 0.59 | <LOD | ***<LOD*** | | | <LOD | ***<LOD*** | | | <LOD | ***<LOD*** | | | <LOD | ***<LOD*** |
| | klebsormidium_1,5 ml tube_b | 11.25 | **RSD** | | **11 %** | 0.92 | **RSD** | | **50 %** | <LOD | | | | <LOD | | | | <LOD | | | | <LOD | |
| | klebsormidium_1,5 ml tube_c | 9.82 | | | | 0.74 | | | | <LOD | | | | <LOD | | | | <LOD | | | | <LOD | |
| | klebsormidium_1,5 ml tube_d | 11.53 | | | | 2.17 | | | | <LOD | | | | <LOD | | | | <LOD | | | | <LOD | |
| **Klebsormidium II.** | klebsormidium_5 ml tube_a | 10.23 | **13.7** | ± | 2.4 | 1.08 | **1.24** | ± | 0.11 | <LOD | ***<LOD*** | | | <LOD | ***<LOD*** | | | <LOD | ***<LOD*** | | | <LOD | ***<LOD*** |
| | klebsormidium_5 ml tube_b | 16.59 | **RSD** | | **18 %** | 1.29 | **RSD** | | **9 %** | <LOD | | | | <LOD | | | | <LOD | | | | <LOD | |
| | klebsormidium_5 ml tube_c | 15.24 | | | | 1.39 | | | | <LOD | | | | <LOD | | | | <LOD | | | | <LOD | |
| | klebsormidium_5 ml tube_d | 12.91 | | | | 1.21 | | | | <LOD | | | | <LOD | | | | <LOD | | | | <LOD | |
| | | pmol*ml$^{-1}$ | | | | | | | | | | | | | | | | | | | | | |
| **medium with alga** | 3NBBM+3V + alga_media_a | 0.22 | **0.19** | ± | 0.02 | 0.27 | **0.23** | ± | 0.02 | <LOD | ***<LOD*** | | | <LOD | ***<LOD*** | | | <LOD | ***<LOD*** | | | <LOD | ***<LOD*** |
| | 3NBBM+3V + alga_media_b | 0.17 | **RSD** | | **10 %** | 0.20 | **RSD** | | **10 %** | <LOD | | | | <LOD | | | | <LOD | | | | <LOD | |
| | 3NBBM+3V + alga_media_c | 0.18 | | | | 0.22 | | | | <LOD | | | | <LOD | | | | <LOD | | | | <LOD | |
| | 3NBBM+3V + alga_media_d | 0.18 | | | | 0.23 | | | | <LOD | | | | <LOD | | | | <LOD | | | | <LOD | |
| **medium control** | 3NBBM+3V control_media_a | 0.008 | **0.013** | ± | 0.005 | 0.012 | **0.007** | ± | 0.005 | <LOD | ***<LOD*** | | | <LOD | ***<LOD*** | | | <LOD | ***<LOD*** | | | <LOD | ***<LOD*** |
| | 3NBBM+3V control_media_b | 0.018 | **RSD** | | **37 %** | 0.001 | **RSD** | | **79 %** | <LOD | | | | <LOD | | | | <LOD | | | | <LOD | |
| | 3NBBM+3V control_media_c | <LOD | | | | <LOD | | | | <LOD | | | | <LOD | | | | <LOD | | | | <LOD | |
| | 3NBBM+3V control_media_d | <LOD | | | | <LOD | | | | <LOD | | | | <LOD | | | | <LOD | | | | <LOD | |

| | |
|---|---|
| *NQ* | level not quantified |
| *<LOD* | level below the detection limit |

**Table S8h. Related to Figure 7. Auxin metabolites detected in Klebsormidium nitens samples in medium with or without the algae.**

| | | pmol/g | | | | | | | | | | | |
|---|---|---|---|---|---|---|---|---|---|---|---|---|---|
| **Name** | **Data file** | **ANT** | **TRP** | **TRA** | **IPyA** | **IAN** | **IAM** | **IAA** | **oxIAA** | **IAA-Asp** | **IAA-Glu** | **IAA-Glc** | **oxIAA-Glc** |
| klebsormidium_1,5 ml tube_a | 211104_21074-57.d | 0.61 | 52656.06 | 67.23 | 558.12 | 51.57 | 4.01 | 8.78 | 0.83 | <LOD | <LOD | <LOD | <LOD |
| klebsormidium_1,5 ml tube_b | 211104_21074-58.d | <LOD | 49070.90 | 59.86 | 507.61 | 70.54 | <LOD | 11.25 | 0.92 | <LOD | <LOD | <LOD | <LOD |
| klebsormidium_1,5 ml tube_c | 211104_21074-59.d | <LOD | 48653.46 | 56.53 | 534.66 | 36.20 | 2.16 | 9.82 | 0.74 | <LOD | <LOD | <LOD | <LOD |
| klebsormidium_1,5 ml tube_d | 211104_21074-60.d | 5.20 | 48998.32 | 52.44 | 574.12 | 43.31 | 2.87 | 11.53 | 2.17 | <LOD | <LOD | <LOD | <LOD |
| klebsormidium_5 ml tube_a | 211104_21074-61.d | <LOD | 58820.53 | 85.92 | 811.88 | 43.05 | 2.49 | 10.23 | 1.08 | <LOD | <LOD | <LOD | <LOD |
| klebsormidium_5 ml tube_b | 211104_21074-62.d | 0.23 | 55329.57 | 79.46 | 782.88 | 40.05 | <LOD | 16.59 | 1.29 | <LOD | <LOD | <LOD | <LOD |
| klebsormidium_5 ml tube_c | 211104_21074-63.d | 6.50 | 61114.80 | 83.88 | 909.00 | 42.85 | <LOD | 15.24 | 1.39 | <LOD | <LOD | <LOD | <LOD |
| klebsormidium_5 ml tube_d | 211104_21074-64.d | 1.56 | 59566.00 | 56.91 | 818.17 | 39.18 | <LOD | 12.91 | 1.21 | <LOD | <LOD | <LOD | <LOD |
| 3NBBM+3V + alga_media_a | 211104_21074_m-57.d | 6.10 | 170.07 | <LOD | 0.83 | <LOD | <LOD | 0.22 | 0.27 | <LOD | <LOD | <LOD | <LOD |
| 3NBBM+3V + alga_media_b | 211104_21074_m-58.d | 5.52 | 115.17 | <LOD | 1.16 | <LOD | <LOD | 0.17 | 0.20 | <LOD | <LOD | <LOD | <LOD |
| 3NBBM+3V + alga_media_c | 211104_21074_m-59.d | 5.76 | 133.05 | <LOD | 1.13 | <LOD | <LOD | 0.18 | 0.22 | <LOD | <LOD | <LOD | <LOD |
| 3NBBM+3V + alga_media_d | 211104_21074_m-60.d | 5.71 | 154.34 | 0.05 | 1.02 | <LOD | <LOD | 0.18 | 0.23 | <LOD | <LOD | <LOD | <LOD |
| 3NBBM+3V control_media_a | 211104_21074_m-61.d | 0.10 | 35.52 | <LOD | 0.03 | <LOD | <LOD | 0.01 | 0.01 | <LOD | <LOD | <LOD | <LOD |
| 3NBBM+3V control_media_b | 211104_21074_m-62.d | 0.08 | 41.37 | 0.28 | 0.10 | <LOD | <LOD | 0.02 | 0.00 | <LOD | <LOD | <LOD | <LOD |
| 3NBBM+3V control_media_c | 211104_21074_m-63.d | <LOD | <LOD | <LOD | 0.21 | <LOD | <LOD | <LOD | <LOD | <LOD | <LOD | <LOD | <LOD |
| 3NBBM+3V control_media_d | 211104_21074_m-64.d | <LOD | 8.34 | <LOD | 0.12 | <LOD | <LOD | <LOD | <LOD | <LOD | <LOD | <LOD | <LOD |
| NQ | level not quantified | | | | | | | | | | | | |
| <LOD | level below the detection limit | | | | | | | | | | | | |

**Table S8i. Related to Figure 7. Four biological replicates for the detection of auxin in two independent samples of Draparnaldia in medium with or without the algae.**

| experiment | sample | | ANT | | | | TRP | | | | TRA | | | | IPyA | | | |
|---|---|---|---|---|---|---|---|---|---|---|---|---|---|---|---|---|---|---|
| ***algae*** | | | pmol*g$^{-1}$ FW | | | | | | | | | | | | | | | |
| | Draparnaldia biomass 1 | CC18_a | 22.94 | **25.82** | ± | 3.11 | 44811.9 | **46,890.8** | ± | 6,486.6 | 98.5 | **101.98** | ± | 7.69 | 111.4 | **75.5** | ± | 25.0 |
| | | CC18_b | 27.61 | **RSD** | | **12 %** | 57993.2 | **RSD** | | **14 %** | 105.4 | **RSD** | | **8 %** | 86.3 | **RSD** | | **33 %** |
| | | CC18_c | 30.01 | | | | 42425.0 | | | | 112.3 | | | | 51.2 | | | |
| | | CC18_d | 22.72 | | | | 42333.1 | | | | 91.7 | | | | 53.0 | | | |
| | Draparnaldia biomass 2 | CC21_a | 23.02 | **26.62** | ± | 3.16 | 123727.8 | **118,136.9** | ± | 6,555.0 | 166.2 | **163.96** | ± | 3.86 | 145.4 | **137.9** | ± | 13.3 |
| | | CC21_b | 24.09 | **RSD** | | **12 %** | 113300.6 | **RSD** | | **6 %** | 169.1 | **RSD** | | **2 %** | 147.5 | **RSD** | | **10 %** |
| | | CC21_c | 30.66 | | | | 125406.2 | | | | 160.9 | | | | 143.6 | | | |
| | | CC21_d | 28.71 | | | | 110113.1 | | | | 159.7 | | | | 115.1 | | | |
| ***media*** | | | pmol*ml$^{-1}$ | | | | | | | | | | | | | | | |
| | Draparnaldia medium 1 | media_CC18_a | 100.77 | **98.37** | ± | 2.81 | 359.5 | **407.2** | ± | 123.5 | 0.97 | **0.99** | ± | 0.08 | 2.34 | **1.93** | ± | 0.28 |
| | | media_CC18_b | 101.02 | **RSD** | | **3 %** | 365.2 | **RSD** | | **30 %** | 1.13 | **RSD** | | **9 %** | 2.03 | **RSD** | | **14 %** |
| | | media_CC18_c | 97.60 | | | | 614.8 | | | | 0.95 | | | | 1.74 | | | |
| | | media_CC18_d | 94.11 | | | | 289.2 | | | | 0.90 | | | | 1.63 | | | |
| | Draparnaldia medium 2 | media_CC21_a | 66.23 | **65.88** | ± | 1.81 | 269.2 | **271.0** | ± | 5.6 | 1.67 | **1.60** | ± | 0.07 | 3.26 | **3.05** | ± | 0.24 |
| | | media_CC21_b | 65.18 | **RSD** | | **3 %** | 274.0 | **RSD** | | **2 %** | 1.63 | **RSD** | | **4 %** | 3.12 | **RSD** | | **8 %** |
| | | media_CC21_c | 63.57 | | | | 277.9 | | | | 1.62 | | | | 2.65 | | | |
| | | media_CC21_d | 68.57 | | | | 263.0 | | | | 1.49 | | | | 3.16 | | | |

| experiment | sample | | IAM | | | | IAN | | | | IAA | | | | oxIAA | | | |
|---|---|---|---|---|---|---|---|---|---|---|---|---|---|---|---|---|---|---|
| ***algae*** | | | | | | | | | | | | | | | | | | |
| | Draparnaldia biomass 1 | CC18_a | <LOD | *<LOD* | | | <LOD | *<LOD* | | | 7.45 | **7.65** | ± | 1.62 | <LOD | *<LOD* | | |
| | | CC18_b | <LOD | | | | <LOD | | | | NQ | **RSD** | | **21 %** | <LOD | | | |
| | | CC18_c | <LOD | | | | <LOD | | | | 9.73 | | | | <LOD | | | |
| | | CC18_d | <LOD | | | | <LOD | | | | 5.78 | | | | <LOD | | | |
| | Draparnaldia biomass 2 | CC21_a | <LOD | *<LOD* | | | <LOD | *<LOD* | | | 12.23 | **11.41** | ± | 2.04 | <LOD | *<LOD* | | |
| | | CC21_b | <LOD | | | | <LOD | | | | 14.10 | **RSD** | | **18 %** | <LOD | | | |
| | | CC21_c | <LOD | | | | <LOD | | | | 10.77 | | | | <LOD | | | |
| | | CC21_d | <LOD | | | | <LOD | | | | 8.53 | | | | <LOD | | | |
| ***media*** | | | | | | | | | | | | | | | | | | |
| | Draparnaldia medium 1 | media_CC18_a | 0.08 | **0.08** | ± | 0.00 | <LOD | *<LOD* | | | 7.76 | **7.64** | ± | 0.08 | 1.06 | **1.11** | ± | 0.08 |
| | | media_CC18_b | 0.08 | **RSD** | | **5 %** | <LOD | | | | 7.67 | **RSD** | | **1 %** | 1.03 | **RSD** | | **7 %** |
| | | media_CC18_c | 0.08 | | | | <LOD | | | | 7.55 | | | | 1.25 | | | |
| | | media_CC18_d | 0.09 | | | | <LOD | | | | 7.60 | | | | 1.11 | | | |
| | Draparnaldia medium 2 | media_CC21_a | 0.09 | **0.09** | ± | 0.01 | <LOD | *<LOD* | | | 12.59 | **12.10** | ± | 0.54 | 1.43 | **1.33** | ± | 0.08 |
| | | media_CC21_b | 0.09 | **RSD** | | **14 %** | <LOD | | | | 12.67 | **RSD** | | **4 %** | 1.21 | **RSD** | | **6 %** |
| | | media_CC21_c | 0.10 | | | | <LOD | | | | 11.49 | | | | 1.34 | | | |
| | | media_CC21_d | 0.07 | | | | <LOD | | | | 11.63 | | | | 1.32 | | | |

| experiment | sample | | oxIAA-Glc | | | | IAA-Glc | | | | IAA-Asp | | | | IAA-Glu | |
|---|---|---|---|---|---|---|---|---|---|---|---|---|---|---|---|---|
| ***algae*** | | | | | | | | | | | | | | | | |
| | Draparnaldia biomass 1 | CC18_a | <LOD | *<LOD* | | | <LOD | *<LOD* | | | <LOD | *<LOD* | | | <LOD | *<LOD* |
| | | CC18_b | <LOD | | | | <LOD | | | | <LOD | | | | <LOD | |
| | | CC18_c | <LOD | | | | <LOD | | | | <LOD | | | | <LOD | |
| | | CC18_d | <LOD | | | | <LOD | | | | <LOD | | | | <LOD | |
| | Draparnaldia biomass 2 | CC21_a | <LOD | *<LOD* | | | <LOD | *<LOD* | | | <LOD | *<LOD* | | | <LOD | *<LOD* |
| | | CC21_b | <LOD | | | | <LOD | | | | <LOD | | | | <LOD | |
| | | CC21_c | <LOD | | | | <LOD | | | | <LOD | | | | <LOD | |
| | | CC21_d | <LOD | | | | <LOD | | | | <LOD | | | | <LOD | |
| ***media*** | | | | | | | | | | | | | | | | |
| | Draparnaldia medium 1 | media_CC18_a | <LOD | *<LOD* | | | <LOD | *<LOD* | | | <LOD | *<LOD* | | | <LOD | *<LOD* |
| | | media_CC18_b | <LOD | | | | <LOD | | | | <LOD | | | | <LOD | |
| | | media_CC18_c | <LOD | | | | <LOD | | | | <LOD | | | | <LOD | |
| | | media_CC18_d | <LOD | | | | <LOD | | | | <LOD | | | | <LOD | |
| | Draparnaldia medium 2 | media_CC21_a | <LOD | *<LOD* | | | <LOD | *<LOD* | | | <LOD | *<LOD* | | | <LOD | *<LOD* |
| | | media_CC21_b | <LOD | | | | <LOD | | | | <LOD | | | | <LOD | |
| | | media_CC21_c | <LOD | | | | <LOD | | | | <LOD | | | | <LOD | |
| | | media_CC21_d | <LOD | | | | <LOD | | | | <LOD | | | | <LOD | |

| | |
|---|---|
| *NQ* | level not quantified |
| *<LOD* | level below the detection limit |

**Table S8j. Related to Figure 7. Four biological replicates for the detection of auxin in two independent samples of Draparnaldia in medium with or without the algae.**

| | | pmol/g | | | | | | | | | | | |
|---|---|---|---|---|---|---|---|---|---|---|---|---|---|
| **Name** | **Data file** | **ANT** | **TRP** | **TRA** | **IPyA** | **IAN** | **IAM** | **IAA** | **oxIAA** | **IAA-Asp** | **IAA-Glu** | **IAA-Glc** | **oxIAA-Glc** |
| Draparnaldia biomass 1_a | 220401_22006_17.d | 22.94 | 44811.91 | 98.45 | 111.42 | <LOD | <LOD | 7.45 | <LOD | <LOD | <LOD | <LOD | <LOD |
| Draparnaldia biomass 1_b | 220401_22006_18.d | 27.61 | 57993.19 | 105.42 | 86.27 | <LOD | <LOD | NQ | <LOD | <LOD | <LOD | <LOD | <LOD |
| Draparnaldia biomass 1_c | 220401_22006_19.d | 30.01 | 42424.95 | 112.32 | 51.16 | <LOD | <LOD | 9.73 | <LOD | <LOD | <LOD | <LOD | <LOD |
| Draparnaldia biomass 1_d | 220401_22006_20.d | 22.72 | 42333.08 | 91.73 | 52.96 | <LOD | <LOD | 5.78 | <LOD | <LOD | <LOD | <LOD | <LOD |
| Draparnaldia biomass 2_a | 220401_22006_21.d | 23.02 | 123727.83 | 166.20 | 145.43 | <LOD | <LOD | 12.23 | <LOD | <LOD | <LOD | <LOD | <LOD |
| Draparnaldia biomass 2_b | 220401_22006_22.d | 24.09 | 113300.64 | 169.12 | 147.55 | <LOD | <LOD | 14.10 | <LOD | <LOD | <LOD | <LOD | <LOD |
| Draparnaldia biomass 2_c | 220401_22006_23.d | 30.66 | 125406.25 | 160.88 | 143.57 | <LOD | <LOD | 10.77 | <LOD | <LOD | <LOD | <LOD | <LOD |
| Draparnaldia biomass 2_d | 220401_22006_24.d | 28.71 | 110113.08 | 159.65 | 115.06 | <LOD | <LOD | 8.53 | <LOD | <LOD | <LOD | <LOD | <LOD |
| | | pmol/ml | | | | | | | | | | | |
| **Name** | **Data file** | **ANT** | **TRP** | **TRA** | **IPyA** | **IAN** | **IAM** | **IAA** | **oxIAA** | **IAA-Asp** | **IAA-Glu** | **IAA-Glc** | **oxIAA-Glc** |
| Draparnaldia medium 1_CC | 220401_22006_m_17.d | 100.77 | 359.50 | 0.97 | 2.34 | <LOD | 0.08 | 7.76 | 1.06 | <LOD | <LOD | <LOD | <LOD |
| Draparnaldia medium 1_CC | 220401_22006_m_18.d | 101.02 | 365.17 | 1.13 | 2.03 | <LOD | 0.08 | 7.67 | 1.03 | <LOD | <LOD | <LOD | <LOD |
| Draparnaldia medium 1_CC | 220401_22006_m_19.d | 97.60 | 614.77 | 0.95 | 1.74 | <LOD | 0.08 | 7.55 | 1.25 | <LOD | <LOD | <LOD | <LOD |
| Draparnaldia medium 1_CC | 220401_22006_m_20.d | 94.11 | 289.22 | 0.90 | 1.63 | <LOD | 0.09 | 7.60 | 1.11 | <LOD | <LOD | <LOD | <LOD |
| Draparnaldia medium 2_CC | 220401_22006_m_21.d | 66.23 | 269.15 | 1.67 | 3.26 | <LOD | 0.09 | 12.59 | 1.43 | <LOD | <LOD | <LOD | <LOD |
| Draparnaldia medium 2_CC | 220401_22006_m_22.d | 65.18 | 273.97 | 1.63 | 3.12 | <LOD | 0.09 | 12.67 | 1.21 | <LOD | <LOD | <LOD | <LOD |
| Draparnaldia medium 2_CC | 220401_22006_m_23.d | 63.57 | 277.90 | 1.62 | 2.65 | <LOD | 0.10 | 11.49 | 1.34 | <LOD | <LOD | <LOD | <LOD |
| Draparnaldia medium 2_CC | 220401_22006_m_24.d | 68.57 | 262.98 | 1.49 | 3.16 | <LOD | 0.07 | 11.63 | 1.32 | <LOD | <LOD | <LOD | <LOD |
| NQ | level not quantified | | | | | | | | | | | | |
| <LOD | level below the detection limit | | | | | | | | | | | | |

**Table S8k. Related to Figure 7. List of compounds for the hormonomics detection.**

| Phytohormone family | # | Compound name | Abbreviation |
|---|---|---|---|
| CYTOKININS | 1 | *trans*-zeatin | *t*Z |
| | 2 | *trans*-zeatin riboside | *t*ZR |
| | 3 | *trans*-zeatin-9-glucoside | *t*Z9G |
| | 4 | *trans*-zeatin-7-glucoside | *t*Z7G |
| | 5 | *trans*-zeatin-O-glucoside | *t*ZOG |
| | 6 | *trans*-zeatin riboside-O-glucoside | *t*ZROG |
| | 7 | dihydrozeatin | DHZ |
| | 8 | dihydrozeatin riboside | DHZR |
| | 9 | dihydrozeatin-9-glucoside | DHZ9G |
| | 10 | dihydrozeatin-7-glucoside | DHZ7G |
| | 11 | dihydrozeatin-O-glucoside | DHZOG |
| | 12 | dihydrozeatin riboside-O-glucoside | DHZROG |
| | 13 | *cis*-zeatin | *c*Z |
| | 14 | *cis*-zeatin riboside | *c*ZR |
| | 15 | *cis*-zeatin-9-glucoside | *c*Z9G |
| | 16 | *cis*-zeatin-O-glucoside | *c*ZOG |
| | 17 | *cis*-zeatin riboside-O-glucoside | cZROG |
| | 18 | isopentenyladenine | iP |
| | 19 | isopentenyladenine riboside | iPR |
| | 20 | isopentenyladenine-9-glucoside | iP9G |
| | 21 | isopentenyladenine-7-glucoside | iP7G |
| | 22 | *para*-topolin | pT |
| | 23 | *para*-topolin riboside | pTR |
| | 24 | *para*-topolin-9-glucoside | pT9G |
| | 25 | *meta*-topolin | mT |
| | 26 | *meta*topolin riboside | mTR |
| | 27 | *meta*topolin-9-glucoside | mT9G |
| | 28 | *ortho*-topolin | oT |
| | 29 | *ortho*-topolin riboside | oTR |
| | 30 | *ortho*-topolin-9-glucoside | oT9G |
| | 31 | benzyladenine | BAP |
| | 32 | benzyladenine riboside | BAPR |
| | 33 | benzyladenine-9-glucoside | BAP9G |
| | 34 | benzyladenine-7-glucoside | BAP7G |
| | 35 | kinetin | K |
| | 36 | kinetin riboside | KR |
| | 37 | kinetin-9-glucoside | K9G |
| 2-METHYLTHIO CYTOKININS | 38 | 2-methylthio-isopentenyladenine | 2MeSiP |
| | 39 | 2-methylthio-isopentenyladenine riboside | 2MeSiPR |
| | 40 | 2-methylthio-*cis*-zeatin | 2MeScZ |
| | 41 | 2-methylthio-*cis*-zeatin riboside | 2MeScZR |
| JASMONATES | 57 | jasmonic acid | JA |
| | 58 | JA-isoleucine | JA-Ile |
| | 59 | *cis*-12-oxo-phytodienoic acid | *cis*OPDA |
| | 60 | dinor-12-oxo-phytodienoic acid | dnOPDA |
| | 61 | JA-tryptophan | JA-Trp |
| | 62 | JA-phenylalanine | JA-Phe |
| | 63 | JA-valine | JA-Val |
| | 64 | 3-oxo-2-(2-(Z)-pentenyl)cyclopentane-1-hexanoic acid | OPC-6 |
| | 65 | 12-hydroxy-jasmonic acid | 12-OH-JA |
| | 66 | 9,10-dihydrojasmonic acid | 9,10-dh-JA |
| | 67 | 3-oxo-2-(2-(Z)-pentenyl)cyclopentane-1-butyric acid | OPC-4 |
| SALICYLIC ACID | 68 | salicylic acid | SA |
| ABSCISATES | 69 | abscisic acid | ABA |
| | 70 | dihydrophaseic acid | DPA |
| | 71 | *neo*-phaseic acid | NeoPA |
| | 72 | phaseic acid | PA |
| | 73 | 7-hydroxy-ABA | 7-OH-ABA |
| GIBBERELLINS | 74 | Gibberellin 8 | GA 8 |
| | 75 | Gibberellin 29 | GA 29 |
| | 76 | Gibberellin 3 | GA 3 |
| | 77 | Gibberellin 1 | GA 1 |
| | 78 | Gibberellin 6 | GA 6 |
| | 79 | Gibberellin 5 | GA 5 |
| | 80 | Gibberellin 19 | GA 19 |
| | 81 | Gibberellin 24 | GA 24 |
| | 82 | Gibberellin 44 | GA 44 |
| | 83 | Gibberellin 34 | GA 34 |
| | 84 | Gibberellin 51 | GA 51 |
| | 85 | Gibberellin 53 | GA 53 |
| | 86 | Gibberellin 4 | GA 4 |
| | 87 | Gibberellin 15 | GA 15 |
| BRASSINOSTEROIDS | 88 | 28-*nor*castasterone | norCS |
| | 89 | dolichosterone | DS |
| | 90 | castasterone | CS |
| | 91 | 24-*epi*castasterone | epiCS |
| | 92 | 28-*nor*brassinolide | norBL |
| | 93 | homodolichosterone | homoDS |
| | 94 | dolicholide | DL |
| | 95 | brassinolide | BL |
| | 96 | 24-*epi*brassinolide | epiBL |
| | 97 | homodolicholide | homoDL |
| | 98 | homobrassinolide | homoBL |
| | 99 | homocastasteron | homoCS |
| | 100 | teasterone | TE |
| | 101 | typhasterol | TY |
| | 102 | anthranilate | ANT |
| | 103 | L-Tryptophan | TRP |
| | 104 | tryptamine | TRA |
| | 105 | indole-3-pyruvic acid | IPyA |
| | 106 | indole-3-acetamide | IAM |
| | 107 | indole-3-acetonitrile | IAN |
| | 108 | indole-3-acetic acid, IAA | IAA |
| | 109 | 2-oxindole-3-acetic acid | oxIAA |
| | 110 | IAA-aspartate | IAAsp |
| | 111 | IAA-glutamate | IAGlu |

**Table S8I. Related to Figure 7. Gene copies encoding members of the synthesis, signalling and transport networks of the phytohormones in the different green species used in this study**

| | gene | TAIR ID | phytohormone | Pfam domains | phylogeny_confirmed | Arabidopsis_thaliana | Oryza_sativa | Selaginella_moellendorffii | Physcomitrium_patens | Marchantia_polymorpha | Mesotaenium_endlicherianum | Spirogloea_muscicola | Chara_braunii | Klebsormidium_nitens | Chlorokybus_atmophyticus | Mesostigma_viride | Draparnaldia | Chlamydomonas_reinhardtii | Volvox_carteri | Picochlorum_SENEW3 | Auxenochlorella_protothecoides | Coccomyxa_obi | Chloropicon_primus | Micromonas_RCC299 | Ostreococcus_tauri | Bathycoccus_prasinos | Cyanophora_paradoxa | Chondrus_crispus | Galdieria_sulphuraria | Cyanidioschyzon_merolae |
|---|---|---|---|---|---|---|---|---|---|---|---|---|---|---|---|---|---|---|---|---|---|---|---|---|---|---|---|---|---|---|
| 1 | PSY | AT5G17230 | ABA | SQS_PSY | NO | 1 | 4 | 1 | 3 | 1 | 2 | 3 | 2 | 1 | 1 | 1 | 1 | 1 | 1 | 1 | 0 | 1 | 1 | 2 | 2 | 1 | 1 | 0 | 1 | 1 |
| 2 | PDS | AT4G14210 | ABA | Amino_oxidase | NO | 2 | 4 | 2 | 4 | 2 | 2 | 0 | 1 | 2 | 1 | 1 | 1 | 2 | 2 | 2 | 2 | 2 | 2 | 2 | 2 | 2 | 1 | 2 | 1 | 2 |
| 3 | LUT5 | AT1G31800 | ABA | p450 | YES | 1 | 3 | 1 | 1 | 1 | 1 | 3 | 1 | 1 | 1 | 1 | 1 | 2 | 1 | 1 | 1 | 1 | 0 | 1 | 1 | 1 | 0 | 0 | 0 | 0 |
| 4 | ZEP | AT5G67030 | ABA | Not checked | NO | 1 | 2 | 0 | 2 | 1 | 1 | 0 | 0 | 0 | 1 | 0 | 0 | 1 | 1 | 0 | 1 | 0 | 0 | 0 | 0 | 0 | 0 | 0 | 0 | 0 |
| 5 | ABA4 | AT1G67080 | ABA | ABA4-like | NO | 1 | 1 | 1 | 2 | 1 | 1 | 3 | 0 | 1 | 0 | 1 | 2 | 0 | 0 | 1 | 0 | 2 | 1 | 2 | 0 | 2 | 0 | 0 | 0 | 0 |
| 6 | NCED3 | AT3G14440 | ABA | RPE65 | NO | 7 | 6 | 12 | 7 | 3 | 1 | 2 | 0 | 2 | 1 | 7 | 1 | 0 | 1 | 0 | 0 | 2 | 1 | 0 | 0 | 0 | 4 | 0 | 0 | 0 |
| 7 | ABA2 | AT1G52340 | ABA | adh_short, adh_short_C2, KR | NO | 11 | 32 | 7 | 4 | 13 | 0 | 2 | 0 | 4 | 2 | 3 | 0 | 1 | 0 | 1 | 1 | 3 | 1 | 0 | 0 | 0 | 2 | 0 | 2 | 0 |
| 8 | ABA3 | AT1G16540 | ABA | Aminotran_5, MOSC | NO | 1 | 1 | 1 | 1 | 1 | 0 | 3 | 1 | 1 | 1 | 2 | 1 | 1 | 1 | 0 | 1 | 1 | 1 | 0 | 0 | 0 | 0 | 0 | 0 | 0 |
| 9 | AAO3 | AT2G27150 | ABA | Not checked | NO | 4 | 10 | 4 | 3 | 2 | 1 | 2 | 0 | 2 | 0 | 3 | 0 | 0 | 1 | 0 | 0 | 2 | 0 | 0 | 0 | 0 | 0 | 0 | 0 | 0 |
| 10 | GPA1 | AT2G26300 | ABA | Arf, G-alpha | NO | 1 | 3 | 1 | 0 | 1 | 0 | 0 | 1 | 1 | 0 | 0 | 0 | 0 | 0 | 0 | 0 | 0 | 0 | 0 | 0 | 0 | 0 | 0 | 0 | 0 |
| 11 | AGB1 | AT4G34460 | ABA | WD40 | NO | 1 | 1 | 1 | 2 | 1 | 0 | 0 | 2 | 1 | 0 | 0 | 0 | 0 | 0 | 0 | 0 | 0 | 0 | 0 | 0 | 0 | 0 | 0 | 0 | 0 |
| 12 | CPK3 | AT4G23650 | ABA | Not checked | YES | 12 | 11 | 4 | 6 | 1 | 1 | 2 | 2 | 0 | 0 | 0 | 0 | 0 | 0 | 0 | 0 | 0 | 0 | 0 | 0 | 0 | 0 | 0 | 0 | 0 |
| 13 | CIPK | AT3G17510 | ABA | Not checked | YES | 10 | 18 | 5 | 8 | 2 | 1 | 0 | 0 | 1 | 1 | 1 | 0 | 0 | 0 | 1 | 1 | 1 | 0 | 0 | 0 | 0 | 1 | 8 | 0 | 0 |
| 14 | ABCG30 | AT4G15230 | ABA | Not checked | YES | 9 | 11 | 3 | 10 | 8 | 0 | 0 | 0 | 0 | 0 | 0 | 0 | 0 | 0 | 0 | 0 | 0 | 0 | 0 | 0 | 0 | 0 | 0 | 0 | 0 |
| 15 | ABCG31 | AT2G29940 | ABA | Not checked | YES | 1 | 1 | 3 | 3 | 2 | 0 | 0 | 0 | 0 | 0 | 0 | 0 | 0 | 0 | 0 | 0 | 0 | 0 | 0 | 0 | 0 | 0 | 0 | 0 | 0 |
| 16 | ABCG40 | AT1G15520 | ABA | Not checked | YES | 5 | 26 | 1 | 0 | 0 | 0 | 0 | 0 | 0 | 0 | 0 | 0 | 0 | 0 | 0 | 0 | 0 | 0 | 0 | 0 | 0 | 0 | 0 | 0 | 0 |
| 17 | DTX50 | AT5G52050 | ABA | MatE | NO | 25 | 52 | 12 | 12 | 12 | 2 | 1 | 14 | 15 | 0 | 0 | 3 | 0 | 0 | 1 | 0 | 1 | 0 | 0 | 0 | 0 | 0 | 0 | 0 | 0 |
| 18 | GTG1 | AT1G64990 | ABA | ABA_GPCR, GPHR_N | NO | 2 | 1 | 1 | 0 | 1 | 1 | 3 | 1 | 1 | 1 | 0 | 1 | 2 | 1 | 1 | 1 | 1 | 0 | 1 | 1 | 1 | 0 | 0 | 0 | 0 |
| 19 | PYR1/PYL1 | AT4G17870, AT5G46790 | ABA | Polyketide_cyc2 | NO | 14 | 20 | 5 | 5 | 4 | 1 | 0 | 0 | 0 | 0 | 0 | 0 | 0 | 0 | 0 | 0 | 0 | 0 | 0 | 0 | 0 | 0 | 0 | 0 | 0 |
| 20 | PP2C | AT4G26080 | ABA | PP2C | NO | 21 | 27 | 8 | 8 | 5 | 2 | 7 | 0 | 4 | 1 | 2 | 5 | 2 | 3 | 1 | 2 | 1 | 0 | 0 | 1 | 0 | 0 | 1 | 0 | 1 |
| 21 | SnRK2.2 | AT3G50500 | ABA | Not checked | YES | 10 | 13 | 3 | 4 | 2 | 1 | 8 | 2 | 5 | 2 | 3 | 23 | 12 | 8 | 11 | 7 | 19 | 4 | 1 | 2 | 2 | 2 | 0 | 0 | 0 |
| 22 | ABF | AT1G49720 | ABA | bZIP_1, bZIP_2 | NO | 5 | 4 | 1 | 0 | 1 | 0 | 0 | 2 | 0 | 0 | 1 | 0 | 0 | 0 | 0 | 0 | 0 | 0 | 0 | 0 | 0 | 0 | 0 | 0 | 0 |
| 23 | ABCG25/14 | AT1G71960, AT1G31770 | ABA, CK | Not checked | YES | 6 | 7 | 2 | 4 | 4 | 3 | 3 | 0 | 3 | 1 | 2 | 3 | 0 | 0 | 1 | 1 | 1 | 3 | 0 | 0 | 0 | 1 | 1 | 2 | 0 |
| 24 | NPF4 | AT1G69850, AT3G25260 | ABA, JA | PTR2 | YES | 7 | 15 | 5 | 3 | 1 | 0 | 0 | 0 | 0 | 0 | 0 | 0 | 0 | 0 | 0 | 0 | 0 | 0 | 0 | 0 | 0 | 0 | 0 | 0 | 0 |
| 25 | TAA1/TAR1/TAR2 | AT1G70560, AT1G23320, AT4G24670 | AUX | Alliinase_C, Aminotran_1_2 | NO | 5 | 3 | 2 | 5 | 1 | 0 | 0 | 0 | 1 | 0 | 0 | 0 | 0 | 0 | 0 | 0 | 0 | 0 | 0 | 0 | 0 | 0 | 0 | 0 | 0 |
| 26 | TIR1 | AT3G62980 | AUX | F-box | YES | 4 | 3 | 1 | 5 | 1 | 0 | 0 | 0 | 0 | 0 | 0 | 0 | 0 | 0 | 0 | 0 | 0 | 0 | 0 | 0 | 0 | 0 | 0 | 0 | 0 |
| 27 | UGT84B1 | AT2G23260 | AUX | UDGPT | YES | 7 | 12 | 0 | 0 | 0 | 0 | 0 | 0 | 0 | 0 | 0 | 0 | 0 | 0 | 0 | 0 | 0 | 0 | 0 | 0 | 0 | 0 | 0 | 0 | 0 |
| 28 | DAO | AT1G14120 | AUX | 2OG-FeII_Oxy, DIOX_N | YES | 3 | 2 | 1 | 0 | 0 | 0 | 0 | 0 | 0 | 0 | 0 | 0 | 0 | 0 | 0 | 0 | 0 | 0 | 0 | 0 | 0 | 0 | 0 | 0 | 0 |
| 29 | UGT74D1 | AT2G31750 | AUX | UDGPT | YES | 10 | 6 | 0 | 0 | 0 | 0 | 0 | 0 | 0 | 0 | 0 | 0 | 0 | 0 | 0 | 0 | 0 | 0 | 0 | 0 | 0 | 0 | 0 | 0 | 0 |
| 30 | ARR | AT3G16857 | AUX | Myb_DNA-binding, Response_reg | NO | 1 | 1 | 0 | 0 | 0 | 1 | 2 | 0 | 0 | 0 | 0 | 0 | 0 | 0 | 0 | 0 | 0 | 0 | 0 | 0 | 0 | 0 | 0 | 0 | 0 |
| 31 | TMK | AT1G66150 | AUX | LRR, Pkinase, PK_Tyr_Ser-Thr | YES | 10 | 23 | 4 | 15 | 16 | 1 | 5 | 3 | 1 | 0 | 0 | 0 | 0 | 0 | 0 | 0 | 0 | 0 | 0 | 0 | 0 | 0 | 0 | 0 | 0 |
| 32 | ABP1 | AT4G02980 | AUX | Auxin_BP, Cupin_2 | NO | 1 | 2 | 1 | 2 | 0 | 1 | 5 | 1 | 1 | 0 | 0 | 0 | 0 | 0 | 0 | 0 | 0 | 0 | 0 | 0 | 0 | 0 | 0 | 0 | 0 |
| 33 | YUCCA1 | AT4G32540 | AUX | Not checked | NO | 11 | 22 | 3 | 6 | 4 | 0 | 0 | 0 | 0 | 0 | 0 | 0 | 0 | 0 | 0 | 0 | 0 | 0 | 0 | 0 | 0 | 0 | 0 | 0 | 0 |
| 34 | GH3.1 | AT2G14960 | AUX | GH3 | YES | 17 | 16 | 9 | 1 | 2 | 0 | 0 | 0 | 3 | 0 | 0 | 0 | 0 | 0 | 0 | 0 | 0 | 0 | 0 | 0 | 0 | 0 | 0 | 0 | 0 |
| 35 | ILR1 | AT3G02875 | AUX | M20_dimer, Peptidase_M20 | NO | 7 | 9 | 3 | 3 | 1 | 1 | 5 | 0 | 2 | 1 | 1 | 1 | 2 | 1 | 1 | 1 | 2 | 1 | 2 | 1 | 1 | 1 | 0 | 0 | 0 |
| 36 | PILS1 | AT1G20925 | AUX | Mem_trans | NO | 7 | 6 | 4 | 6 | 3 | 1 | 4 | 0 | 2 | 0 | 0 | 1 | 1 | 0 | 0 | 1 | 1 | 0 | 0 | 0 | 0 | 0 | 0 | 0 | 0 |
| 37 | PIN1 | AT1G73590 | AUX | Mem_trans | NO | 7 | 13 | 6 | 4 | 5 | 1 | 3 | 1 | 1 | 0 | 1 | 0 | 0 | 0 | 0 | 0 | 0 | 0 | 0 | 0 | 0 | 0 | 0 | 0 | 0 |
| 38 | AUX1/LAX1 | AT2G38120, AT5G01240 | AUX | Aa_trans | NO | 4 | 6 | 19 | 7 | 1 | 0 | 0 | 0 | 1 | 0 | 0 | 0 | 0 | 0 | 0 | 0 | 3 | 0 | 0 | 0 | 0 | 0 | 0 | 0 | 0 |
| 39 | ABCB1 | AT2G36910 | AUX | AAA_16, ABC_membrane, ABC_tran, SMC_N | NO | 6 | 8 | 10 | 4 | 5 | 3 | 3 | 3 | 4 | 0 | 1 | 0 | 0 | 0 | 0 | 0 | 1 | 2 | 0 | 0 | 1 | 4 | 0 | 0 | 0 |
| 40 | ARF | AT1G59750 | AUX | Auxin_resp, AUX_IAA, B3 | NO | 18 | 36 | 6 | 11 | 3 | 0 | 1 | 0 | 0 | 1 | 0 | 0 | 0 | 0 | 0 | 0 | 0 | 0 | 0 | 0 | 0 | 0 | 0 | 0 | 0 |
| 41 | DET2 | AT2G38050 | BR | DUF1295, Steroid_dh | NO | 1 | 3 | 2 | 1 | 1 | 0 | 0 | 0 | 1 | 1 | 0 | 1 | 1 | 1 | 1 | 1 | 0 | 0 | 0 | 0 | 0 | 0 | 0 | 1 | 1 |
| 42 | CPD | AT5G05690 | BR | p450 | YES | 2 | 2 | 0 | 0 | 0 | 0 | 0 | 0 | 0 | 0 | 0 | 0 | 0 | 0 | 0 | 0 | 0 | 0 | 0 | 0 | 0 | 0 | 0 | 0 | 0 |
| 43 | BIM2 | AT1G69010 | BR | HLH | NO | 3 | 7 | 1 | 0 | 1 | 0 | 0 | 0 | 0 | 0 | 0 | 0 | 0 | 0 | 0 | 0 | 0 | 0 | 0 | 0 | 0 | 0 | 0 | 0 | 0 |
| 44 | BKI1 | AT5G42750 | BR | Not checked | NO | 1 | 0 | 0 | 0 | 0 | 0 | 0 | 0 | 0 | 0 | 0 | 0 | 0 | 0 | 0 | 0 | 0 | 0 | 0 | 0 | 0 | 0 | 0 | 0 | 0 |
| 45 | BSK1 | AT4G35230 | BR | Pkinase, PK_Tyr_Ser-Thr | YES | 31 | 31 | 9 | 12 | 6 | 3 | 4 | 2 | 2 | 0 | 0 | 0 | 0 | 0 | 0 | 0 | 0 | 0 | 0 | 0 | 0 | 0 | 0 | 0 | 0 |
| 46 | DWF4 | AT3G50660 | BR | p450 | YES | 2 | 2 | 1 | 0 | 0 | 0 | 0 | 0 | 0 | 0 | 0 | 0 | 0 | 0 | 0 | 0 | 0 | 0 | 0 | 0 | 0 | 0 | 0 | 0 | 0 |
| 47 | CYP85A2 | AT3G30180 | BR | p450 | YES | 6 | 12 | 0 | 0 | 0 | 0 | 0 | 0 | 0 | 0 | 0 | 0 | 0 | 0 | 0 | 0 | 0 | 0 | 0 | 0 | 0 | 0 | 0 | 0 | 0 |
| 48 | BRI1/BIN1 | AT4G39400 | BR | LRR, Pkinase, PK_Tyr_Ser-Thr | YES | 16 | 24 | 10 | 11 | 5 | 1 | 0 | 1 | 0 | 0 | 0 | 0 | 0 | 0 | 0 | 0 | 0 | 0 | 0 | 0 | 0 | 0 | 0 | 0 | 0 |
| 49 | BAK1 | AT4G33430 | BR | LRR, Pkinase, PK_Tyr_Ser-Thr | YES | 14 | 20 | 4 | 7 | 3 | 0 | 4 | 0 | 0 | 0 | 0 | 0 | 0 | 0 | 0 | 0 | 0 | 0 | 0 | 0 | 0 | 0 | 0 | 0 | 0 |
| 50 | BSU1 | AT1G03445 | BR | Kelch | NO | 4 | 4 | 2 | 5 | 2 | 1 | 3 | 1 | 1 | 1 | 5 | 1 | 2 | 1 | 1 | 1 | 1 | 1 | 1 | 1 | 1 | 0 | 0 | 0 | 0 |
| 51 | BES1/BZR1 | AT1G19350, AT1G75080 | BR | BES1_N | NO | 6 | 2 | 1 | 0 | 0 | 0 | 0 | 0 | 0 | 0 | 0 | 0 | 0 | 0 | 0 | 0 | 0 | 0 | 0 | 0 | 0 | 0 | 0 | 0 | 0 |
| 52 | BIN2 | AT4G18710 | BR | Pkinase, PK_Tyr_Ser-Thr | YES | 10 | 11 | 2 | 7 | 1 | 0 | 3 | 2 | 2 | 1 | 3 | 2 | 8 | 4 | 1 | 1 | 2 | 3 | 3 | 1 | 1 | 5 | 0 | 1 | 1 |
| 53 | TCP1 | AT1G67260 | BR | TCP | NO | 1 | 0 | 0 | 0 | 0 | 0 | 0 | 0 | 0 | 0 | 0 | 0 | 0 | 0 | 0 | 0 | 0 | 0 | 0 | 0 | 0 | 0 | 0 | 0 | 0 |
| 54 | AHK4/CHK | AT2G01830 | CK | CHASE, HATPase_c, HisKA, Response_reg | NO | 3 | 6 | 2 | 5 | 1 | 2 | 7 | 1 | 4 | 0 | 0 | 0 | 0 | 0 | 0 | 0 | 0 | 0 | 0 | 0 | 0 | 0 | 0 | 0 | 0 |
| 55 | CYP735A | AT1G67110 | CK | p450 | YES | 19 | 38 | 37 | 6 | 5 | 2 | 5 | 1 | 6 | 2 | 1 | 1 | 1 | 1 | 1 | 0 | 1 | 1 | 2 | 2 | 2 | 0 | 2 | 0 | 0 |
| 56 | HPT | AT3G54720 | CK | PA, Peptidase_M28, TFR_dimer | NO | 2 | 3 | 6 | 4 | 1 | 0 | 2 | 0 | 1 | 1 | 1 | 0 | 0 | 0 | 0 | 0 | 0 | 0 | 0 | 0 | 0 | 2 | 0 | 0 | 0 |
| 57 | IPT | AT1G68460 | CK | IPT | NO | 8 | 10 | 0 | 3 | 0 | 0 | 0 | 0 | 0 | 0 | 0 | 0 | 0 | 0 | 0 | 0 | 0 | 0 | 0 | 0 | 0 | 0 | 0 | 0 | 0 |
| 58 | LOG | AT2G28305 | CK | LDcluster4, Lysine_decarbox | NO | 8 | 11 | 2 | 9 | 1 | 1 | 3 | 1 | 1 | 1 | 0 | 1 | 1 | 0 | 1 | 0 | 1 | 1 | 1 | 1 | 1 | 1 | 1 | 0 | 2 |
| 59 | ENT3/6/7 | AT4G05120, AT4G05110, AT1G61630 | CK | Nucleoside_tran | NO | 7 | 8 | 4 | 4 | 3 | 0 | 0 | 1 | 0 | 1 | 1 | 0 | 0 | 0 | 0 | 0 | 0 | 0 | 0 | 0 | 0 | 0 | 0 | 0 | 0 |
| 60 | PUP1/2/14 | AT1G28230, AT2G33750, AT1G19770 | CK | PUNUT | NO | 21 | 19 | 6 | 0 | 1 | 0 | 0 | 0 | 0 | 0 | 0 | 0 | 0 | 0 | 0 | 0 | 0 | 0 | 0 | 0 | 0 | 0 | 0 | 0 | 0 |
| 61 | RRA | HMMsearch of Pfam domain PF00072 | CK | Response_reg | YES | 10 | 18 | 2 | 7 | 1 | 1 | 0 | 0 | 1 | 0 | 0 | 1 | 1 | 0 | 1 | 1 | 0 | 0 | 1 | 1 | 1 | 1 | 0 | 0 | 0 |
| 62 | RRB | HMMsearch of Pfam domain PF00072 and PF00249 | CK | Response_reg (PF00072), Myb_DNA-binding (PF00249) | YES | 7 | 9 | 5 | 5 | 1 | 2 | 6 | 0 | 1 | 1 | 1 | 1 | 0 | 0 | 0 | 0 | 1 | 0 | 1 | 1 | 1 | 0 | 0 | 0 | 0 |
| 63 | ACS1 | AT3G61510 | ETH | Aminotran_1_2 | NO | 11 | 6 | 2 | 2 | 2 | 2 | 4 | 2 | 1 | 0 | 0 | 1 | 0 | 0 | 1 | 1 | 0 | 2 | 1 | 0 | 0 | 2 | 0 | 1 | 0 |
| 64 | ACO1 | AT2G19590 | ETH | 2OG-FeII_Oxy, DIOX_N | YES | 5 | 9 | 0 | 0 | 2 | 0 | 0 | 0 | 0 | 0 | 0 | 0 | 0 | 0 | 0 | 0 | 0 | 0 | 0 | 0 | 0 | 0 | 0 | 0 | 0 |
| 65 | CTR1 | AT5G03730 | ETH | EDR1, Pkinase, PK_Tyr_Ser-Thr | NO | 6 | 11 | 2 | 1 | 3 | 2 | 8 | 2 | 1 | 1 | 1 | 1 | 1 | 1 | 0 | 0 | 1 | 1 | 0 | 1 | 1 | 0 | 0 | 0 | 0 |
| 66 | EBF1 | AT2G25490 | ETH | F-box | NO | 2 | 2 | 3 | 2 | 1 | 0 | 0 | 1 | 0 | 0 | 0 | 0 | 0 | 0 | 0 | 0 | 0 | 0 | 0 | 0 | 0 | 0 | 0 | 0 | 0 |
| 67 | LHT1 | AT5G40780 | ETH | Aa_trans, Trp_Tyr_perm | NO | 7 | 4 | 6 | 3 | 1 | 0 | 0 | 0 | 0 | 0 | 0 | 0 | 0 | 0 | 1 | 0 | 0 | 0 | 0 | 0 | 0 | 0 | 0 | 0 | 0 |
| 68 | ETR1 | AT1G66340 | ETH | GAF, HATPase_c, HisKA, Response_reg | NO | 2 | 3 | 2 | 7 | 3 | 4 | 10 | 0 | 6 | 0 | 0 | 0 | 0 | 0 | 0 | 0 | 0 | 0 | 0 | 0 | 0 | 2 | 0 | 0 | 0 |
| 69 | EIN2 | AT5G03280 | ETH | Nramp | NO | 1 | 7 | 0 | 1 | 1 | 0 | 0 | 0 | 0 | 0 | 0 | 0 | 0 | 0 | 0 | 0 | 0 | 0 | 0 | 0 | 0 | 0 | 0 | 0 | 0 |
| 70 | EIN3 | AT3G20770 | ETH | EIN3 | NO | 4 | 3 | 0 | 2 | 1 | 1 | 0 | 1 | 1 | 0 | 0 | 0 | 0 | 0 | 0 | 0 | 0 | 0 | 0 | 0 | 0 | 0 | 0 | 0 | 0 |
| 71 | CPS | AT4G02780 | GA | Terpene_synth, Terpene_synth_C | YES | 1 | 4 | 11 | 2 | 5 | 0 | 0 | 0 | 0 | 0 | 0 | 0 | 0 | 0 | 0 | 0 | 0 | 0 | 0 | 0 | 0 | 0 | 0 | 0 | 0 |
| 72 | KS | AT1G79460 | GA | Terpene_synth, Terpene_synth_C | YES | 2 | 25 | 1 | 1 | 2 | 0 | 0 | 0 | 0 | 0 | 0 | 0 | 0 | 0 | 0 | 0 | 0 | 0 | 0 | 0 | 0 | 0 | 0 | 0 | 0 |
| 73 | KO1 | AT5G25900 | GA | p450 | NO | 3 | 12 | 5 | 2 | 7 | 0 | 0 | 0 | 0 | 0 | 0 | 0 | 0 | 0 | 0 | 0 | 0 | 0 | 0 | 0 | 0 | 0 | 0 | 0 | 0 |
| 74 | GA20ox3 | AT5G07200 | GA | 2OG-FeII_Oxy, DIOX_N | YES | 5 | 4 | 1 | 0 | 0 | 0 | 0 | 0 | 0 | 0 | 0 | 0 | 0 | 0 | 0 | 0 | 0 | 0 | 0 | 0 | 0 | 0 | 0 | 0 | 0 |
| 75 | GA3ox1 | AT1G15550 | GA | 2OG-FeII_Oxy, DIOX_N | YES | 4 | 2 | 1 | 13 | 2 | 0 | 0 | 0 | 0 | 0 | 0 | 0 | 0 | 0 | 0 | 0 | 0 | 0 | 0 | 0 | 0 | 0 | 0 | 0 | 0 |
| 76 | RGA1/DELLA | AT2G01570 | GA | DELLA, GRAS | NO | 17 | 39 | 18 | 28 | 7 | 1 | 8 | 0 | 0 | 0 | 0 | 0 | 0 | 0 | 0 | 0 | 0 | 0 | 0 | 0 | 0 | 0 | 0 | 0 | 0 |
| 77 | KAO | AT1G05160 | GA | p450 | YES | 2 | 3 | 2 | 0 | 3 | 0 | 0 | 0 | 0 | 0 | 0 | 0 | 0 | 0 | 0 | 0 | 0 | 0 | 0 | 0 | 0 | 0 | 0 | 0 | 0 |
| 78 | SLY1/GID2 | AT4G24210 | GA | F-box, F-box-like | NO | 2 | 2 | 1 | 0 | 1 | 0 | 0 | 0 | 0 | 0 | 0 | 0 | 0 | 0 | 0 | 0 | 0 | 0 | 0 | 0 | 0 | 0 | 0 | 0 | 0 |
| 79 | SLR | AT4G14550 | GA | AUX_IAA | NO | 20 | 39 | 4 | 2 | 1 | 0 | 0 | 1 | 0 | 0 | 0 | 0 | 0 | 0 | 0 | 0 | 0 | 0 | 0 | 0 | 0 | 0 | 0 | 0 | 0 |
| 80 | GID1 | AT3G05120 | GA | Abhydrolase_3 | NO | 7 | 3 | 9 | 2 | 1 | 0 | 0 | 0 | 0 | 0 | 0 | 0 | 0 | 0 | 0 | 0 | 0 | 0 | 0 | 0 | 0 | 0 | 0 | 0 | 0 |
| 81 | SWEET13/14 | AT5G50800, AT4G25010 | GA | MtN3_slv | NO | 17 | 22 | 16 | 6 | 9 | 1 | 4 | 2 | 2 | 1 | 0 | 0 | 0 | 0 | 0 | 0 | 0 | 1 | 0 | 0 | 1 | 0 | 0 | 0 | 0 |
| 82 | AOS | AT5G42650 | JA | p450 | NO | 2 | 5 | 11 | 3 | 2 | 0 | 2 | 0 | 1 | 0 | 0 | 0 | 0 | 0 | 0 | 0 | 0 | 0 | 0 | 0 | 0 | 0 | 0 | 0 | 0 |
| 83 | AOC | AT1G13280 | JA | Allene_ox_cyc | NO | 4 | 1 | 1 | 5 | 1 | 0 | 0 | 0 | 0 | 0 | 0 | 0 | 0 | 0 | 0 | 0 | 0 | 0 | 0 | 0 | 0 | 0 | 0 | 0 | 0 |
| 84 | OPR3 | AT2G06050 | JA | Oxidored_FMN | NO | 6 | 21 | 7 | 8 | 3 | 2 | 4 | 0 | 1 | 0 | 1 | 6 | 3 | 2 | 1 | 1 | 2 | 0 | 0 | 0 | 0 | 2 | 2 | 1 | 0 |
| 85 | JAR1 | AT2G46370 | JA | GH3 | YES | 2 | 6 | 11 | 1 | 0 | 0 | 0 | 0 | 0 | 0 | 0 | 0 | 0 | 0 | 0 | 0 | 0 | 0 | 0 | 0 | 0 | 0 | 0 | 0 | 0 |
| 86 | MYC2 | AT1G32640 | JA | bHLH-MYC_N, HLH | NO | 9 | 3 | 0 | 5 | 1 | 0 | 0 | 0 | 0 | 0 | 0 | 0 | 0 | 0 | 0 | 0 | 0 | 0 | 0 | 0 | 0 | 0 | 0 | 0 | 0 |
| 87 | NINJA | AT4G28910 | JA | EAR, TDBD | NO | 1 | 2 | 0 | 0 | 0 | 0 | 0 | 0 | 0 | 0 | 0 | 0 | 0 | 0 | 0 | 0 | 0 | 0 | 0 | 0 | 0 | 0 | 0 | 0 | 0 |
| 88 | JAZ | AT1G19180 | JA | Jas_motif, tify | NO | 4 | 2 | 0 | 0 | 0 | 0 | 0 | 0 | 0 | 0 | 0 | 0 | 0 | 0 | 0 | 0 | 0 | 0 | 0 | 0 | 0 | 0 | 0 | 0 | 0 |
| 89 | COI1 | AT2G39940 | JA | F-box | YES | 3 | 10 | 6 | 8 | 1 | 0 | 0 | 0 | 0 | 0 | 0 | 0 | 0 | 0 | 0 | 0 | 0 | 0 | 0 | 0 | 0 | 0 | 0 | 0 | 0 |
| 90 | ABCG16/JAT1 | AT3G55090 | JA | Not checked | YES | 13 | 11 | 3 | 12 | 9 | 2 | 12 | 4 | 4 | 1 | 1 | 0 | 0 | 0 | 0 | 0 | 0 | 0 | 0 | 0 | 0 | 0 | 0 | 0 | 0 |
| 91 | ABCD1/CTS/PXA1 | AT4G39850 | JA | AAA_21, AAA_29, ABC_membrane_2, ABC_tran | NO | 1 | 2 | 1 | 2 | 1 | 1 | 2 | 0 | 0 | 0 | 0 | 0 | 0 | 0 | 0 | 0 | 0 | 0 | 0 | 0 | 0 | 0 | 0 | 0 | 0 |
| 92 | NPF2/3 | AT3G47960, AT1G68570 | JA, GA | PTR2 | YES | 17 | 23 | 6 | 2 | 3 | 0 | 0 | 0 | 0 | 0 | 0 | 0 | 0 | 0 | 0 | 0 | 0 | 0 | 0 | 0 | 0 | 0 | 0 | 0 | 0 |
| 93 | ICS1 | AT1G74710 | SA | Chorismate_bind | NO | 2 | 2 | 1 | 1 | 1 | 0 | 0 | 0 | 1 | 0 | 1 | 0 | 1 | 0 | 1 | 1 | 0 | 1 | 0 | 1 | 0 | 0 | 0 | 0 | 0 |
| 94 | EDS5 | AT4G39030 | SA | MatE, Polysacc_synt_C | NO | 2 | 1 | 5 | 3 | 1 | 1 | 3 | 1 | 1 | 0 | 2 | 0 | 1 | 0 | 0 | 0 | 0 | 1 | 0 | 0 | 0 | 0 | 0 | 0 | 0 |
| 95 | NPR1/3/4 | AT1G64280, AT5G45110 | SA | NPR1_like_C | NO | 6 | 6 | 3 | 5 | 1 | 0 | 0 | 0 | 0 | 0 | 0 | 0 | 0 | 0 | 0 | 0 | 0 | 0 | 0 | 0 | 0 | 0 | 0 | 0 | 0 |
| 96 | TGA | AT5G65210 | SA | bZIP_1, bZIP_2, DOG1 | NO | 10 | 68 | 3 | 4 | 1 | 1 | 3 | 4 | 2 | 0 | 2 | 0 | 0 | 0 | 0 | 0 | 0 | 0 | 0 | 0 | 0 | 0 | 0 | 0 | 0 |
| 97 | WRKYs | AT4G26640 | SA | WRKY | NO | 6 | 19 | 0 | 3 | 1 | 1 | 1 | 0 | 0 | 0 | 0 | 0 | 0 | 0 | 0 | 0 | 0 | 0 | 0 | 0 | 0 | 0 | 0 | 0 | 0 |
| 98 | D27 | AT1G03055 | SL | D27-like_C | NO | 2 | 4 | 3 | 4 | 3 | 2 | 8 | 1 | 3 | 2 | 3 | 1 | 4 | 1 | 2 | 0 | 3 | 1 | 2 | 1 | 0 | 0 | 0 | 1 | 0 |
| 99 | CCD7(MAX3) | AT2G44990 | SL | RPE65 | NO | 1 | 1 | 1 | 1 | 1 | 0 | 0 | 0 | 0 | 0 | 0 | 0 | 0 | 0 | 0 | 0 | 0 | 0 | 0 | 0 | 0 | 0 | 0 | 0 | 0 |
| 100 | CCD8(MAX4) | AT4G32810 | SL | RPE65 | NO | 1 | 3 | 1 | 1 | 0 | 0 | 0 | 0 | 0 | 1 | 0 | 0 | 1 | 2 | 1 | 0 | 1 | 0 | 0 | 0 | 0 | 0 | 0 | 0 | 0 |
| 101 | MAX1 | AT2G26170 | SL | p450 | NO | 1 | 9 | 1 | 0 | 0 | 1 | 3 | 0 | 4 | 4 | 3 | 0 | 0 | 0 | 0 | 0 | 0 | 0 | 0 | 0 | 0 | 0 | 0 | 0 | 0 |
| 102 | D14/D14L/KAI2 | AT3G03990, AT4G37470 | SL | Abhydrolase_1, Abhydrolase_6, Hydrolase_4 | NO | 3 | 6 | 2 | 12 | 2 | 0 | 3 | 1 | 1 | 2 | 0 | 0 | 0 | 0 | 0 | 0 | 0 | 0 | 0 | 0 | 0 | 0 | 0 | 0 | 0 |
| 103 | MAX2 | AT2G42620 | SL | F-box | NO | 1 | 2 | 1 | 1 | 1 | 0 | 2 | 0 | 0 | 0 | 0 | 0 | 0 | 0 | 0 | 0 | 0 | 0 | 0 | 0 | 0 | 0 | 0 | 0 | 0 |
| 104 | SMAX1 | AT5G57710 | SL | AAA_2 | NO | 1 | 0 | 0 | 1 | 1 | 0 | 0 | 0 | 0 | 0 | 0 | 0 | 0 | 0 | 0 | 0 | 0 | 0 | 0 | 0 | 0 | 0 | 0 | 0 | 0 |
